# Accelerated evolution of whole gene clusters by an engineered lytic phage system in *E. coli*

**DOI:** 10.1101/2025.04.21.649713

**Authors:** Shujian Ong, Pramila Ghode, Ashvinath Narenderan, Fabian Willenborg, Tobias V. Eden, Shuxuan Lao, Carl O. Marsh, Wen Shan Yew, Julius Fredens

## Abstract

Directed evolution excels at optimizing individual proteins, but simultaneous evolution of multiple genes remains a significant challenge. Here, we establish a robust phage-based system for accelerated evolution of large gene clusters of up to 39 kilobases. Lytic Selection and Evolution (LySE) selectively replicates and mutagenizes the target gene cluster, carried on a phagemid through alternating cycles of lysis and transduction in *Escherichia coli*. We engineered a hypermutagenic T7 DNA polymerase (T7 DNAP) that increases mutation rates of the replicating phagemid 7,000-fold during the lytic cycle. We further optimized the mutational spectrum by fusing the T7 DNAP to a dual adenine-cytosine deaminase to install all possible transition mutations at similar frequencies. LySE-mediated transduction enables selection for desired metabolic functions by coupling gene cluster expression to host fitness, while eliminating genomic off-target mutations by refreshing the host in each cycle. Using LySE, we evolved a 25-fold increase in tigecycline resistance in 5 cycles, and a 50.9% increase in end-point biomass of a bacterial strain that utilizes the PET monomer, ethylene glycol, as its sole carbon source.

## Introduction

Directed evolution enables discovery of novel protein variants and is an indispensable tool for bioengineering^1^. Classical directed evolution platforms based on display technologies and *in vitro* mutant library generation are time-consuming to iterate, hard to scale up, and limited in their ability to explore the full sequence space of a gene of interest (GOI)^2, 3^. To overcome these limitations, continuous evolution systems have been developed to selectively mutagenize target GOIs inside cells, enabling evolutionary cycles to run without manually initiated steps^4^. Most prominently, phage-assisted continuous evolution (PACE) has been applied for successful evolution of various targets^5–7^. PACE utilizes the M13 bacteriophage as a gene delivery vector, enabling seamless transition between mutagenesis and selection of GOIs entirely *in vivo*, thereby greatly expanding the scale and depth of the evolutionary search. Despite its utility, PACE is constrained by the M13 phage system: The small capsid, together with the need to maintain essential phage genes, restricts GOI size to 5,000 base pairs during evolution cycles^8^. This limits the applicability of PACE to the evolution of ‘simple’ phenotypes encoded by only one or a few GOIs.

However, most phenotypes in living organisms emerge from interactions between multiple genes^9^, including important cellular processes like small molecule synthesis^10^, carbon assimilation^11^, protein folding and complex assembly^12^, and bacterial secretion systems^13^. Evolution of such complex processes depends on methods with capacity to evolve multiple genes simultaneously. Furthermore, these complex phenotypes are often slow manifesting, resulting in high fitness cost and reduced phage titers if they were to be evolved through PACE^14^. Recently, several systems for orthogonal replication have emerged^15–18^, offering expanded capacity and more direct coupling between GOI function and cellular metabolism. However, adaptive evolution with orthogonal replication systems risks accumulating two classes of undesirable mutations: (i) genomic off-target mutations that make non-GOI genes contribute to the phenotype, which complicates the attribution of phenotypic effects to specific genetic changes^19^, and (ii) ‘cheater mutations’ that allow the evolution system to circumvent selection pressure altogether, especially in biosensor-based selection^20^.

To address both the size limitation of PACE and the risk of off-target mutations in orthogonal replication systems, we developed Lytic Selection and Evolution (LySE), a robust T7 phage-based system for accelerated evolution of whole Gene Clusters Of Interest (GCOI) up to 40 kilobases. LySE employs a T7 phagemid that is maintained as a stable plasmid during the cell cycle but packaged into the phage capsid upon infection^21^. The system selectively replicates and mutagenizes the GCOI phagemid and GCOI while carrying it through alternating cycles of lysis and transduction in *Escherichia coli*. Each LySE cycle completely eliminates the *E. coli* culture after selection, effectively removing all off-target mutations. We engineered a hypermutagenic T7 DNA polymerase (T7 DNAP), that achieves a 7,000-fold increase in phagemid mutation rate during the lytic cycle. The mutational spectrum was further optimized by fusing the T7 DNAP with a dual adenine-cytosine deaminase to assure an even incorporation of all transition (purine-to-purine and pyrimidine-to-pyrimidine) mutations.

LySE-mediated transduction enables selection of slow-manifesting metabolic functions by coupling large gene cluster expression to host fitness. We demonstrate this by evolving a pathway that allows the host bacterium to utilize and grow on the polyethylene terephthalate (PET) monomer, ethylene glycol (EG), as its sole carbon source. LySE uniquely combines continuous evolution with discrete evolution cycles, enabling directed evolution of large gene clusters while maintaining stringent control over mutational trajectories.

## Results

### Evolution through a lytic cycle

Bacteriophage T7 is a lytic phage that infects *E. coli*, replicates its 40 kb genome, and lyses the host cell within 17 minutes, releasing approximately 180 progeny phages (**Extended Data** Fig. 1a)^22^. We exploited the ability of the T7 phage for rapid multiplication and distribution of large genetic material to develop a system for continuous, accelerated evolution of large Gene Clusters of Interest (GCOI). First, we engineered a T7 phage variant lacking T7 DNAP, by *in vitro* assembly of the complete genome except *gp5*, and rebooting the phage in a cell-free extract from *E. coli* (**Supplementary Data 1**)^23^. This phage, T7ΔDNAP, efficiently propagates only in hosts that carry an accessory plasmid (AP) expressing the T7 DNAP under a T7 promoter (**Extended Data** Fig. 1c,d). The absence of phage propagation without accessory plasmid demonstrates the strict biocontainment of the system.

Phage T7 can replicate, package, and transduce phagemids (PM); circular plasmids containing the T7 origin of replication, a T7 terminal repeat, and a host origin of choice^24^. Importantly, the phagemid is replicated by the host replication machinery during cell cycle, and only by the phage replication machinery during phage infection. We constructed a phagemid with a p15a host origin and demonstrated that it was efficiently packaged and transduced by phage T7ΔDNAP in cells containing accessory plasmid expressing wildtype or error-prone T7 DNAP (**Fig. 1a, Extended Data** Fig. 1e, **Supplementary Data 2**, **Supplementary Table 1**).

**Fig. 1.**
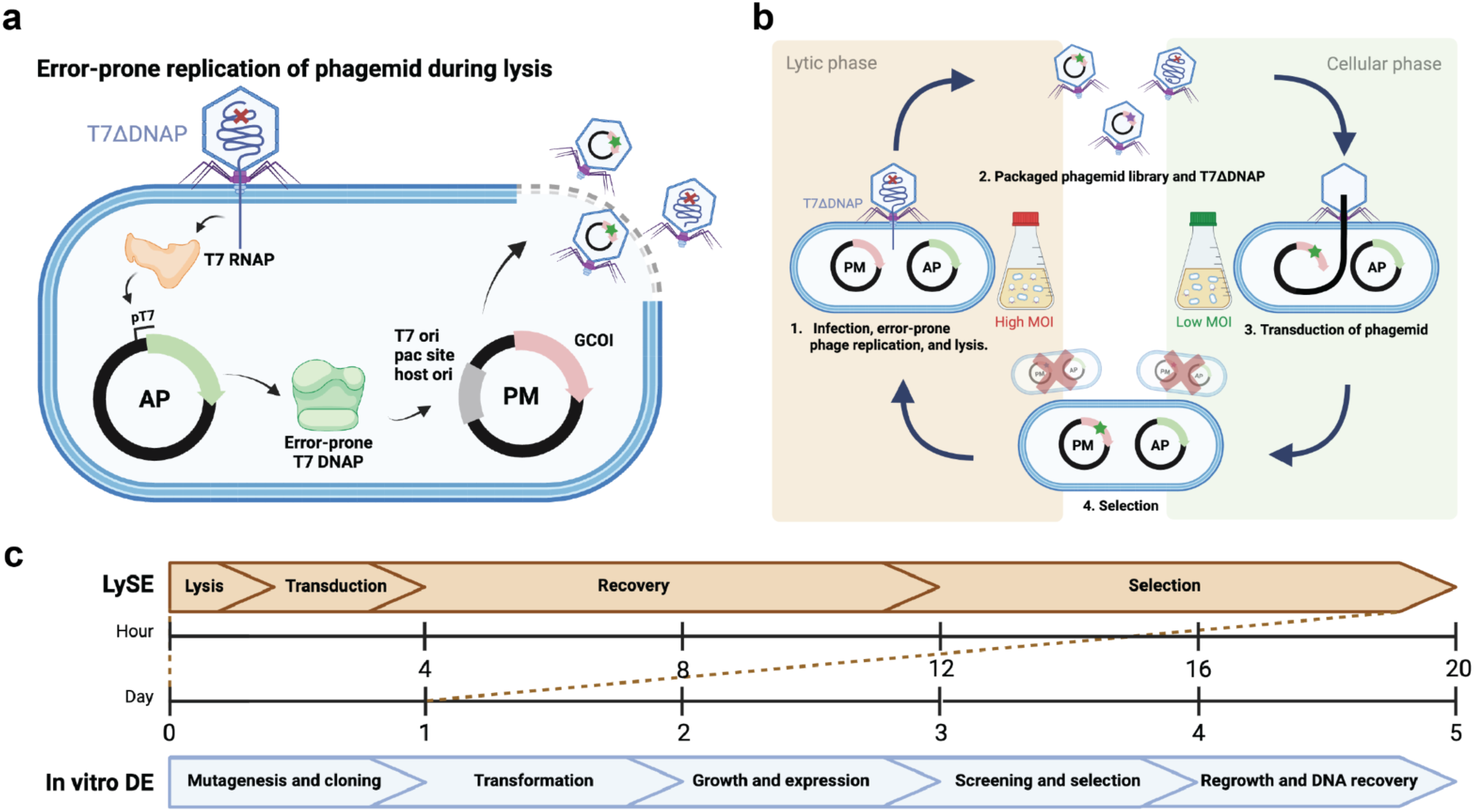
LySE evolves gene clusters through cycles of lysis and transduction. **a.** Controlled replication and phage packaging. T7 DNAP expression from the accessory plasmid (AP) is tightly regulated under a T7 promoter during the cell cycle. Upon T7 phage infection, the T7 RNA polymerase (T7 RNAP) induces expression of error-prone T7 DNAP, which replicates the phagemid (PM) for subsequent packaging and transduction. **b.** Schematic of the LySE workflow. At high multiplicity of infection (MOI), the system enters a lytic phase where the gene cluster of interest (GCOI), carried on a phagemid, undergoes error-prone replication by error-prone T7 DNAP. Mutated variants are packaged into T7 phage particles and released. During subsequent transduction at low MOI, these mutated phagemids are introduced into new host cells, enabling fitness-based selection through cell growth. The evolved gene cluster pool is then cycled back to the lytic phase by reintroducing phage T7ΔDNAP. **c.** Temporal comparison of LySE versus conventional directed evolution methods, demonstrating acceleration from week-long to day-scale evolution cycles through exploitation of the short life cycle of T7 phage.

When equipped with an error-prone T7 DNAP variant, this system will enable cyclic evolution, where the phagemid undergoes error-prone replication and packaging during a lytic phase (high Multiplicity Of Infection, MOI), followed by transduction to fresh hosts for growth and selection during a cellular phase (low MOI) (**Fig. 1b**). During the cellular phase, T7 DNAP expression will cease and host DNAPIII maintains the phagemid with high fidelity. The selection of improved GCOI variants is achieved by linking phagemid-encoded functions to host cell fitness during the cellular phase. The short lysis time and large burst size of T7 facilitate quick turnover of large genetic pools, thus reducing the time for directed evolution from days to hours (**Fig. 1c**).

### Engineering hypermutagenic T7 DNA polymerases

We engineered and tested several error-prone T7 DNAP variants to drive targeted mutagenesis of phagemid-encoded GCOIs (see **Supplementary Table 2** for all variants). Our design strategy incorporated complementary error-inducing mechanisms for additive effects. We began with a previously characterized error-prone variant, T7 DNAP v1, containing mutations in both the thumb domain (S399T) and exonuclease domain (Y64C/F120L)^25^. S399T has been proposed to increase error rates by disturbing the nucleic acid binding cleft, while Y64C and F120L likely reduce proof-reading activity. This variant exhibited a 13.5-fold increased error rate compared to wild type T7 DNAP (**Fig. 2a**), in line with previous reports.

**Fig. 2.**
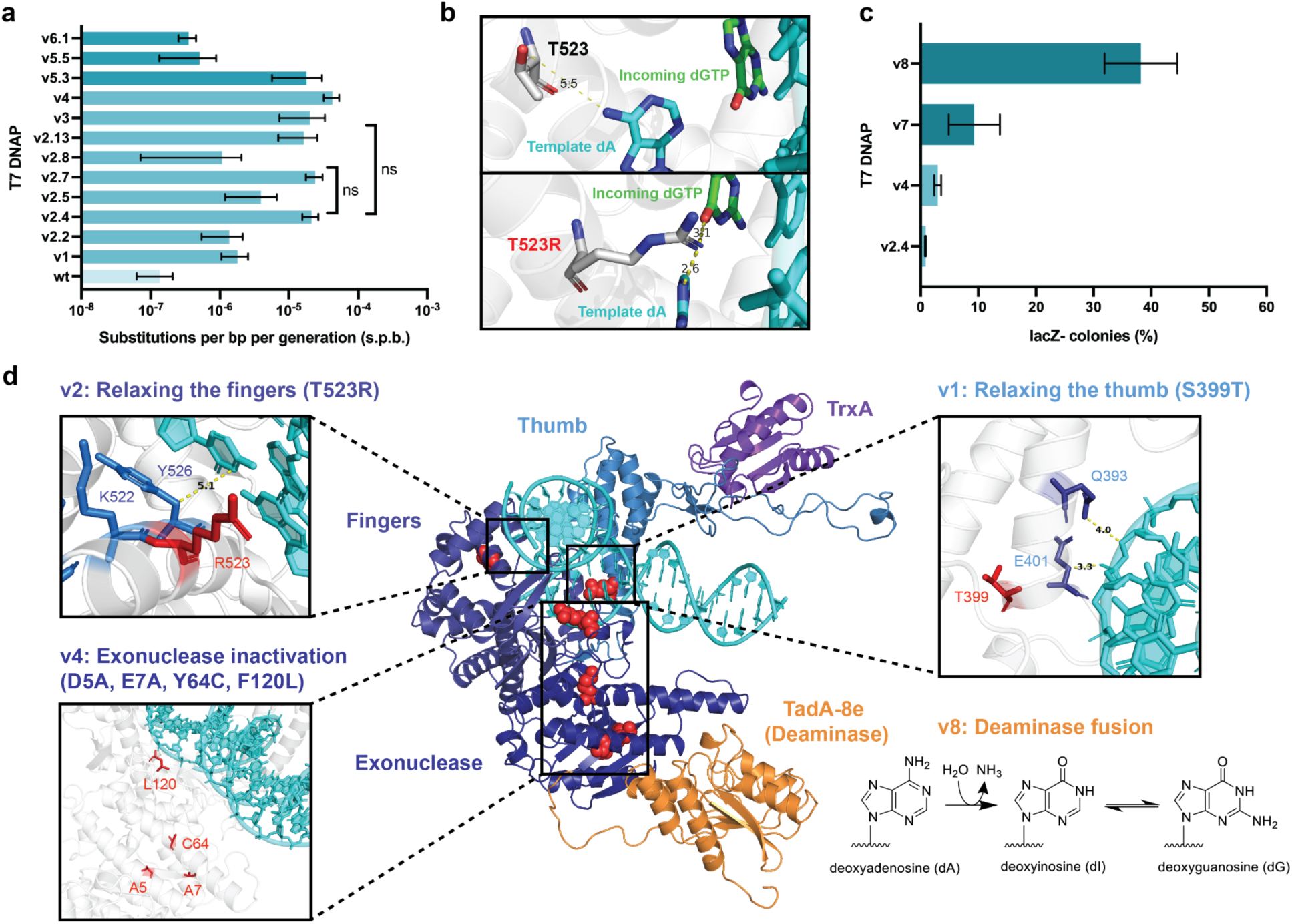
Engineering and characterization of hypermutagenic T7 DNA polymerase variants **a.** Mutational frequencies of engineered T7 DNAP variants measured by a fluctuation assay using a phagemid-encoded chloramphenicol resistance gene containing an internal stop codon. Wild type T7 DNAP (wt), mutant T7 DNAPs (v1-v4), and wild type T7 DNAP-deaminase fusions (v5-v6). Data represent mean ± SD from three independent experiments. **b.** Snapshots from molecular dynamics simulations of T7 DNAP (PDB: 1T7P): wildtype (top) and T523R variant mutated *in silico* (bottom) with A-G nucleotide mismatch. Hydrogens not displayed. Distances measured in Ångström. **c.** Mutational frequencies of advanced T7 DNAP variants determined through a *lacZ* inactivation assay using a phagemid-encoded *lacZ* reporter. Mutations were quantified by screening for white or light blue colonies indicative of loss-of-function mutations. Mutant T7 DNAPs (v2-v4), mutant T7 DNAP-deaminase fusions (v7-v8). Data represent mean ± SD from three independent experiments. **d.** AlphaFold3-predicted structure of variant v8, highlighting key engineered modifications: v1 (S399T) - thumb domain relaxation affecting template minor groove interactions; v2 (T523R) - fingers domain modification proximal to active site; v4 (D5A, E7A, Y64C, F120L) - exonuclease domain inactivation; v8 - fusion with TadA-8e adenosine deaminase enabling concurrent DNA deamination during replication.

Mutation rates were quantified by fluctuation analysis using a chloramphenicol resistance gene (*CmR*) containing a premature stop codon (Q38TAG)^26, 27^. After one generation of phagemid replication and transduction, we measured the frequency of cells acquiring chloramphenicol resistance through point mutations converting the TAG to sense codons. We calculated apparent mutation rates in substitutions per base pair per generation (s.p.b) by correcting for multiple phagemid copies per phage particle.

To further enhance mutagenesis, we employed homology-guided design based on fidelity-reduced *E. coli* DNAP I variants^28^. We identified several potential mutations in the fingers domain of T7 DNAP (L479N/H506Y/T523R/P560H), of which some are positioned near the polymerase active site. Through systematic experimental testing, we determined T523R to significantly increase error rates, and when combined with v1 (v2.4: Y64C/F120L/S399T/T523R) achieved error rates 157-fold higher than wild type (wt). Molecular dynamics simulations of the T7 DNAP crystal structure^29^ revealed that T523R significantly alters nucleotide positioning in the active site (**Fig 2b**, **Extended Data** Fig 2, **Supplementary Data 3**). An incoming triphosphate nucleotide is pulled deeper into the active site in presence of the arginine substitution. Thus, we propose that the arginine stabilizes incorrect base-pairing.

Complete inactivation of the exonuclease function through D5A/E7A mutations further reduced replication fidelity^30^. Incorporating D5A/E7A into T7 DNAP v2.4 generated variant v4 with a 2-fold increased error rate (**Fig 2a**). Despite successfully engineering multiple error-prone variants, we observed a ceiling at 4.27 × 10^-5 s.p.b in the fluctuation assay, whereby increased mutation rates were accompanied by declining phagemid transduction efficiency (**Extended Data** Fig 3a). We hypothesized that higher mutation rates might be obscured if inactivating mutations elsewhere in the *CmR* gene counteracted TAG reversion effects.

We therefore employed a *lacZ*-inactivation assay capable of detecting higher mutation frequencies (**Fig. 2c**)^31^. We encoded *lacZ* on the phagemid, yielding blue colonies in presence of X-Gal. At high error rates, *lacZ* disruption produces white or light blue colonies. Using variants v2.4 and v4 as benchmarks, we observed 0.86% and 2.96% *lacZ* inactivation rates respectively—a 3.4-fold difference consistent with the fluctuation assay results.

In parallel we explored error-prone replication by fusing T7 DNAP to DNA deaminases for concurrent deamination during replication. We fused the adenosine deaminase TadA-8e to the N-terminus of wild type T7 DNAP with linkers of varying lengths^32^ (v5.1-5.5, **Supplementary Table 2b**). A 24-residue linker (v5.3) resulted in the highest error rate of 1.79 × 10^-5^, comparable to the rate of v2.4 (**Fig 2a**). The cytosine deaminase PmCDA1 fusion (v6.1) proved less effective than TadA-8e.

Finally, we combined the two complementary approaches by fusing TadA-8e to the most error-prone T7 DNAP variants with mutations in the thumb, fingers and exonuclease domain (v2.4) as well as exonuclease inactivation (v6), creating v7 and v8, respectively. These fusion variants resulted in 10-fold and 40-fold increased lacZ inactivation compared to the mutant variants (**Fig. 2c**). T7 DNAP v8 achieved error rates of 9.40 × 10^-4^ s.p.b—nearly one substitution per one thousand base pairs. Variant v8 had negligible impact on host growth kinetics (**Extended Data** Fig 3b), demonstrating tight control of error-prone replication. The final engineered T7 DNAP v8 comprises a hypermutagenic T7 DNAP suitable for mutagenizing and diversifying phagemid libraries for continuous evolution with LySE (**Fig 2d**).

### Multiplicity tuning for controlled lysis

Next, we assessed how our error-prone T7 DNAP variants affect T7 phage replication. *E. coli* cultures harboring T7 DNAP variants were infected with T7ΔDNAP at varying MOIs while we monitored cell density over time (**Fig. 3a**). Wild type T7 DNAP caused efficient cell lysis independent of the initial MOI, due to rapid phage propagation^22^. However, as the error rate of replication increased (v2.4-v8), the resulting lysis became highly MOI-dependent. In cultures with T7 DNAP v8, the cell density increased during infection at low MOIs, while it decreased at high MOI (**Fig. 3b**). These findings demonstrate that lysis by T7 phage—a strictly lytic phage in nature—can be tuned by multiplicity of infection under error-prone replication.

**Fig. 3.**
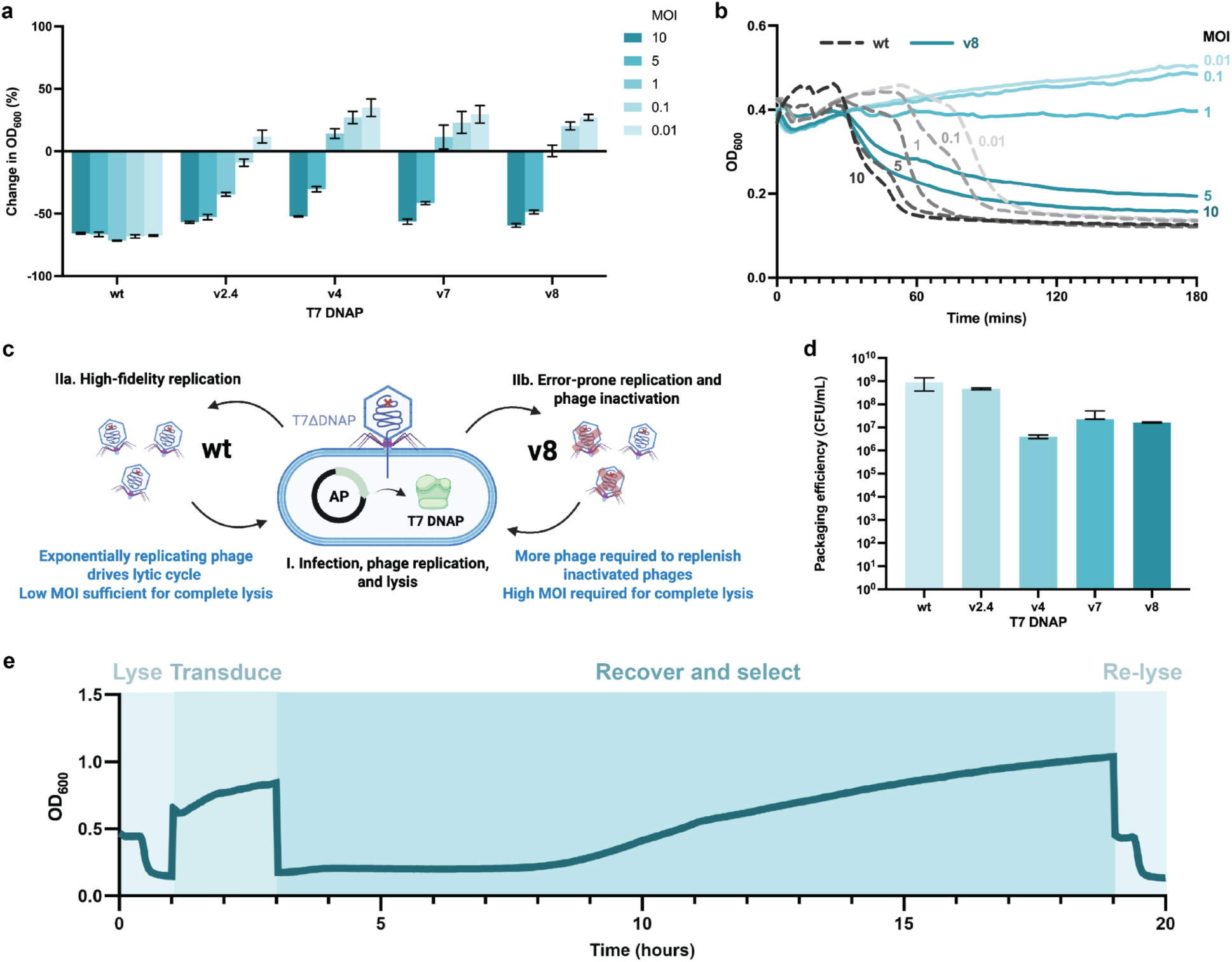
Continuous lytic cycling of phagemids by multiplicity tuning **a.** Change in optical density of *E. coli* cultures expressing T7 DNAP wild type (wt) and variants (v2.4-v8) when infected with T7ΔDNAP over 3 hours at varying multiplicities of infection (MOI: 0.01-10). Error bars represent mean ± SD (n = 3). **b.** Lysis kinetics of *E. coli* cultures expressing wild type (wt, dashed lines) or hypermutagenic (v8, solid lines) T7 DNAP at different MOIs (0.01-10). **c.** Schematic representation of proposed multiplicity tuning mechanism: After infection (step I), wild type T7 DNAP enables efficient phage replication at low MOI (step IIa), while error-prone variant v8 requires higher MOI due to increased phage inactivation during replication (step IIb). **d.** Quantification of phagemid packaging efficiency across T7 DNAP variants, demonstrating maintained library diversity despite reduced transduction rates in error-prone variants. Data shown as mean ± SD (n = 3). **e**. Representative 20-hour time course of a complete LySE cycle, showing distinct phases of bacterial growth and phage-mediated lysis.

During faithful replication with wild type T7 DNAP, the phage propagates and lyses the cells exponentially, independent of MOIs (**Fig. 3c**). Conversely, error-prone variants like v8 compromise replication fidelity, resulting in unsustainable propagation and MOI-dependent lysis^25^. We verified that this compromised propagation still permits substantial phagemid packaging and transduction even with highly error-prone DNAP v8 (**Fig. 3d**).

The ability to tune the degree of culture lysis by adjusting T7 phage multiplicity enables controlled and distinct phases of the LySE cycle. At high MOI, mutated phagemids are packaged into virions and released through efficient cell lysis (**Fig. 3e**). Upon lowering the MOI by infecting a fresh culture, phagemid libraries get transduced to fresh cells without widespread lysis. This library of transductants can be grown under selective conditions to enrich for improved GCOI performance before initiating another round of lysis. These distinct LySE phases allow adjustment of growth and selection duration, facilitating evolution of slow-manifesting phenotypes such as low-rate metabolic pathways^33^ or slow-folding protein complexes^34^.

### Broadening the mutagenesis spectrum

Having established a highly error-prone T7 DNAP variant and methods to control the LySE cycle, we next sought to broaden its mutagenesis spectrum to introduce all transition mutations at comparable frequencies. Variant v8 contains TadA-8e^32^, which catalyzes adenosine-to-inosine deamination, resulting in A:T→G:C transitions during DNA replication (**Fig. 4a**). Illumina next-generation sequencing (NGS) confirmed that transition mutation frequencies were skewed toward A:T→G:C transitions at a ratio of 1:0.8 (**Fig. 4b**). To address this bias, we installed a recently engineered dual adenine-cytosine deaminase, TadDE (v9), capable of performing both adenosine and cytidine deamination^7^, facilitating A:T→G:C and C:G→T:A transitions, respectively (**Fig. 4a**). This modification improved the transition ratio to 1:0.92, demonstrating that v9 introduces all transition mutations at nearly equivalent frequencies (**Fig. 4b**) while maintaining a high overall error rate (**Extended Data** Fig 4a).

**Fig. 4.**
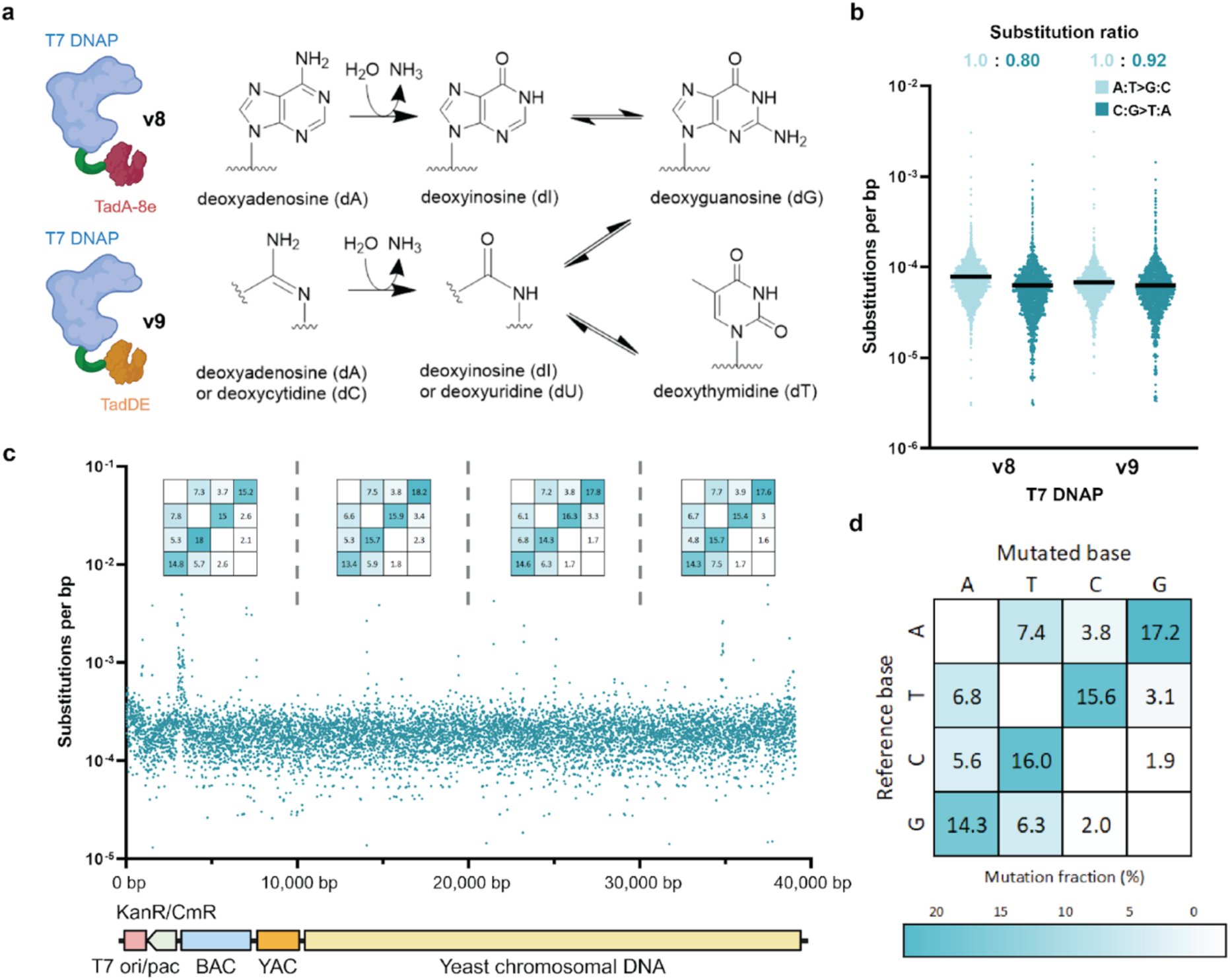
A hybrid deaminase enables a broad mutagenesis spectrum **a.** Biochemical mechanism of nucleobase modifications by deaminases. Fusion of error-prone T7 DNAP with the adenosine deaminase TadA-8e (v8) catalyzes adenosine-to-inosine deamination resulting in A:T→G:C transitions. Installation of the dual adenine-cytosine deaminase TadDE (v9) enables both adenosine and cytidine deamination, facilitating A:T→G:C and C:G→T:A transitions. **b.** Substitution frequencies and ratios of A:T→G:C and C:G→T:A transitions for v8 and v9 variants derived from NGS data (Illumina sequencing, mean coverage >14,000x per base pair). **c.** Distribution of substitution frequencies across a 39-kb BAC-phagemid (mean coverage = 14,098x per base pair). Mutational spectra were calculated for every 10,000 bp. **d.** Mutational spectra of LySE v9.

To probe the maximum capacity of LySE to evolve large GCOIs, we assembled a 39 kb T7 phagemid with a bacterial artificial chromosome (BAC) origin by homologous recombination in *S. cerevisiae* (**Supplementary Data 4**). After performing one generation of LySE with this BAC-phagemid, nanopore sequencing of transductants confirmed that the intact vector was successfully transferred into fresh host cells (**Extended Data** Fig 4b). NGS analysis revealed a uniform distribution of mutations and base transitions throughout the entire 39 kb BAC-phagemid (**Fig. 4c**). LySE exhibited a broad mutagenesis spectrum with all transitions occurring at similar frequencies, while maintaining the biologically relevant preference for transitions over transversions^35^ (**Fig. 4d**). Taken together, LySE presents a significant improvement over existing continuous evolution tools by enabling the potential evolution of large gene clusters up to 40 kb, equivalent to the size of the complete T7 genome^21^.

### Accelerated evolution of genes and gene clusters

To demonstrate the utility of LySE for accelerated gene evolution, we first evolved the tetracycline resistance gene *tetA* to confer resistance to tigecycline. We expressed *tetA* from a phagemid and subjected it to 5 generations of LySE with progressively increasing tigecycline concentrations (**Fig. 5a**, **Supplementary Data 5**). For comparison, we performed adaptive laboratory evolution (ALE) under identical selection conditions. Following LySE evolution, we obtained cells tolerating 2.5 μg/mL of tigecycline—a 25-fold increase over the wild type tolerance of 0.1 μg/mL (E5 LySE, **Fig. 5b**). In contrast, cells evolved by ALE tolerated only up to 1 μg/mL (E5 ALE). Furthermore, when the ALE-evolved phagemid was transferred to fresh cells, the acquired resistance was lost (E5T ALE, **Fig. 5b**). This indicates that the evolved resistance was caused by genomic mutations rather than mutations in the target phagemid, and illustrates that ALE is highly sensitive to off-target and cheater mutations^19^. LySE overcomes these issues by refreshing the host in each cycle, thus only allowing evolution through on-target mutations of the phagemid.

**Fig. 5.**
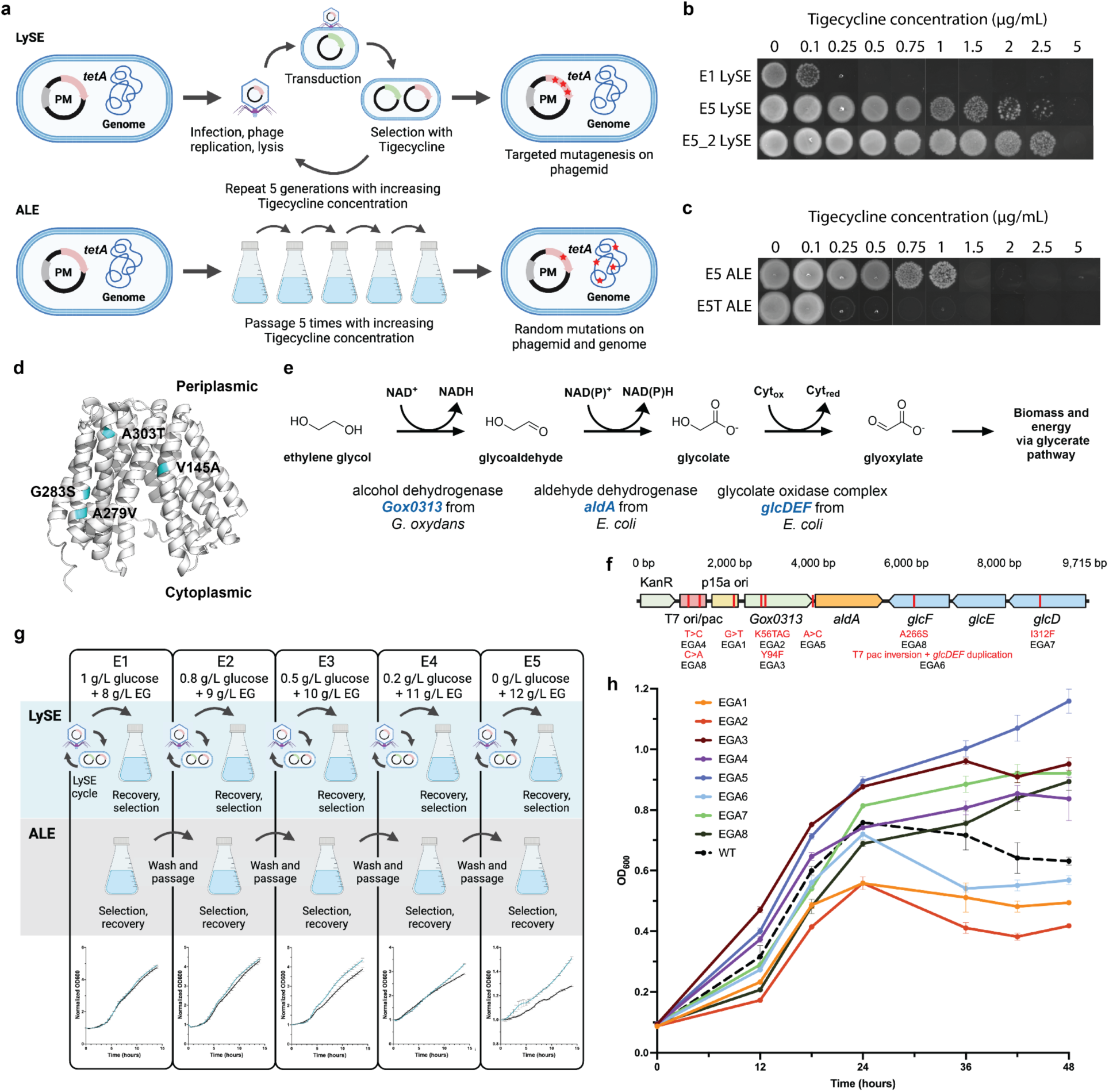
Phagemid-targeted evolution of genes and gene clusters **a.** Comparative workflows of LySE versus adaptive laboratory evolution (ALE). LySE confines mutations to the phagemid through phage-mediated cycling, while ALE allows accumulation of genome-wide mutations during serial passaging. **b.** Tigecycline resistance profiles of an evolved pool after five generations of LySE (E5 LySE), compared to the culture before evolution (E1 LySE). A representative single clone from E5 (E5_2 LySE) is also shown. **c.** The same phagemid and *tetA* evolved by ALE. Resistance profile of pool after five passages (E5 ALE). The acquired resistance was almost entirely lost following when transferring the phagemid to a fresh host cell (E5T ALE), indicating genome-dependent rather than phagemid-encoded adaptation. **d.** Structure of TetA and identified mutations in the evolved pool by Sanger sequencing of 32 clones. **e.** Metabolic pathway for ethylene glycol (EG) assimilation in *E. coli*. **f.** Linearized representation of the phagemid with the EG assimilation pathway for evolution. Identified mutations from 8 clones are labelled in red, with corresponding clone number (EGA1-8). **g.** Comparative workflows for semi-relaxing evolution of EG assimilation pathway using LySE versus ALE. Both approaches employed progressive selection from 1 g/L glucose + 8 g/L EG (E1) to 0 g/L glucose + 12 g/L EG (E5). The LySE protocol included cell recovery in ampicillin and kanamycin to maintain phagemid and accessory plasmid, followed by washing and selection in minimal medium. Similarly, the ALE protocol comprised initial growth, washing, minimal medium selection, antibiotic recovery to maintain phagemid and accessory plasmid, and final washing before the next selection cycle. Growth curves for each selection cycle are shown. **h.** Growth curves of eight isolated evolved phagemids (EGA1-8), and wild type phagemid (WT) in M9 media with 10 g/L EG and no glucose. For panel g and h, data are the mean ± standard deviation from three (n = 3) independent biological replicates.

From evolved clones of E5 LySE, we identified four point mutations in *tetA* (**Fig. 5d**, **Extended Data** Fig. 5a,b), and a mutation in the promoter that increased *tetA* expression approximately 200-fold (**Extended Data** Fig. 5c). This demonstrates the ability of LySE to simultaneously evolve both regulatory and coding regions, which is particularly useful for optimizing complex phenotypes^36^.

We next explored the capability to evolve a complex process—a multigene cluster constituting a complete metabolic pathway for assimilation of ethylene glycol (EG), a monomer derived from PET degradation. This pathway was chosen due to its significant implications for plastic recycling and its potential contribution towards developing a sustainable circular economy for plastic waste management^33, 37^.

We constructed the pathway starting with EG reduction to glycolaldehyde using *Gluconobacter oxydans* alcohol dehydrogenase (*Gox0313*), selected for its superior performance and usage of only NAD+ as cofactor^38^ (**Fig. 5e**). Glycolaldehyde is further reduced to glycolate and then to glyoxlylate by endogenous aldehyde dehydrogenase (*aldA*) and glycolate oxidase complex (*glcDEF*) from *E. coli*. The resulting glyoxylate enters the glycerate pathway for biomass and energy production^39^ (**Fig. 5e**). We cloned all five genes into a T7 phagemid, creating a 9,715 bp plasmid (**Fig. 5f**, **Supplementary Data 6**), and subjected the pathway to 5 generations of evolution by LySE or ALE under the same selection regime. We implemented a semi-relaxing selection protocol with gradual glucose withdrawal from the culture medium, resulting in improved normalized growth rates for LySE-evolved cells compared to ALE-evolved cells (**Fig. 5g, Extended Data** Fig. 6). Sequencing of clones evolved by LySE, revealed mutations in the T7 ori/pac site, the p15a ori, near the *aldA* promoter, and nonsynonymous mutations in three of the five genes (**Fig. 5f**, **Supplementary Data 7**). Five clones exhibited accelerated growth on EG compared to wild type, featuring a novel Y94F substitution in Gox0313 (EGA3), and I312F substitution in *glcD* (EGA7), with improved endpoint biomass by 50.9% and 46.1% respectively after just 5 generations (**Fig. 5h**, **Extended Data** Fig. 7). These experiments demonstrate LySE as a powerful directed evolution platform that enables rapid, targeted genetic optimization across both single genes and complex gene clusters.

## Discussion

Here, we have developed LySE, a robust T7 phage-based system enabling the accelerated evolution of gene clusters up to 40 kb in size. Unlike previous phage-based continuous evolution methods such as PACE^5^ and T7AE^40^ that insert GOIs directly into the phage genome — thereby limiting the capacity to less than 5 kb^8^ — our approach employs a T7 phagemid design utilizing T7 origin and packaging signals to direct the phage to package the GCOI as-is into its capsid. This strategy leverages the full capacity of the T7 capsid to enable evolution of 40 kb constructs, of which 29 kb of GCOI was tested in this work (**Extended Data** Fig. 4b, **Supplementary Data 4**). This substantially exceeds the capabilities of existing phage-assisted and *in vivo* continuous evolution methods^5, 28, 40–42^. The expanded capacity of LySE significantly broadens potential applications for continuous evolution. It enables work with anabolic pathways for small molecule synthesis^10^, catabolic pathways for waste assimilation^33, 38, 39^ (**Fig. 5**) or carbon capture^11^, and evolution of protein complexes^12^ — many of which exceed 10 kb when including regulatory elements. This versatility eliminates size constraints that have previously limited directed evolution approaches.

While orthogonal replication systems have gained traction for handling larger constructs of up to a 16.5 kb replicon^16^, they remain vulnerable to off-target mutations that can accumulate as genomic hitchhikers, complicating phenotypic attribution^19^. These systems also risk enabling cheater mutations that bypass selection pressure, especially in biosensor-based evolution^20^. These vulnerabilities stem from the fundamental mechanism of orthogonal replication, where the GOI replicates with the host, allowing off-target mutations to be carried over to subsequent generations. Similarly, P1 phage-based methods like IDE, despite accommodating large inserts of up to 36 kb^43^, are known to transfer flanking genomic regions after integration—a property that, while useful for genome engineering^44^, presents challenges for precise directed evolution. LySE overcomes these limitations through its lytic cycle. T7 phage-mediated cell lysis completely eliminates the *E. coli* culture after each selection cycle, effectively removing all off-target mutations. There remains a risk of off-target mutations within the phagemid itself, such as alterations to the *E. coli* origin that could increase plasmid copy number instead of evolving the GCOI (**Fig. 5f**). This can be addressed through stringent selection pressures or biosensor-based approaches, particularly when evolving enzyme specificity for synthetic metabolism^45^.

Furthermore, as an orthogonal DNA polymerase, T7 DNAP can be engineered to achieve extremely high error rates without affecting cell viability^18^. Our implementation of controlled error-prone replication and multiplicity tuning redirects pressure away from maintaining T7 genome fidelity, enabling us to exceed standard T7 phage error thresholds for the first time^25^ (**Fig. 2a**), achieving an estimated 9.40 × 10^-4^ s.p.b – one of the highest error rates in continuous evolution^46^.

By leveraging T7 as an efficient gene shuttle between cells, LySE uniquely combines continuous evolution with controlled discrete cycles. Unlike other phage-assisted evolution methods, LySE employs intracellular selection rather than viral fitness coupling. This enables direct coupling of GCOI function to cellular fitness, more appropriate for applications ultimately deployed in cellular contexts, such as strain or therapeutic cell engineering^47^, while also enabling selections based on physical cell properties through techniques like cell sorting^48^.

While manually performed in this study, the LySE workflow (lysis-transduction-recovery-selection) is straightforward and amenable to automation via liquid handling systems for increased throughput^49^. Our results establish a generalizable method for accelerated evolution of whole gene clusters, providing researchers with a robust tool for engineering large metabolic pathways and protein complexes without the complications of off-target mutations or mutations that circumvent selection.

## Acknowledgements

We thank L.F.H. Funke for helpful comments on the manuscript, and B. Xue for helpful discussions on molecular dynamics simulations. This work was supported by the National University of Singapore Presidential Young Professorship (NUHSRO/2021/071/Startup/06) and the National Research Foundation Fellowship (NRFF14-2022-0096) all to J.F., while S.O. was supported by the National University of Singapore President’s Graduate Fellowship, Integrative Science and Engineering Programme. Illustrations were created in BioRender.com under full license to publish.

## Author contributions

S.O. and J.F. conceived and designed the project. P.B.G. established T7 phagemid packaging and transduction. S.O., J.F. and F.W. designed the error-prone T7 DNAP variants. S.O., P.B.G., A., F.W. and S.L. performed cloning and strain engineering. S.O. performed experimental characterizations and NGS analysis. T.V.E. and C.M. performed molecular dynamics simulations. S.O., A. and F.W. performed accelerated evolution experiments. W.S.Y. supervised C.M, J.F. supervised the project. S.O. and J.F. wrote the paper with input from all authors.

