## Supplementary Materials for "Accelerated evolution of whole gene clusters by an engineered lytic phage system in *E. coli*"

### Extended Data Figures

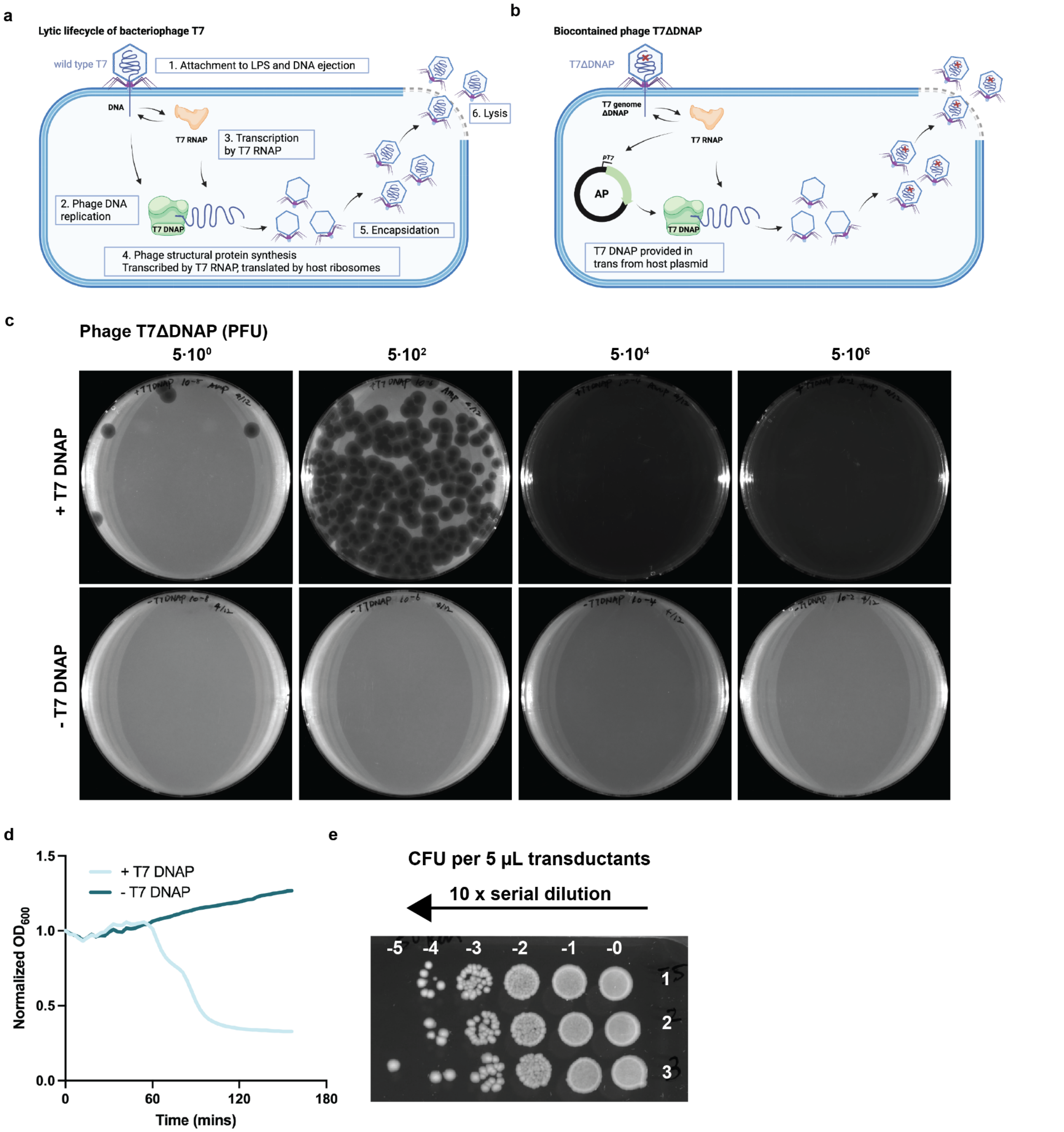


**Extended Data Fig. 1 | T7 phagemid packaging during lysis.**

**a.** Schematic of the T7 lytic cycle. T7 bacteriophage attaches to specific lipopolysaccharides (LPS) on the outer membrane of the *E. coli* host and ejects its DNA. Early genes like T7 DNA polymerase (T7 DNAP) and T7 RNA polymerase (T7 RNAP) are expressed first. T7 DNAP replicates the T7 genome in a bidirectional manner, producing concatemers. Meanwhile, T7 RNAP transcribes phage genes, which are then translated by host ribosomes to produce phage structural proteins. Towards the end of the lytic cycle, new phage particles are assembled, the T7 genome is packaged, and the cell lyses to release 180 progeny phages. The whole process takes 17 minutes. **b.** Biocontainment of phage T7ΔDNAP. Phage T7ΔDNAP is engineered to lack T7 DNAP, and the accessory plasmid (AP) carries T7 DNAP under the control of a T7 promoter. Upon infection, phage T7ΔDNAP expresses T7 RNAP that induces expression of T7 DNAP, leading to phage genome replication, packaging, and production of phage progeny. Phage T7ΔDNAP is biocontained and propagates only in *E. coli* cells harboring AP. **c.** Plaque assay of T7ΔDNAP infecting *E. coli* cells with or without AP carrying T7 DNAP. For cells expressing T7 DNAP, the bacteria lawn was completed cleared upon infection with T7ΔDNAP of $5x10^4$ PFU or higher. No plaques were observed on cells not expressing T7 DNAP. **d.** Lysis kinetics of T7ΔDNAP infecting *E. coli* cells with or without AP carrying T7 DNAP. MOI = 0.1. **e.** Transduction of a phagemid into new host cells by phage T7ΔDNAP. *E. coli* cells containing the AP and phagemid were lysed by addition of phage T7ΔDNAP. The lysate was washed with chloroform, transduced 1:100 into fresh *E. coli* cells and directly spotted with dilution on LB agar with 50 µg/mL kanamycin to select for cells that received the phagemid. Three independent replicates are shown.

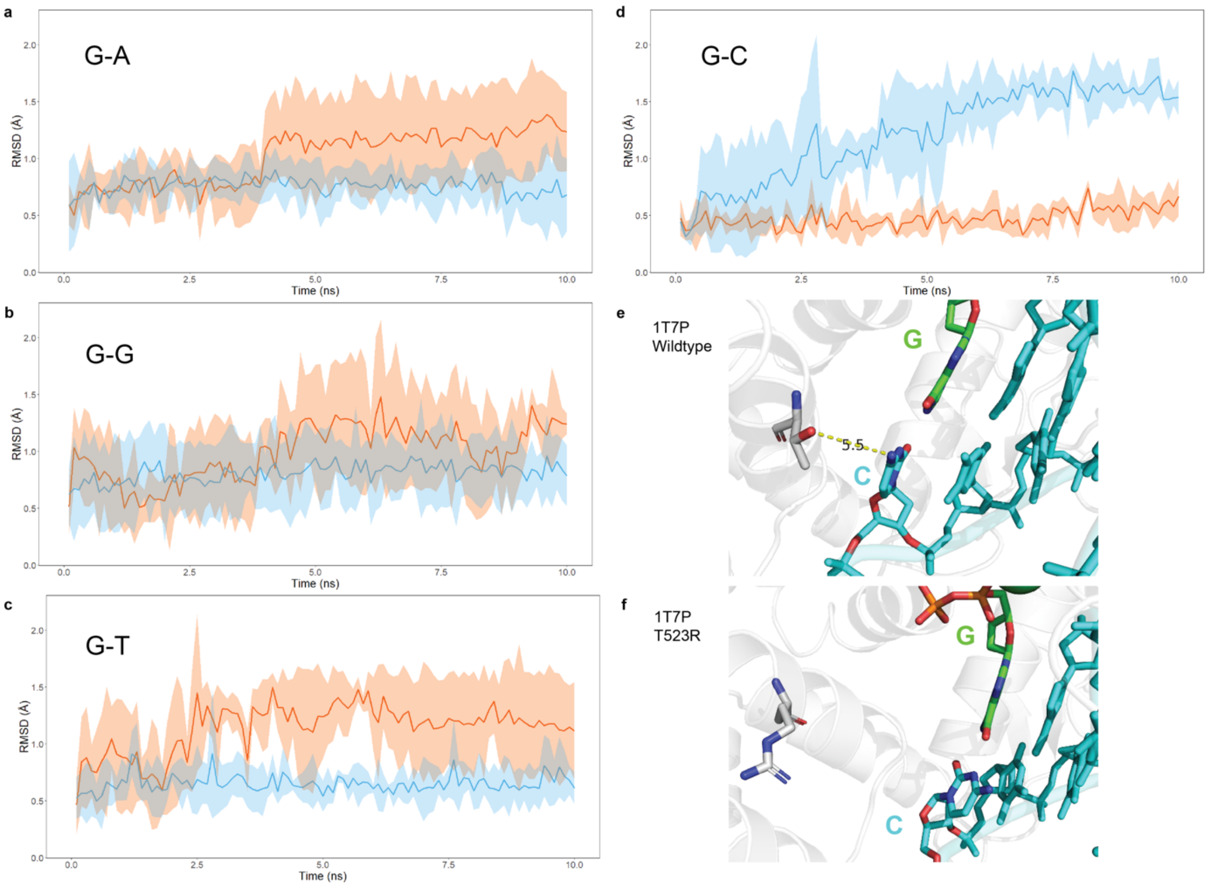


**Extended Data Fig. 2 | Molecular dynamics simulations of T7 DNAP crystal structure**

**a.** Root Mean Square Deviation (RMSD) values of dGTP incorrectly paired with adenine in wild type T7 DNAP (blue) and T523R mutant (orange) in triplicate. dGTP shows increased movement with R523 compared to T523 throughout the simulation. **b.** RMSD values of dGTP incorrectly paired with guanine in wild type T7 DNAP (blue) and T523R mutant (orange) in triplicate. dGTP shows slightly increased movement with R523 compared to T523, but this is less pronounced than with other mispairings. **c.** RMSD values of dGTP incorrectly paired with thymine in wild type T7 DNAP (blue) and T523R mutant (orange) in triplicate. dGTP shows increased movement with R523 compared to T523 throughout the simulation. **d.** RMSD values of dGTP correctly paired with cytosine in wild type T7 DNAP (blue) and T523R mutant (orange) in triplicate. Correctly paired dGTP shows notably low RMSD values in the T523R mutant, while RMSD increases in wild type compared to other conditions. This contrast correlates with a conformational change in the protein when matching nucleotides align in the mutated structure. R523 may interact strongly only with mismatched nucleotides, potentially contributing to increased error rates. **e.** Snapshot of wild type T7 DNAP with threonine in position 523 (grey) with correct pairing between incoming dGTP (green) and template cytosine (cyan). The α-helix with T523 maintains the confirmation observed for base mispairings in wild type and T523R T7 DNAP. **f.** Snapshot of T7 DNAP with T523R mutation (grey) and correct base pairing between incoming dGTP (green) and template cytosine (cyan). Correct base pairing prevents R523 from positioning between incoming and template nucleotides, resulting in an entirely different helix conformation.

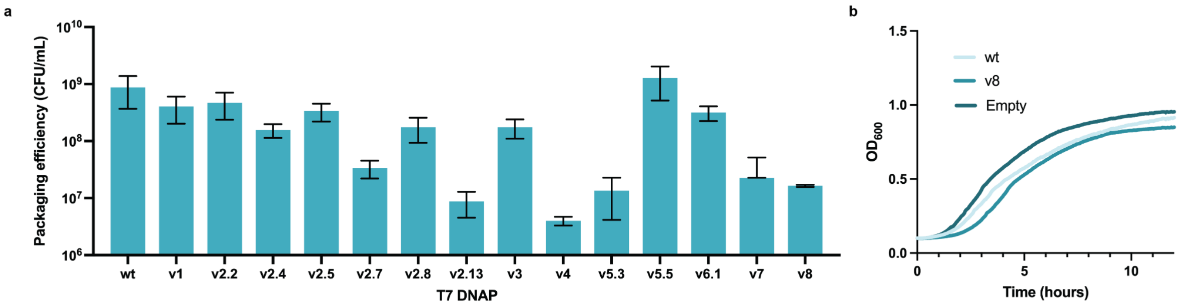


**Extended Data Fig. 3 | Transduction efficiency and growth rates for T7 DNAP variants.**

**a.** Quantification of phagemid packaging efficiency across wild type (wt) and engineered (v1-8) T7 DNAP variants by selection with kanamycin after transduction. Data shown as mean ± SD (n = 3). **b.** Growth kinetics of *E. coli* strains harboring no T7 DNAP (Empty), wild type T7 DNAP (wt), or the hypermutagenic T7 DNAP variant v8.

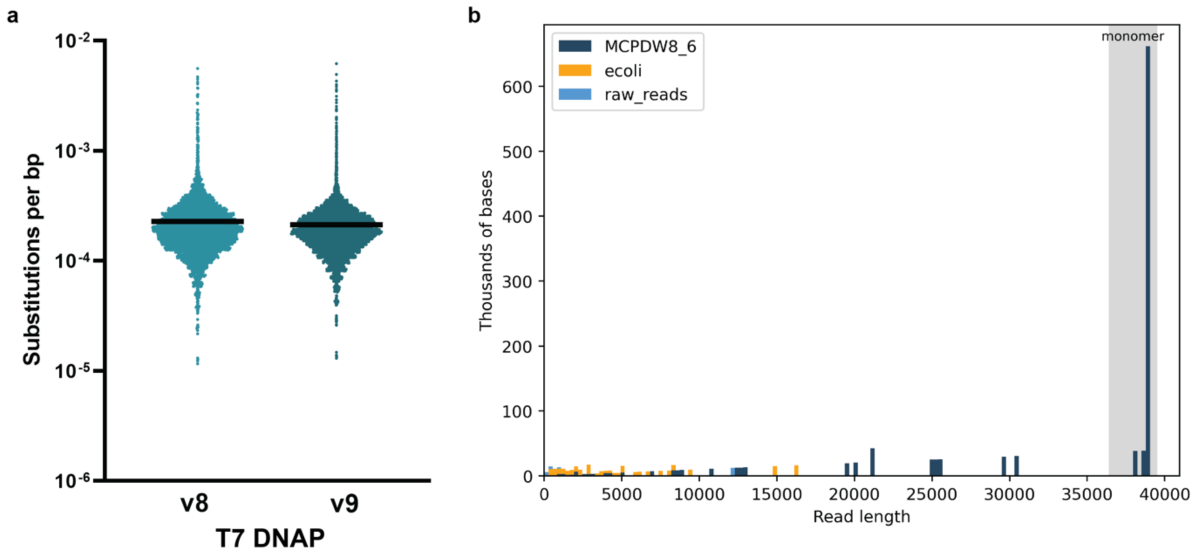


**Extended Data Fig. 4 | LySE replication, packaging and transduction of a 39 kb BAC-phagemid.**

**a.** Overall base substitution frequencies for T7 DNAP variants v8 and v9 derived from Illumina next-generation sequencing data (mean coverage >14,000x per base pair). **b.** Read length distribution from nanopore sequencing of transductants after one generation of LySE with 39 kb BAC-phagemid. *E. coli* cells containing accessory plasmid (WT T7 DNAP) and BAC-phagemid were lysed by addition of phage T7ΔDNAP. The lysate was washed with chloroform, transduced 1:100 into fresh *E. coli* cells and recovered overnight in LB medium with 50 µg/mL kanamycin to select for the BAC-phagemid. Phagemids were extracted by miniprep and sequenced by nanopore.

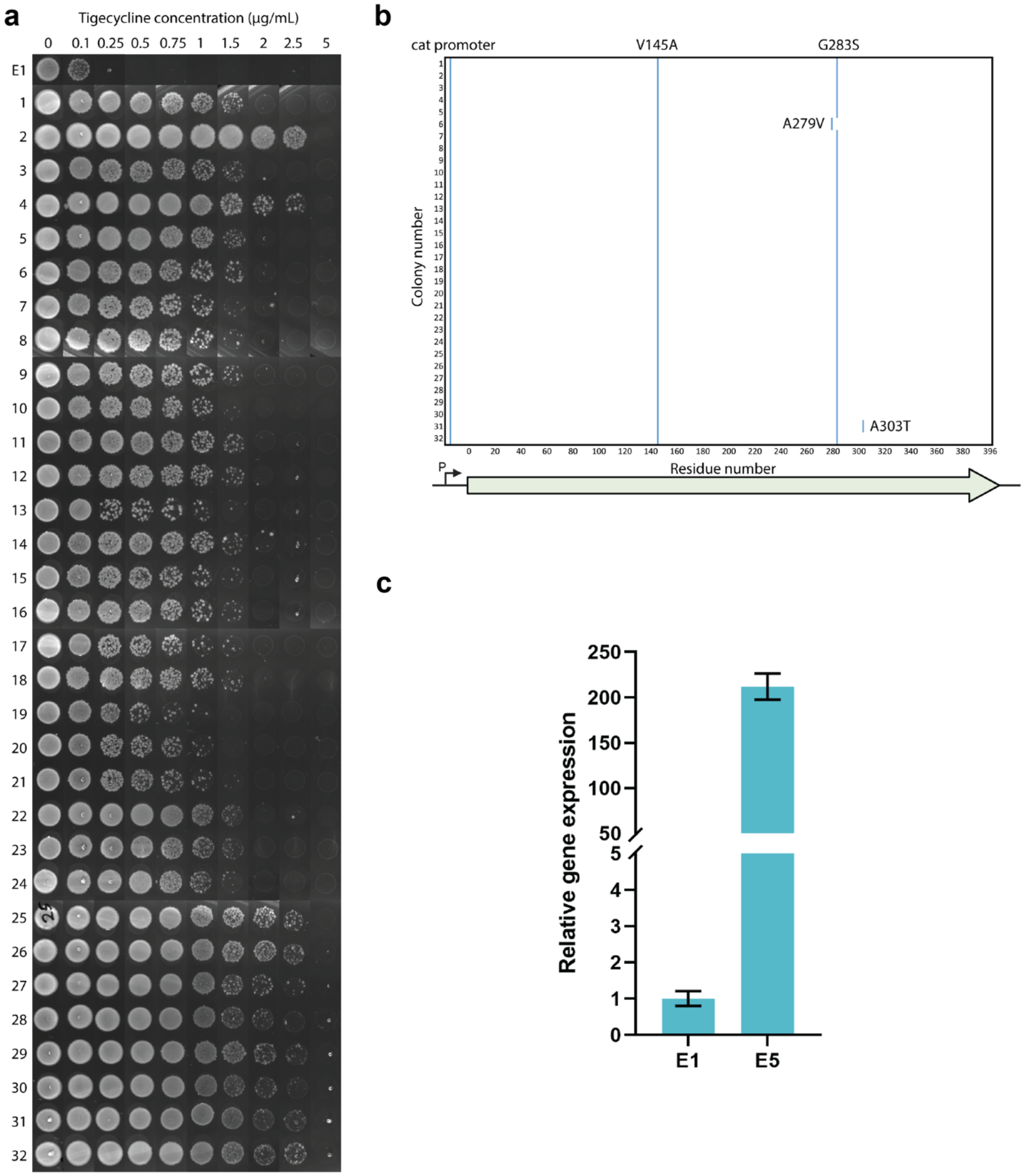


**Extended Data Fig. 5 | *tetA* mutants obtained from LySE evolution for tigecycline resistance.**

**a.** Screening of 32 clones from LySE E5 evolution of *tetA* gene for tigecycline resistance.
**b.** Sanger sequencing of the *tetA* gene in the 32 evolved clones. Convergence of sequences in the promoter region (pCAT) as well as in the coding sequence (V145A,

G283S). **c.** Normalized *tetA* expression levels in E5 LySE compared to E1 pools quantified by RT-qPCR (n = 3; mean ± SD).

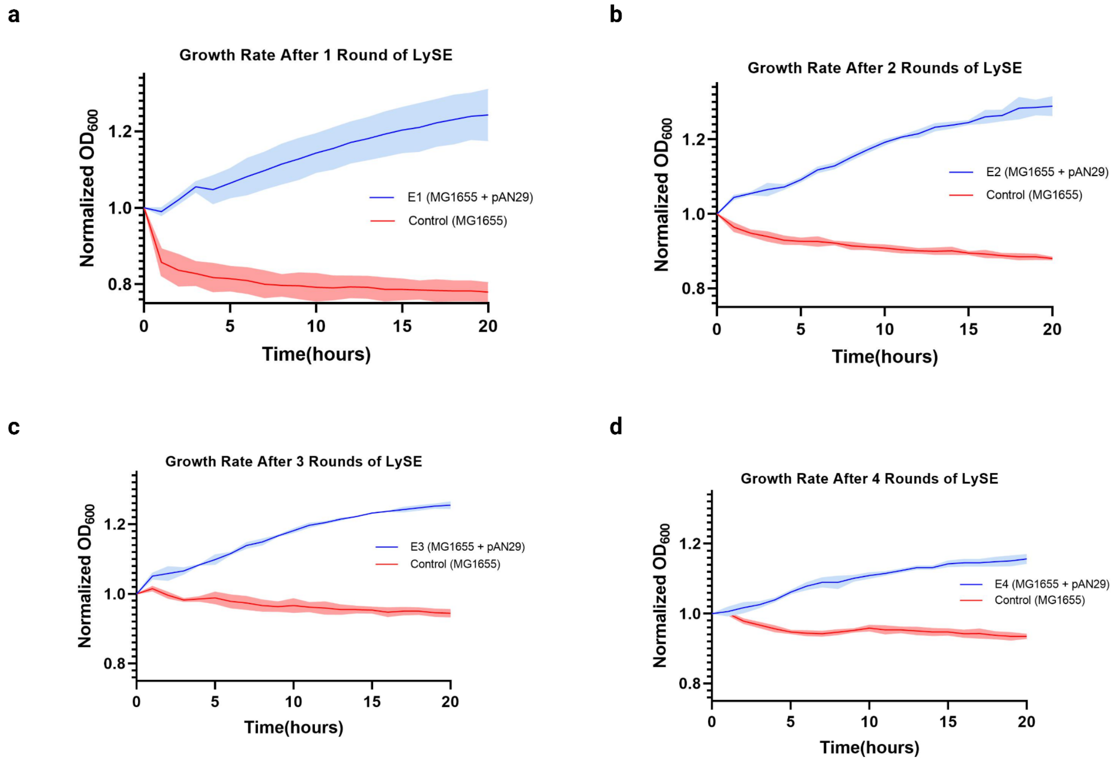


**Extended Data Fig. 6 | LySE evolution of ethylene glycol (EG) assimilation pathway and selection with only EG.** Initial LySE evolution with pure EG in M9 minimal medium showed no improvements, likely due to combined mutagenesis and carbon restriction stresses causing loss of potential mutants. We therefore implemented a semi-relaxing selection protocol with initial glucose supplementation followed by gradual glucose withdrawal. Shown here are growth rates of cells containing EG assimilation pathway phagemid after each round of LySE evolution in *E. coli*. Control is *E. coli* with no phagemid. Cultures were grown in M9 minimal medium with 10 g/L EG. Data are the mean ± standard deviation from three (n = 3) independent biological replicates. The best replicate from each round was picked for subsequent rounds of LySE.

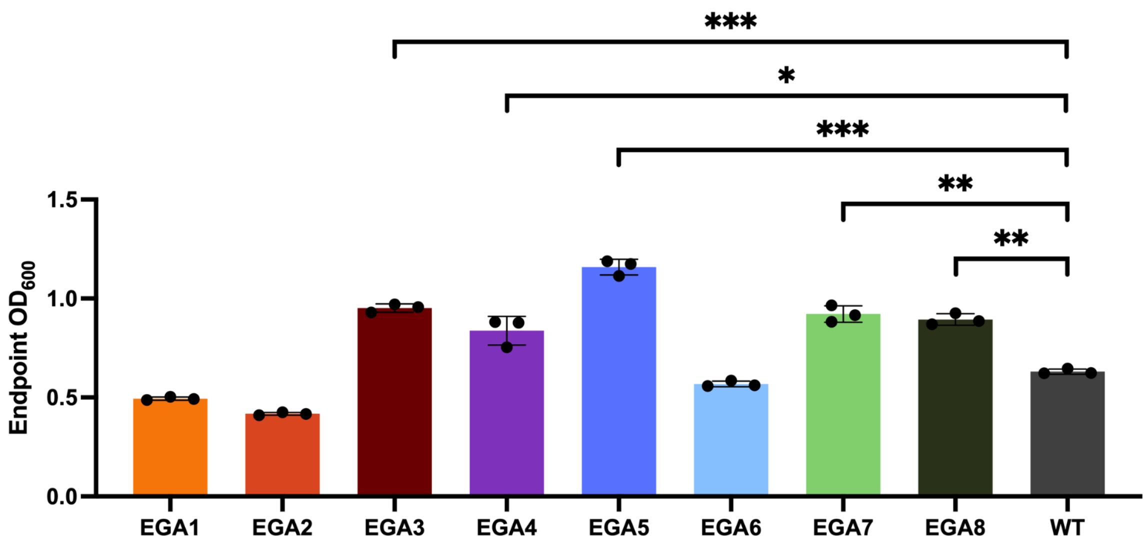


**Extended Data Fig. 7 | Endpoint biomass (t = 48 hours) of evolved ethylene glycol (EG) assimilation pathway phagemid clones after five generations of LySE.** Cultures were grown in M9 minimal medium with 10 g/L EG. Five clones exhibited accelerated growth on EG compared to the wild type (Starting from best: EGA5, EGA3, EGA7, EGA8, EGA4). Data are the mean ± standard deviation from three ($n = 3$) independent biological replicates. Significance was tested via two-tailed unpaired two-sample t-test. * $p < 0.01$; ** $p < 0.001$; *** $p < 0.0001$.

### Supplementary Data

**Supplementary Data 1: Phage T7ΔDNAP**
Phage genome map and DNA sequence

**Supplementary Data 2: Phagemid-p15A**
Phagemid map and DNA sequence

**Supplementary Data 3: Simulation MDP**
Molecular dynamics simulations

**Supplementary Data 4: Phagemid-BAC, pSJ77**
Phagemid map and DNA sequence

**Supplementary Data 5: Phagemid *tetA*, pSJ78**
Phagemid map and DNA sequence

**Supplementary Data 6: Phagemid EG pathway, pAN29**
Phagemid map and DNA sequence

**Supplementary Data 7: EGA1-8, nanopore sequencing**
DNA sequences

**Supplementary Data 8: Recombination plasmid, pPBG01**
Plasmid map and DNA sequence

**Supplementary Table 1: Strains, plasmids, oligos**
Lists of all strains, plasmids, and oligos used in this study

**Supplementary Table 2: T7 DNAP variants**
Lists of all T7 DNA variants used in this study

### Methods

General methods

All *E. coli* strains and plasmids created in this study are listed in **Supplementary Table 1**, together with all oligos. Phage T7 was obtained from ATCC (BAA-1025-B2). Cloning of all plasmids was carried out using MDS42 cells (Scarab Genomics, E-6265-05K). Plasmids were constructed with NEBuilder HiFi DNA Assembly (New England Biolabs) unless otherwise stated. Native *E. coli* and T7 genes were amplified by PCR directly from genomic DNA. Other genes were synthesized as gBlocks Gene Fragments (Integrated DNA Technologies), unless otherwise stated. PCR reactions were performed using PrimeSTAR GXL DNA Polymerase (Takara) for cloning and using Rapid Taq DNA polymerase (Vazyme) for genotyping. All oligos were synthesized by Integrated DNA Technologies. Sanger sequencing was performed by 1st BASE, Singapore. Nanopore sequencing was performed by Plasmidsaurus, CA, U.S.A. Sample preparation for sequencing was done according to each companies' protocol.

Culture media

*E. coli* was grown in LB medium or on LB agar (Bio Basic Asia Pacific, Singapore) added kanamycin (50 µg/mL), ampicillin (100 µg/mL), streptomycin (100 µg/mL), chloramphenicol (20 µg/mL), tetracycline (10 µg/mL), or hygromycin (200 µg/mL) where appropriate. Ethylene glycol (EG) selection and growth experiments were carried out without antibiotics in standard M9 minimal media (50 mM Na2HPO4, 20 mM KH2PO4, 1 mM NaCl, 20 mM NH4Cl, 2 mM MgSO4 and 100 µM CaCl2, 134 µM EDTA, 13 µM FeCl3·6H2O, 6.2 µM ZnCl2, 0.76 µM CuCl2·2H2O, 0.42 µM CoCl2·2H2O, 1.62 µM H3BO3, 0.081 µM MnCl2·4H2O). Carbon sources were used as indicated in the text.

Electrocompetent cell preparation

*E. coli* cells were inoculated in 10 mL LB and grown overnight at 37 °C and shaking at 225 rpm, with appropriate antibiotics if applicable. The next morning, cells were diluted 1:50 fold in the same media to a final volume of 500 mL, grown to $OD_{600}$ = 0.5, and incubated on ice for 20 min. The cells were spun down at 4000 g for 10 min at 4 °C. The supernatant was discarded, and the cells resuspended in chilled 50 mL sterile $dH_2O$. The cells were further washed once again with sterile $dH_2O$ followed by chilled 16% (w/v) glycerol. The cells were then resuspended in 1 mL chilled 16% glycerol and used immediately for transformation by electroporation, or flash frozen and stored at -80°C.

Scarless gene deletion by homologous recombination

Gene deletions were performed as previously described[50]. We created plasmid pPBG01 (**Supplementary Data 8**) for arabinose-inducible lambdaRED recombination by cloning genes *gam*, *beta*, and *exo* from *E. coli* C321.ΔA.opt (Addgene #87359) into a CloDF13 vector (Addgene #69669).

*E. coli* MG1655$^{rpslK43R}$ was used for all gene deletions. It was created by PCR amplification of *rpsL*K43R from *E. coli* DH10b, which we electroporated into competent and induced MG1655 cells harboring pPBG01. After recovery, the cells were plated on LB agar with streptomycin to select for successful recombination. We further created a double-selection cassette of negative selection gene *rpsL* (from MG1655) and positive selection gene *hygR* (from Addgene #104405) combined by overhang extension PCR. For gene deletions, we amplified the *rpsL-hygR* cassette by PCR using primers containing homology to the regions flanking the target gene to be deleted. We electroporated the the PCR product into competent and induced MG1655$^{rpslK43R}$ harbouring pPBG01 and selected for genomic integration with hygromycin. Successful gene deletions were verified by genotyping. To achieve scarless gene deletion, the *rpsL-hygR* cassette was subsequently removed by a second round of recombineering, by electroporating 10 µg of a 90-base oligo covering the genomic homology regions targeting the lagging strand. Loss of the *rpsL-hygR* cassette was selected for with streptomycin and verified by genotyping and Sanger sequencing.

<u>Phage plaque assay</u>

Phage from clarified phage lysate (or from rebooting in cell-free TXTL extract, see below) were serially diluted ten-fold in LB medium. Overnight cultures of *E. coli* host strains were prepared, diluted 4-fold in LB, and incubated with shaking for 1 hour at 37 °C. For each 100 mm petri dish, 100 µL of cells were mixed with 1 µL of serially diluted phage and incubated for 5 mins at 37 °C. The phage-cell mixtures were then mixed with 4 mL pre-warmed molten 0.35% top LB agar and immediately poured uniformly on to 100 mm petri dishes containing 15 mL of solidified 1.5% bottom LB agar. The top agar was left to solidify for 1 hour, then incubated for 4 hours at 37 °C for plaque formation.

<u>Phage titer determination</u>

Overnight cultures of *E. coli* were diluted and embedded in top agar as above, but without phage. Plates were dried for 1 hour. Each phage was serially diluted and 5 µL were spotted onto the top agar and left to dry for 15 min. The plates were incubated at 37 °C, facing down, for 4 h. The number of plaques was counted at each dilution to determine plaque forming units (PFU). The MOI was determined by taking the ratio of PFU of phage added to the colony forming units (CFU) of cells used for phage infection. We determined empirically that on average, approximate phage lysates are $10^{13}$ PFU/mL and diluted overnight cell cultures are $10^{11}$ CFU/mL.

<u>T7ΔDNAP phage assembly and rebooting in TXTL</u>

T7 genomic DNA excluding T7 DNAP was PCR amplified in five fragments of 10.0 kb (F1), 4.4 kb (F2), 3.5 kb (F4), 10.0 kb (F5) and 10.0 kb (F6), with 25-30 bp overlapping sequences. Fragment F3 containing the *trxA* gene was amplified from the *E. coli* genome, with overlapping sequences with F2 and F4 to replace T7 DNAP. Purified PCR fragments were mixed, including 300 ng of F3, and 100 ng of each of the other fragments. They were then assembled by Gibson assembly using NEBuilder HiFi DNA Assembly (New England Biolabs). The assembly mix, together with 50 ng of T7 genomic DNA, was mixed with myTXTL Linear DNA Expression Kit (Arbor Biosciences) according to the manufacturer's protocol and incubated overnight at 37 °C to generate a mixture of rebooted wild type T7 and T7ΔDNAP phage. To select for the mutant phage, a plaque assay was performed by infecting MG1655Δ*trxA* containing pSJ55, which expresses wild type T7 DNAP. Wild type T7 not expressing trxA would not propagate in this strain. Only T7ΔDNAP *trxA* would be able to propagate in this strain and form plaques. We confirmed successful generation of T7ΔDNAP *trxA* by genotyping and Sanger sequencing the genomic region where T7 DNAP was deleted from. Sequences of the fragments and primers used are listed in **Supplementary Data** 2.

<u>Phage infection kinetic assay</u>

Infection kinetics were carried out in 96 well plates in BioTek Synergy HTX Microplate Reader (Agilent). Overnight cultures of *E. coli* host strains were prepared, diluted by half with LB medium, and 100 µL added to each well. In each well, 20 µL of different serial phage dilutions were added to tune MOI. Each condition was replicated in three different wells. A lid was added to the 96 well plates to reduce evaporation during acquisition. The microplate reader was set to 37 °C with continuous orbital shaking at 300 rpm. $OD_{600}$ was measured every 3 mins and monitored for at least 2 hours.

<u>Yeast assembly</u>

Large phagemids were assembled by transformation-associated recombination in yeast[51] as previously described[50]. *S. cerevisiae* BY4741 (MATα his3Δ1 leu2Δ0 met15Δ0 ura3Δ0) was obtained from ATCC (#201388). We modified the cell-wall digestion step slightly: We only used 1 µL zymolyase, and we measured $OD_{660}$ every 10 min after 30 min zymolyase digestion. Digested cells were resuspended by swirling and inverting the tubes.

Phagemid assembly was assessed by genotyping of the resulting yeast colonies, amplifying 350-1000 bp across the intersections of assembled fragments. Positive yeast colonies were cultured, phagemids were isolated with Zymoprep Yeast Plasmid Miniprep Kit (Zymo Research, USA) and electroporated into *E. coli* MDS42 with selection. Individual colonies were genotyped again with the same primers, and phagemids were isolated from positive clones using Monarch Plasmid Miniprep Kit and verified by Nanopore sequencing.

<u>General LySE strain preparation</u>

All LySE experiments were conducted with *E. coli* MG1655. Cells were first transformed with accessory plasmids (APs) containing different T7 DNAP variants by electroporation as described above and selected with ampicillin. The protein sequences of the T7 DNAP variants are given in **Supplementary Table 2**. Cells containing the AP are then transformed a second time with the phagemid (PM) by electroporation and selected with both ampicillin and kanamycin. Genotyping was performed at each step to confirm successful transformation.

LySE cycle

To propagate T7ΔDNAP phage, MG1655 with pSJ55 was grown overnight in LB medium with ampicillin, then diluted with equal volumes of LB the next day. T7ΔDNAP phage lysates were mixed with the diluted cells at a volume ratio of 1:1000 (approximately MOI = 0.1) and incubated at 37 °C with shaking for 2 hours until there was no further reduction in $OD_{600}$. The phage lysates were then washed once with equal volumes of chloroform to remove residual cells and debris.

The LySE cycle begins with an overnight culture of MG1655 cells containing AP and PM. Overnight culture cells were diluted with equal volumes of LB the next day. To 100 µL of diluted cells, 20 µL of phage is added (approximately MOI = 10; High MOI) and incubated at 37 °C with shaking for 2 hours until there was no further reduction in $OD_{600}$. The phage lysates were washed once with equal volumes of chloroform. Next, to 1 mL overnight culture of MG1655 cells containing only AP diluted with equal volumes of LB, we mixed 10 µL phage lysates containing phagemids and incubated at 37 °C with shaking for 1 hour for complete transduction of phagemids (approximately MOI = 1; Low MOI). Phagemid packaging efficiency was determined by serial dilution of the transduced cells, followed by spot plating on LB agar with kanamycin to determine CFU of transduced phagemid. To continue the LySE cycle, transduced cells were diluted in selection media and grown to confluency. The exact protocol for selection and culture recovery is specific to each evolution experiment, but should include addition of ampicillin to maintain the AP. The next LySE cycle is continued by adding phage at high MOI to lyse cultured cells.

Fluctuation assay

We performed Luria-Delbrück fluctuation analysis to quantify the mutation rate per generation of LySE for each T7 DNAP variant. We cloned pSJ51, a phagemid constitutively expressing a chloramphenicol resistance gene (CmR) containing a premature stop codon (Q38TAG). MG1655 containing pSJ51 and an AP encoding the T7 DNAP variant to be tested was grown overnight, diluted and lysed as per standard LySE protocols. After one generation of phagemid replication and transduction, transduced cells were serially diluted ten-fold and spotted on LB agar with kanamycin to quantify total transduction CFU, and on LB agar with chloramphenicol to quantify stop codon reversion CFU respectively. After overnight incubation at 37 °C, CFUs from three independent replicates were counted. Mutation per generation per base pair µ (s.p.b.) was calculated using µ (s.p.b.) = m / (R × C). Mutation frequency (m) was calculated based on the ratio of cells grown on chloramphenicol to that grown on kanamycin. For the parameter R, which is the number of distinct mutation sites that make the resistance gene effective, we found that 8/9 possible single base substitutions (which yield sense codons) can result in chloramphenicol resistance. So, R = 8/3. C is the gene copy number. Since T7 packages concatemeric phagemid DNA via a head-full mechanism, multiple CmR copies are expected per phage, while only a single reverted TAG is sufficient to confer chloramphenicol resistance. We therefore determined C by taking the fraction of phagemid to T7 genome size, so C = 39937/3469 = 11.5.

LacZ inactivation assay

LacZ inactivation assay was performed to quantify the mutation rates of more error-prone T7 DNAP variants. We performed scarless deletion of *lacIZYA* from *E coli* MG1655 as described above. We cloned pSJ78, a phagemid constitutively expressing wild type *lacZ.* MG1655Δ*lacIZYA* containing pSJ78 and an AP encoding the T7 DNAP variant to be tested was grown overnight, diluted and lysed as per standard LySE protocols. After one generation of phagemid replication and transduction, transduced cells were serially diluted ten-fold and spotted on LB agar with kanamycin and 200 µg/mL X-Gal (Thermo Scientific). After overnight incubation at 37 °C, CFUs from three independent replicates were counted. The fraction of white or light blue colonies (lacZ- phenotype) was counted as a function of all colonies (blue+light blue+white) and used as a measure of mutation frequency for the lacZ cassette.

Molecular dynamics simulation

We used the crystal structure published by Doublié *et al*. (PDB: 1T7P)[29] as the basis for our molecular dynamics simulations of the wild type and T523R mutant T7 DNAP. 1T7P contains a growing DNA strand terminated with a dideoxy cytosine nucleotide, and the incoming nucleotide is dideoxyguanosine triphosphate (ddGTP). To represent the real biomolecules as closely as possible, we manually added 3'-hydroxyl groups to the chain-terminating nucleotide of the growing DNA strand and ddGTP. A wild type and mutant T523R T7 DNAP variants were created *in silico*, and subsets of these containing DNA substitutions of the leading cytosine nucleotide on the template strand were implemented to study the effect T523R has on base mispairing. The structure is dissolved in water under physiological conditions using the solution builder on CHARMM-GUI. We placed each protein structure in a 130Å × 130Å × 130Å simulation box with water containing 215/216 sodium ions and 183 chloride ions (150 mM) to balance protein charges at pH 7.0. For each condition, we ran a 5000 step steepest descent energy minimisation. This was followed by an NVT equilibration with a simulation time of 125 ps (125,000 steps) at 303.15 K. The subsequent NPT production simulation is 10 ns (5,000,000 steps) at 303.15 K. All simulations used the CHARMM36m force field and were run using CUDA supported GROMACS, version 2023.3, on high performance computing facilities (National University of Singapore HPC).

Illumina NGS and data analysis

We cloned pSJ77, a 39 kb BAC-phagemid by yeast assembly. *E. coli* MG1655 containing pSJ77 and an AP encoding either T7 DNAP variant v8 or v9 was grown overnight, diluted and lysed as per standard LySE protocols. After one generation of phagemid replication and transduction into MG1655 with no plasmids, transduced cells were diluted 10-fold in LB with kanamycin and recovered overnight. The recovered mutated BAC-phagemid library was purified by ZymoPURE Plasmid Miniprep Kit (Zymo Research, USA). Illumina Next-Generation Sequencing (NGS) of the BAC-phagemid library was performed by Bio Basic Asia Pacific. Next generation sequencing library preparations were constructed following the manufacturer's protocol. For each sample, 200 µg DNA was randomly fragmented by Covaris to an average size of 300-350 bp. The fragments were treated with End Prep Enzyme Mix for end repairing, 5' Phosphorylation and 3' adenylated, to add adaptors to both ends. Size selection of Adaptor-ligated DNA was then performed by DNA Cleanup beads. Each sample was then amplified by PCR for 8 cycles using P5 and P7 primers, with both primers carrying sequences that can anneal with flowcell to perform bridge PCR and P7 primer carrying a six-base index allowing for multiplexing. The PCR products were cleaned up and validated using an Agilent 2100 Bioanalyzer. The qualified libraries were sequenced pair end PE150 on the Illumina Novaseq System. Fastp (v0.23.0) was used for quality control and preprocessing, including removal of adaptor sequences, PCR primers, reads with more than 14 N bases, and reads with less than 40% bases above Q20. The cleaned data was then mapped to the reference genome using the Sentieon pipeline (v202112.02). A custom python script using the pysam (v0.23.0) module was used to align the NGS reads with Q score ⩾30 to the wild type sequence and count the nucleotide positions from which the experimental sample deviates from the wild type sequence. General mismatch rates and A:T>G:C and C:G>T:A mutation rates per 5 bp were calculated and plotted. Overall mutational spectra, and for every 10,000 bp, were calculated. We yielded an average of > 14,000 reads per position for each of the sequenced samples.

LySE evolution of tetA

We cloned pSJ78, a phagemid constitutively expressing the *tetA* tetracycline efflux gene. *E. coli* MG1655 containing pSJ78 and pSJ139 (T7 DNAP v9) was grown overnight, diluted and lysed as per standard LySE protocols. After one generation of phagemid replication and transduction into MG1655 with pSJ139, transduced cells were diluted 10-fold with LB with ampicillin, kanamycin, and 0.1 µg/mL tigecycline. The cells were incubated overnight at 37 °C with shaking at 225 rpm. The next day, the cells were diluted with equal volumes of LB, and T7ΔDNAP phages were added to lyse the culture, starting another round of

LySE. Evolution cycles were repeated another four times for a total of five evolution cycles, using the best growing cultures from the previous cycle as the starting material for the next cycle. Tigecycline concentrations were incrementally increased from LySE E1 to E5, with concentrations 0.1 µg/mL, 0.25 µg/mL, 0.5 µg/mL, 0.75 µg/mL and 1 µg/mL respectively. After the 5$^{th}$ LySE cycle, the best growing culture was streaked on LB agar with kanamycin and 1 µg/mL tigecycline. Individual colonies were picked and inoculated separately in 1 mL LB with kanamycin in 24-well plates and grown overnight. 32 evolved clones were spotted on LB agar with increasing concentrations of tigecycline to test resistance. The 32 clones were PCR amplified for the *tetA* cassette using PrimeSTAR GXL DNA Polymerase and the amplicon was sequenced by Sanger sequencing.

For ALE, *E. coli* MG1655 containing pSJ78 and pSJ55 (WT T7 DNAP) was grown overnight in LB with kanamycin and ampicillin, then diluted 10-fold with LB with kanamycin, ampicillin, and 0.1 µg/mL tigecycline. The cells were incubated overnight at 37 °C with shaking at 225 rpm. The next day, the cells were diluted another 10-fold to continue ALE. A total of 5 passages were performed, each time with increasing concentrations of tigecycline identical to the LySE schedule. Cells at the 5$^{th}$ passage were spotted on LB agar with increasing concentrations of tigecycline to test resistance. The cells were also lysed by addition of T7ΔDNAP, the lysate washed with chloroform, and phagemids transduced to fresh MG1655 with no plasmids. The transduced cells were diluted 10-fold with LB with kanamycin, then grown overnight. The recovered E5T ALE cells were then spotted on LB agar with tigecycline to test resistance after transduction.

Quantitative real-time PCR (qPCR)

Total RNA was first isolated from the *E. coli* cells. Cells were grown overnight and diluted 1:50 with LB and appropriate antibiotics and cultured until $OD_{600}$ = 0.5. Then, 500 µL of cells were transferred into an Eppendorf tube, spun down, and the pellet dried by dabbing on a paper towel. The pellet was resuspended in 100 µL of Tris-EDTA buffer pH 8.5 with 15 mg/mL lysozyme, vortex for 10 s and incubated at room temperature for 5 mins with shaking. To the mixture, 400 µL of TRK Lysis buffer (Omega Biotek) with 4 µL 2-mercaptoethanol was added and total RNA was extracted immediately with E.Z.N.A RNA isolation kit (Omega Biotek). Purified RNA was quantified with NanoDrop (Thermo Fisher Scientific). One microgram of RNA was converted to cDNA using the GoScript™ Reverse Transcriptase (Promega). qPCR was performed with GoTaq qPCR (Promega) using the CFX Opus 96 Real-Time PCR System (Bio Rad). Fold changes were normalized to 16s rRNA and are based on relative expression values calculated using the 2−ΔΔCT method.

LySE evolution of EG assimilation pathway

The *gox0313* gene (UniProt: Q5FU50) was synthesized by GentleGen, China. We cloned pAN29, a phagemid containing metabolic pathway genes for complete assimilation of EG.

*E. coli* MG1655 containing pAN29 and pSJ139 (T7 DNAP v9) was grown overnight, diluted and lysed as per standard LySE protocols. After one generation of phagemid replication and transduction into MG1655 with pSJ139, transduced cells were diluted 10-fold with LB with ampicillin and kanamycin and incubated overnight at 37 °C with shaking at 225 rpm. The next day, 5 mL of recovered cells were pelleted by centrifugation at 3900 x *g* for 3 mins, washed three times with M9 medium, and then resuspended in M9 medium with EG, with or without glucose at concentrations as indicated in the text until the $OD_{600}$ is approximately 0.2. No antibiotics were used for selection. To each well in a 24-well plate, 1 mL of the cell suspension was added and cultured overnight at 37 °C with shaking at 300 rpm in a Thermo-Shaker PST-60HL-4 Microplate Reader (BioSan, Latvia). Every hour, $OD_{600}$ was measured to monitor cell growth. Evolution cycles were repeated another four times for a total of five evolution cycles, using the best growing cultures from the previous cycle as the starting material for the next cycle. After the $5^{th}$ LySE cycle, the best growing culture was streaked on M9 minimal agar plates supplemented with 2 g/L of glucose and 10 g/L of EG. Individual colonies were picked and inoculated separately in 1 mL LB with kanamycin in 24-well plates and grown overnight. The following day, the cultures were washed with M9 media at 3900 x *g* for 1 min. Subsequently, the cells were inoculated in M9 medium supplemented with 10 g/L EG in 12 mL culture tubes for 6 hours to reach OD600=0.1. 200 μL of each culture was transferred into a sterile 96-well microplate as triplicates and incubated overnight with continuous orbital shaking at 600 rpm. Growth rates were determined by measuring $OD_{600}$ at 12, 18, 24, 36, 42, and 48 hours post-inoculation using the microplate reader. Eight clones were PCR amplified for the phagemid using PrimeSTAR GXL DNA Polymerase and the amplicon was sequenced by nanopore.

For ALE, 5 mL *E. coli* MG1655 containing pAN29 was grown overnight in LB with kanamycin, washed three times with M9 medium, and then resuspended in M9 medium with EG, with or without glucose at concentrations as indicated in the text until the $OD_{600}$ is approximately 0.2. To each well in a 24-well plate, 1 mL of the cell suspension was added and cultured overnight. The next day, 1 μL of the cells were added to 10 mL of LB with kanamycin and grown overnight before 5 mL was taken and washed, following the same washing, resuspension procedure as above for the next round of ALE. Evolution cycles were repeated another four times for a total of five evolution cycles, using the best growing cultures from the previous cycle as the starting material for the next cycle.

**Supplementary Data 1: T7 delta DNAP**

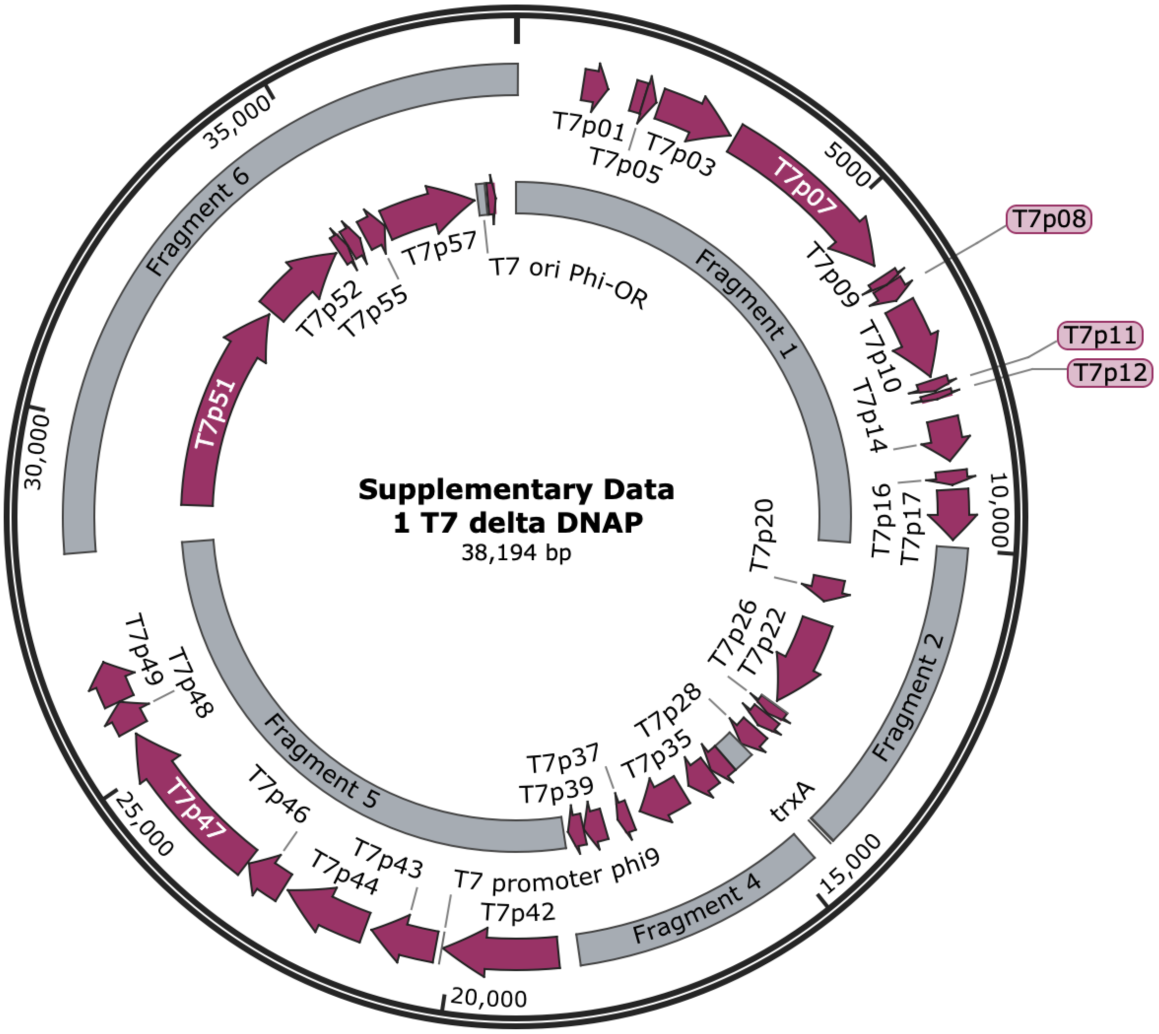


TCTCTGTGTCCCTTCTCACAGTGTACGGACCTAAAGTTCCCCCATAGGGGGTACCTAAAGCC
CAGCCAATCACCTAAAGTCAACCTTCGGTTGACCTTGAGGGTTCCCTAAGGGTTGGGGATGA
CCCTTGGGTTTGTCTTTGGGTGTTACCTTGAGTGTCTCTCTGTGTCCCTATCTGTTACAGTC
TCCTAAAGTATCCTCCTAAAGTCACCTCCTAACGTCCATCCTAAAGCCAACACCTAAAGCCT
ACACCTAAAGACCCATCAAGTCAACGCCTATCTTAAAGTTTAAACATAAAGACCAGACCTAA
AGACCAGACCTAAAGACACTACATAAAGACCAGACCTAAAGACGCCTTGTTGTTAGCCATAA
AGTGATAACCTTTAATCATTGTCTTTATTAATACAACTCACTATAAGGAGAGACAACTTAAA
GAGACTTAAAAGATTAATTTAAAATTTATCAAAAAGAGTATTGACTTAAAGTCTAACCTATA
GGATACTTACAGCCATCGAGAGGGACACGGCGAATAGCCATCCCAATCGACACCGGGGTCAA
CCGGATAAGTAGACAGCCTGATAAGTCGCACGAAAAACAGGTATTGACAACATGAAGTAACA
TGCAGTAAGATACAAATCGCTAGGTAACACTAGCAGCGTCAACCGGGCGCACAGTGCCTTCT
AGGTGACTTAAGCGCACCACGGCACATAAGGTGAAACAAAACGGTTGACAACATGAAGTAAA
CACGGTACGATGTACCACATGAAACGACAGTGAGTCACCACACTGAAAGGTGATGCGGTCTA
ACGAAACCTGACCTAAGACGCTCTTTAACAATCTGGTAAATAGCTCTTGAGTGCATGACTAG
CGGATAACTCAAGGGTATCGCAAGGTGCCCTTTATGATATTCACTAATAACTGCACGAGGTA
ACACAAGATGGCTATGTCTAACATGACTTACAACAACGTTTTCGACCACGCTTACGAAATGC
TGAAAGAAAACATCCGTTATGATGACATCCGTGACACTGATGACCTGCACGATGCTATTCAC

22 ATGGCTGCCGATAATGCAGTTCCGCACTACTACGCTGACATCTTTAGCGTAATGGCAAGTGA
23 GGGCATTGACCTTGAGTTCGAAGACTCTGGTCTGATGCCTGACACCAAGGACGTAATCCGCA
24 TCCTGCAAGCGCGTATCTATGAGCAATTAACGATTGACCTCTGGGAAGACGCAGAAGACTTG
25 CTCAATGAATACTTGGAGGAAGTCGAGGAGTACGAGGAGGATGAAGAGTAATGTCTACTACC
26 AACGTGCAATACGGTCTGACCGCTCAAACTGTACTTTTCTATAGCGACATGGTGCGCTGTGG
27 CTTTAACTGGTCACTCGCAATGGCACAGCTCAAAGAACTGTACGAAAACAACAAGGCAATAG
28 CTTTAGAATCTGCTGAGTGATAGACTCAAGGTCGCTCCTAGCGAGTGGCCTTTATGATTATC
29 ACTTTACTTATGAGGGAGTAATGTATATGCTTACTATCGGTCTACTCACCGCTCTAGGTCTA
30 GCTGTAGGTGCATCCTTTGGGAAGGCTTTAGGTGTAGCTGTAGGTTCCTACTTTACCGCTTG
31 CATCATCATAGGAATCATCAAAGGGGCACTACGCAAATGATGAAGCACTACGTTATGCCAAT
32 CCACACGTCCAACGGGGCAACCGTATGTACACCTGATGGGTTCGCAATGAAACAACGAATCG
33 AACGCCTTAAGCGTGAACTCCGCATTAACCGCAAGATTAACAAGATAGGTTCCGGCTATGAC
34 AGAACGCACTGATGGCTTAAAGAAAGGTTATATGCCCAATGGCACACTATACGCTGCAAATC
35 GGCGAATAGTGAGAACTTGGCGAGAGAACAACCTCGAACGCCGCAAGGACAAGAGAGGGCGG
36 CGTGGCATAGACGAAAGGAAAAGGTTAAAGCCAAGAAACTCGCCGCACTTGAACAGGCACTA
37 GCCAACACACTGAACGCTATCTCATAACGAACATAAAGGACACAATGCAATGAACATTACCG
38 ACATCATGAACGCTATCGACGCAATCAAAGCACTGCCAATCTGTGAACTTGACAAGCGTCAA
39 GGTATGCTTATCGACTTACTGGTCGAGATGGTCAACAGCGAGACGTGTGATGGCGAGCTAAC
40 CGAACTAAATCAGGCACTTGAGCATCAAGATTGGTGGACTACCTTGAAGTGTCTCACGGCTG
41 ACGCAGGGTTCAAGATGCTCGGTAATGGTCACTTCTCGGCTGCTTATAGTCACCCGCTGCTA
42 CCTAACAGAGTGATTAAGGTGGGCTTTAAGAAAGAGGATTCAGGCGCAGCCTATACCGCATT
43 CTGCCGCATGTATCAGGGTCGTCCTGGTATCCCTAACGTCTACGATGTACAGCGCCACGCTG
44 GATGCTATACGGTGGTACTTGACGCACTTAAGGATTGCGAGCGTTTCAACAATGATGCCCAT
45 TATAAATACGCTGAGATTGCAAGCGACATCATTGATTGCAATTCGGATGAGCATGATGAGTT
46 AACTGGATGGGATGGTGAGTTTGTTGAAACTTGTAAACTAATCCGCAAGTTCTTTGAGGGCA
47 TCGCCTCATTCGACATGCATAGCGGGAACATCATGTTCTCAAATGGAGACGTACCATACATC
48 ACCGACCCGGTATCATTCTCGCAGAAGAAAGACGGTGGCGCATTCAGCATCGACCCTGAGGA
49 ACTCATCAAGGAAGTCGAGGAAGTCGCACGACAGAAAGAAATTGACCGCGCTAAGGCCCGTA
50 AAGAACGTCACGAGGGGCGCTTAGAGGCACGCAGATTCAAACGTCGCAACCGCAAGGCACGT
51 AAAGCACACAAAGCTAAGCGCGAAAGAATGCTTGCTGCGTGGCGATGGGCTGAACGTCAAGA
52 ACGGCGTAACCATGAGGTAGCTGTAGATGTACTAGGAAGAACCAATAACGCTATGCTCTGGG
53 TCAACATGTTCTCTGGGGACTTTAAGGCGCTTGAGGAACGAATCGCGCTGCACTGGCGTAAT
54 GCTGACCGGATGGCTATCGCTAATGGTCTTACGCTCAACATTGATAAGCAACTTGACGCAAT
55 GTTAATGGGCTGATAGTCTTATCTTACAGGTCATCTGCGGGTGGCCTGAATAGGTACGATTT
56 ACTAACTGGAAGAGGCACTAAATGAACACGATTAACATCGCTAAGAACGACTTCTCTGACAT
57 CGAACTGGCTGCTATCCCGTTCAACACTCTGGCTGACCATTACGGTGAGCGTTTAGCTCGCG
58 AACAGTTGGCCCTTGAGCATGAGTCTTACGAGATGGGTGAAGCACGCTTCCGCAAGATGTTT
59 GAGCGTCAACTTAAAGCTGGTGAGGTTGCGGATAACGCTGCCGCCAAGCCTCTCATCACTAC
60 CCTACTCCCTAAGATGATTGCACGCATCAACGACTGGTTTGAGGAAGTGAAAGCTAAGCGCG
61 GCAAGCGCCCGACAGCCTTCCAGTTCCTGCAAGAAATCAAGCCGGAAGCCGTAGCGTACATC
62 ACCATTAAGACCACTCTGGCTTGCCTAACCAGTGCTGACAATACAACCGTTCAGGCTGTAGC
63 AAGCGCAATCGGTCGGGCCATTGAGGACGAGGCTCGCTTCGGTCGTATCCGTGACCTTGAAG
64 CTAAGCACTTCAAGAAAAACGTTGAGGAACAACTCAACAAGCGCGTAGGGCACGTCTACAAG
65 AAAGCATTTATGCAAGTTGTCGAGGCTGACATGCTCTCTAAGGGTCTACTCGGTGGCGAGGC
66 GTGGTCTTCGTGGCATAAGGAAGACTCTATTCATGTAGGAGTACGCTGCATCGAGATGCTCA
67 TTGAGTCAACCGGAATGGTTAGCTTACACCGCCAAAATGCTGGCGTAGTAGGTCAAGACTCT
68 GAGACTATCGAACTCGCACCTGAATACGCTGAGGCTATCGCAACCCGTGCAGGTGCGCTGGC
69 TGGCATCTCTCCGATGTTCCAACCTTGCGTAGTTCCTCCTAAGCCGTGGACTGGCATTACTG
70 GTGGTGGCTATTGGGCTAACGGTCGTCGTCCTCTGGCGCTGGTGCGTACTCACAGTAAGAAA
71 GCACTGATGCGCTACGAAGACGTTTACATGCCTGAGGTGTACAAAGCGATTAACATTGCGCA
72 AAACACCGCATGGAAAATCAACAAGAAAGTCCTAGCGGTCGCCAACGTAATCACCAAGTGGA

73 AGCATTGTCCGGTCGAGGACATCCCTGCGATTGAGCGTGAAGAACTCCCGATGAAACCGGAA
74 GACATCGACATGAATCCTGAGGCTCTCACCGCGTGGAAACGTGCTGCCGCTGCTGTGTACCG
75 CAAGGACAAGGCTCGCAAGTCTCGCCGTATCAGCCTTGAGTTCATGCTTGAGCAAGCCAATA
76 AGTTTGCTAACCATAAGGCCATCTGGTTCCCTTACAACATGGACTGGCGCGGTCGTGTTTAC
77 GCTGTGTCAATGTTCAACCCGCAAGGTAACGATATGACCAAAGGACTGCTTACGCTGGCGAA
78 AGGTAAACCAATCGGTAAGGAAGGTTACTACTGGCTGAAAATCCACGGTGCAAACTGTGCGG
79 GTGTCGATAAGGTTCCGTTCCCTGAGCGCATCAAGTTCATTGAGGAAAACCACGAGAACATC
80 ATGGCTTGCGCTAAGTCTCCACTGGAGAACACTTGGTGGGCTGAGCAAGATTCTCCGTTCTG
81 CTTCCTTGCGTTCTGCTTTGAGTACGCTGGGGTACAGCACCACGGCCTGAGCTATAACTGCT
82 CCCTTCCGCTGGCGTTTGACGGGTCTTGCTCTGGCATCCAGCACTTCTCCGCGATGCTCCGA
83 GATGAGGTAGGTGGTCGCGCGGTTAACTTGCTTCCTAGTGAAACCGTTCAGGACATCTACGG
84 GATTGTTGCTAAGAAAGTCAACGAGATTCTACAAGCAGACGCAATCAATGGGACCGATAACG
85 AAGTAGTTACCGTGACCGATGAGAACACTGGTGAAATCTCTGAGAAAGTCAAGCTGGGCACT
86 AAGGCACTGGCTGGTCAATGGCTGGCTTACGGTGTTACTCGCAGTGTGACTAAGCGTTCAGT
87 CATGACGCTGGCTTACGGGTCCAAAGAGTTCGGCTTCCGTCAACAAGTGCTGGAAGATACCA
88 TTCAGCCAGCTATTGATTCCGGCAAGGGTCTGATGTTCACTCAGCCGAATCAGGCTGCTGGA
89 TACATGGCTAAGCTGATTTGGGAATCTGTGAGCGTGACGGTGGTAGCTGCGGTTGAAGCAAT
90 GAACTGGCTTAAGTCTGCTGCTAAGCTGCTGGCTGCTGAGGTCAAAGATAAGAAGACTGGAG
91 AGATTCTTCGCAAGCGTTGCGCTGTGCATTGGGTAACTCCTGATGGTTTCCCTGTGTGGCAG
92 GAATACAAGAAGCCTATTCAGACGCGCTTGAACCTGATGTTCCTCGGTCAGTTCCGCTTACA
93 GCCTACCATTAACACCAACAAAGATAGCGAGATTGATGCACACAAACAGGAGTCTGGTATCG
94 CTCCTAACTTTGTACACAGCCAAGACGGTAGCCACCTTCGTAAGACTGTAGTGTGGGCACAC
95 GAGAAGTACGGAATCGAATCTTTTGCACTGATTCACGACTCCTTCGGTACCATTCCGGCTGA
96 CGCTGCGAACCTGTTCAAAGCAGTGCGCGAAACTATGGTTGACACATATGAGTCTTGTGATG
97 TACTGGCTGATTTCTACGACCAGTTCGCTGACCAGTTGCACGAGTCTCAATTGGACAAAATG
98 CCAGCACTTCCGGCTAAAGGTAACTTGAACCTCCGTGACATCTTAGAGTCGGACTTCGCGTT
99 CGCGTAACGCCAAATCAATACGACTCACTATAGAGGGACAAACTCAAGGTCATTCGCAAGAG
100 TGGCCTTTATGATTGACCTTCTTCCGGTTAATACGACTCACTATAGGAGAACCTTAAGGTTT
101 AACTTTAAGACCCTTAAGTGTTAATTAGAGATTTAAATTAAAGAATTACTAAGAGAGGACTT
102 TAAGTATGCGTAACTTCGAAAAGATGACCAAACGTTCTAACCGTAATGCTCGTGACTTCGAG
103 GCAACCAAAGGTCGCAAGTTGAATAAGACTAAGCGTGACCGCTCTCACAAGCGTAGCTGGGA
104 GGGTCAGTAAGATGGGACGTTTATATAGTGGTAATCTGGCAGCATTCAAGGCAGCAACAAAC
105 AAGCTGTTCCAGTTAGACTTAGCGGTCATTTATGATGACTGGTATGATGCCTATACAAGAAA
106 AGATTGCATACGGTTACGTATTGAGGACAGGAGTGGAAACCTGATTGATACTAGCACCTTCT
107 ACCACCACGACGAGGACGTTCTGTTCAATATGTGTACTGATTGGTTGAACCATATGTATGAC
108 CAGTTGAAGGACTGGAAGTAATACGACTCAGTATAGGGACAATGCTTAAGGTCGCTCTCTAG
109 GAGTGGCCTTAGTCATTTAACCAATAGGAGATAAACATTATGATGAACATTAAGACTAACCC
110 GTTTAAAGCCGTGTCTTTCGTAGAGTCTGCCATTAAGAAGGCTCTGGATAACGCTGGGTATC
111 TTATCGCTGAAATCAAGTACGATGGTGTACGCGGGAACATCTGCGTAGACAATACTGCTAAC
112 AGTTACTGGCTCTCTCGTGTATCTAAAACGATTCCGGCACTGGAGCACTTAAACGGGTTTGA
113 TGTTCGCTGGAAGCGTCTACTGAACGATGACCGTTGCTTCTACAAAGATGGCTTTATGCTTG
114 ATGGGGAACTCATGGTCAAGGGCGTAGACTTTAACACAGGGTCCGGCCTACTGCGTACCAAA
115 TGGACTGACACGAAGAACCAAGAGTTCCATGAAGAGTTATTCGTTGAACCAATCCGTAAGAA
116 AGATAAAGTTCCCTTTAAGCTGCACACTGGACACCTTCACATAAAACTGTACGCTATCCTCC
117 CGCTGCACATCGTGGAGTCTGGAGAAGACTGTGATGTCATGACGTTGCTCATGCAGGAACAC
118 GTTAAGAACATGCTGCCTCTGCTACAGGAATACTTCCCTGAAATCGAATGGCAAGCGGCTGA
119 ATCTTACGAGGTCTACGATATGGTAGAACTACAGCAACTGTACGAGCAGAAGCGAGCAGAAG
120 GCCATGAGGGTCTCATTGTGAAAGACCCGATGTGTATCTATAAGCGCGGTAAGAAATCTGGC
121 TGGTGGAAAATGAAACCTGAGAACGAAGCTGACGGTATCATTCAGGGTCTGGTATGGGGTAC
122 AAAAGGTCTGGCTAATGAAGGTAAAGTGATTGGTTTTGAGGTGCTTCTTGAGAGTGGTCGTT
123 TAGTTAACGCCACGAATATCTCTCGCGCCTTAATGGATGAGTTCACTGAGACAGTAAAAGAG

124 GCCACCCTAAGTCAATGGGGATTCTTTAGCCCATACGGTATTGGCGACAACGATGCTTGTAC
125 TATTAACCCTTACGATGGCTGGGCGTGTCAAATTAGCTACATGGAGGAAACACCTGATGGCT
126 CTTTGCGGCACCCATCGTTCGTAATGTTCCGTGGCACCGAGGACAACCCTCAAGAGAAAATG
127 TAATCACACTGGCTCACCTTCGGGTGGGCCTTTCTGCGTTTATAAGGAGACACTTTATGTTT
128 AAGAAGGTTGGTAAATTCCTTGCGGCTTTGGCAGCTATCCTGACGCTTGCGTATATTCTTGC
129 GGTATACCCTCAAGTAGCACTAGTAGTAGTTGGCGCTTGTTACTTAGCGGCAGTGTGTGCTT
130 GCGTGTGGAGTATAGTTAACTGGTAATACGACTCACTAAAGGAGGTACACACCATGATGTAC
131 TTAATGCCATTACTCATCGTCATTGTAGGATGCCTTGCGCTCCACTGTAGCGATGATGATAT
132 GCCAGATGGTCACGCTTAATACGACTCACTAAAGGAGACACTATATGTTTCGACTTCATTAC
133 AACAAAAGCGTTAAGAATTTCACGGTTCGCCGTGCTGACCGTTCAATCGTATGTGCGAGCGA
134 GCGCCGAGCTAAGATACCTCTTATTGGTAACACAGTTCCTTTGGCACCGAGCGTCCACATCA
135 TTATCACCCGTGGTGACTTTGAGAAAGCAATAGACAAGAAACGTCCGGTTCTTAGTGTGGCA
136 GTGACCCGCTTCCCGTTCGTCCGTCTGTTACTCAAACGAATCAAGGAGGTGTTCTGATGGGA
137 CTGTTAGATGGTGAAGCCTGGGAAAAAGAAAACCCGCCAGTACAAGCAACTGGGTGTATAGC
138 TTGCTTAGAGAAAGATGACCGTTATCCACACACCTGTAACAAAGGAGCTAACGATATGACCG
139 AACGTGAACAAGAGATGATCATTAAGTTGATAGACAATAATGAAGGTCGCCCAGATGATTTG
140 AATGGCTGCGGTATTCTCTGCTCCAATGTCCCTTGCCACCTCTGCCCCGCAAATAACGATCA
141 AAAGATAACCTTAGGTGAAATCCGAGCGATGGACCCACGTAAACCACATCTGAATAAACCTG
142 AGGTAACTCCTACAGATGACCAGCCTTCCGCTGAGACAATCGAAGGTGTCACTAAGCCTTCC
143 CACTACATGCTGTTTGACGACATTGAGGCTATCGAAGTGATTGCTCGTTCAATGACCGTTGA
144 GCAGTTCAAGGGATACTGCTTCGGTAACATCTTAAAGTACAGACTACGTGCTGGTAAGAAGT
145 CAGAGTTAGCGTACTTAGAGAAAGACCTAGCGAAAGCAGACTTCTATAAAGAACTCTTTGAG
146 AAACATAAGGATAAATGTTATGCATAACTTCAAGTCAACCCCACCTGCCGACAGCCTATCTG
147 ATGACTTCACATCTTGCTCAGAGTGGTGCCGAAAGATGTGGGAAGAGACATTCGACGATGCG
148 TACATCAAGCTGTATGAACTTTGGAAATCGAGAGGTCAATGACTATGTCAAACGTAAATACA
149 GGTTCACTTAGTGTGGACAATAAGAAGTTTTGGGCTACCGTAGAGTCCTCGGAGCATTCCTT
150 CGAGGTTCCAATCTACGCTGAGACCCTAGACGAAGCTCTGGAGTTAGCCGAATGGCAATACG
151 TTCCGGCTGGCTTTGAGGTTACTCGTGTGCGTCCTTGTGTAGCACCGAAGTAATACGACTCA
152 CTATTAGGGAAGACTCCCTCTGAGAAACCAAACGAAACCTAAAGGAGATTAACATTATGGCT
153 AAGAAGATTTTCACCTCTGCGCTGGGTACCGCTGAACCTTACGCTTACATCGCCAAGCCGGA
154 CTACGGCAACGAAGAGCGTGGCTTTGGGAACCCTCGTGGTGTCTATAAAGTTGACCTGACTA
155 TTCCCAACAAAGACCCGCGCTGCCAGCGTATGGTCGATGAAATCGTGAAGTGTCACGAAGAG
156 GCTTATGCTGCTGCCGTTGAGGAATACGAAGCTAATCCACCTGCTGTAGCTCGTGGTAAGAA
157 ACCGCTGAAACCGTATGAGGGTGACATGCCGTTCTTCGATAACGGTGACGGTACGACTACCT
158 TTAAGTTCAAATGCTACGCGTCTTTCCAAGACAAGAAGACCAAAGAGACCAAGCACATCAAT
159 CTGGTTGTGGTTGACTCAAAAGGTAAGAAGATGGAAGACGTTCCGATTATCGGTGGTGGCTC
160 TAAGCTGAAAGTTAAATATTCTCTGGTTCCATACAAGTGGAACACTGCTGTAGGTGCGAGCG
161 TTAAGCTGCAACTGGAATCCGTGATGCTGGTCGAACTGGCTACCTTTGGTGGCGGTGAAGAC
162 GATTGGGCTGACGAAGTTGAAGAGAACGGCTATGTTGCCTCTGGTTCTGCCAAAGCGAGCAA
163 ACCACGCGACGAAGAAAGCTGGGACGAAGACGACGAAGAGTCCGAGGAAGCAGACGAAGACG
164 GAGACTTCTAAGTGGAACTGCGGGAGAAAATCCTTGAGCGAATCAAGGTGACTTCCTCTGGG
165 TGTTGGGAGTGGCAGGGCGCTACGAACAATAAAGGGTACGGGCAGGTGTGGTGCAGCAATAC
166 CGGAAAGGTTGTCTACTGTCATCGCGTAATGTCTAATGCTCCGAAAGGTTCTACCGTCCTGC
167 ACTCCTGTGATAATCCATTATGTTGTAACCCTGAACACCTATCCATAGGAACTCCAAAAGAG
168 AACTCCACTGACATGGTAAATAAGGGTCGCTCACACAAGGGGTATAAACTTTCAGACGAAGA
169 CGTAATGGCAATCATGGAGTCCAGCGAGTCCAATGTATCCTTAGCTCGCACCTATGGTGTCT
170 CCCAACAGACTATTTGTGATATACGCAAAGGGAGGCGACATGGCAGGTTACGGCGCTAAAGG
171 AATCCGAAAGGTTGGAGCGTTTCGCTCTGGCCTAGAGGACAAGGTTTCAAAGCAGTTGGAAT
172 CAAAAGGTATTAAATTCGAGTATGAAGAGTGGAAAGTGCCTTATGTAATTCCGGCGAGCAAT
173 CACACTTACACTCCAGACTTCTTACTTCCAAACGGTATATTCGTTGAGACAAAGGGTCTGTG
174 GGAAAGCGATGATAGAAAGAAGCACTTATTAATTAGGGAGCAGCACCCCGAGCTAGACATCC

175 GTATTGTCTTCTCAAGCTCACGTACTAAGTTATACAAAGGTTCTCCAACGTCTTATGGAGAG
176 TTCTGCGAAAAGCATGGTATTAAGTTCGCTGATAAACTGATACCTGCTGAGTGGATAAAGGA
177 ACCCAAGAAGGAGGTCCCCTTTGATAGATTAAAAAGGAAAGGAGGAAAGAAATAATGGCTCG
178 TGTACAGTTTAAACAACGTGAATCTACTGACGCAATCTTTGTTCACTGCTCGGCTACCAAGC
179 CAAGTCAGAATGTTGGTGTCCGTGAGATTCGCCAGTGGCACAAAGAGCAGGGTTGGCTCGAT
180 GTGGGATACCACTTTATCATCAAGCGAGACGGTACTGTGGAGGCAGGACGAGATGAGATGGC
181 TGTAGGCTCTCACGCTAAGGGTTACAACCACAACTCTATCGGCGTCTGCCTTGTTGGTGGTA
182 TCGACGATAAAGGTAAGTTCGACGCTAACTTTACGCCAGCCCAAATGCAATCCCTTCGCTCA
183 CTGCTTGTCACACTGCTGGCTAAGTACGAAGGCGCTGTGCTTCGCGCCCATCATGAGGTGGC
184 GCCGAAGGCTTGCCCTTCGTTCGACCTTAAGCGTTGGTGGGAGAAGAACGAACTGGTCACTT
185 CTGACCGTGGATAATTAATTGAACTCACTAAAGGGAGACCACAGCGGTTTCCCTTTGTTCGC
186 ATTGGAGGTCAAATAATGCGCAAGTCTTATAAACAATTCTATAAGGCTCCGAGGAGGCATAT
187 CCAAGTGTGGGAGGCAGCCAATGGGCCTATACCAAAAGGTTATTATATAGACCACATTGACG
188 GCAATCCACTCAACGACGCCTTAGACAATCTCCGTCTGGCTCTCCCAAAAGAAAACTCATGG
189 AACATGAAGACTCCAAAGAGCAATACCTCAGGACTAAAGGGACTGAGTTGGAGCAAGGAAAG
190 GGAGATGTGGAGAGGCACTGTAACAGCTGAGGGTAAACAGCATAACTTTCGTAGTAGAGATC
191 TATTGGAAGTCGTTGCGTGGATTTATAGAACTAGGAGGGAATTGCATGGACAATTCGCACGA
192 TTCCGATAGTGTATTTCTTTACCACATTCCTTGTGACAACTGTGGGAGTAGTGATGGGAACT
193 CGCTGTTCTCTGACGGACACACGTTCTGCTACGTATGCGAGAAGTGGACTGCTGGTAATGAA
194 GACACTAAAGAGAGGGCTTCAAAACGGAAACCCTCAGGAGGTAAACCAATGACTTACAACGT
195 GTGGAACTTCGGGGAATCCAATGGACGCTACTCCGCGTTAACTGCGAGAGGAATCTCCAAGG
196 AAACCTGTCAGAAGGCTGGCTACTGGATTGCCAAAGTAGACGGTGTGATGTACCAAGTGGCT
197 GACTATCGGGACCAGAACGGCAACATTGTGAGTCAGAAGGTTCGAGATAAAGATAAGAACTT
198 TAAGACCACTGGTAGTCACAAGAGTGACGCTCTGTTCGGGAAGCACTTGTGGAATGGTGGTA
199 AGAAGATTGTCGTTACAGAAGGTGAAATCGACATGCTTACCGTGATGGAACTTCAAGACTGT
200 AAGTATCCTGTAGTGTCGTTGGGTCACGGTGCCTCTGCCGCTAAGAAGACATGCGCTGCCAA
201 CTACGAATACTTTGACCAGTTCGAACAGATTATCTTAATGTTCGATATGGACGAAGCAGGGC
202 GCAAAGCAGTCGAAGAGGCTGCACAGGTTCTACCTGCTGGTAAGGTACGAGTGGCAGTTCTT
203 CCGTGTAAGGATGCAAACGAGTGTCACCTAAATGGTCACGACCGTGAAATCATGGAGCAAGT
204 GTGGAATGCTGGTCCTTGGATTCCTGATGGTGTGGTATCGGCTCTTTCGTTACGTGAACGAA
205 TCCGTGAGCACCTATCGTCCGAGGAATCAGTAGGTTTACTTTTCAGTGGCTGCACTGGTATC
206 AACGATAAGACCTTAGGTGCCCGTGGTGGTGAAGTCATTATGGTCACTTCCGGTTCCGGTAT
207 GGGTAAGTCAACGTTCGTCCGTCAACAAGCTCTACAATGGGGCACAGCGATGGGCAAGAAGG
208 TAGGCTTAGCGATGCTTGAGGAGTCCGTTGAGGAGACCGCTGAGGACCTTATAGGTCTACAC
209 AACCGTGTCCGACTGAGACAATCCGACTCACTAAAGAGAGAGATTATTGAGAACGGTAAGTT
210 CGACCAATGGTTCGATGAACTGTTCGGCAACGATACGTTCCATCTATATGACTCATTCGCCG
211 AGGCTGAGACGGATAGACTGCTCGCTAAGCTGGCCTACATGCGCTCAGGCTTGGGCTGTGAC
212 GTAATCATTCTAGACCACATCTCAATCGTCGTATCCGCTTCTGGTGAATCCGATGAGCGTAA
213 GATGATTGACAACCTGATGACCAAGCTCAAAGGGTTCGCTAAGTCAACTGGGGTGGTGCTGG
214 TCGTAATTTGTCACCTTAAGAACCCAGACAAAGGTAAAGCACATGAGGAAGGTCGCCCCGTT
215 TCTATTACTGACCTACGTGGTTCTGGCGCACTACGCCAACTATCTGATACTATTATTGCCCT
216 TGAGCGTAATCAGCAAGGCGATATGCCTAACCTTGTCCTCGTTCGTATTCTCAAGTGCCGCT
217 TTACTGGTGATACTGGTATCGCTGGCTACATGGAATACAACAAGGAAACCGGATGGCTTGAA
218 CCATCAAGTTACTCAGGGGAAGAAGAGTCACACTCAGAGTCAACAGACTGGTCCAACGACAC
219 TGACTTCTGACAGGATTCTTGATGACTTTCCAGACGACTACGAGAAGTTTCGCTGGAGAGTC
220 CCATTCTAATACGACTCACTAAAGGAGACACACCATGTTCAAACTGATTAAGAAGTTAGGCC
221 AACTGCTGGTTCGTATGTACAACGTGGAAGCCAAGCGACTGAACGATGAGGCTCGTAAAGAG
222 GCCACACAGTCACGCGCTCTGGCGATTCGCTCCAACGAACTGGCTGACAGTGCATCCACTAA
223 AGTTACCGAGGCTGCCCGTGTGGCAAACCAAGCTCAACAGCTTTCCAAATTCTTTGAGTAAT
224 CAAACAGGAGAAACCATTATGTCTAACGTAGCTGAAACTATCCGTCTATCCGATACAGCTGA
225 CCAGTGGAACCGTCGAGTCCACATCAACGTTCGCAACGGTAAGGCGACTATGGTTTACCGCT

226 GGAAGGACTCTAAGTCCTCTAAGAATCACACTCAGCGTATGACGTTGACAGATGAGCAAGCA
227 CTGCGTCTGGTCAATGCGCTTACCAAAGCTGCCGTGACAGCAATTCATGAAGCTGGTCGCGT
228 CAATGAAGCTATGGCTATCCTCGACAAGATTGATAACTAAGAGTGGTATCCTCAAGGTCGCC
229 AAAGTGGTGGCCTTCATGAATACTATTCGACTCACTATAGGAGATATTACCATGCGTGACCC
230 TAAAGTTATCCAAGCAGAAATCGCTAAACTGGAAGCTGAACTGGAGGACGTTAAGTACCATG
231 AAGCTAAGACTCGCTCCGCTGTTCACATCTTGAAGAACTTAGGCTGGACTTGGACAAGACAG
232 ACTGGCTGGAAGAAACCAGAAGTTACCAAGCTGAGTCATAAGGTGTTCGATAAGGACACTAT
233 GACCCACATCAAGGCTGGTGATTGGGTTAAGGTTGACATGGGAGTTGTTGGTGGATACGGCT
234 ACGTCCGCTCAGTTAGTGGCAAATATGCACAAGTGTCATACATCACAGGTGTTACTCCACGC
235 GGTGCAATCGTTGCCGATAAGACCAACATGATTCACACAGGTTTCTTGACAGTTGTTTCATA
236 TGAAGAGATTGTTAAGTCACGATAATCAATAGGAGAAATCAATATGAGCGATAAAATTATTC
237 ACCTGACTATAGGAGAAATCAATATGAGCGATAAAATTATTCACCTGACTGACGACAGTTTT
238 GACACGGATGTACTCAAAGCGGACGGGGCGATCCTCGTCGATTTCTGGGCAGAGTGGTGCGG
239 TCCGTGCAAAATGATCGCCCCGATTCTGGATGAAATCGCTGACGAATATCAGGGCAAACTGA
240 CCGTTGCAAAACTGAACATCGATCAAAACCCTGGCACTGCGCCGAAATATGGCATCCGTGGT
241 ATCCCGACTCTGCTGCTGTTCAAAAACGGTGAAGTGGCGGCAACCAAAGTGGGTGCACTGTC
242 TAAAGGTCAGTTGAAAGAGTTCCTCGACGCTAACCTGGCGTAATACAGGAGGCTACTCATGA
243 ACGAAAGACACTTAACAGGTGCTGCTTCTGAAATGCTAGTAGCCTACAAATTTACCAAAGCT
244 GGGTACACTGTCTATTACCCTATGCTGACTCAGAGTAAAGAGGACTTGGTTGTATGTAAGGA
245 TGGTAAATTTAGTAAGGTTCAGGTTAAAACAGCCACAACGGTTCAAACCAACACAGGAGATG
246 CCAAGCAGGTTAGGCTAGGTGGATGCGGTAGGTCCGAATATAAGGATGGAGACTTTGACATT
247 CTTGCGGTTGTGGTTGACGAAGATGTGCTTATTTTCACATGGGACGAAGTAAAAGGTAAGAC
248 ATCCATGTGTGTCGGCAAGAGAAACAAAGGCATAAAACTATAGGAGAAATTATTATGGCTAT
249 GACAAAGAAATTTAAAGTGTCCTTCGACGTTACCGCAAAGATGTCGTCTGACGTTCAGGCAA
250 TCTTAGAGAAAGATATGCTGCATCTATGTAAGCAGGTCGGCTCAGGTGCGATTGTCCCCAAT
251 GGTAAACAGAAGGAAATGATTGTCCAGTTCCTGACACACGGTATGGAAGGATTGATGACATT
252 CGTAGTACGTACATCATTTCGTGAGGCCATTAAGGACATGCACGAAGAGTATGCAGATAAGG
253 ACTCTTTCAAACAATCTCCTGCAACAGTACGGGAGGTGTTCTGATGTCTGACTACCTGAAAG
254 TGCTGCAAGCAATCAAAAGTTGCCCTAAGACTTTCCAGTCCAACTATGTACGGAACAATGCG
255 AGCCTCGTAGCGGAGGCCGCTTCCCGTGGTCACATCTCGTGCCTGACTACTAGTGGACGTAA
256 CGGTGGCGCTTGGGAAATCACTGCTTCCGGTACTCGCTTTCTGAAACGAATGGGAGGATGTG
257 TCTAATGTCTCGTGACCTTGTGACTATTCCACGCGATGTGTGGAACGATATACAGGGCTACA
258 TCGACTCTCTGGAACGTGAGAACGATAGCCTTAAGAATCAACTAATGGAAGCTGACGAATAC
259 GTAGCGGAACTAGAGGAGAAACTTAATGGCACTTCTTGACCTTAAACAATTCTATGAGTTAC
260 GTGAAGGCTGCGACGACAAGGGTATCCTTGTGATGGACGGCGACTGGCTGGTCTTCCAAGCT
261 ATGAGTGCTGCTGAGTTTGATGCCTCTTGGGAGGAAGAGATTTGGCACCGATGCTGTGACCA
262 CGCTAAGGCCCGTCAGATTCTTGAGGATTCCATTAAGTCCTACGAGACCCGTAAGAAGGCTT
263 GGGCAGGTGCTCCAATTGTCCTTGCGTTCACCGATAGTGTTAACTGGCGTAAAGAACTGGTT
264 GACCCGAACTATAAGGCTAACCGTAAGGCCGTGAAGAAACCTGTAGGGTACTTTGAGTTCCT
265 TGATGCTCTCTTTGAGCGCGAAGAGTTCTATTGCATCCGTGAGCCTATGCTTGAGGGTGATG
266 ACGTTATGGGAGTTATTGCTTCCAATCCGTCTGCCTTCGGTGCTCGTAAGGCTGTAATCATC
267 TCTTGCGATAAGGACTTTAAGACCATCCCTAACTGTGACTTCCTGTGGTGTACCACTGGTAA
268 CATCCTGACTCAGACCGAAGAGTCCGCTGACTGGTGGCACCTCTTCCAGACCATCAAGGGTG
269 ACATCACTGATGGTTACTCAGGGATTGCTGGATGGGGTGATACCGCCGAGGACTTCTTGAAT
270 AACCCGTTCATAACCGAGCCTAAAACGTCTGTGCTTAAGTCCGGTAAGAACAAAGGCCAAGA
271 GGTTACTAAATGGGTTAAACGCGACCCTGAGCCTCATGAGACGCTTTGGGACTGCATTAAGT
272 CCATTGGCGCGAAGGCTGGTATGACCGAAGAGGATATTATCAAGCAGGGCCAAATGGCTCGA
273 ATCCTACGGTTCAACGAGTACAACTTTATTGACAAGGAGATTTACCTGTGGAGACCGTAGCG
274 TATATTGGTCTGGGTCTTTGTGTTCTCGGAGTGTGCCTCATTTCGTGGGGCCTTTGGGACTT
275 AGCCAGAATAATCAAGTCGTTACACGACACTAAGTGATAAACTCAAGGTCCCTAAATTAATA
276 CGACTCACTATAGGGAGATAGGGGCCTTTACGATTATTACTTTAAGATTTAACTCTAAGAGG

277 AATCTTTATTATGTTAACACCTATTAACCAATTACTTAAGAACCCTAACGATATTCCAGATG
278 TACCTCGTGCAACCGCTGAGTATCTACAGGTTCGATTCAACTATGCGTACCTCGAAGCGTCT
279 GGTCATATAGGACTTATGCGTGCTAATGGTTGTAGTGAGGCCCACATCTTGGGTTTCATTCA
280 GGGCCTACAGTATGCCTCTAACGTCATTGACGAGATTGAGTTACGCAAGGAACAACTAAGAG
281 ATGATGGGGAGGATTGACACTATGTGTTTCTCACCGAAAATTAAAACTCCGAAGATGGATAC
282 CAATCAGATTCGAGCCGTTGAGCCAGCGCCTCTGACCCAAGAAGTGTCAAGCGTGGAGTTCG
283 GTGGGTCTTCTGATGAGACGGATACCGAGGGCACCGAAGTGTCTGGACGCAAAGGCCTCAAG
284 GTCGAACGTGATGATTCCGTAGCGAAGTCTAAAGCCAGCGGCAATGGCTCCGCTCGTATGAA
285 ATCTTCCATCCGTAAGTCCGCATTTGGAGGTAAGAAGTGATGTCTGAGTTCACATGTGTGGA
286 GGCTAAGAGTCGCTTCCGTGCAATCCGGTGGACTGTGGAACACCTTGGGTTGCCTAAAGGAT
287 TCGAAGGACACTTTGTGGGCTACAGCCTCTACGTAGACGAAGTGATGGACATGTCTGGTTGC
288 CGTGAAGAGTACATTCTGGACTCTACCGGAAAACATGTAGCGTACTTCGCGTGGTGCGTAAG
289 CTGTGACATTCACCACAAAGGAGACATTCTGGATGTAACGTCCGTTGTCATTAATCCTGAGG
290 CAGACTCTAAGGGCTTACAGCGATTCCTAGCGAAACGCTTTAAGTACCTTGCGGAACTCCAC
291 GATTGCGATTGGGTGTCTCGTTGTAAGCATGAAGGCGAGACAATGCGTGTATACTTTAAGGA
292 GGTATAAGTTATGGGTAAGAAAGTTAAGAAGGCCGTGAAGAAAGTCACCAAGTCCGTTAAGA
293 AAGTCGTTAAGGAAGGGGCTCGTCCGGTTAAACAGGTTGCTGGCGGTCTAGCTGGTCTGGCT
294 GGTGGTACTGGTGAAGCACAGATGGTGGAAGTACCACAAGCTGCCGCACAGATTGTTGACGT
295 ACCTGAGAAAGAGGTTTCCACTGAGGACGAAGCACAGACAGAAAGCGGACGCAAGAAAGCTC
296 GTGCTGGCGGTAAGAAATCCTTGAGTGTAGCCCGTAGCTCCGGTGGCGGTATCAACATTTAA
297 TCAGGAGGTTATCGTGGAAGACTGCATTGAATGGACCGGAGGTGTCAACTCTAAGGGTTATG
298 GTCGTAAGTGGGTTAATGGTAAACTTGTGACTCCACATAGGCACATCTATGAGGAGACATAT
299 GGTCCAGTTCCAACAGGAATTGTGGTGATGCATATCTGCGATAACCCTAGGTGCTATAACAT
300 AAAGCACCTTACGCTTGGAACTCCAAAGGATAATTCCGAGGACATGGTTACCAAAGGTAGAC
301 AGGCTAAAGGAGAGGAACTAAGCAAGAAACTTACAGAGTCAGACGTTCTCGCTATACGCTCT
302 TCAACCTTAAGCCACCGCTCCTTAGGAGAACTGTATGGAGTCAGTCAATCAACCATAACGCG
303 AATACTACAGCGTAAGACATGGAGACACATTTAATGGCTGAGAAACGAACAGGACTTGCGGA
304 GGATGGCGCAAAGTCTGTCTATGAGCGTTTAAAGAACGACCGTGCTCCCTATGAGACACGCG
305 CTCAGAATTGCGCTCAATATACCATCCCATCATTGTTCCCTAAGGACTCCGATAACGCCTCT
306 ACAGATTATCAAACTCCGTGGCAAGCCGTGGGCGCTCGTGGTCTGAACAATCTAGCCTCTAA
307 GCTCATGCTGGCTCTATTCCCTATGCAGACTTGGATGCGACTTACTATATCTGAATATGAAG
308 CAAAGCAGTTACTGAGCGACCCCGATGGACTCGCTAAGGTCGATGAGGGCCTCTCGATGGTA
309 GAGCGTATCATCATGAACTACATTGAGTCTAACAGTTACCGCGTGACTCTCTTTGAGGCTCT
310 CAAACAGTTAGTCGTAGCTGGTAACGTCCTGCTGTACCTACCGGAACCGGAAGGGTCAAACT
311 ATAATCCCATGAAGCTGTACCGATTGTCTTCTTATGTGGTCCAACGAGACGCATTCGGCAAC
312 GTTCTGCAAATGGTGACTCGTGACCAGATAGCTTTTGGTGCTCTCCCTGAGGACATCCGTAA
313 GGCTGTAGAAGGTCAAGGTGGTGAGAAGAAAGCTGATGAGACAATCGACGTGTACACTCACA
314 TCTATCTGGATGAGGACTCAGGTGAATACCTCCGATACGAAGAGGTCGAGGGTATGGAAGTC
315 CAAGGCTCCGATGGGACTTATCCTAAAGAGGCTTGCCCATACATCCCGATTCGGATGGTCAG
316 ACTAGATGGTGAATCCTACGGTCGTTCGTACATTGAGGAATACTTAGGTGACTTACGGTCCC
317 TTGAAAATCTCCAAGAGGCTATCGTCAAGATGTCCATGATTAGCTCTAAGGTTATCGGCTTA
318 GTGAATCCTGCTGGTATCACCCAGCCACGCCGACTGACCAAAGCTCAGACTGGTGACTTCGT
319 TACTGGTCGTCCAGAAGACATCTCGTTCCTCCAACTGGAGAAGCAAGCAGACTTTACTGTAG
320 CTAAAGCCGTAAGTGACGCTATCGAGGCTCGCCTTTCGTTTGCCTTTATGTTGAACTCTGCG
321 GTTCAGCGTACAGGTGAACGTGTGACCGCCGAAGAGATTCGGTATGTAGCTTCTGAACTTGA
322 AGATACTTTAGGTGGTGTCTACTCTATCCTTTCTCAAGAATTACAATTGCCTCTGGTACGAG
323 TGCTCTTGAAGCAACTACAAGCCACGCAACAGATTCCTGAGTTACCTAAGGAAGCCGTAGAG
324 CCAACCATTAGTACAGGTCTGGAAGCAATTGGTCGAGGACAAGACCTTGATAAGCTGGAGCG
325 GTGTGTCACTGCGTGGGCTGCACTGGCACCTATGCGGGACGACCCTGATATTAACCTTGCGA
326 TGATTAAGTTACGTATTGCCAACGCTATCGGTATTGACACTTCTGGTATTCTACTCACCGAA
327 GAACAGAAGCAACAGAAGATGGCCCAACAGTCTATGCAAATGGGTATGGATAATGGTGCTGC

TGCGCTGGCTCAAGGTATGGCTGCACAAGCTACAGCTTCACCTGAGGCTATGGCTGCTGCCG
CTGATTCCGTAGGTTTACAGCCGGGAATTTAATACGACTCACTATAGGGAGACCTCATCTTT
GAAATGAGCGATGACAAGAGGTTGGAGTCCTCGGTCTTCCTGTAGTTCAACTTTAAGGAGAC
AATAATAATGGCTGAATCTAATGCAGACGTATATGCATCTTTTGGCGTGAACTCCGCTGTGA
TGTCTGGTGGTTCCGTTGAGGAACATGAGCAGAACATGCTGGCTCTTGATGTTGCTGCCCGT
GATGGCGATGATGCAATCGAGTTAGCGTCAGACGAAGTGGAAACAGAACGTGACCTGTATGA
CAACTCTGACCCGTTCGGTCAAGAGGATGACGAAGGCCGCATTCAGGTTCGTATCGGTGATG
GCTCTGAGCCGACCGATGTGGACACTGGAGAAGAAGGCGTTGAGGGCACCGAAGGTTCCGAA
GAGTTTACCCCACTGGGCGAGACTCCAGAAGAACTGGTAGCTGCCTCTGAGCAACTTGGTGA
GCACGAAGAGGGCTTCCAAGAGATGATTAACATTGCTGCTGAGCGTGGCATGAGTGTCGAGA
CCATTGAGGCTATCCAGCGTGAGTACGAGGAGAACGAAGAGTTGTCCGCCGAGTCCTACGCT
AAGCTGGCTGAAATTGGCTACACGAAGGCTTTCATTGACTCGTATATCCGTGGTCAAGAAGC
TCTGGTGGAGCAGTACGTAAACAGTGTCATTGAGTACGCTGGTGGTCGTGAACGTTTTGATG
CACTGTATAACCACCTTGAGACGCACAACCCTGAGGCTGCACAGTCGCTGGATAATGCGTTG
ACCAATCGTGACTTAGCGACCGTTAAGGCTATCATCAACTTGGCTGGTGAGTCTCGCGCTAA
GGCGTTCGGTCGTAAGCCAACTCGTAGTGTGACTAATCGTGCTATTCCGGCTAAACCTCAGG
CTACCAAGCGTGAAGGCTTTGCGGACCGTAGCGAGATGATTAAAGCTATGAGTGACCCTCGG
TATCGCACAGATGCCAACTATCGTCGTCAAGTCGAACAGAAAGTAATCGATTCGAACTTCTG
ATAGACTTCGAAATTAATACGACTCACTATAGGGAGACCACAACGGTTTCCCTCTAGAAATA
ATTTTGTTTAACTTTAAGAAGGAGATATACATATGGCTAGCATGACTGGTGGACAGCAAATG
GGTACTAACCAAGGTAAAGGTGTAGTTGCTGCTGGAGATAAACTGGCGTTGTTCTTGAAGGT
ATTTGGCGGTGAAGTCCTGACTGCGTTCGCTCGTACCTCCGTGACCACTTCTCGCCACATGG
TACGTTCCATCTCCAGCGGTAAATCCGCTCAGTTCCCTGTTCTGGGTCGCACTCAGGCAGCG
TATCTGGCTCCGGGCGAGAACCTCGACGATAAACGTAAGGACATCAAACACACCGAGAAGGT
AATCACCATTGACGGTCTCCTGACGGCTGACGTTCTGATTTATGATATTGAGGACGCGATGA
ACCACTACGACGTTCGCTCTGAGTATACCTCTCAGTTGGGTGAATCTCTGGCGATGGCTGCG
GATGGTGCGGTTCTGGCTGAGATTGCCGGTCTGTGTAACGTGGAAAGCAAATATAATGAGAA
CATCGAGGGCTTAGGTACTGCTACCGTAATTGAGACCACTCAGAACAAGGCCGCACTTACCG
ACCAAGTTGCGCTGGGTAAGGAGATTATTGCGGCTCTGACTAAGGCTCGTGCGGCTCTGACC
AAGAACTATGTTCCGGCTGCTGACCGTGTGTTCTACTGTGACCCAGATAGCTACTCTGCGAT
TCTGGCAGCACTGATGCCGAACGCAGCAAACTACGCTGCTCTGATTGACCCTGAGAAGGGTT
CTATCCGCAACGTTATGGGCTTTGAGGTTGTAGAAGTTCCGCACCTCACCGCTGGTGGTGCT
GGTACCGCTCGTGAGGGCACTACTGGTCAGAAGCACGTCTTCCCTGCCAATAAAGGTGAGGG
TAATGTCAAGGTTGCTAAGGACAACGTTATCGGCCTGTTCATGCACCGCTCTGCGGTAGGTA
CTGTTAAGCTGCGTGACTTGGCTCTGGAGCGCGCTCGCCGTGCTAACTTCCAAGCGGACCAG
ATTATCGCTAAGTACGCAATGGGCCACGGTGGTCTTCGCCCAGAAGCTGCTGGTGCAGTGGT
TTTCAAAGTGGAGTAATGCTGGGGGTGGCCTCAACGGTCGCTGCTAGTCCCGAAGAGGCGAG
TGTTACTTCAACAGAAGAAACCTTAACGCCAGCACAGGAGGCCGCACGCACCCGCGCTGCTA
ACAAAGCCCGAAAGGAAGCTGAGTTGGCTGCTGCCACCGCTGAGCAATAACTAGCATAACCC
CTTGGGGCCTCTAAACGGGTCTTGAGGGGTTTTTTGCTGAAAGGAGGAACTATATGCGCTCA
TACGATATGAACGTTGAGACTGCCGCTGAGTTATCAGCTGTGAACGACATTCTGGCGTCTAT
CGGTGAACCTCCGGTATCAACGCTGGAAGGTGACGCTAACGCAGATGCAGCGAACGCTCGGC
GTATTCTCAACAAGATTAACCGACAGATTCAATCTCGTGGATGGACGTTCAACATTGAGGAA
GGCATAACGCTACTACCTGATGTTTACTCCAACCTGATTGTATACAGTGACGACTATTTATC
CCTAATGTCTACTTCCGGTCAATCCATCTACGTTAACCGAGGTGGCTATGTGTATGACCGAA
CGAGTCAATCAGACCGCTTTGACTCTGGTATTACTGTGAACATTATTCGTCTCCGCGACTAC
GATGAGATGCCTGAGTGCTTCCGTTACTGGATTGTCACCAAGGCTTCCCGTCAGTTCAACAA
CCGATTCTTTGGGGCACCGGAAGTAGAGGGTGTACTCCAAGAAGAGGAAGATGAGGCTAGAC
GTCTCTGCATGGAGTATGAGATGGACTACGGTGGGTACAATATGCTGGATGGAGATGCGTTC
ACTTCTGGTCTACTGACTCGCTAACATTAATAAATAAGGAGGCTCTAATGGCACTCATTAGC
CAATCAATCAAGAACTTGAAGGGTGGTATCAGCCAACAGCCTGACATCCTTCGTTATCCAGA

379 CCAAGGGTCACGCCAAGTTAACGGTTGGTCTTCGGAGACCGAGGGCCTCCAAAAGCGTCCAC
380 CTCTTGTTTTCTTAAATACACTTGGAGACAACGGTGCGTTAGGTCAAGCTCCGTACATCCAC
381 CTGATTAACCGAGATGAGCACGAACAGTATTACGCTGTGTTCACTGGTAGCGGAATCCGAGT
382 GTTCGACCTTTCTGGTAACGAGAAGCAAGTTAGGTATCCTAACGGTTCCAACTACATCAAGA
383 CCGCTAATCCACGTAACGACCTGCGAATGGTTACTGTAGCAGACTATACGTTCATCGTTAAC
384 CGTAACGTTGTTGCACAGAAGAACACAAAGTCTGTCAACTTACCGAATTACAACCCTAATCA
385 AGACGGATTGATTAACGTTCGTGGTGGTCAGTATGGTAGGGAACTAATTGTACACATTAACG
386 GTAAAGACGTTGCGAAGTATAAGATACCAGATGGTAGTCAACCTGAACACGTAAACAATACG
387 GATGCCCAATGGTTAGCTGAAGAGTTAGCCAAGCAGATGCGCACTAACTTGTCTGATTGGAC
388 TGTAAATGTAGGGCAAGGGTTCATCCATGTGACCGCACCTAGTGGTCAACAGATTGACTCCT
389 TCACGACTAAAGATGGCTACGCAGACCAGTTGATTAACCCTGTGACCCACTACGCTCAGTCG
390 TTCTCTAAGCTGCCACCTAATGCTCCTAACGGCTACATGGTGAAAATCGTAGGGGACGCCTC
391 TAAGTCTGCCGACCAGTATTACGTTCGGTATGACGCTGAGCGGAAAGTTTGGACTGAGACTT
392 TAGGTTGGAACACTGAGGACCAAGTTCTATGGGAAACCATGCCACACGCTCTTGTGCGAGCC
393 GCTGACGGTAATTTCGACTTCAAGTGGCTTGAGTGGTCTCCTAAGTCTTGTGGTGACGTTGA
394 CACCAACCCTTGGCCTTCTTTTGTTGGTTCAAGTATTAACGATGTGTTCTTCTTCCGTAACC
395 GCTTAGGATTCCTTAGTGGGGAGAACATCATATTGAGTCGTACAGCCAAATACTTCAACTTC
396 TACCCTGCGTCCATTGCGAACCTTAGTGATGACGACCCTATAGACGTAGCTGTGAGTACCAA
397 CCGAATAGCAATCCTTAAGTACGCCGTTCCGTTCTCAGAAGAGTTACTCATCTGGTCCGATG
398 AAGCACAATTCGTCCTGACTGCCTCGGGTACTCTCACATCTAAGTCGGTTGAGTTGAACCTA
399 ACGACCCAGTTTGACGTACAGGACCGAGCGAGACCTTTTGGGATTGGGCGTAATGTCTACTT
400 TGCTAGTCCGAGGTCCAGCTTCACGTCCATCCACAGGTACTACGCTGTGCAGGATGTCAGTT
401 CCGTTAAGAATGCTGAGGACATTACATCACACGTTCCTAACTACATCCCTAATGGTGTGTTC
402 AGTATTTGCGGAAGTGGTACGGAAAACTTCTGTTCGGTACTATCTCACGGGGACCCTAGTAA
403 AATCTTCATGTACAAATTCCTGTACCTGAACGAAGAGTTAAGGCAACAGTCGTGGTCTCATT
404 GGGACTTTGGGGAAAACGTACAGGTTCTAGCTTGTCAGAGTATCAGCTCAGATATGTATGTG
405 ATTCTTCGCAATGAGTTCAATACGTTCCTAGCTAGAATCTCTTTCACTAAGAACGCCATTGA
406 CTTACAGGGAGAACCCTATCGTGCCTTTATGGACATGAAGATTCGATACACGATTCCTAGTG
407 GAACATACAACGATGACACATTCACTACCTCTATTCATATTCCAACAATTTATGGTGCAAAC
408 TTCGGGAGGGGCAAAATCACTGTATTGGAGCCTGATGGTAAGATAACCGTGTTTGAGCAACC
409 TACGGCTGGGTGGAATAGCGACCCTTGGCTGAGACTCAGCGGTAACTTGGAGGGACGCATGG
410 TGTACATTGGGTTCAACATTAACTTCGTATATGAGTTCTCTAAGTTCCTCATCAAGCAGACT
411 GCCGACGACGGGTCTACCTCCACGGAAGACATTGGGCGCTTACAGTTACGCCGAGCGTGGGT
412 TAACTACGAGAACTCTGGTACGTTTGACATTTATGTTGAGAACCAATCGTCTAACTGGAAGT
413 ACACAATGGCTGGTGCCCGATTAGGCTCTAACACTCTGAGGGCTGGGAGACTGAACTTAGGG
414 ACCGGACAATATCGATTCCCTGTGGTTGGTAACGCCAAGTTCAACACTGTATACATCTTGTC
415 AGATGAGACTACCCCTCTGAACATCATTGGGTGTGGCTGGGAAGGTAACTACTTACGGAGAA
416 GTTCCGGTATTTAATTAAATATTCTCCCTGTGGTGGCTCGAAATTAATACGACTCACTATAG
417 GGAGAACAATACGACTACGGGAGGGTTTTCTTATGATGACTATAAGACCTACTAAAAGTACA
418 GACTTTGAGGTATTCACTCCGGCTCACCATGACATTCTTGAAGCTAAGGCTGCTGGTATTGA
419 GCCGAGTTTCCCTGATGCTTCCGAGTGTGTCACGTTGAGCCTCTATGGGTTCCCTCTAGCTA
420 TCGGTGGTAACTGCGGGGACCAGTGCTGGTTCGTTACGAGCGACCAAGTGTGGCGACTTAGT
421 GGAAAGGCTAAGCGAAAGTTCCGTAAGTTAATCATGGAGTATCGCGATAAGATGCTTGAGAA
422 GTATGATACTCTTTGGAATTACGTATGGGTAGGCAATACGTCCCACATTCGTTTCCTCAAGA
423 CTATCGGTGCGGTATTCCATGAAGAGTACACACGAGATGGTCAATTTCAGTTATTTACAATC
424 ACGAAAGGAGGATAACCATATGTGTTGGGCAGCCGCAATACCTATCGCTATATCTGGCGCTC
425 AGGCTATCAGTGGTCAGAACGCTCAGGCCAAAATGATTGCCGCTCAGACCGCTGCTGGTCGT
426 CGTCAAGCTATGGAAATCATGAGGCAGACGAACATCCAGAATGCTGACCTATCGTTGCAAGC
427 TCGAAGTAAACTTGAGGAAGCGTCCGCCGAGTTGACCTCACAGAACATGCAGAAGGTCCAAG
428 CTATTGGGTCTATCCGAGCGGCTATCGGAGAGAGTATGCTTGAAGGTTCCTCAATGGACCGC
429 ATTAAGCGAGTCACAGAAGGACAGTTCATTCGGGAAGCCAATATGGTAACTGAGAACTATCG

430 CCGTGACTACCAAGCAATCTTCGCACAGCAACTTGGTGGTACTCAAAGTGCTGCAAGTCAGA
431 TTGACGAAATCTATAAGAGCGAACAGAAACAGAAGAGTAAGCTACAGATGGTTCTGGACCCA
432 CTGGCTATCATGGGGTCTTCCGCTGCGAGTGCTTACGCATCCGGTGCGTTCGACTCTAAGTC
433 CACAACTAAGGCACCTATTGTTGCCGCTAAAGGAACCAAGACGGGGAGGTAATGAGCTATGA
434 GTAAAATTGAATCTGCCCTTCAAGCGGCACAACCGGGACTCTCTCGGTTACGTGGTGGTGCT
435 GGAGGTATGGGCTATCGTGCAGCAACCACTCAGGCCGAACAGCCAAGGTCAAGCCTATTGGA
436 CACCATTGGTCGGTTCGCTAAGGCTGGTGCCGATATGTATACCGCTAAGGAACAACGAGCAC
437 GAGACCTAGCTGATGAACGCTCTAACGAGATTATCCGTAAGCTGACCCCTGAGCAACGTCGA
438 GAAGCTCTCAACAACGGGACCCTTCTGTATCAGGATGACCCATACGCTATGGAAGCACTCCG
439 AGTCAAGACTGGTCGTAACGCTGCGTATCTTGTGGACGATGACGTTATGCAGAAGATAAAAG
440 AGGGTGTCTTCCGTACTCGCGAAGAGATGGAAGAGTATCGCCATAGTCGCCTTCAAGAGGGC
441 GCTAAGGTATACGCTGAGCAGTTCGGCATCGACCCTGAGGACGTTGATTATCAGCGTGGTTT
442 CAACGGGGACATTACCGAGCGTAACATCTCGCTGTATGGTGCGCATGATAACTTCTTGAGCC
443 AGCAAGCTCAGAAGGGCGCTATCATGAACAGCCGAGTGGAACTCAACGGTGTCCTTCAAGAC
444 CCTGATATGCTGCGTCGTCCAGACTCTGCTGACTTCTTTGAGAAGTATATCGACAACGGTCT
445 GGTTACTGGCGCAATCCCATCTGATGCTCAAGCCACACAGCTTATAAGCCAAGCGTTCAGTG
446 ACGCTTCTAGCCGTGCTGGTGGTGCTGACTTCCTGATGCGAGTCGGTGACAAGAAGGTAACA
447 CTTAACGGAGCCACTACGACTTACCGAGAGTTGATTGGTGAGGAACAGTGGAACGCTCTCAT
448 GGTCACAGCACAACGTTCTCAGTTTGAGACTGACGCGAAGCTGAACGAGCAGTATCGCTTGA
449 AGATTAACTCTGCGCTGAACCAAGAGGACCCAAGGACAGCTTGGGAGATGCTTCAAGGTATC
450 AAGGCTGAACTAGATAAGGTCCAACCTGATGAGCAGATGACACCACAACGTGAGTGGCTAAT
451 CTCCGCACAGGAACAAGTTCAGAATCAGATGAACGCATGGACGAAAGCTCAGGCCAAGGCTC
452 TGGACGATTCCATGAAGTCAATGAACAAACTTGACGTAATCGACAAGCAATTCCAGAAGCGA
453 ATCAACGGTGAGTGGGTCTCAACGGATTTTAAGGATATGCCAGTCAACGAGAACACTGGTGA
454 GTTCAAGCATAGCGATATGGTTAACTACGCCAATAAGAAGCTCGCTGAGATTGACAGTATGG
455 ACATTCCAGACGGTGCCAAGGATGCTATGAAGTTGAAGTACCTTCAAGCGGACTCTAAGGAC
456 GGAGCATTCCGTACAGCCATCGGAACCATGGTCACTGACGCTGGTCAAGAGTGGTCTGCCGC
457 TGTGATTAACGGTAAGTTACCAGAACGAACCCCAGCTATGGATGCTCTGCGCAGAATCCGCA
458 ATGCTGACCCTCAGTTGATTGCTGCGCTATACCCAGACCAAGCTGAGCTATTCCTGACGATG
459 GACATGATGGACAAGCAGGGTATTGACCCTCAGGTTATTCTTGATGCCGACCGACTGACTGT
460 TAAGCGGTCCAAAGAGCAACGCTTTGAGGATGATAAAGCATTCGAGTCTGCACTGAATGCAT
461 CTAAGGCTCCTGAGATTGCCCGTATGCCAGCGTCACTGCGCGAATCTGCACGTAAGATTTAT
462 GACTCCGTTAAGTATCGCTCGGGGAACGAAAGCATGGCTATGGAGCAGATGACCAAGTTCCT
463 TAAGGAATCTACCTACACGTTCACTGGTGATGATGTTGACGGTGATACCGTTGGTGTGATTC
464 CTAAGAATATGATGCAGGTTAACTCTGACCCGAAATCATGGGAGCAAGGTCGGGATATTCTG
465 GAGGAAGCACGTAAGGGAATCATTGCGAGCAACCCTTGGATAACCAATAAGCAACTGACCAT
466 GTATTCTCAAGGTGACTCCATTTACCTTATGGACACCACAGGTCAAGTCAGAGTCCGATACG
467 ACAAAGAGTTACTCTCGAAGGTCTGGAGTGAGAACCAGAAGAAACTCGAAGAGAAAGCTCGT
468 GAGAAGGCTCTGGCTGATGTGAACAAGCGAGCACCTATAGTTGCCGCTACGAAGGCCCGTGA
469 AGCTGCTGCTAAACGAGTCCGAGAGAAACGTAAACAGACTCCTAAGTTCATCTACGGACGTA
470 AGGAGTAACTAAAGGCTACATAAGGAGGCCCTAAATGGATAAGTACGATAAGAACGTACCAA
471 GTGATTATGATGGTCTGTTCCAAAAGGCTGCTGATGCCAACGGGGTCTCTTATGACCTTTTA
472 CGTAAAGTCGCTTGGACAGAATCACGATTTGTGCCTACAGCAAAATCTAAGACTGGACCATT
473 AGGCATGATGCAATTTACCAAGGCAACCGCTAAGGCCCTCGGTCTGCGAGTTACCGATGGTC
474 CAGACGACGACCGACTGAACCCTGAGTTAGCTATTAATGCTGCCGCTAAGCAACTTGCAGGT
475 CTGGTAGGGAAGTTTGATGGCGATGAACTCAAAGCTGCCCTTGCGTACAACCAAGGCGAGGG
476 ACGCTTGGGTAATCCACAACTTGAGGCGTACTCTAAGGGAGACTTCGCATCAATCTCTGAGG
477 AGGGACGTAACTACATGCGTAACCTTCTGGATGTTGCTAAGTCACCTATGGCTGGACAGTTG
478 GAAACTTTTGGTGGCATAACCCCAAAGGGTAAAGGCATTCCGGCTGAGGTAGGATTGGCTGG
479 AATTGGTCACAAGCAGAAAGTAACACAGGAACTTCCTGAGTCCACAAGTTTTGACGTTAAGG
480 GTATCGAACAGGAGGCTACGGCGAAACCATTCGCCAAGGACTTTTGGGAGACCCACGGAGAA

481 ACACTTGACGAGTACAACAGTCGTTCAACCTTCTTCGGATTCAAAAATGCTGCCGAAGCTGA
482 ACTCTCCAACTCAGTCGCTGGGATGGCTTTCCGTGCTGGTCGTCTCGATAATGGTTTTGATG
483 TGTTTAAAGACACCATTACGCCGACTCGCTGGAACTCTCACATCTGGACTCCAGAGGAGTTA
484 GAGAAGATTCGAACAGAGGTTAAGAACCCTGCGTACATCAACGTTGTAACTGGTGGTTCCCC
485 TGAGAACCTCGATGACCTCATTAAATTGGCTAACGAGAACTTTGAGAATGACTCCCGCGCTG
486 CCGAGGCTGGCCTAGGTGCCAAACTGAGTGCTGGTATTATTGGTGCTGGTGTGGACCCGCTT
487 AGCTATGTTCCTATGGTCGGTGTCACTGGTAAGGGCTTTAAGTTAATCAATAAGGCTCTTGT
488 AGTTGGTGCCGAAAGTGCTGCTCTGAACGTTGCATCCGAAGGTCTCCGTACCTCCGTAGCTG
489 GTGGTGACGCAGACTATGCGGGTGCTGCCTTAGGTGGCTTTGTGTTTGGCGCAGGCATGTCT
490 GCAATCAGTGACGCTGTAGCTGCTGGACTGAAACGCAGTAAACCAGAAGCTGAGTTCGACAA
491 TGAGTTCATCGGTCCTATGATGCGATTGGAAGCCCGTGAGACAGCACGAAACGCCAACTCTG
492 CGGACCTCTCTCGGATGAACACTGAGAACATGAAGTTTGAAGGTGAACATAATGGTGTCCCT
493 TATGAGGACTTACCAACAGAGAGAGGTGCCGTGGTGTTACATGATGGCTCCGTTCTAAGTGC
494 AAGCAACCCAATCAACCCTAAGACTCTAAAAGAGTTCTCCGAGGTTGACCCTGAGAAGGCTG
495 CGCGAGGAATCAAACTGGCTGGGTTCACCGAGATTGGCTTGAAGACCTTGGGGTCTGACGAT
496 GCTGACATCCGTAGAGTGGCTATCGACCTCGTTCGCTCTCCTACTGGTATGCAGTCTGGTGC
497 CTCAGGTAAGTTCGGTGCAACAGCTTCTGACATCCATGAGAGACTTCATGGTACTGACCAGC
498 GTACTTATAATGACTTGTACAAAGCAATGTCTGACGCTATGAAAGACCCTGAGTTCTCTACT
499 GGCGGCGCTAAGATGTCCCGTGAAGAAACTCGATACACTATCTACCGTAGAGCGGCACTAGC
500 TATTGAGCGTCCAGAACTACAGAAGGCACTCACTCCGTCTGAGAGAATCGTTATGGACATCA
501 TTAAGCGTCACTTTGACACCAAGCGTGAACTTATGGAAAACCCAGCAATATTCGGTAACACA
502 AAGGCTGTGAGTATCTTCCCTGAGAGTCGCCACAAAGGTACTTACGTTCCTCACGTATATGA
503 CCGTCATGCCAAGGCGCTGATGATTCAACGCTACGGTGCCGAAGGTTTGCAGGAAGGGATTG
504 CCCGCTCATGGATGAACAGCTACGTCTCCAGACCTGAGGTCAAGGCCAGAGTCGATGAGATG
505 CTTAAGGAATTACACGGGGTGAAGGAAGTAACACCAGAGATGGTAGAGAAGTACGCTATGGA
506 TAAGGCTTATGGTATCTCCCACTCAGACCAGTTCACCAACAGTTCCATAATAGAAGAGAACA
507 TTGAGGGCTTAGTAGGTATCGAGAATAACTCATTCCTTGAGGCACGTAACTTGTTTGATTCG
508 GACCTATCCATCACTATGCCAGACGGACAGCAATTCTCAGTGAATGACCTAAGGGACTTCGA
509 TATGTTCCGCATCATGCCAGCGTATGACCGCCGTGTCAATGGTGACATCGCCATCATGGGGT
510 CTACTGGTAAAACCACTAAGGAACTTAAGGATGAGATTTTGGCTCTCAAAGCGAAAGCTGAG
511 GGAGACGGTAAGAAGACTGGCGAGGTACATGCTTTAATGGATACCGTTAAGATTCTTACTGG
512 TCGTGCTAGACGCAATCAGGACACTGTGTGGGAAACCTCACTGCGTGCCATCAATGACCTAG
513 GGTTCTTCGCTAAGAACGCCTACATGGGTGCTCAGAACATTACGGAGATTGCTGGGATGATT
514 GTCACTGGTAACGTTCGTGCTCTAGGGCATGGTATCCCAATTCTGCGTGATACACTCTACAA
515 GTCTAAACCAGTTTCAGCTAAGGAACTCAAGGAACTCCATGCGTCTCTGTTCGGGAAGGAGG
516 TGGACCAGTTGATTCGGCCTAAACGTGCTGACATTGTGCAGCGCCTAAGGGAAGCAACTGAT
517 ACCGGACCTGCCGTGGCGAACATCGTAGGGACCTTGAAGTATTCAACACAGGAACTGGCTGC
518 TCGCTCTCCGTGGACTAAGCTACTGAACGGAACCACTAACTACCTTCTGGATGCTGCGCGTC
519 AAGGTATGCTTGGGGATGTTATTAGTGCCACCCTAACAGGTAAGACTACCCGCTGGGAGAAA
520 GAAGGCTTCCTTCGTGGTGCCTCCGTAACTCCTGAGCAGATGGCTGGCATCAAGTCTCTCAT
521 CAAGGAACATATGGTACGCGGTGAGGACGGGAAGTTTACCGTTAAGGACAAGCAAGCGTTCT
522 CTATGGACCCACGGGCTATGGACTTATGGAGACTGGCTGACAAGGTAGCTGATGAGGCAATG
523 CTGCGTCCACATAAGGTGTCCTTACAGGATTCCCATGCGTTCGGAGCACTAGGTAAGATGGT
524 TATGCAGTTTAAGTCTTTCACTATCAAGTCCCTTAACTCTAAGTTCCTGCGAACCTTCTATG
525 ATGGATACAAGAACAACCGAGCGATTGACGCTGCGCTGAGCATCATCACCTCTATGGGTCTC
526 GCTGGTGGTTTCTATGCTATGGCTGCACACGTCAAAGCATACGCTCTGCCTAAGGAGAAACG
527 TAAGGAGTACTTGGAGCGTGCACTGGACCCAACCATGATTGCCCACGCTGCGTTATCTCGTA
528 GTTCTCAATTGGGTGCTCCTTTGGCTATGGTTGACCTAGTTGGTGGTGTTTTAGGGTTCGAG
529 TCCTCCAAGATGGCTCGCTCTACGATTCTACCTAAGGACACCGTGAAGGAACGTGACCCAAA
530 CAAACCGTACACCTCTAGAGAGGTAATGGGCGCTATGGGTTCAAACCTTCTGGAACAGATGC
531 CTTCGGCTGGCTTTGTGGCTAACGTAGGGGCTACCTTAATGAATGCTGCTGGCGTGGTCAAC

532 TCACCTAATAAAGCAACCGAGCAGGACTTCATGACTGGTCTTATGAACTCCACAAAAGAGTT
533 AGTACCGAACGACCCATTGACTCAACAGCTTGTGTTGAAGATTTATGAGGCGAACGGTGTTA
534 ACTTGAGGGAGCGTAGGAAATAATACGACTCACTATAGGGAGAGGCGAAATAATCTTCTCCC
535 TGTAGTCTCTTAGATTTACTTTAAGGAGGTCAAATGGCTAACGTAATTAAAACCGTTTTGAC
536 TTACCAGTTAGATGGCTCCAATCGTGATTTTAATATCCCGTTTGAGTATCTAGCCCGTAAGT
537 TCGTAGTGGTAACTCTTATTGGTGTAGACCGAAAGGTCCTTACGATTAATACAGACTATCGC
538 TTTGCTACACGTACTACTATCTCTCTGACAAAGGCTTGGGGTCCAGCCGATGGCTACACGAC
539 CATCGAGTTACGTCGAGTAACCTCCACTACCGACCGATTGGTTGACTTTACGGATGGTTCAA
540 TCCTCCGCGCGTATGACCTTAACGTCGCTCAGATTCAAACGATGCACGTAGCGGAAGAGGCC
541 CGTGACCTCACTACGGATACTATCGGTGTCAATAACGATGGTCACTTGGATGCTCGTGGTCG
542 TCGAATTGTGAACCTAGCGAACGCCGTGGATGACCGCGATGCTGTTCCGTTTGGTCAACTAA
543 AGACCATGAACCAGAACTCATGGCAAGCACGTAATGAAGCCTTACAGTTCCGTAATGAGGCT
544 GAGACTTTCAGAAACCAAGCGGAGGGCTTTAAGAACGAGTCCAGTACCAACGCTACGAACAC
545 AAAGCAGTGGCGCGATGAGACCAAGGGTTTCCGAGACGAAGCCAAGCGGTTCAAGAATACGG
546 CTGGTCAATACGCTACATCTGCTGGGAACTCTGCTTCCGCTGCGCATCAATCTGAGGTAAAC
547 GCTGAGAACTCTGCCACAGCATCCGCTAACTCTGCTCATTTGGCAGAACAGCAAGCAGACCG
548 TGCGGAACGTGAGGCAGACAAGCTGGAAAATTACAATGGATTGGCTGGTGCAATTGATAAGG
549 TAGATGGAACCAATGTGTACTGGAAAGGAAATATTCACGCTAACGGGCGCCTTTACATGACC
550 ACAAACGGTTTTGACTGTGGCCAGTATCAACAGTTCTTTGGTGGTGTCACTAATCGTTACTC
551 TGTCATGGAGTGGGGAGATGAGAACGGATGGCTGATGTATGTTCAACGTAGAGAGTGGACAA
552 CAGCGATAGGCGGTAACATCCAGTTAGTAGTAAACGGACAGATCATCACCCAAGGTGGAGCC
553 ATGACCGGTCAGCTAAAATTGCAGAATGGGCATGTTCTTCAATTAGAGTCCGCATCCGACAA
554 GGCGCACTATATTCTATCTAAAGATGGTAACAGGAATAACTGGTACATTGGTAGAGGGTCAG
555 ATAACAACAATGACTGTACCTTCCACTCCTATGTACATGGTACGACCTTAACACTCAAGCAG
556 GACTATGCAGTAGTTAACAAACACTTCCACGTAGGTCAGGCCGTTGTGGCCACTGATGGTAA
557 TATTCAAGGTACTAAGTGGGGAGGTAAATGGCTGGATGCTTACCTACGTGACAGCTTCGTTG
558 CGAAGTCCAAGGCGTGGACTCAGGTGTGGTCTGGTAGTGCTGGCGGTGGGGTAAGTGTGACT
559 GTTTCACAGGATCTCCGCTTCCGCAATATCTGGATTAAGTGTGCCAACAACTCTTGGAACTT
560 CTTCCGTACTGGCCCCGATGGAATCTACTTCATAGCCTCTGATGGTGGATGGTTACGATTCC
561 AAATACACTCCAACGGTCTCGGATTCAAGAATATTGCAGACAGTCGTTCAGTACCTAATGCA
562 ATCATGGTGGAGAACGAGTAATTGGTAAATCACAAGGAAAGACGTGTAGTCCACGGATGGAC
563 TCTCAAGGAGGTACAAGGTGCTATCATTAGACTTTAACAACGAATTGATTAAGGCTGCTCCA
564 ATTGTTGGGACGGGTGTAGCAGATGTTAGTGCTCGACTGTTCTTTGGGTTAAGCCTTAACGA
565 ATGGTTCTACGTTGCTGCTATCGCCTACACAGTGGTTCAGATTGGTGCCAAGGTAGTCGATA
566 AGATGATTGACTGGAAGAAAGCCAATAAGGAGTGATATGTATGGAAAAGGATAAGAGCCTTA
567 TTACATTCTTAGAGATGTTGGACACTGCGATGGCTCAGCGTATGCTTGCGGACCTTTCGGAC
568 CATGAGCGTCGCTCTCCGCAACTCTATAATGCTATTAACAAACTGTTAGACCGCCACAAGTT
569 CCAGATTGGTAAGTTGCAGCCGGATGTTCACATCTTAGGTGGCCTTGCTGGTGCTCTTGAAG
570 AGTACAAAGAGAAAGTCGGTGATAACGGTCTTACGGATGATGATATTTACACATTACAGTGA
571 TATACTCAAGGCCACTACAGATAGTGGTCTTTATGGATGTCATTGTCTATACGAGATGCTCC
572 TACGTGAAATCTGAAAGTTAACGGGAGGCATTATGCTAGAATTTTTACGTAAGCTAATCCCT
573 TGGGTTCTCGCTGGGATGCTATTCGGGTTAGGATGGCATCTAGGGTCAGACTCAATGGACGC
574 TAAATGGAAACAGGAGGTACACAATGAGTACGTTAAGAGAGTTGAGGCTGCGAAGAGCACTC
575 AAAGAGCAATCGATGCGGTATCTGCTAAGTATCAAGAAGACCTTGCCGCGCTGGAAGGGAGC
576 ACTGATAGGATTATTTCTGATTTGCGTAGCGACAATAAGCGGTTGCGCGTCAGAGTCAAAAC
577 TACCGGAACCTCCGATGGTCAGTGTGGATTCGAGCCTGATGGTCGAGCCGAACTTGACGACC
578 GAGATGCTAAACGTATTCTCGCAGTGACCCAGAAGGGTGACGCATGGATTCGTGCGTTACAG
579 GATACTATTCGTGAACTGCAACGTAAGTAGGAAATCAAGTAAGGAGGCAATGTGTCTACTCA
580 ATCCAATCGTAATGCGCTCGTAGTGGCGCAACTGAAAGGAGACTTCGTGGCGTTCCTATTCG
581 TCTTATGGAAGGCGCTAAACCTACCGGTGCCCACTAAGTGTCAGATTGACATGGCTAAGGTG
582 CTGGCGAATGGAGACAACAAGAAGTTCATCTTACAGGCTTTCCGTGGTATCGGTAAGTCGTT

CATCACATGTGCGTTCGTTGTGTGGTCCTTATGGAGAGACCCTCAGTTGAAGATACTTATCG
TATCAGCCTCTAAGGAGCGTGCAGACGCTAACTCCATCTTTATTAAGAACATCATTGACCTG
CTGCCATTCCTATCTGAGTTAAAGCCAAGACCCGGACAGCGTGACTCGGTAATCAGCTTTGA
TGTAGGCCCAGCCAATCCTGACCACTCTCCTAGTGTGAAATCAGTAGGTATCACTGGTCAGT
TAACTGGTAGCCGTGCTGACATTATCATTGCGGATGACGTTGAGATTCCGTCTAACAGCGCA
ACTATGGGTGCCCGTGAGAAGCTATGGACTCTGGTTCAGGAGTTCGCTGCGTTACTTAAACC
GCTGCCTTCCTCTCGCGTTATCTACCTTGGTACACCTCAGACAGAGATGACTCTCTATAAGG
AACTTGAGGATAACCGTGGGTACACAACCATTATCTGGCCTGCTCTGTACCCAAGGACACGT
GAAGAGAACCTCTATTACTCACAGCGTCTTGCTCCTATGTTACGCGCTGAGTACGATGAGAA
CCCTGAGGCACTTGCTGGGACTCCAACAGACCCAGTGCGCTTTGACCGTGATGACCTGCGCG
AGCGTGAGTTGGAATACGGTAAGGCTGGCTTTACGCTACAGTTCATGCTTAACCCTAACCTT
AGTGATGCCGAGAAGTACCCGCTGAGGCTTCGTGACGCTATCGTAGCGGCCTTAGACTTAGA
GAAGGCCCCAATGCATTACCAGTGGCTTCCGAACCGTCAGAACATCATTGAGGACCTTCCTA
ACGTTGGCCTTAAGGGTGATGACCTGCATACGTACCACGATTGTTCCAACAACTCAGGTCAG
TACCAACAGAAGATTCTGGTCATTGACCCTAGTGGTCGCGGTAAGGACGAAACAGGTTACGC
TGTGCTGTACACACTGAACGGTTACATCTACCTTATGGAAGCTGGAGGTTTCCGTGATGGCT
ACTCCGATAAGACCCTTGAGTTACTCGCTAAGAAGGCAAAGCAATGGGGAGTCCAGACGGTT
GTCTACGAGAGTAACTTCGGTGACGGTATGTTCGGTAAGGTATTCAGTCCTATCCTTCTTAA
ACACCACAACTGTGCGATGGAAGAGATTCGTGCCCGTGGTATGAAAGAGATGCGTATTTGCG
ATACCCTTGAGCCAGTCATGCAGACTCACCGCCTTGTAATTCGTGATGAGGTCATTAGGGCC
GACTACCAGTCCGCTCGTGACGTAGACGGTAAGCATGACGTTAAGTACTCGTTGTTCTACCA
GATGACCCGTATCACTCGTGAGAAAGGCGCTCTGGCTCATGATGACCGATTGGATGCCCTTG
CGTTAGGCATTGAGTATCTCCGTGAGTCCATGCAGTTGGATTCCGTTAAGGTCGAGGGTGAA
GTACTTGCTGACTTCCTTGAGGAACACATGATGCGTCCTACGGTTGCTGCTACGCATATCAT
TGAGATGTCTGTGGGAGGAGTTGATGTGTACTCTGAGGACGATGAGGGTTACGGTACGTCTT
TCATTGAGTGGTGATTTATGCATTAGGACTGCATAGGGATGCACTATAGACCACGGATGGTC
AGTTCTTTAAGTTACTGAAAAGACACGATAAATTAATACGACTCACTATAGGGAGAGGAGGG
ACGAAAGGTTACTATATAGATACTGAATGAATACTTATAGAGTGCATAAAGTATGCATAATG
GTGTACCTAGAGTGACCTCTAAGAATGGTGATTATATTGTATTAGTATCACCTTAACTTAAG
GACCAACATAAAGGGAGGAGACTCATGTTCCGCTTATTGTTGAACCTACTGCGGCATAGAGT
CACCTACCGATTTCTTGTGGTACTTTGTGCTGCCCTTGGGTACGCATCTCTTACTGGAGACC
TCAGTTCACTGGAGTCTGTCGTTTGCTCTATACTCACTTGTAGCGATTAGGGTCTTCCTGAC
CGACTGATGGCTCACCGAGGGATTCAGCGGTATGATTGCATCACACCACTTCATCCCTATAG
AGTCAAGTCCTAAGGTATACCCATAAAGAGCCTCTAATGGTCTATCCTAAGGTCTATACCTA
AAGATAGGCCATCCTATCAGTGTCACCTAAAGAGGGTCTTAGAGAGGGCCTATGGAGTTCCT
ATAGGGTCCTTTAAAATATACCATAAAAATCTGAGTGACTATCTCACAGTGTACGGACCTAA
AGTTCCCCCATAGGGGGTACCTAAAGCCCAGCCAATCACCTAAAGTCAACCTTCGGTTGACC
TTGAGGGTTCCCTAAGGGTTGGGGATGACCCTTGGGTTTGTCTTTGGGTGTTACCTTGAGTG
TC

**Supplementary Data 2: Phagemid-p15A**

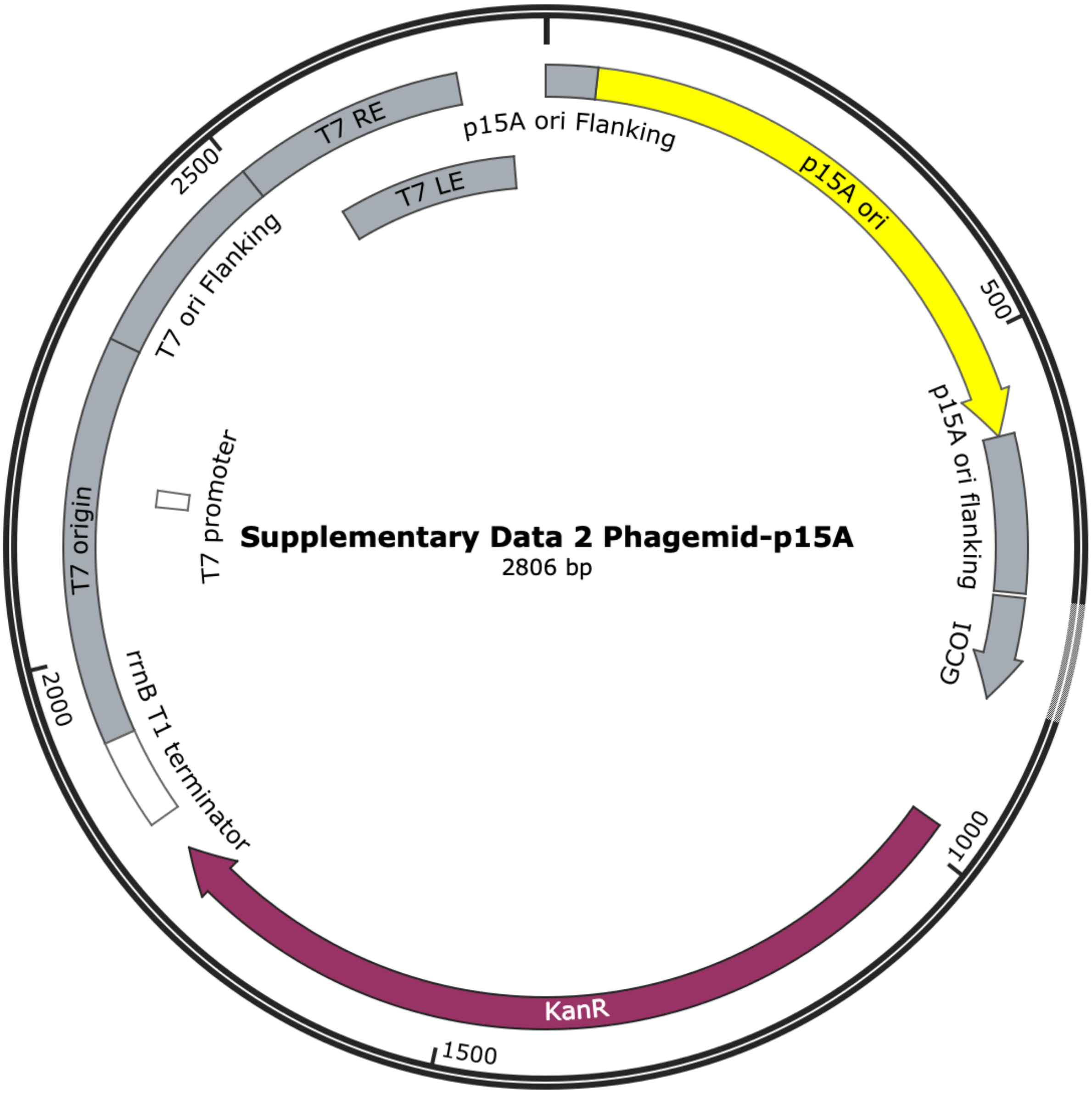


ACTTCGGGCTCATGAGCAAATATTTTATCTGATTAATAAGATGATCTTCTTGAGATCGTTTT
GGTCTGCGCGTAATCTCTTGCTCTGAAAACGAAAAAACCGCCTTGCAGGGCGGTTTTTCGAA
GGTTCTCTGAGCTACCAACTCTTTGAACCGAGGTAACTGGCTTGGAGGAGCGCAGTCACCAA
AACTTGTCCTTTCAGTTTAGCCTTAACCGGCGCATGACTTCAAGACTAACTCCTCTAAATCA
ATTACCAGTGGCTGCTGCCAGTGGTGCTTTTGCATGTCTTTCCGGGTTGGACTCAAGACGAT
AGTTACCGGATAAGGCGCAGCGGTCGGACTGAACGGGGGGTTCGTGCATACAGTCCAGCTTG
GAGCGAACTGCCTACCCGGAACTGAGTGTCAGGCGTGGAATGAGACAAACGCGGCCATAACA
GCGGAATGACACCGGTAAACCGAAAGGCAGGAACAGGAGAGCGCACGAGGGAGCCGCCAGGG
GGAAACGCCTGGTATCTTTATAGTCCTGTCGGGTTTCGCCACCACTGATTTGAGCGTCAGAT
TTCGTGATGCTTGTCAGGGGGGCGGAGCCTATGGAAAAACGGCTTTGCCGCGGCCCTCTCAC
TTCCCTGTTAAGTATCTTCCTGGCATCTTCCAGGAAATCTCCGCCCCGTTCGTAAGCCATTT
CCGCTCGCCGCAGTCGAACGACCGAGCGTAGCGAGTCAGTGAGCGAGGAAGCGGAATATATC
CTAGGNNNNNNNNNNNNNNNNNNNNNNNNNNNNNNNNNNNNNNNNNNNNNNNNNNNNNNN
NNNNNNNNNNNNNNNNNNNNNNNNNNNNNNNNNNNNNNNNNTTTTTTTAAGGCAGTTATT
GGTGCCCTTAAACGTCTGACGCTCAGTGGAACGAAAACTCACGTTAAGGATTTTGGTCATG

19 AACAATAAAACTGTCTGCTTACATAAACAGTAATACAAGGGGTGTTATGAGCCATATTCAAC
20 GGGAAACGTCTTGCTCTAGGCCGCGATTAAATTCCAACATGGATGCTGATTTATATGGGTAT
21 AAATGGGCTCGCGATAATGTCGGGCAATCAGGTGCGACAATCTATCGATTGTATGGGAAGCC
22 CGATGCGCCAGAGTTGTTTCTGAAACATGGCAAAGGTAGCGTTGCCAATGATGTTACAGATG
23 AGATGGTCAGACTAAACTGGCTGACGGAATTTATGCCTCTTCCGACCATCAAGCATTTTATC
24 CGTACTCCTGATGATGCATGGTTACTCACCACTGCGATCCCCGGGAAAACAGCATTCCAGGT
25 ATTAGAAGAATATCCTGATTCAGGTGAAAATATTGTTGATGCGCTGGCAGTGTTCCTGCGCC
26 GGTTGCATTCGATTCCTGTTTGTAATTGTCCTTTTAACAGCGATCGCGTATTTCGTCTCGCT
27 CAGGCGCAATCACGAATGAATAACGGTTTGGTTGATGCGAGTGATTTTGATGACGAGCGTAA
28 TGGCTGGCCTGTTGAACAAGTCTGGAAAGAAATGCATAAACTTTTGCCATTCTCACCGGATT
29 CAGTCGTCACTCATGGTGATTTCTCACTTGATAACCTTATTTTTGACGAGGGGAAATTAATA
30 GGTTGTATTGATGTTGGACGAGTCGGAATCGCAGACCGATACCAGGATCTTGCCATCCTATG
31 GAACTGCCTCGGTGAGTTTTCTCCTTCATTACAGAAACGGCTTTTTCAAAAATATGGTATTG
32 ATAATCCTGATATGAATAAATTGCAGTTTCATTTGATGCTCGATGAGTTTTTCTAAAAGCTT
33 AATTAGCTGATCTAGACGCGTGCTAGAGGCATCAAATAAAACGAAAGGCTCAGTCGAAAGAC
34 TGGGCCTTTCGTTTTATCTGTTGTTTGTCGGTGAACGCTCTCCTGAGTAGGACAAATAGGTC
35 GAGGGTGAAGTACTTGCTGACTTCCTTGAGGAACACATGATGCGTCCTACGGTTGCTGCTAC
36 GCATATCATTGAGATGTCTGTGGGAGGAGTTGATGTGTACTCTGAGGACGATGAGGGTTACG
37 GTACGTCTTTCATTGAGTGGTGATTTATGCATTAGGACTGCATAGGGATGCACTATAGACCA
38 CGGATGGTCAGTTCTTTAAGTTACTGAAAAGACACGATAAATTAATACGACTCACTATAGGG
39 AGAGGAGGGACGAAAGGTTACTATATAGATACTGAATGAATACTTATAGAGTGCATAAAGTA
40 TGCATAATGGTGTACCTAGAGTGACCTCTAAGAATGGTGATTATATTGTATTAGTATCACCT
41 TAACTTAAGGCGGGATCGTCACCCTCAGCAGCGAAAGACAGCTGTCGGTCAGAGCGTCATTG
42 CGAAGCTGAGTGTGATCGATGCCATCAGCGAAGGGCCCAAACTCCGAGCGATTAAGCGTTTG
43 CTGGCTGTCACGCCTGCCTGTTGCTTGCTTGGACTTGCGATGTACGTGCTCAGCTGTCTTTC
44 GCTGCTGAGGGTGACGATCCCGCGAGGGCCTATGGAGTTCCTATAGGGTCCTTTAAAATATA
45 CCATAAAAATCTGAGTGACTATCTCACAGTGTACGGACCTAAAGTTCCCCCATAGGGGGTAC
46 CTAAAGCCCAGCCAATCACCTAAAGTCAACCTTCGGTTGACCTTGAGGGTTCCCTAAGGGTT
47 GGGGATGACCCTTGGGTTTGTCTTTGGGTGTTACCTTGAGTGTCTCTCTGTGTCCCTATCTG
48 TTACAGTCTCCTAAAGTATCCTCCTAAAGTCACCTCCTAACGCACATTTCCCCGAAAAGTGC
49 CACCTGGGTCCTTTTC

**Supplementary Data 3: Simulation MDP**

**T7_step4.0_minimization.mdp**

```
define                 = -DPOSRES -DPOSRES_FC_BB=400.0 -DPOSRES_FC_SC=40.0
integrator             = steep
emtol                  = 1000.0
nsteps                 = 5000
nstlist                = 10
cutoff-scheme          = Verlet
rlist                  = 1.2
vdwtype                = Cut-off
vdw-modifier           = Force-switch
rvdw_switch            = 1.0
rvdw                   = 1.2
coulombtype            = PME
rcoulomb               = 1.2
;
constraints            = h-bonds
constraint_algorithm   = LINCS
```

**T7_step4.1_equilibration.mdp**

```
define                = -DPOSRES -DPOSRES_FC_BB=400.0 -
DPOSRES_FC_SC=40.0
integrator            = md
dt                    = 0.001
nsteps                = 125000
nstxout-compressed    = 5000
nstxout               = 0
nstvout               = 0
nstfout               = 0
nstcalcenergy         = 100
nstenergy             = 1000
nstlog                = 1000
;
cutoff-scheme         = Verlet
nstlist               = 20
rlist                 = 1.2
vdwtype               = Cut-off
vdw-modifier          = Force-switch
rvdw_switch           = 1.0
rvdw                  = 1.2
coulombtype           = PME
rcoulomb              = 1.2
;
tcoupl                = v-rescale
tc_grps               = SOLU SOLV
tau_t                 = 1.0 1.0
ref_t                 = 303.15 303.15
;
constraints           = h-bonds
constraint_algorithm  = LINCS
;
nstcomm               = 100
comm_mode             = linear
comm_grps             = SOLU SOLV
;
gen-vel               = yes
gen-temp              = 303.15
gen-seed              = -1
```

**T7_step5_production.mdp**

```
integrator              = md
dt                      = 0.002
nsteps                  = 5000000  ; 10 ns
nstxout-compressed      = 50000
nstxout                 = 0
nstvout                 = 0
nstfout                 = 0
nstcalcenergy           = 100
nstenergy               = 5000
nstlog                  = 5000
;
cutoff-scheme           = Verlet
nstlist                 = 40
vdwtype                 = Cut-off
vdw-modifier            = Force-switch
rvdw_switch             = 1.0
rvdw                    = 1.2
rlist                   = 1.2
rcoulomb                = 1.2
coulombtype             = PME
;
tcoupl                  = v-rescale
tc_grps                 = SOLU SOLV
tau_t                   = 1.0 1.0
ref_t                   = 303.15 303.15
;
pcoupl                  = C-rescale
pcoupltype              = isotropic
tau_p                   = 5.0
compressibility         = 4.5e-5
ref_p                   = 1.0
;
constraints             = h-bonds
constraint_algorithm    = LINCS
continuation            = no
lincs-order             = 6
lincs-warnangle         = 30
;
nstcomm                 = 100
comm_mode               = linear
comm_grps               = SOLU SOLV
;
gen_vel                 = yes
```

### Supplementary Data 4: Phagemid-BAC, pSJ77

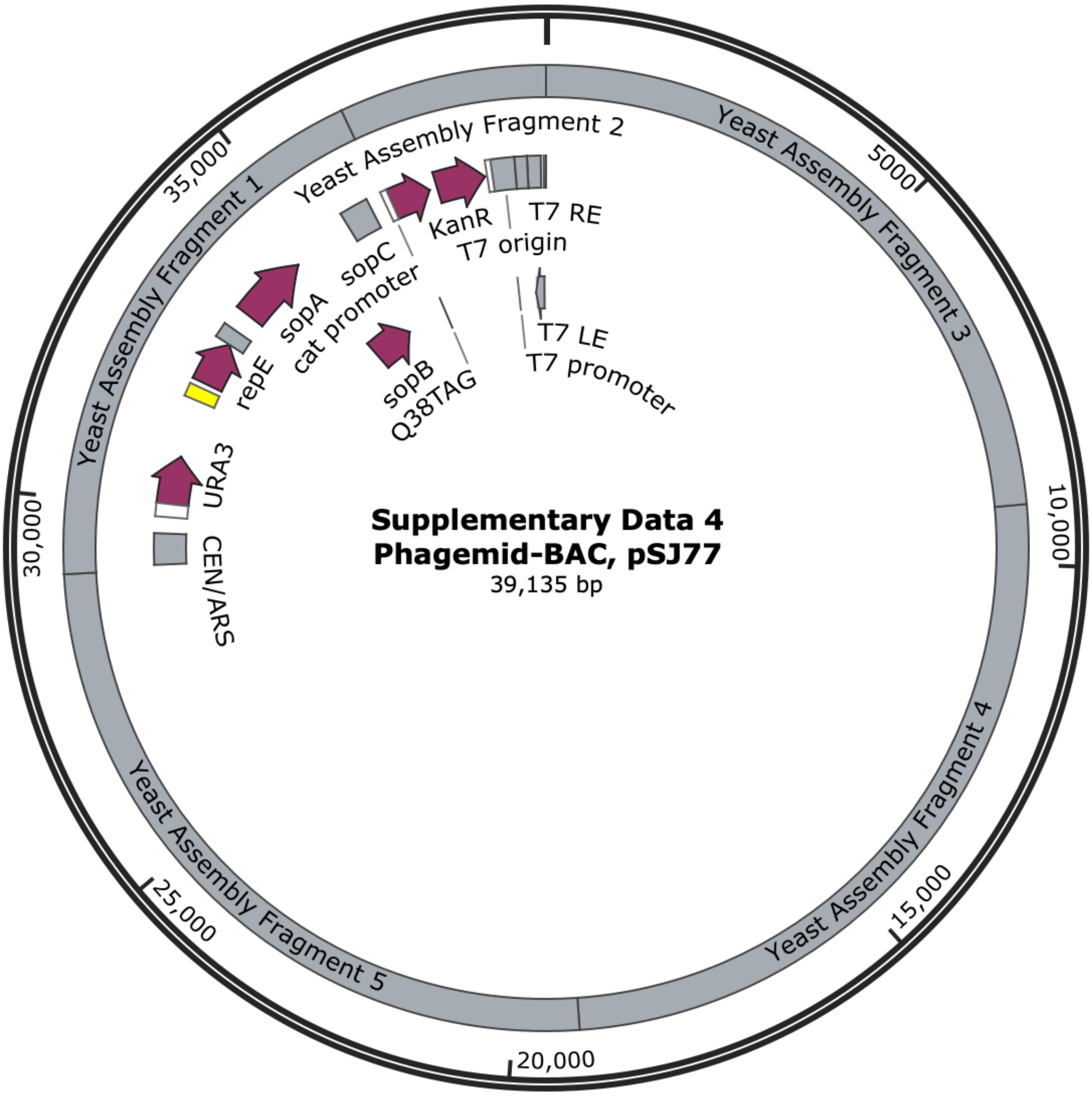


AAGAAGCGCTTTAGAAATCAAAGCACAACGTAACAATTTGTCGACAACCGAGCCTTTGAAGA
AAAAATTTTTCACATTGTCGCCTCTAAATAAATAGTTTAAGGTTATCTACCCACTATATTTA
GTTGGTTCTTTTTTTTTTCCTTCTACTCTTTATCTTTTTACCTCATGCTTTCTACCTTTCAG
CACTGAAGAGTCCAACCGAATATATACACACATAATGGCATCCACCGATTTCTCCAAGATTG
AAACTTTGAAACAATTAAACGCTTCTTTGGCTGACAAGTCATACATTGAAGGGTATGTTCCG
ATTTAGTTTACTTTATAGATCGTTGTTTTTCTTTCTTTTTTTTTTTTCCTATGGTTACATGT
AAAGGGAAGTTAACTAATAATGATTACTTTTTTTCGCTTATGTGAATGATGAATTTAATTCT
TTGGTCCGTGTTTATGATGGGAAGTAAGACCCCCGATATGAGTGACAAAAGAGATGTGGTTG
ACTATCACAGTATCTGACGATAGCACAGAGCAGAGTATCATTATTAGTTATCTGTTATTTTT
TTTTCCTTTTTTGTTCAAAAAAAGAAAGACAGAGTCTAAAGATTGCATTACAAGAAAAAAGT
TCTCATTACTAACAAGCAAAATGTTTTGTTTCTCCTTTTAAAATAGTACTGCTGTTTCTCAA
GCTGACGTCACTGTCTTCAAGGCTTTCCAATCTGCTTACCCAGAATTCTCCAGATGGTTCAA
CCACATCGCTTCCAAGGCCGATGAATTCGACTCTTTCCCAGCTGCCTCTGCTGCCGCTGCCG
AAGAAGAAGAAGATGACGATGTCGATTTATTCGGTTCCGACGATGAAGAAGCTGACGCTGAA
GCTGAAAAGTTGAAGGCTGAAAGAATTGCCGCATACAACGCTAAGAAGGCTGCTAAGCCAGC
TAAGCCAGCTGCTAAGTCCATTGTCACTCTAGATGTCAAGCCATGGGATGATGAAACCAATT

20 TGGAAGAAATGGTTGCTAACGTCAAGGCCATCGAAATGGAAGGTTTGACCTGGGGTGCTCAC
21 CAATTTATCCCAATTGGTTTCGGTATCAAGAAGTTGCAAATTAACTGTGTTGTCGAAGATGA
22 CAAGGTTTCCTTGGATGACTTGCAACAAAGCATTGAAGAAGACGAAGACCACGTCCAATCTA
23 CCGATATTGCTGCTATGCAAAAATTATAAAAGGCTTTTTTATAAACTTTTTATAATTAACAT
24 TAAAGCAAAAACAACATTGTAAAGATTAACAAATAAATGAAAAAAACAACGAAATAACTTAG
25 GTTTTAGGCTAAAAAAAACAGAAGGAATTTTGAACGATAAACTTTTCGACTGCACACGAAAC
26 ATTATTACTAATTTGTGTAACCACTATATAAGGAATCGTGTTTATTAATTGAATTTATTCCG
27 GGAATATTCAAGTTATGTATATCTCTTTTCATATTCTTAAATACACATACTCATAATATCTT
28 GTCGAAAATACGCGGTGTAGGGAGTTATGGTGGATAACTTTTTCACGATTAGAAGAAAAGGA
29 AAATTTCATTATTCGTAGCTTAACATGGCAAAAACGAGAAAGACATATAATCAAAACGTGAG
30 TTTCCTGTGGAAAAAAAAAAAAAGGGAACCTCTGGTTACGATGATATACCTGCGTGAAAAAGG
31 ACAGTTATTACCAATACATACAAAGGCTTAATAAGTGTAAAATATATATCTGCCGAGACCAT
32 TACTCATTACACCTAGAATGGAGCAAAATGGCCTTGACCACGACAGCAGATCTAGCATCGAT
33 ACGACTATTAATGACACTCAAAAGACTTTCCTAGAATTTAGATCGTATACCCAATTAAGTGA
34 AAAACTGGCATCTAGTTCTTCATATACGGCACCTCCCCTGAACGAAGATGGTCCTAAAGGGG
35 TAGCTTCTGCAGTGTCACAAGGCTCCGAATCCGTAGTCTCATGGACAACTTTAACACACGTA
36 TATTCCATCCTGGGTGCTTATGGAGGGCCCACGTGCTTGTATCCGACAGCCACGTATTTTTT
37 GATGGGCACTTCTAAAGGATGCGTACTCATTTTTAATTATAATGAACATTTGCAGACAATCC
38 TAGTGCCGACCTTATCTGAGGACCCTTCTATTCACTCAATAAGAAGTCCAGTGAAATCAATT
39 GTCATATGTTCCGATGGTACTCATGTAGCTGCCTCATACGAGACCGGAAATATATGCATTTG
40 GAACTTGAACGTAGGGTATAGAGTGAAACCCACTTCTGAACCAACAAATGGTATGACCCCAA
41 CGCCTGCCTTACCGGCAGTCTTACACATCGATGACCATGTGAACAAGGAAATCACAGGGTTA
42 GACTTTTTTGGTGCTCGGCATACAGCCCTGATTGTTAGTGATAGGACAGGTAAAGTATCACT
43 CTATAACGGTTACAGAAGAGGCTTTTGGCAGTTGGTGTATAATTCAAAAAAAATTTTAGATG
44 TGAACTCTTCCAAGGAGAAATTAATAAGGTCAAAGTTGTCTCCACTAATATCACGGGAGAAA
45 ATTTCCACTAATTTGTTGAGTGTACTCACAACTACACATTTTGCCCTTATTTTATTATCGCC
46 ACACGTTTCTTTGATGTTTCAAGAAACTGTTGAACCCTCAGTACAAAATTCTCTAGTCGTGA
47 ATAGCTCTATTTCATGGACTCAAAACTGTTCCAGGGTTGCTTATTCCGTAAATAATAAAATT
48 TCTGTTATTTCCATATCTTCATCAGACTTCAATGTTCAGTCCGCTAGCCATTCTCCTGAATT
49 TGCAGAATCTATATTATCCATTCAATGGATTGACCAGCTCCTACTTGGTGTTTTAACCATAT
50 CGCACCAATTTTTGGTATTGCACCCCCAACATGACTTCAAGATCCTGTTAAGATTGGATTTT
51 CTGATTCACGATTTGATGATCCCACCTAATAAATATTTTGTAATAAGTAGAAGAAGTTTCTA
52 CCTGTTAACAAACTACTCATTTAAAATTGGCAAATTTGTGTCTTGGTCAGATATTACTTTAA
53 GACATATTTTGAAAGGCGACTACTTGGGTGCATTGGAGTTCATAGAATCACTTTTGCAACCT
54 TACTGTCCACTGGCAAACTTGTTGAAGCTAGATAATAATACGGAAGAGAGGACTAAGCAACT
55 TATGGAACCATTTTACAATCTGTCCTTGGCTGCCCTAAGGTTTCTTATAAAAAAAGATAATG
56 CCGACTACAATAGGGTTTACCAATTATTAATGGTAGTTGTTCGTGTTTTGCAGCAATCTTCC
57 AAAAAACTAGACTCAATTCCTTCTCTAGACGTCTTTTTGGAACAGGGTCTGGAGTTCTTTGA
58 ATTGAAGGACAACGCGGTATATTTTGAAGTTGTAGCAAATATTGTTGCCCAAGGATCAGTTA
59 CGTCAATTTCCCCAGTTCTTTTCAGGTCCATAATTGATTACTATGCTAAGGAGGAGAATTTA
60 AAAGTAATTGAAGACTTAATCATCATGTTAAATCCTACTACGCTTGATGTTGATCTTGCCGT
61 CAAACTATGCCAAAAGTATAATTTGTTCGATTTATTAATATATATTTGGAACAAGATCTTTG
62 ATGATTATCAAACCCCAGTGGTGGACTTGATATACAGGATTTCTAACCAAAGTGAAAAATGT
63 GTGATCTTCAATGGTCCTCAAGTACCTCCTGAAACGACTATATTTGATTACGTAACGTATAT
64 CCTTACTGGCAGGCAATATCCACAAAACTTGTCTATATCACCAAGTGATAAATGCTCCAAAA
65 TACAAAGGGAACTTTCAGCATTTATTTTTAGTGGCTTCTCCATAAAATGGCCGTCGAACAGC
66 AATCATAAACTTTACATATGCGAAAATCCAGAAGAAGAGCCAGCATTTCCTTACTTTCACCT
67 TTTATTGAAATCGAATCCGAGTAGGTTCTTAGCAATGCTCAATGAAGTGTTTGAAGCGTCCT
68 TGTTTAACGATGACAATGACATGGTTGCATCAGTTGGAGAAGCAGAATTGGTAAGTAGGCAA
69 TATGTTATTGATCTACTATTGGATGCTATGAAAGATACGGGAAATTCAGACAACATCAGGGT
70 ACTTGTTGCAATTTTCATTGCAACTAGTATATCAAAATATCCTCAATTTATTAAAGTGTCTA

ACCAAGCCCTCGACTGCGTTGTTAATACCATATGCTCCTCTAGGGTTCAAGGTATATATGAA
ATTTCTCAAATAGCTCTGGAGTCGCTTTTACCCTATTATCATTCAAGAACAACAGAAAATTT
TATACTGGAACTAAAAGAAAAAAATTTCAATAAAGTTCTTTTCCATATCTATAAAAGTGAAA
ATAAGTACGCCAGTGCGCTTTCACTTATTTTAGAAACTAAGGACATCGAAAAAGAATATAAC
ACGGACATTGTATCCATAACCGACTACATACTCAAAAAATGCCCACCTGGAAGTTTAGAATG
TGGCAAAGTTACTGAAGTTATCGAGACGAACTTTGATCTTCTTCTCTCAAGGATCGGTATCG
AAAAATGCGTCACAATTTTTTCTGACTTTGACTATAATCTTCATCAAGAAATCCTGGAAGTA
AAGAATGAGGAGACTCAGCAAAAGTATTTGGATAAGCTTTTTTCTACGCCAAATATCAACAA
TAAGGTCGATAAGCGTTTAAGAAATTTACACATCGAATTGAACTGTAAATACAAGAGCAAAA
GGGAAATGATTCTTTGGCTTAATGGTACAGTTCTCAGCAACGCTGAGAGCTTACAAATTCTG
GACTTATTGAATCAAGACTCTAATTTTGAAGCTGCAGCTATAATTCACGAACGCTTGGAAAG
TTTTAACCTAGCAGTCAGGGATTTATTAAGTTTTATTGAACAATGTCTAAATGAAGGGAAAA
CAAATATATCTACTTTATTGGAATCTTTGAGGAGGGCCTTTGATGATTGTAATTCTGCTGGT
ACCGAGAAAAAATCGTGTTGGATATTATTGATTACATTCCTGATCACTCTATATGGGAAATA
TCCTTCACACGATGAAAGGAAAGATTTATGTAATAAACTACTTCAAGAAGCATTTTTGGGAT
TGGTTAGGTCCAAGAGTTCCTCTCAGAAGGATTCAGGTGGGGAATTCTGGGAAATAATGTCT
TCTGTTCTTGAGCACCAAGACGTTATTTTAATGAAAGTTCAGGATTTAAAGCAACTGCTACT
GAATGTTTTTAATACTTATAAATTGGAAAGATCTCTTTCTGAGTTGATTCAAAAGATTATAG
AGGATTCTTCGCAAGATCTTGTTCAACAGTATAGAAAATTTCTGAGTGAAGGGTGGTCTATA
CACACCGACGACTGCGAAATCTGCGGGAAAAAAATATGGGGAGCTGGTCTGGACCCATTACT
TTTTCTAGCTTGGGAAAATGTACAGCGCCACCAAGATATGATTAGTGTAGATCTCAAAACTC
CCCTTGTCATATTCAAATGTCACCATGGCTTTCACCAGACTTGCCTCGAAAACTTGGCCCAG
AAACCCGATGAATATTCTTGTTTAATTTGCCAGACGGAATCTAACCCAAAAATAGTATAACA
TTTCTAAATATTTAATACAACTTTGGTTACATAAAAGTAAAATTTATACACCTCATTTCATT
ATGTAGATTCATATATAGAATACCAATTATGATTGACCCAATAGCCATCAAAATCAGTAGTT
ATTAATACTTGTCTTTCTAGGAGCCATTTGCATATTTCTGATATTTCATGAAGCGAAAGTAC
TTCACGACACCTAGATTGCAATCTACTCAATGTTATCCCTGGATGAAATATTATTTCGTTAA
CGACCATAGTAACTACCTGCTTCCATATGTTTGGCCTAATGGAACCAGATCCATTCACCCAT
AAACGAGAAAATGGTTTGCCCAGTGGAACTTTGACAGCAGACTTCCTTGCTGTATTCAATTT
TGTCTGAGAATTGGCATATATAATCAGAGGGGGAGTTAATGTTCGTATTTCAAATCTCCTTG
AAGTATACGTTAAAGGTCGAACATTTCTCACCATTGGAATTACATCCATATTCAATAGCTCT
CCCGAAATCAAATCAATTAAAACCCAAGAGGATATATCGGACGGCTCTTGATTGATAACAAT
AGCGTTTCCGGCCTCCAATAATTCATTAACCTTACATCTATACTGAAAAGCTACACCAAAAT
CTTTATAATTTCCTCTATTTTCCAAAATGTCTGGTAAAGTATCAGTACATTCAAGTTTTGAG
CCATGGAGATAAATTTGCTTTTCCTTAGCCATATCCATGATGACGTTATCTATTGATTCGTT
TCCAACGTTCTTCAACGCCTCTATTTCATTTCTAGTGGTCGAAGGACTTTCTATTAATATGG
ACCGGATCACTGTGCGAATATAATCGTCGCTTTGACTCTTCGATAAGTCCTTAGTAGAAGCG
GAAATCTTTCTAGTGTAAGTTTTTTTAAAGAAGAGATCTCTCTTTGAATCATAGAAGACAT
GGCCAGATCGTTGCCAGAATGTGTAAGTGTGTCATCACGTACCAGAGTAAATTTTTTTCTAT
TCTCTTCGTAGTTCTTATAAAGAAAGAGCGGCCTTTTTATTTCCTTTTCATCAAAAGAGGTC
CAAATATCAAGCAATTTGATAAGATCTAGTTCTTCAACATCCCTCAGTGAAATCTTTTCACT
TTTAATGGCTAGAACGAGCATTTTTTTCCACTTATCGACATATGCCCTCCAACCACTATGAC
CCATTCTTACTCGCCGTGCCGTCCATTTCTTTTTTAGGTTATCTAAAGAATTATTAGGAAAT
AATTTTGTTATTTTGTCCCACATTATTTCATTTTTAATACTTTTAGTAACTACAACAGCTCT
GATTAAAGCCTGAACACCATCTTTAGTGCCTGCATGATAGACAGTTTTGTCTTCTTTAGTAT
TTTCCACGACCACCGTTGTCCTACCAGCAGACACTTTTTTGTCTCTCCTTTTGATCTTTCCA
TCTGATACGTTGACCGACGTACTCTTCTTAGAGATAGCGTCATCTGAAGCTTTGATCTTAGC
ATTCTTTTGGCGGTTCTGAAACTTACGAATTCTTTCAACTGATTTCTTTCCTCTGTGAAACC
TATTTTTTTCCGTTTGGTCAAAGAAGTATATTTCCGTATCATGTATAACATCAGTAAAGGTT
GCTTTTTTACTATCTTTTTCTTTCAGGATGTACCTTTGGATAGCGTCCTCTCCAACAGTGGG
CAAAAAAATAATTTTTCTTCCTGATACAGGCTCTGTTCTGGCTCCTAATTTTTCGCTTTCTA

122 CCATCAAATCAACATCACCACGGACAGTCTTTTTATCTAATGTCGTTGTGGAGCCCATATAT
123 TTAGAAACGCTTTCGTAAAATTGTTCTCTCAGGTATGCTACCCCACCAATCGTATTCATAAC
124 TTTCAAAATGGCTCTCTGTCTCTGTAGTGAACGCAAAGAGCGGGCAGAAAAGCCGCCGAAGT
125 TAACCACCTTGCCTTTGACAACAGTGCCTCCGTTTGAACTTGGACTATCTTCAGCAGATTTC
126 GGCTCCTGTGCAGTGCTGACATGCTGCTCTAGCTTAATCCTTTTGGGATTCGAAATGTTTCC
127 TGCAACAGAAGCATTAGTACTGTTTTTAACCTGCCTCTTCCGTTTGTTTTTATTCGGAGTTT
128 TTTTTGAGTTTGGGGGAATTTTTAATTCACCGTGCCAGAAGAATATATCCTGTCCATCGCTG
129 TCCGTTGTAAATCTAACAGTGTTGTTGAGTGCGACGAAATTATCCTCGTTGAGAGTTTTCAA
130 ATCGGTACGAGATTTGCCTAGCTCATCAAACCCTTTTGGAACGGATATTTCGTCTTCCGCAT
131 TTGTTAACTTTTGAAAGTTCTGAGCTGTGAACAGCCTAAAAAACTTCTTCTTTCCCTCAAAA
132 TCGTATATGCGAAAAAGCCTATACCCCCCTGTATTTTCTTTTTGCTTATCCACACTTTCTAA
133 ATAATATTCGCTTGATTTGGTAAAAGCTCGCTGAAATTCTTTTCCGGTAATTCGATTTACAA
134 CATCCATAGTTGAAATTCCTTTAAGGCCAGACTTATCTGCAATGTCATAAGTCTGATTTTGA
135 AGTGGATAAAATCGATTAAGAAGAACTTCATTCTTTACAGCATCCTCTTTCTCTTCCATAAC
136 AAGGCCTTGATTTTGTAATAAATCAGTCGCATTGAAATTATCTAAACCTTCGACTAAGTCTT
137 CATCTTCGAAAGCTGCCTTGCTATCTGATACAGAATCTTCATCCGCGCTATTGCTATCATAC
138 TCAAATGAAGGCGAGCCTTTAGAGTCTGGAATATCTTTCACGTATTTTACACATCTGATTTT
139 AATGGCAGGATTCTTGGGTGATACTACAAGCACTTTCTTTAAGTACTCCTTTTCATCTAACC
140 ATGCAATAGCTGCAATAAAAGCTTTAGAAAGTCTTTTCTCTTTGTCAAATTTCAATTCACGC
141 TTTAAATCAATTATCTGGCGAATACCATTTTTTGATCGTTTTACCACCTCAACTATTGTTGC
142 TAAATGATCCCTAATATTAATATAGGGATTACTATCCACCCCGTCATGGCTGAATTTTTTTA
143 GCTTCAATTGCTTCACGACGTGTCCCTTATAAATCAGTTGTGAACTTGTTAACAGGTGGTTT
144 ATTTTCTTGATACGTCCAGTCACACTTCTAGGATCTTGCCCAGTTACCTGCGCCAAATCCAT
145 AGTATTGATCCCTTTTTCTCCTGATTTGGCAACTTCGAGAAGTAGTTCAAATGCAGAATTTC
146 CAATAGTTGACTCCTTTTTTGTGTATCCCGTTAATAATGTCCATAGGCTGTCCTCAGTAATC
147 CCAACCGAGTATGAATGATTAGCGTCGCCTATAATATCAGTCACATTTTTAGTTGTTATAGC
148 ACCATCACAATACACCTCAATGTCCTTTTTCAATATCACGCATGAAAGCACGAACTGTTTAA
149 CTTTTTTATCAGACAAATCAAAATATTTACCAGATATATCCCACAGCTGATTCAAAGTGATT
150 TCTTCAATGTGTCGTTAGTAAATAATCTTTCACAATATAGTACGTTTATCACCTAAACGGAG
151 CCGAAAAGGAGAATGAGACATGAACATACTTCCCTTATTTGAAGCAATTTTATCAGACACTA
152 TTTGTACGAGTTCGTCAGGATAAATCGTCAGTACCATTTTTCGGTGAGTAACTGAGTAAAAA
153 CATTAGGCAAATTTCTTCCAAAAAATCATTGACATTGCCTGTCAATTGAGTAAATGAATAAA
154 TAAATAAATGCAAAAAAGCCAACCTAACTCTCCTGCAATGATACAATGAGTACTGTATCAAC
155 GACCATTAGTAAGTATCATGAAGTGTTTTGTATATTTGGAAAAAACCAGGCTGCATTGCCCC
156 AGTGGTGTATTTATATCTACTGCAGTATGTTCTATTCTGTGAAGTTCCTAAAAGACACCGAG
157 GTTTCTATGGCCTAACCATATTGTTGTTCTAACGCCTTTTCTATGTAAGGGTCCTTATTGTT
158 CAAAGCCCCACCGCTCCTTCGCGCGTTGTGCCTATCATTGTGTCAGTAAATGTTAGGATACG
159 GGAAGTGTATATATTAATTGAGTGGACACAGAAAACGCTCTAGAAAGGAATATCTCACAAAG
160 AACTGTAATATAAGCCTACATTATTGATTTTTTTAGTTCTCTTGCCCTTATATTACCGTAAG
161 TCTGCGGTATAATCACCTGGCCTAGTGCTTTTTCAATCATGTTCATTGAATACTCTAGATTA
162 CCAGGCTTCGAAAGCATCAATATATCATTTTCTAGAGGCATGCTAAGATTGGCTAAATTTAC
163 TAATTTCGCTACATACAAACAAAAATTAGAGTACTTCAGGCTGTTGGCAGGTTCAAACAAAT
164 ACATCCAACGAATCAGCGTTGCTGACTTTGAGAGACATCCAGATGAAATCAATTATATTTAC
165 ATTATACTAATATCCATTCTCCAAATGGAAGAGTGTATGCCAGTCCTAGTTCTATGTCCAAC
166 TGTTTATTGGGTACGATTCCACTGGCCTGGAAAATGTTCCGTGAACAGTTTGAACTTTACGA
167 ATGAAACGTTAAAAAGCGCTTTCCATGCTGTCTTCACTCCCTATTTTGCACTAATGAAAAAA
168 GTTCTTGGAAGGATTAAAAACAACATGTTGCTATTTGCAGAACCTCATGCAAATTTAAATAA
169 TTTGTTTGTCAAGCATTTCCACGATCTCATATATAAGAGCGTAAAGGATGAAAAAACCGGCG
170 AGGCAATTCTGTATTTGAGAACTAACGTTAATGTTCCAAATGTTTTTATTGACGATAAAAGA
171 GCAGTTTTTCATGGTGACGGTATGAAAATTGGAAAATTTACCGGCAAATTTTTATGCTTTTC
172 CTTCAAACGAACAATAAGATGGTCAAAATTAGATTCAGTGGATTCTTTTGCGGTTACGACAG

173 TAAATTATAGAGTATCTGTAAATTGGGAAAAGACACCAAGAAAGACTTTCCTTTCTCTAGAC
174 AGTGACACCAAAAATTTACACTATATATCGAAAAAGATATTGAATAAAAAGGGAAAGAACGC
175 CACTACATCAAAAACAACAAAAAGTTCATGCACAAGTGAGAACGTTTGCGATGATAAAACAT
176 TCTCCGTCGAATTCCCACTTACAACGTCAGCAAAAACTGAATATTTGTTGAGAAGTAATTTT
177 TCTTTAGAAAAAATTAATGAAAGCAACAATCCTACCCTTCAAGAACTTACTTTAAACCGTAC
178 ACATCGCTTATACCGATCAAATTTTCGCAATGAACAAAGCACCACACAGAGGAAGTTTGAAA
179 AAATCGGTAGAACTGTCTCTACTGATTCAGGCAATAAACTGCTTACGTTTCCGGAACAAAAG
180 GCCACAAGAGATAGTAACCCCTTCTCAATCGAATTGACACATGCAACAGTAATTTCCAGCGA
181 CGAAAGTGCCTTAAAAGACACTACTAATCAAGCGATAGCCGAAATGCAACGTATAACTCCTG
182 CTATTGCAAAGACAATCTCGAGAAGAACAGCAAATTGGGTCTGCTCAAACCCAGCTCCTGAT
183 CCGTATGGGGAACCTTCAACGTGGTCCAGAATTTTGACCCCTAATTTGAAAATAATATCGGA
184 ATCATCACCATATTATCCAGTCCATCTAGCTTCACCAAATTCTACCTTTAGTAGAGATCAAA
185 GTGTAAGAAGTGTTGTCATGAGGAGAAGTTCTGTATGTGTTGAAAAACAAAATAGTTTCTTC
186 AGAAATTACGAACATTTCAAAAATATATTGTCGAGAAGAACTATTAAGGTTAAGACATCATG
187 TCCTAGGCTGAGCGTTGACGTAAGTGACAACAAAAGAGAAAATTTATCACAAGAACACCTCA
188 TTTTACCGAACAAATCTAGAGAAAAAGTCAACCGATTTAAAAATTGCCTACATAGAGTTGCA
189 GAAGCGCTCAGAGCTGCCAAAGAAAACTGGGATCAGCACAATCCCAGGAACTCTATTCATTA
190 AGTACTGTTAGTTTTTATTTTGGGACACCCTTGACATATAGTTTAGAACTTTACCTACTTTA
191 CCCTTGTAAAGTTTCTTTCCTTACTTAGCGAAACTGTACTGAATAAGTTGTGTAAGACAGAA
192 TCTTTGATTTTTTCCAGGAATAATCATCACTGCCTACTGCTACAACAGACTAGTTTACTCTC
193 CCTTTTTTCCGTTTTTATTCCGACAAGGGGCATGAACATTCAATACTAAATTCTTTTGTGCT
194 TTTTTTGGGAAGCCTTTTCTAATAAGTACCAGAAGACACATTTTTATGTGACACTACTTTAA
195 GCCTCGAATGTTTTTTTGTGCCCTATTATTTCTAAGCAAAATATTAAACAATCTAGGCCGTA
196 ATTAAAACGCCTATCTTGGTACAGTGTTACAGCGGCTTATACAGTTTATGTTAAACTTAAAA
197 TTACATAAAGTTTCGAGAACTTCCCCTTAAGAAGGGTCCGGGCGCAGGCAACTGAGCCTACT
198 CGGAATAAACATCGATGACCGTTTTTGGAAAAAGGGTTTGTCGCGGCATATAATGGATACTG
199 AAGAGGTCGCCTGAAAACCTGAAAATTTTTTAGCCACCATAAATAATAAATAGTTATATATA
200 TTAAAACATCATTTGCTATTTTATCAAAATTCCGTTGTAGCGTGGTCAATTGTTCAACTATT
201 TCAATTGTTCAGAAAAGTGAAGCTACTTAAAAGTAGAACAAAAAAGAGTTACTCGAACGTTG
202 AACTCTTTGATTATTTAAAGATTTCAGTTTAACAAAAAAAGTAAAGTAACGTGAGAATGTCT
203 GATTCAACAACTTCCCCCACAACCACATTTTCTACTTCATTATCACCAAATGCGACCACATC
204 ATTTTTTTTGTGGTAACATGGGAGTGATGTATACGGCCATGTCTGGATACGAAACAGAAGAC
205 GCACAAGCCTACTGGGCATGCGGTAGAGCATATGAATCAGCTTTTGCAACGTTGACTAAAAA
206 GGTTCCAGGAACCACATTCTCAGCTGATATGCCAACATCTACTTGGCATGGTGTCCTGGATT
207 GCGGCTATTCCTCTTCAATTAACGTGGCAGAAAATAAAAGTTCGCCCATCGATTATTGGAAT
208 TGTGGTAGAACATATGCCAGGAATTATGCTCTGTCCGATGCGCTTTCACTGAAGCCAACGAA
209 TATGTTACAGTACTTTCTATTAGTATTATTTTTTATCTGCATTATACTATGAGAAAAGCTCA
210 TTAATATTTGAGCACTTAGATAAAGTTGCATATGTGTATGAATACATTCTGAAGCAGCCATT
211 AATTATTAATTTATTCAAGGGCATAATCAGTCCAAAATTTATGTTTTTACAATTATTTTTTT
212 TCTCGTTGTTGGTTTCTAAATGACCTAGGAGAATTATGTATAGAGATTAGAAGCTTTTCGAC
213 AATATAGACTTCATAATAACGAAATGATTTTACTTATGTCTGCAATTATTTCCTCATGCCTT
214 CCTGTATTATCGCTTGTTAGTCCTAGTACGTTATTTGCACGCCTGTTCCGACTGAATTCGAT
215 GAGAGTTGGAATCTATGTCACATTTAGCACCGTATAGGCTAATATTCTAATAAGCGTGACAA
216 AAAATTGGAGTGTTTTACTTGTAATTGATGAATATCAATTCAATGCCCCCATAGCTTTTTTT
217 TACGGGTGATTCTTTTTTTATTGTATTTTTCCACCACGAACCGTAGATGGCTTATTGCATTT
218 CTACATGAAAGAATAGTAAATGTGTAGCCTTTGTTACGTGAATAACTTACTTATCCAAGTAT
219 TAAAAGAATTTTGGACCAGATTTCTTCTTCTATTACTATCCCTCTAATTGAAATATGAGTGA
220 TTTTGAGATAATTGTTGGAATTTCATCGTTGTTACAGGTTATAATATTAAATATACAAAATA
221 TGCTGGAAGTTCTTCTTGAGTATATAGGAATTCACAAAAGGAGAGTCGATATATCTACATAT
222 TATGATAATGTTTACTTCTTCTTCCATTTTATATGTTATCATTTATTATCCTATTGCATTAT
223 CAACCTTAGGATTTCTGCTTCCTTTATATGTGATAGCTGTTCCTTAATCATTATGTCACCTT

224 CCTATGCCGTATGTGATAATATACTAGTAACATGAATACCAATAATAAGTGGATGGTACTAG
225 AATTACATTTCAACATAGGCTGAGTTGCTTTGGAAGGCCATGATGTGTATTATAAATTCTGA
226 GTCTTTTCATGGTTCTCAGAAAAGATCGGGCGTGTGGTCTAGTGGTATGATTCTCGCTTTGG
227 GCGACTTCCTGATTAACAGGGGGACAAAGCATGCGAGAGGCCCTGGGTTCAATTCCCAGCTC
228 GCCCCCTTTTTTACTATTGAAAAGTATTCGGTCAGACGTAGCTGATCGATGTAGCATGTGTA
229 CGCATTTCTAGATATCGAGTAATTTCCGCTGATGTATAATATATATATATGTGTGTGTGTGT
230 GTGTTTTTCATCCAGTAGTTTTTTTCTTTAAAAAAAACTATGTATAATATAAAACATGCAAT
231 CTAGATATGCTGGATAAATTTATCTGGTATGACAATTTTCTTGGCAATTTGCTCGCCCCCGT
232 CTCCATGCAGAGAATTTACTTCGTGATTTTCTATTGTTAATTTGACAGGAATACCATTTTCT
233 TTCTCATCTGAATTTGTGATCAAAAATTCGTAGGCTGATCGGCTTGACCCAACAGAATTATC
234 ACCAGTTTTATGTAATGGGAACGTATACCACGTAGGAGATGGTGTAGAGGTTGACCTTGTTC
235 TCATTGCTAATGTGTCTATATTTTGAGTAGGCACTATCCTTATATCAGGGGAAGTTGTTCTT
236 GTGGGAAGTAGATCTACTTCGTTTGCGAAACCTTGTTGATCTGTCGCTATCCCAGTGCTATC
237 TGTTACGGTGGTCTGTGTCGTGATATCACTATTGGAAGTGAAAGTCGTTTGCACAACATTAC
238 CTTTTTCATCAAAGACAGGCAATCTACAAATAGGACAAGTCTGAGAACGTTCCATCCAATTC
239 TTTAAACACGACAAATGAAGTATGTGGCCACAAGGTAACCTTTTGGGTTTCTTGTTTTTATT
240 CTTCCACGTCTGCTGGTTTGGAGAATGTATTAACTCATCCATACAAATGATACAAATATTGT
241 CATCATTTGCAGAATTTTGTAGCTGTTCTACGGTGACAGTGACAAGAGTGTCGTCGAGCTGT
242 TTGTTATTTCTCCAGATTTTCCACAAACTTGTGCCACTTTGATATAGTGCCAAGATATCCCA
243 CACCACATCTTTCAAAAGCATCATAGGCATCCTAAATGGTATTAGCATAGACAAATGAAGTG
244 CCGTTTTTAAGAATCTTGTGAATACGTCAATTGCTTTTTCATACATGAATTTACCCTCCAGG
245 CCGGTAAATTGTCTATCATCATCGTCGTCATCGTCGTCGTCATTCAGCACTGGCTGAGATTG
246 ATCAGACTCAACCGTGTTTTCATCTGTAGGATCGCCATGGACAATATGGTTGTTCTCATTAG
247 ACAGACTTTGTTGTGAGCGATAAAATTCCCAGAAATTCAAACAAGTCTGTAGGAATAAATTT
248 AGCAAATCAATCAAAAGCATGGTAAACTCCATTACTTGTATCAGGTAAAGGGATGTGGATTC
249 AATATCACTCTTTTGGTTTGTATATATGGAGGAGATGCATCGTGTTATTATCTGGTAGTCTA
250 CAACCGCCAATAGTACGAGGTTAAATGAGAATCTACTAAAGATAAGGGTTTTCATTGTGGTG
251 GAATCATTTATTGACTGTAATAAGGCCTCCAGCCTATCCTTTAAAATCCAATGGAAAACTTT
252 CAGATAGAGTAGTAATAGTCCAAAAAATGCCACTGTGAAAAAATACCGTTCGTGGAACAGTG
253 AGGACATAAACAAGGTGTTTATAATGGTAAATGGTAACCTTTCAAAAATGTGCTCATGCTCA
254 ATAAGCCTCAGTTCACCAAATAATAGTTTCGTTAGGAGTTGCCATAGTAAGGTAGAATTTAA
255 TAAGATGAATATCGACAAAACCATTAGATTGAAGCCTTCATTTAGCTTCAGTGTTACTTGCA
256 AAAAGGAAACGCTTGTCTTGGTGGCTGAATACACGCAATAAAATGTGAGCAAATATGTGACA
257 ACTACAAAAATTGCCAACTGTTTCCTTCTATTTTCTGGCACCATATTATTATTTTGTCTTTC
258 TCCCCTTCTCTTACAAATTGCAATTGATAGATGATTTTCTGGTTATAAAAAGGAACCAACAA
259 ATCGGTAGAAAGTCGACAGTATAACTAGTGGTGAAGCTGTTATTAGAAATGGTAGGGGGGTT
260 TTTTTTTTGTCCTGTCGGCACGTATGGAAATTTATCGTCTATTAAACCTTTTCTTTGATTTG
261 ATTTCTACAGTGTACTTTTGTAGGTGTCCTTAATTTCCATTTTTCCGGGTATGCAAAGTTCA
262 ACAAAAAGAAAGCCCTGACCAAATTTCTGCGTACATAACGGCTACTGCGCTATCGTTGGGAA
263 TATTGTGTGAGAGTATATTTTCATGCTATATATTATAGTAAAAACATAAGAGATTAGACGTG
264 AAAGGAGAGATAGATTACATATATGTATGTATATAACTCGGGTACGGAGAGACAAATTTCAA
265 CTTATAATAATGATACAGACGTTAATATGGATAGAATGAGGATACAGGAGCAGGGAGAATTA
266 CGGGAAATGGGAAAGAAAAACTATTCTTCTTATTTCTTACTTCCCTTTTTTTCCACTTTCAA
267 TAATAATGCTTTATTTATATGAGGCAAAACACCACCTGAAGCAATTGTAGCCCTGATCAAAG
268 AATCTAACTCATCGTCACCTCTAATGGCCAATTGCAGATGTCTTGGGGTAATCCTCTTGACT
269 TTAAGATCCTTGGCTGCATTACCAGCCAACTCCAGCACTTCAGCAGTCAAATATTCCAACAC
270 AGCAGTCAAATAAATGGCAGCTTTGGATCCTACTCTTGTTCGGCCGGTAGCGTGCCTTTTCA
271 GGTAACGCTTAATCCTACCGACAGGAAACTGCAAGCCAGCCCTTGCTGAAGAAGATTGAGAT
272 CTCAATGAACCACTGTCTTTAGCGCCGGATTTACCTTTACCTCCATGAGCTTTTCCTGACAT
273 TATGTTAAGTTATTTTTACGTGCGGCTATAGTGCGAAATTGAATTTAACGATATTTTTTCCG
274 ACTGAAGTATAATAGAGTCCGCAGTGTTTTTTAGTTGGGGGTTGTGACTCTATTACACTGGA

275 TAATAGTTCAGCTGTTAAATAGAGCTGTATGCTTTATTTGTTGTTTTTTTTGTTTTGTTGTT
276 TTCGTGCACGGAGCGAGCGCGGCGGGTTACGGAACACCAGCGGCCAGTTCTACGAGTAGTAA
277 GTTACGGGCGTAAAGTTACTGTGTGCTAATCTAGCATGGATGCATTCAGTAAACCTTTTCAA
278 TTGAGTCTAACGTTCTCTAAAGAAAGAAGAATACTGCTATTTTAGGTATGTGCTTACTACCG
279 CTTGTGCGGACCTGCCCTGATTCCAAGGTAAGAGACAGTGATTCTTGTCCTAAGCTCTTTTC
280 TCCGGGGCGGGCGGGCGGCTGGCAGACATATTGATTGCGGGTAACACCTCCGAGGTTAATTA
281 ATCAAGCAACGTGCGGAGTTTTATCACATGGATGTTTCTTATCACCGGGTGCGCGCCACGCA
282 ATATATAAAACGGTAAACATACGCTGTTGCAAGGCAAGACACAACAGCGGCAAACACTTCAG
283 AGAATACAGACGATAGTCATCAGCACTCCTAATAATCGTATAATGAATACTATACGAAGCTT
284 TGTGCGTTGCAAAAAGAGCATACCAGGCGAAGAGAGCGCAGTTAGTAGGTGATCGGGAAATT
285 GCTGAGTACAGTTGTTTGCCTCCTGTGATACCGCGTCACATGACCATTTAAAAGTGTCCATA
286 ATGTCATCGGTAACGTCACACTCACATGAGTTTTCCTGACTTCAGCCTCCTTCATCTCGCGT
287 TAATACGCCAGCATCGCGCACCATATTACTGCCCGCGTAATCCAATTCCTCCTTTCGTTCTC
288 AATGACTAAAACGCAAATCATTTTTCCTTTTTTTTTGAAACCGGTGGACGCCCCATAGAACT
289 TCCAAATCAGGAAATACCGCAACTTCATCCGTTTTCGTTTCGCTTTAGTCCTCCACGAAAAT
290 ATCTTGCCGGTTCCGCGAATGTTGTACTGGTTCTTCTGTGCAATGAGCCTTTTTCTCTTCTT
291 TCTTTTTCCCTGCTTTTAAGATGGCTTGGCTATGGAATTACTCAATTTGGCACTTCTTTTGC
292 AAGATCCAGACTTGCAGGGATTCTATCACCATTATTACCACACTACTATGTTATTGTTACGC
293 TTTTCATCCACTTTTATATAAGTTATACGTGGTTTTACGAGTTCTAATATTTCACTTTTCAG
294 CTTGGAATACGCCGTTTTAAGAGGACTAAAAGCAAGGAAATTACACACATAATAATATAAGT
295 AATGATACGTCCATTATGTTCAAAAATTATTATCAGTTACATATTCGCAATTTCTCAGTTTC
296 TACTGGCCGCTAATGCGTGGTCGCCCACAGATAGTTATGTTCCTGGCACCGTGTCGTGTCCC
297 GATGACATAAATCTGGTAAGAGAGGCTACGTCTATATCTCAGAATGAGAGCGCATGGTTGGA
298 AAAGAGGAATAAAGTCACTAGTGTAGCTTTAAAAGATTTCTTGACTAGGGCTACTGCAAATT
299 TTTCAGATAGCTCAGAAGTTTTGTCGAAGCTATTTAATGATGGCAACAGCGAAAACCTGCCG
300 AAAATTGCTGTCGCCGTTTCAGGTGGGGGCTATCGGTCCATGCTAACAGGTGCGGGTGTTCT
301 AGCAGCAATGGATAACAGAACTGAAGGTGCTTATGAGCATGGGCTGGGTGGACTTTTACAAA
302 GCACAACATATTTATCTGGTGCCTCGGGCGGCAACTGGCTAGTTGGTACATTAGCCTTGAAC
303 AATTGGACATCCGTGCAAGACATTCTTAATAATATGCAGAACGACGATTCTATTTGGGATTT
304 GTCAGATTCTATTGTTACCCCCGGCGGCATTAATATATTCAAAACAGCCAAAAGGTGGGATC
305 ATATCTCTAATGCTGTCGAATCTAAGCAGAACGCCGATTACAATACTTCTTTGGCCGATATT
306 TGGGGCAGAGCCTTGGCGTATAATTTTTTCCCTTCTCTAAATAGAGGGGGTATAGGCCTGAC
307 TTGGTCTTCCATTAGGGATTTCCCAGTGTTTCAAAATGCTGAAATGCCTTTTCCAATTTCTG
308 TTGCGGACGGTAGGTATCCTGGAACAAAAGTCATCAATTTGAACGCAACGGTTTTTGAATTT
309 AATCCCTTTGAAATGGGATCCTGGGATCCCTCTTTGAACTCTTTTGCCAACGTTAAGTACCT
310 TGGAACGAATGTCTCCAATGGTGTACCATTGGAAAGGGGAAAATGTACCGCAGGCTTTGATA
311 ATGCAGGTTTTATCATGGGTACTTCCTCCACCTTATTTAACCAGTTTCTTTTGAGAATAAAC
312 TCCACTCACTTACCTAGTTTCATCACAAGATTAGCAAGATCAGATCTAAATCTCTGCTGTAA
313 TTTACAGCGGCATAAGAAGGCACGTCCCCTGAAAGTAGGCTAGCCAGAAAACAAAAAGGCCA
314 GAATCGGCGCGGAACTTCTCGTATCGAGAGCAAATGAGGGGTGCGGCACACAGTAGGAAAGG
315 GTAACTGATTAACATACGTACACCCCGGCATAAAAGCTTCTGCAGGCTTTGTAGGCAAGAGA
316 CACTGCGGAGAATAAAAGGTTGTGACGCCCAATACCTTCGCCGGACTCCTTCTTGGCGTTCA
317 CAAAAGACGATGCTGCAGCATTTTTTTCTTGCGTCGAACTCGCGCCCACAGCAACGAGACGG
318 TTCCGTTCCTTCCCGTAATGTTGCTTTGATTAACAAAATTCTCGTTTCCTCAGTTCTGTGAG
319 CCCGTTTGTCAGCAGAGGAATCCCAATGAAGAAAATTTTTTCTGGAAAGTAAAAGCGAGCGA
320 AAAGTTTAAGGTTCAGATTGCATCACTATTCGTTCAGTTGATAGCGAATGTAGACCGCTAGC
321 GGCTTGATTAGATAAATAGAGTGGGTTAAAAAAAAGGAAAAAAAAAATGTGAATTGCACATG
322 CCCAGCGCGCTCGCAAAACATATAAAGTTGTCCTCACATGCGGCGTGCATTGTTCATAGCGG
323 GGCAAACATACTTGTGGCTTAATTTGACACATCTACTTTTAATCTTTTCTTGGTCTAGTACA
324 ATGGCTTTTTCCCAAAGTAGAAGGCTTCTTACTCCAACCGTACCCTGTCCTACTTTACTGGG
325 CATTGATTTTTTAATTCTTGTCTTGAGGCATTTCGACGAGATTTTCATTTGAAAAATATACC

TCCGATATGAAACGTGCAGTACTTAACTTTTATTTACCTTTATAAACAAATTGGGAAAGCAA
GGAGATAGATCTGACTGCCGGCCGAGCTTGGCTTGACTTAGTAGTTTCGATGTCCCTTTTCT
TACTTTCCTCATGCAAAATTGGTGATGAATTGAATGATTTTCGTTCATCTACGCTTTTCATG
TCAATCACAATTCTCTGTGACCTGGCACCAATAAGGCTTTTTCGCAGTGATGGCCTTTGCGT
GATTTGATTTTTACCATCGTTATCGTCGCAATCATCATCTCTATTTTGATTGACGTATACCA
TTCCTCTACTTTTTTGATTAACGTGCCGAGGAAATAGTGGTTGTCTATTTGCATTAACTCCG
CTGAATGAAGATGAATATCTAAACTTTTGTAAAACTTTTGTGCACCCCGATATACTCCTCTG
GGATGACCCGGGAAATGAGGTGGTGCTGCTCGTGGGAACTCTTGATTCCGCAGTTTCGCTAT
AATCCTCGCTAGTATCGCCATCACCGTCATCATCTGTATTATAGATTTCGTTACCGATAGTC
TTTCCTTCAGAATTCTTTCGGTCCTCTTCAATGTCACTGACAACTTCAGAAACCTCTTCACT
CAGGTTGTCGTCATCCTCTAAATTGGTTGTAATAAAAGAACTAGGTATTTGTGTAGATAGTT
GAGACTCGGAAACAGGCTCTTCCAAAGATTGTGAGTTTGTGTCCTCTTTAAAATCCTGTATG
CTGTCGCCTTTTTCTTTGAAGTTTTCCACAACAGTCATGGAGGGTTTATTAAAAAATTTGCT
TCTTGTATCCTTCAACGTTTCTTGGACTACATTGGCATTACTTAATTTCCTACGCTTAATTG
CCATTGTCAGTTTGTCTTCCGCTTGTCTTAGACTATGGATGTTATTTGCAACTTTTGGGTTA
TGATCCAATTTTGTCATTTTTTGGAAGAACGATTCTATTGGTTTGACTTTGGCTTCAGAATT
AGTAGTATTAGTTTTCCCGAACTCTATATTTGACTTGGATGCCAGCTGTAATTTGTGCTCTC
TGTTGGCTAGAGGTTGGTGAAAATCATATGGATTCAAGTCACCTTGAGCAATTTTTAAATGC
AAGTGGTGATCTATATCGTCGTCGAAATGAACAATTTGCTTTTTCTGAGTTTCTCTGTGTAT
GACAAAACCAATGCACGCGTAGAGTCTTTTTCTTTTACTTTCGGTATCTTTCAAGTACAGCG
GAATTTCATTCAAGCTCACTATTTTCTTCCGAATAGGGCAAAATACCCTTTGGAATTGAAAT
GCTAAAACTGCAGCTTCATATTCATTAATGTATGTGTCTGGTATCATCAGTTTTCCCTCCCG
CTGTATACTCAGAATTATCCTTTCAATAGTATTGAATCTTCTAACTAATTTCATTGCGGTAA
TCAGGCCAACCTTGGGAATTCCATTTGTATAGTCACAACCGGATAAACAAACCATTGTTATG
ATTTCTTCATTAGTTAAGGATCCCAACGGAAACTTTTTAGGCAGTTTAATAAAATTATCGCG
ACATATTTCTAAACATTCTCCATAATCATTCAGCTTCGTAATGAGACGTCGACATCCGAAGA
CGAGGAGGTCAGAATCTTCGGATATTATTCCTTGCACAATGTTTTTCTGTTCTAAATATACC
ATTTGAGAGTCAGCCTCAAACGGAGCCACTATGTACCGAATACCGTTTAGCTTACAGTAGCA
TATGATACATTTTGCCATTTCAGGCGTTATGTCGACACATTTTTGAAAATAGTCCATAGCAT
TTTTCTTTTCGCCACAGGCCCACAGTCTTTCAGCTATGGCTTTGTTTTCTTTTCTCTTATCC
CTTCTTTTAGATTCAGTAGACTTTTTAACTGGAATGGCATCACCATCGAAGACCAAATACGG
TTCAACTTTAAAGGTTTTCAATAAACTAAATCTTTTTATGAAAAACTGCAGGTACTTATCAG
TTGGTTTTCCCATTGCAAGTTCATAAGCACAAGAGCAGGCTGCTCTATGTAGCCATGCATAG
CCATCAATGGCTAACACTTCTCCTTCATACCTACGTAGTGATACTGGATTCTGTATGGGCTT
TAACTGAGGAAGAAGACCTTGGATACCCATTCCTTTCTACGCCTTTCAGTTTTTTCGAGCTC
CTTTTATTTTAATGTGGTCCAAAAAGCAATATCAAATATAATTCTATGAGAGCATATAGATA
TACCTGAAGGAGCAAGGACCTGATATTTTTTGTTAAGGTTTTTTACACGGGCCAGTAATAC
AAATTTCTCGAAAGATCCTTTTGCCTTCTATCAATCACGCAAATTTTTGTTGTGCACAAAAC
TAGTTATTGTACTTCCCGCCTTTGCTAAAGACGCGTAAGAAAAAAAAAGTACAAATAATGCCC
TATAAAGAAAAAAATTTAAATAAAACGCGAACTTAGTTTGGACGTAATACTTCTCCTTCTCG
GGCCGGATATTCGTATAGCTCGTAGTAGTCAACCTCAGTTCCACCCAACATAGCTTTTGAAG
TATCCGAAGAATAGAAAAATCGTTGCCACAAGGGCGCTGTTACATCACTATTAAGCCAAAAA
TCCGTCAAATTCGTCTTCCAGTGATTTGTCACTAAGAAACTTCGAAGCAATGAACTGTGATG
TTGTTGTGGAACCTGTCCATGCAAAATGTGGGTTATACCTACTATTTCCTTAATAGTAACGA
GCCATTGATTAATACCCATCATTGAAAAGCAAGCGCTAAGAATGGCGCGGCGCCTATTTCTT
GGTTCCAAACCATCAATAGAAATTGTGTGAGTAATGGTGGTCGCACTTGGACCTTCAGGAAT
CATATCACTATCTTCAACACCTTTACTTTCACTGGTATATATAGAGTTTTCGAAGGGTATCA
GATTGATAAATTTTGCCTTGTGCTCCTCCTTGATTAAGATGAACAATTCAGGAGTCGTGATT
CCTTTTAATATCTGGAAGAACTGGACAATCAACATAGCCCCCAGCCAGATGAAGTTTACGCT
AATCCAAATGAATAAAAAAAAGGTAGAGGGATCGTAATTAGAACCCTTACACAAAGTCTCAT
TCTTGAGCAGGGCACATCTTGCATATTCTTCAACTTGTTCGTAAATATAGTTCGTTTTCTTA

377 AAATATGCCAGACATAACCACATGAACAAAAACATGTGATATTGGACCGTCACAGCAAAGAA
378 GACAAAAAGTTTGTGATTTTTTAAGCCAACATCATTATAAATCCATGGACAATAATGGTCAT
379 ATCTGGCCACCAGGGCGCCGCTAAAGAAAGAATATTTACTCCTCAAAGGCTTCCTTTCTAAT
380 GTCTCAACACAAAAGTTTTCCCTATCAAACTTCCCTAAGTCAATCAGTTGTTTTATTGTTTC
381 TTGTATCGAAGTCAAACTATCATCCGTTTTCAGACAACCAGGATCTGACCGTACTAATCTCA
382 GAAATAGTACAACTGTGAGAAAGGAAGTAACCAAAAACTGGACGTTTTTCATGGTATAGTCA
383 GATACACTATATGGATAGAGTTTCTTCGTCCATATATATATTAAAAAACAGAAAGTGGACAG
384 AAAGAGGCCACTGAAAAAGGGAGTTCTAGTCAAAGAAACTTTGTAAGTGTTTTTTCTGGGAA
385 GGCATGGTAATACAAACTTCTTTAATGTGTTGACCATGACAACAGTGACCAGTAGGGAAAGC
386 ATAATGGCTAGCACTGGAGAAAGAACGAGAGAAATTAAGTATATGTAAACCATGAGTAAAAA
387 TGGTGACAAAAAAATTGTGAATTGTGCATGCGAGCTTTTCTTGAACAGTTTGTCTTTCTGAT
388 TTCCAAGCCTGTCATATCCATGATGTTGCAACGCTTGTTCAAAAAGTGATTTGGTTCCCATT
389 CCTTCGGCTAGATCAAAACAGTCTTGCTTCTGGTTATTCCTTTCGTAGAAATTTGCACCGCT
390 AAGTATTAAGTCACATATGACCCTTTGATCACCTCCAACTAGAGCACAATGGAGAGCATTGA
391 ACCCTCTGTTGTCCGTCCATGCAACTGTGGAGCCAAATTTCAATAAAAGTTCTACAGTGAGA
392 AAATCTCCTTGATAAGCAGCCCAAAGTAGGGGTGTTCTGTTATTGTTATCTTTTGAATCGAT
393 ATCGACGTTGTCATTGTTGTTTACGACAAAATAAAGAACATAAACAACAAGTAAAATGTTAG
394 AACTGTAAACGCTAAAATGCATGATGTTAAGACCCTGCTCATCTTTGAGCGTCGGATCAGCG
395 CCATGTTTGAGAAGCAGGTCAACAATGTAGATATTACCGTACCTTGCGGCCCAGTGCAAAGC
396 AGTAGCCCCTCCGGGGCCTGCCGCCTGGTTAGGATTTGCTCCCCTAAGTAATAGGAACTTTG
397 CTACAGAAAACCTATTGTTTATACAGGCCCAGTGTAAGCCGGATAATTCATCAATGCGGTCG
398 TTATTAATATCCACTGCTCCACTTTCGACCACGTCCTTCACCACTTTTAAATCCCCATCCTT
399 AGCGGCTTCAATGAAAATATCAACCACATATTCAGTGTTGGATCCATCGCCTAGAAGTTGCG
400 CTTTGTTATTGTCCGAATCACTTACAGCGTGTTGAGCCACATCAGTGACAACAGCTGCCCCA
401 TTAGAAGTTTTTTTCACGTTCTCATCATCAATAATTGACATACTGGTCATAAGAAACAGTAG
402 AGTATTTCACAGGTAATTAAGATGTTTATCTGACCGATGAACTTTACAACTTGCTAGTAAAA
403 AGTTGGTGGAGTTCAGTTATTATTAAATGAAAAATGCAAAGTTTTGACGCTTTCTAATTGTA
404 AACGACAAAGTGAATGATAAAAAGATCGCTGAACGAAAGATGATTCCAAATACTCAAAAATA
405 GCAGCCATACAAACTGGAACTACTAAATTCAACTAAGATTAAGCGATTAAGAAGTCAGGCTC
406 GTTTAACAACAAGCTTGGTACATCCCAGGCTGCACGAAGTCGTAACTTAAAAAAAAATAAAG
407 AAGCGAAAAAGAGAAGATCATTTCCTTTGGCATCTATTGTTTGCGAAGCCATCGGACCGAGG
408 CACCTTTGAAAAAAAAAAAAAAGGAGCACTTTCTTCCTTTTTCTCTCACCCAGATTCTTTTAG
409 AACACGTAGGGTACCAGACATAATTTAGAGTAGTGTGATACTTCTGGTGTAAAAATAGAACA
410 AAATATATAAAGGGCACCACTACCAGAGGATATTCTCTGGCTACGAAAATCTTACATAGTTC
411 TTGTGTGTTCATTATCAGGGTGCTGTTGTATTTCTTAGTGAAAATTAAGAAGAGTTAAACAT
412 GGCGATTCTTTCCATATTAGACTTTAATTTTAGCAAGGATAACGGGAATTGCCGCTGAAAAC
413 TCTGGTCCTGAATCGCCTCTCTTTAAAGACATATTTAAAGCATCCTTTAATTTTTGATTTTC
414 TTGCAGCAACGGATACACCTCGTTAGTGCCATTATCCGGTATCACTTCGAATAATCCAAAAA
415 AAAGCATAAGTAACCTTTGTCTTAACTCAATGTCATCAATTTGATCCGCAAATACCTGAATA
416 GCATTTTCGATTAGTTCTTTCTTGGTTAACAGTTCCTTAGCTATCAATGGAATCGTGGTGGC
417 AATGTTAGCAAAGATAGCAGCCACTGCTCGTTGAGACTCAACGTCGGATAACTGAAGAAGTT
418 TCACCAATATATTAAAATTTCTCAAACTCTGAGGATTTTCCAAGTTAAAGAACTTAGCAGCA
419 ATCGTTAATGGATGACTCATCATATTACTTATGAGCTCTAATGTTGATCTTTGCAAAGGAAC
420 GTTTTCGTCTAGCATCAGATTCTCAATCGTGGACCAATATACTTTTGTAGATACGATATGCT
421 TGCAAACTTCTTCACCATCAGAGGTCTCCGATGATGCCAAGTTCGTCAGAGCTAATAGTGCC
422 TCGTAATTGTCTGTCAGCTTTATTTGTTCGTCGTTATGTAACGGATTGTCATCCACTGGCGT
423 AGACCTTGGCAACAATTCGAACAAGAAGGGGATAGCGTTTAATGCAGAGTATTTTTTAAATA
424 TCAAACCCGGATTAGTGAATATTAGCATTCTTGTTAATGCCCGACACCCTAGAATCCGTATT
425 GGCTCCCCTATGTCCTGCTTATTGGCCAAATATTCTAAGATGATAGTTGTACCACCTTGTGA
426 AATGCATTGAGGAATGAAGTTTTTCGAGCGGGTTATGTTATATATAACCCTAACCACTTGTT
427 GTTTACAATTGGGGCTCAAATTGTGCATTTCCCTCTTCAAAAAGGAAATCAGCTCGGTCCTT

AATATGTACTTTTCGTTAAACAAAAGAATATCTTCCTTTGATTCTTTCTCTGCGCCTACCTT
GTCTGCGCCTGGGCCCTTCAAATCTGCGTAATTTTTTAAGTCATTGATAGACTGTGAACTAC
CATTGGATTCCTCTGGTAAAGTGGACAAATTTGCCATGATAACTAAGAGACCATATAAACAG
TGCGTCATTTTCTGGCTTTTAATCATAGTTAATAGAATTTCAGTAAAGCTCTCGTTACTCCT
AATCATAATCTTAACCGAGGCCTTTAGACTTAAATATGCCAGCGCTTCAACGGACATTTCAA
CTTTAGGAACCTCCTCGAGCTTAACCGCACTTTCATTGACATTTTCAATTTTTGGCATGATA
CGTCTTGAAATAGCGTTTATAAAGATTTCGGACAGTTGCTTCAAGTTTATACACGTTAATTT
TGTGAAAGACCATGTCTTAACTAACACCAATGCAGAATATATTTGTACATCTTCCACGTTTA
GTGATCTTTCCAATAGTTGAAGGTAATTTTCTGTAATATACGTTCTCATGGTCTCATCAATA
CACGCGGAAGATAATAACCGCAGCAGTTCTTTTGTAAACTGTAAATCCTGCTCTTCGAATAC
TCTCTTCTTAAAAAGCTTACTTAGTCCTTTGGTCAAAAATATTTCAGAACAAAGTGTTGTCA
ATGAAGGATATAATTCTGATAGGGTTTTGACTATGATGGACAATGGATCATTTCCAACGTCT
ATTTCTGCCTCCACAATTAAGCTACTCATGAAATCGACCACTGCTTTATCAAAGTCTTTTTG
GAAGCTTGATTGTAATTCAGCAAAAATTATAAGCATCATCGATTTTACTTCATCTTCGCTTA
TGCGCAAAATAAGTTCCTTCACTAAAAATCGCACCTCCTTGAAATTGTACTTGAACTTATTC
AATAGTTGTAGAATTATGCTCAGTAAATACGTGACGTGGGTGTCCTCGCCATAGTGTATCCT
AATTTGCAGCTCATTTAACAATGCAATTGTTGCGTCTTTATTGGAAAAGCAACCAACGAAAA
CGTGGATGGATGATCTTGATAATATTTCAAATAAATTGAGTGCGAAGGGAATATCATTCTGT
ATAAGTTTAGATAAATGTACGCGGGACTCAGAATGGTCTTGGTAAGACCTGGTCAATATTTC
TTCGATTTTCTGACGATCTTTAAGGGCTTCACTATTCGTTAAATCAGCCGGCATTTTCCCAG
ATTCAGATTTTTGCCTTAGCTGGAAAATTGTATTTATAGCATCATTGTATTTTTGGACATCA
GGGGTGGATTTCAAAGTCTTGTCAAATGCAGCGCAGAGGCTATCAATAGTAGAGCTATCGAT
TGGATCATTCCCTTTCTCACACAGTGGCATCGTGATTCTAATTTTTTAGTAATCTTTTCATG
TGAGTAGTTTAGATAAAACGTCTTCACTATTAAGTAGAAGATTCGTAATTGGATTTAAGCCT
TCGTGCCTTATCGCGGCTGATAATCATCATGATCGACGATGAACTACTGCGCCAGAGTATAA
TCCCCTATAAAACAGGCCTGCTATCACTCCAAAGAGAATGCTTTTATAGAAGAATATAACGT
AAATTACTACAATAATTGTGTTGAGATGTCGGAAGACGAATTTTTTGGTGGTGATAATGAAG
CCGTTTGGAACGGTTCCAGATTCAGTGATTCACCTGAGTTCCAAACGTTGAAAGAAGAGGTT
GCTGCAGAGTTATTTGAAATAAATGGGCAAATAAGCACGCTGCAGCAGTTTACCGCGACACT
TAAGTCATTTATAGATCGGGGAGATGTTAGTGCGAAAGTTGTGGAAAGAATTAATAAGAGAT
CTGTGGCAAAGATAGAAGAAATAGGCGGGCTCATTAAAAAGGTCAATACATCAGTGAAAAAG
ATGGATGCGATTGAGGAAGCTAGCCTGGATAAGACTCAAATAATAGCGAGAGAGAAACTTGT
GAGGGATGTCAGTTACTCTTTTCAAGAGTTTCAAGGTATTCAGCGGCAGTTTACCCAAGTAA
TGAAACAAGTTAATGAAAGAGCAAAAGAATCTCTTGAAGCAAGTGAGATGGCAAATGATGCT
GCTTTATTAGATGAAGAACAAAGGCAGAATAGCTCAAAAAGTACTCGAATACCAGGCAGCCA
AATAGTCATTGAGAGAGACCCGATAAATAACGAAGAGTTTGCTTATCAGCAAAATCTTATCG
AGCAAAGAGACCAGGAAATCAGCAATATTGAAAGAGGTATAACGGAACTGAACGAAGTTTTT
AAAGATTTGGGAAGTGTTGTTCAACAGCAGGGTGTGCTAGTTGACAATATTGAAGCAAATAT
TTATACAACGTCAGATAACACTCAATTGGCTTCAGACGAGCTAAGGAAGGCCATGCGGTACC
AAAAACGTACGAGCAGATGGAGGGTGTATTTGTTGATTGTGCTTCTCGTAATGCTTCTTTTT
ATTTTTCTCATTATGAAATTGTAAACCAAACCAACACACGTCGTCAAATATCATATACATAA
TATATACACTTAAAACAGATTCGTGTTAATGAACTTGTTTTGCACTTTCACTGTACAATGCA
TGGGCTCTCTCATAATATTTCTACGCCGTCGCATCTTAACCAGCCGCAAAGAGAGAGTATAT
TCCATAATTCAGTTTTTCGCCTTTCCTTGCCCTTCTGAAATGTTGTTAATATCATCATACGT
ATTTAGAAAAATAAACAAATAGGGGTTCCGCGCACATTTCCCCGAAAAGTGCCACCTGGGTC
CTTTTCATCACGTGCTATAAAAATAATTATAATTTAAATTTTTTAATATAAATATATAAATT
AAAAATAGAAAGTAAAAAAAGAAATTAAAGAAAAAATAGTTTTTGTTTTCCGAAGATGTAAA
AGACTCTAGGGGGATCGCCAACAAATACTACCTTTTATCTTGCTCTTCCTGCTCTCAGGTAT
TAATGCCGAATTGTTTCATCTTGTCTGTGTAGAAGACCACACACGAAAATCCTGTGATTTTA
CATTTTACTTATCGTTAATCGAATGTATATCTATTTAATCTGCTTTTCTTGTCTAATAAATA
TATATGTAAAGTACGCTTTTTGTTGAAATTTTTTAAACCTTTGTTTATTTTTTTTTCTTCAT

479 TCCGTAACTCTTCTACCTTCTTTATTTACTTTCTAAAATCCAAATACAAAACATAAAAATAA
480 ATAAACACAGAGTAAATTCCCAAATTATTCCATCATTAAAAGATACGAGGCGCGTGTAAGTT
481 ACAGGCAAGCGATCCGTCCTAAGAAACCATTATTATCATGACATTAACCTATAAAAATAGGC
482 GTATCACGAGGCCCTTTCGTCTCGCGCGTTTCGGTGATGACGGTGAAAACCTCTGACACATG
483 CAGCTCCCGGAGACGGTCACAGCTTGTCTGTAAGCGGATGCCGGGAGCAGACAAGCCCGTCA
484 GGGCGCGTCAGCGGGTGTTGGCGGGTGTCGGGGCTGGCTTAACTATGCGGCATCAGAGCAGA
485 TTGTACTGAGAGTGCACCATACCACAGCTTTTCAATTCAATTCATCATTTTTTTTTTATTCT
486 TTTTTTTGATTTCGGTTTCTTTGAAATTTTTTTGATTCGGTAATCTCCGAACAGAAGGAAGA
487 ACGAAGGAAGGAGCACAGACTTAGATTGGTATATATACGCATATGTAGTGTTGAAGAAACAT
488 GAAATTGCCCAGTATTCTTAACCCAACTGCACAGAACAAAAACCTGCAGGAAACGAAGATAA
489 ATCATGTCGAAAGCTACATATAAGGAACGTGCTGCTACTCATCCTAGTCCTGTTGCTGCCAA
490 GCTATTTAATATCATGCACGAAAAGCAAACAAACTTGTGTGCTTCATTGGATGTTCGTACCA
491 CCAAGGAATTACTGGAGTTAGTTGAAGCATTAGGTCCCAAAATTTGTTTACTAAAAACACAT
492 GTGGATATCTTGACTGATTTTTCCATGGAGGGCACAGTTAAGCCGCTAAAGGCATTATCCGC
493 CAAGTACAATTTTTTACTCTTCGAAGACAGAAAATTTGCTGACATTGGTAATACAGTCAAAT
494 TGCAGTACTCTGCGGGTGTATACAGAATAGCAGAATGGGCAGACATTACGAATGCACACGGT
495 GTGGTGGGCCCAGGTATTGTTAGCGGTTTGAAGCAGGCGGCAGAAGAAGTAACAAAGGAACC
496 TAGAGGCCTTTTGATGTTAGCAGAATTGTCATGCAAGGGCTCCCTATCTACTGGAGAATATA
497 CTAAGGGTACTGTTGACATTGCGAAGAGCGACAAAGATTTTGTTATCGGCTTTATTGCTCAA
498 AGAGACATGGGTGGAAGAGATGAAGGTTACGATTGGTTGATTATGACACCCGGTGTGGGTTT
499 AGATGACAAGGGAGACGCATTGGGTCAACAGTATAGAACCGTGGATGATGTGGTCTCTACAG
500 GATCTGACATTATTATTGTTGGAAGAGGACTATTTGCAAAGGGAAGGGATGCTAAGGTAGAG
501 GGTGAACGTTACAGAAAAGCAGGCTGGGAAGCATATTTGAGAAGATGCGGCCAGCAAAACTA
502 AAAAACTGTATTATAAGTAAATGCATGTATACTAAACTCACGAATTCGCCAAAAGTTGGCCC
503 AGGGCTTCCCGGTATCAACAGGGACACCAGGATTTATTTATTCTGCGAAGTGATCTTCCGTC
504 ACAGGTATTTATTCGCGATAAGCTCATGGAGCGGCGTAACCGTCGCACAGGAAGGACAGAGA
505 AAGCGCGGATCTGGGAAGTGACGGACAGAACGGTCAGGACCTGGATTGGGGAGGCGGTTGCC
506 GCCGCTGCTGCTGACGGTGTGACGTTCTCTGTTCCGGTCACACCACATACGTTCCGCCATTC
507 CTATGCGATGCACATGCTGTATGCCGGTATACCGCTGAAAGTTCTGCAAAGCCTGATGGGAC
508 ATAAGTCCATCAGTTCAACGGAAGTCTACACGAAGGTTTTTGCGCTGGATGTGGCTGCCCGG
509 CACCGGGTGCAGTTTGCGATGCCGGAGTCTGATGCGGTTGCGATGCTGAAACAATTATCCTG
510 AGAATAAATGCCTTGGCCTTTATATGGAAATGTGGAACTGAGTGGATATGCTGTTTTTGTCT
511 GTTAAACAGAGAAGCTGGCTGTTATCCACTGAGAAGCGAACGAAACAGTCGGGAAAATCTCC
512 CATTATCGTAGAGATCCGCATTATTAATCTCAGGAGCCTGTGTAGCGTTTATAGGAAGTAGT
513 GTTCTGTCATGATGCCTGCAAGCGGTAACGAAAACGATTTGAATATGCCTTCAGGAACAATA
514 GAAATCTTCGTGCGGTGTTACGTTGAAGTGGAGCGGATTATGTCAGCAATGGACAGAACAAC
515 CTAATGAACACAGAACCATGATGTGGTCTGTCCTTTTACAGCCAGTAGTGCTCGCCGCAGTC
516 GAGCGACAGGGCGAAGCCCTCGAGTGAGCGAGGAAGCACCAGGGAACAGCACTTATATATTC
517 TGCTTACACACGATGCCTGAAAAAACTTCCCTTGGGGTTATCCACTTATCCACGGGGATATT
518 TTTATAATTATTTTTTTTATAGTTTTTAGATCTTCTTTTTTAGAGCGCCTTGTAGGCCTTTA
519 TCCATGCTGGTTCTAGAGAAGGTGTTGTGACAAATTGCCCTTTCAGTGTGACAAATCACCCT
520 CAAATGACAGTCCTGTCTGTGACAAATTGCCCTTAACCCTGTGACAAATTGCCCTCAGAAGA
521 AGCTGTTTTTTCACAAAGTTATCCCTGCTTATTGACTCTTTTTTATTTAGTGTGACAATCTA
522 AAAACTTGTCACACTTCACATGGATCTGTCATGGCGGAAACAGCGGTTATCAATCACAAGAA
523 ACGTAAAAATAGCCCGCGAATCGTCCAGTCAAACGACCTCACTGAGGCGGCATATAGTCTCT
524 CCCGGGATCAAAAACGTATGCTGTATCTGTTCGTTGACCAGATCAGAAAATCTGATGGCACC
525 CTACAGGAACATGACGGTATCTGCGAGATCCATGTTGCTAAATATGCTGAAATATTCGGATT
526 GACCTCTGCGGAAGCCAGTAAGGATATACGGCAGGCATTGAAGAGTTTCGCGGGGAAGGAAG
527 TGGTTTTTTATCGCCCTGAAGAGGATGCCGGCGATGAAAAAGGCTATGAATCTTTTCCTTGG
528 TTTATCAAACGTGCGCACAGTCCATCCAGAGGGCTTTACAGTGTACATATCAACCCATATCT
529 CATTCCCTTCTTTATCGGGTTACAGAACCGGTTTACGCAGTTTCGGCTTAGTGAAACAAAAG

530 AAATCACCAATCCGTATGCCATGCGTTTATACGAATCCCTGTGTCAGTATCGTAAGCCGGAT
531 GGCTCAGGCATCGTCTCTCTGAAAATCGACTGGATCATAGAGCGTTACCAGCTGCCTCAAAG
532 TTACCAGCGTATGCCTGACTTCCGCCGCCGCTTCCTGCAGGTCTGTGTTAATGAGATCAACA
533 GCAGAACTCCAATGCGCCTCTCATACATTGAGAAAAAGAAAGGCCGCCAGACGACTCATATC
534 GTATTTTCCTTCCGCGATATCACTTCCATGACGACAGGATAGTCTGAGGGTTATCTGTCACA
535 GATTTGAGGGTGGTTCGTCACATTTGTTCTGACCTACTGAGGGTAATTTGTCACAGTTTTGC
536 TGTTTCCTTCAGCCTGCATGGATTTTCTCATACTTTTTGAACTGTAATTTTTAAGGAAGCCA
537 AATTTGAGGGCAGTTTGTCACAGTTGATTTCCTTCTCTTTCCCTTCGTCATGTGACCTGATA
538 TCGGGGGTTAGTTCGTCATCATTGATGAGGGTTGATTATCACAGTTTATTACTCTGAATTGG
539 CTATCCGCGTGTGTACCTCTACCTGGAGTTTTTCCCACGGTGGATATTTCTTCTTGCGCTGA
540 GCGTAAGAGCTATCTGACAGAACAGTTCTTCTTTGCTTCCTCGCCAGTTCGCTCGCTATGCT
541 CGGTTACACGGCTGCGGCGAGCGCTAGTGATAATAAGTGACTGAGGTATGTGCTCTTCTTAT
542 CTCCTTTTGTAGTGTTGCTCTTATTTTAAACAACTTTGCGGTTTTTTGATGACTTTGCGATT
543 TTGTTGTTGCTTTGCAGTAAATTGCAAGATTTAATAAAAAAACGCAAAGCAATGATTAAAGG
544 ATGTTCAGAATGAAACTCATGGAAACACTTAACCAGTGCATAAACGCTGGTCATGAAATGAC
545 GAAGGCTATCGCCATTGCACAGTTTAATGATGACAGCCCGGAAGCGAGGAAAATAACCCGGC
546 GCTGGAGAATAGGTGAAGCAGCGGATTTAGTTGGGGTTTCTTCTCAGGCTATCAGAGATGCC
547 GAGAAAGCAGGGCGACTACCGCACCCGGATATGGAAATTCGAGGACGGGTTGAGCAACGTGT
548 TGGTTATACAATTGAACAAATTAATCATATGCGTGATGTGTTTGGTACGCGATTGCGACGTG
549 CTGAAGACGTATTTCCACCGGTGATCGGGGTTGCTGCCCATAAAGGTGGCGTTTACAAAACC
550 TCAGTTTCTGTTCATCTTGCTCAGGATCTGGCTCTGAAGGGGCTACGTGTTTTGCTCGTGGA
551 AGGTAACGACCCCCAGGGAACAGCCTCAATGTATCACGGATGGGTACCAGATCTTCATATTC
552 ATGCAGAAGACACTCTCCTGCCTTTCTATCTTGGGGAAAAGGACGATGTCACTTATGCAATA
553 AAGCCCACTTGCTGGCCGGGGCTTGACATTATTCCTTCCTGTCTGGCTCTGCACCGTATTGA
554 AACTGAGTTAATGGGCAAATTTGATGAAGGTAAACTGCCCACCGATCCACACCTGATGCTCC
555 GACTGGCCATTGAAACTGTTGCTCATGACTATGATGTCATAGTTATTGACAGCGCGCCTAAC
556 CTGGGTATCGGCACGATTAATGTCGTATGTGCTGCTGATGTGCTGATTGTTCCCACGCCTGC
557 TGAGTTGTTTGACTACACCTCCGCACTGCAGTTTTTCGATATGCTTCGTGATCTGCTCAAGA
558 ACGTTGATCTTAAAGGGTTCGAGCCTGATGTACGTATTTTGCTTACCAAATACAGCAATAGT
559 AATGGCTCTCAGTCCCCGTGGATGGAGGAGCAAATTCGGGATGCCTGGGGAAGCATGGTTCT
560 AAAAAATGTTGTACGTGAAACGGATGAAGTTGGTAAAGGTCAGATCCGGATGAGAACTGTTT
561 TTGAACAGGCCATTGATCAACGCTCTTCAACTGGTGCCTGGAGAAATGCTCTTTCTATTTGG
562 GAACCTGTCTGCAATGAAATTTTCGATCGTCTGATTAAACCACGCTGGGAGATTAGATAATG
563 AAGCGTGCGCCTGTTATTCCAAAACATACGCTCAATACTCAACCGGTTGAAGATACTTCGTT
564 ATCGACACCAGCTGCCCCGATGGTGGATTCGTTAATTGCGCGCGTAGGAGTAATGGCTCGCG
565 GTAATGCCATTACTTTGCCTGTATGTGGTCGGGATGTGAAGTTTACTCTTGAAGTGCTCCGG
566 GGTGATAGTGTTGAGAAGACCTCTCGGGTATGGTCAGGTAATGAACGTGACCAGGAGCTGCT
567 TACTGAGGACGCACTGGATGATCTCATCCCTTCTTTTCTACTGACTGGTCAACAGACACCGG
568 CGTTCGGTCGAAGAGTATCTGGTGTCATAGAAATTGCCGATGGGAGTCGCCGTCGTAAAGCT
569 GCTGCACTTACCGAAAGTGATTATCGTGTTCTGGTTGGCGAGCTGGATGATGAGCAGATGGC
570 TGCATTATCCAGATTGGGTAACGATTATCGCCCAACAAGTGCTTATGAACGTGGTCAGCGTT
571 ATGCAAGCCGATTGCAGAATGAATTTGCTGGAAATATTTCTGCGCTGGCTGATGCGGAAAAT
572 ATTTCACGTAAGATTATTACCCGCTGTATCAACACCGCCAAATTGCCTAAATCAGTTGTTGC
573 TCTTTTTTCTCACCCCGGTGAACTATCTGCCCGGTCAGGTGATGCACTTCAAAAAGCCTTTA
574 CAGATAAAGAGGAATTACTTAAGCAGCAGGCATCTAACCTTCATGAGCAGAAAAAAGCTGGG
575 GTGATATTTGAAGCTGAAGAAGTTATCACTCTTTTAACTTCTGTGCTTAAAACGTCATCTGC
576 ATCAAGAACTAGTTTAAGCTCACGACATCAGTTTGCTCCTGGAGCGACAGTATTGTATAAGG
577 GCGATAAAATGGTGCTTAACCTGGACAGGTCTCGTGTTCCAACTGAGTGTATAGAGAAAATT
578 GAGGCCATTCTTAAGGAACTTGAAAAGCCAGCACCCTGATGCGACCACGTTTTAGTCTACGT
579 TTATCTGTCTTTACTTAATGTCCTTTGTTACAGGCCAGAAAGCATAACTGGCCTGAATATTC
580 TCTCTGGGCCCACTGTTCCACTTGTATCGTCGGTCTGATAATCAGACTGGGACCACGGTCCC

581 ACTCGTATCGTCGGTCTGATTATTAGTCTGGGACCACGGTCCCACTCGTATCGTCGGTCTGA
582 TTATTAGTCTGGGACCACGGTCCCACTCGTATCGTCGGTCTGATAATCAGACTGGGACCACG
583 GTCCCACTCGTATCGTCGGTCTGATTATTAGTCTGGGACCATGGTCCCACTCGTATCGTCGG
584 TCTGATTATTAGTCTGGGACCACGGTCCCACTCGTATCGTCGGTCTGATTATTAGTCTGGAA
585 CCACGGTCCCACTCGTATCGTCGGTCTGATTATTAGTCTGGGACCACGGTCCCACTCGTATC
586 GTCGGTCTGATTATTAGTCTGGGACCACGATCCCACTCGTGTTGTCGGTCTGATTATCGGTC
587 TGGGACCACGGTCCCACTTGTATTGTCGATCAGACTATCAGCGTGAGACTACGATTCCATCA
588 ATGCCTGTCAAGGGCAAGTATTGACATGTCGTCGTAACCTGTAGAACGGAGTAACCTCGGTG
589 TGCGGTTGTATGCCTGCTGTGGATTGCTGCTGTGTCCTGCTTATCCACAACATTTTGCGCAC
590 GGTTATGTGGACAAAATACCTGGTTACCCAGGCCGTGCCGGCACGTTAACCGGGCTGCATCC
591 GATGCAAGTGTGTCGCTTAGGTGATCGGCACGTAAGAGGTTCCAACTTTCACCATAATGAAA
592 TAAGATCACTACCGGGCGTATTTTTTGAGTTATCGAGATTTTCAGGAGCTAAGGAAGCTAAA
593 ATGGAGAAAAAAATCACTGGATATACCACCGTTGATATATCCCAATGGCATCGTAAAGAACA
594 TTTTGAGGCATTTCAGTCAGTTGCTCAATGTACCTATAACCAGACCGTTTAGCTGGATATTA
595 CGGCCTTTTTAAAGACCGTAAAGAAAAATAAGCACAAGTTTTATCCGGCCTTTATTCACATT
596 CTTGCCCGCCTGATGAATGCTCATCCGGAATTCCGTATGGCAATGAAAGACGGTGAGCTGGT
597 GATATGGGATAGTGTTCACCCTTGTTACACCGTTTTCCATGAGCAAACTGAAACGTTTTCAT
598 CGCTCTGGAGTGAATACCACGACGATTTCCGGCAGTTTCTACACATATATTCGCAAGATGTG
599 GCGTGTTACGGTGAAAACCTGGCCTATTTCCCTAAAGGGTTTATTGAGAATATGTTTTTCGT
600 CTCAGCCAATCCCTGGGTGAGTTTCACCAGTTTTGATTTAAACGTGGCCAATATGGACAACT
601 TCTTCGCCCCCGTTTTCACCATGGGCAAATATTATACGCAAGGCGACAAGGTGCTGATGCCG
602 CTGGCGATTCAGGTTCATCATGCCGTTTGTGATGGCTTCCATGTCGGCAGAATGCTTAATGA
603 ATTACAACAGTACTGCGATGAGTGGCAGGGCGGGGCGTAATTTTTTTAAGGCAGTTATTGGT
604 GCCCTTAAACGTCTGACGCTCAGTGGAACGAAAACTCACGTTAAGGGATTTTGGTCATGAAC
605 AATAAAACTGTCTGCTTACATAAACAGTAATACAAGGGGTGTTATGAGCCATATTCAACGGG
606 AAACGTCTTGCTCTAGGCCGCGATTAAATTCCAACATGGATGCTGATTTATATGGGTATAAA
607 TGGGCTCGCGATAATGTCGGGCAATCAGGTGCGACAATCTATCGATTGTATGGGAAGCCCGA
608 TGCGCCAGAGTTGTTTCTGAAACATGGCAAAGGTAGCGTTGCCAATGATGTTACAGATGAGA
609 TGGTCAGACTAAACTGGCTGACGGAATTTATGCCTCTTCCGACCATCAAGCATTTTATCCGT
610 ACTCCTGATGATGCATGGTTACTCACCACTGCGATCCCCGGGAAAACAGCATTCCAGGTATT
611 AGAAGAATATCCTGATTCAGGTGAAAATATTGTTGATGCGCTGGCAGTGTTCCTGCGCCGGT
612 TGCATTCGATTCCTGTTTGTAATTGTCCTTTTAACAGCGATCGCGTATTTCGTCTCGCTCAG
613 GCGCAATCACGAATGAATAACGGTTTGGTTGATGCGAGTGATTTTGATGACGAGCGTAATGG
614 CTGGCCTGTTGAACAAGTCTGGAAAGAAATGCATAAACTTTTGCCATTCTCACCGGATTCAG
615 TCGTCACTCATGGTGATTTCTCACTTGATAACCTTATTTTTGACGAGGGGAAATTAATAGGT
616 TGTATTGATGTTGGACGAGTCGGAATCGCAGACCGATACCAGGATCTTGCCATCCTATGGAA
617 CTGCCTCGGTGAGTTTTCTCCTTCATTACAGAAACGGCTTTTTCAAAAATATGGTATTGATA
618 ATCCTGATATGAATAAATTGCAGTTTCATTTGATGCTCGATGAGTTTTTCTAAAAGCTTAAT
619 TAGCTGATCTAGACGCGTGCTAGAGGCATCAAATAAAACGAAAGGCTCAGTCGAAAGACTGG
620 GCCTTTCGTTTTATCTGTTGTTTGTCGGTGAACGCTCTCCTGAGTAGGACAAATAGGTCGAG
621 GGTGAAGTACTTGCTGACTTCCTTGAGGAACACATGATGCGTCCTACGGTTGCTGCTACGCA
622 TATCATTGAGATGTCTGTGGGAGGAGTTGATGTGTACTCTGAGGACGATGAGGGTTACGGTA
623 CGTCTTTCATTGAGTGGTGATTTATGCATTAGGACTGCATAGGGATGCACTATAGACCACGG
624 ATGGTCAGTTCTTTAAGTTACTGAAAAGACACGATAAATTAATACGACTCACTATAGGGAGA
625 GGAGGGACGAAAGGTTACTATATAGATACTGAATGAATACTTATAGAGTGCATAAAGTATGC
626 ATAATGGTGTACCTAGAGTGACCTCTAAGAATGGTGATTATATTGTATTAGTATCACCTTAA
627 CTTAAGGCGGGATCGTCACCCTCAGCAGCGAAAGACAGCTGTCGGTCAGAGCGTCATTGCGA
628 AGCTGAGTGTGATCGATGCCATCAGCGAAGGGCCCAAACTCCGAGCGATTAAGCGTTTGCTG
629 GCTGTCACGCCTGCCTGTTGCTTGCTTGGACTTGCGATGTACGTGCTCAGCTGTCTTTCGCT
630 GCTGAGGGTGACGATCCCGCGAGGGCCTATGGAGTTCCTATAGGGTCCTTTAAAATATACCA
631 TAAAAATCTGAGTGACTATCTCACAGTGTACGGACCTAAAGTTCCCCCATAGGGGGTACCTA

AAGCCCAGCCAATCACCTAAAGTCAACCTTCGGTTGACCTTGAGGGTTCCCTAAGGGTTGGG
GATGACCCTTGGGTTTGTCTTTGGGTGTTACCTTGAGTGTCTCTCTGTGTCCCTATCTGTTA
CAGTCTCCTAAAGTATCCTCCTAAAGTCACCTCCTAACGCACATTTCCCCGAAAAGTGCCAC
CTGGGTCCTTTTC

**Supplementary Data 5: Phagemid with *tetA*, pSJ78**

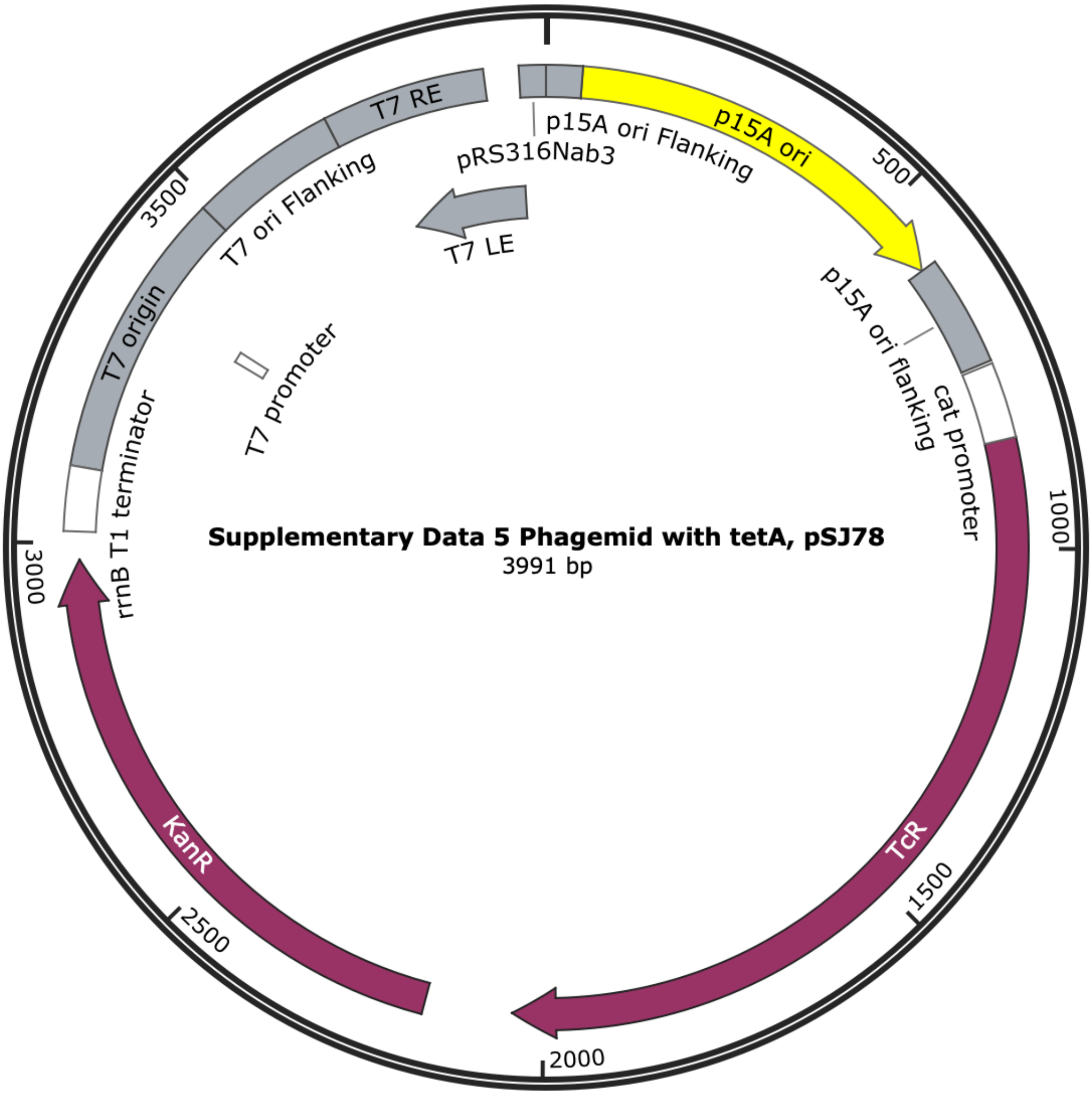


ACTTCGGGCTCATGAGCAAATATTTTATCTGATTAATAAGATGATCTTCTTGAGATCGTTTT
GGTCTGCGCGTAATCTCTTGCTCTGAAAACGAAAAAACCGCCTTGCAGGGCGGTTTTTCGAA
GGTTCTCTGAGCTACCAACTCTTTGAACCGAGGTAACTGGCTTGGAGGAGCGCAGTCACCAA
AACTTGTCCTTTCAGTTTAGCCTTAACCGGCGCATGACTTCAAGACTAACTCCTCTAAATCA
ATTACCAGTGGCTGCTGCCAGTGGTGCTTTTGCATGTCTTTCCGGGTTGGACTCAAGACGAT
AGTTACCGGATAAGGCGCAGCGGTCGGACTGAACGGGGGGTTCGTGCATACAGTCCAGCTTG
GAGCGAACTGCCTACCCGGAACTGAGTGTCAGGCGTGGAATGAGACAAACGCGGCCATAACA
GCGGAATGACACCGGTAAACCGAAAGGCAGGAACAGGAGAGCGCACGAGGGAGCCGCCAGGG
GGAAACGCCTGGTATCTTTATAGTCCTGTCGGGTTTCGCCACCACTGATTTGAGCGTCAGAT
TTCGTGATGCTTGTCAGGGGGGCGGAGCCTATGGAAAAACGGCTTTGCCGCGGCCCTCTCAC
TTCCCTGTTAAGTATCTTCCTGGCATCTTCCAGGAAATCTCCGCCCCGTTCGTAAGCCATTT
CCGCTCGCCGCAGTCGAACGACCGAGCGTAGCGAGTCAGTGAGCGAGGAAGCGGAATATATC
CTAGGTGATCGGCACGTAAGAGGTTCCAACTTTCACCATAATGAAATAAGATCACTACCGGG
CGTATTTTTTGAGTTATCGAGATTTTCAGGAGCTAAGGAACTAAAATGAAATCTAACAATGC
GCTCATCGTCATCCTCGGCACCGTCACCCTGGATGCTGTAGGCATAGGCTTGGTTATGCCGG

TACTGCCGGGCCTCTTGCGGGATATCGTCCATTCCGACAGCATCGCCAGTCACTATGGCGTG
CTGCTAGCGCTATATGCGTTGATGCAATTTCTATGCGCACCCGTTCTCGGAGCACTGTCCGA
CCGCTTTGGCCGCCGCCCAGTCCTGCTCGCTTCGCTACTTGGAGCCACTATCGACTACGCGA
TCATGGCGACCACACCCGTCCTGTGGATCCTCTACGCCGGACGCATCGTGGCCGGCATCACC
GGCGCCACAGGTGCGGTTGCTGGCGCCTATATCGCCGACATCACCGATGGGGAAGATCGGGC
TCGCCACTTCGGGCTCATGAGCGCTTGTTTCGGCGTGGGTATGGTGGCAGGCCCCGTGGCCG
GGGGACTGTTGGGCGCCATCTCCTTGCATGCACCATTCCTTGCGGCGGCGGTGCTCAACGGC
CTCAACCTACTACTGGGCTGCTTCCTAATGCAGGAGTCGCATAAGGGAGAGCGTCGACCGAT
GCCCTTGAGAGCCTTCAACCCAGTCAGCTCCTTCCGGTGGGCGCGGGGCATGACTATCGTCG
CCGCACTTATGACTGTCTTCTTTATCATGCAACTCGTAGGACAGGTGCCGGCAGCGCTCTGG
GTCATTTTCGGCGAGGACCGCTTTCGCTGGAGCGCGACGATGATCGGCCTGTCGCTTGCGGT
ATTCGGAATCTTGCACGCCCTCGCTCAAGCCTTCGTCACTGGTCCCGCCACCAAACGTTTCG
GCGAGAAGCAGGCCATTATCGCCGGCATGGCGGCCGACGCGCTGGGCTACGTCTTGCTGGCG
TTCGCGACGCGAGGCTGGATGGCCTTCCCCATTATGATTCTTCTCGCTTCCGGCGGCATCGG
GATGCCCGCGTTGCAGGCCATGCTGTCCAGGCAGGTAGATGACGACCATCAGGGACAGCTTC
AAGGATCGCTCGCGGCTCTTACCAGCCTAACTTCGATCATTGGACCGCTGATCGTCACGGCG
ATTTATGCCGCCTCGGCGAGCACATGGAACGGGTTGGCATGGATTGTAGGCGCCGCCCTATA
CCTTGTCTGCCTCCCCGCGTTGCGTCGCGGTGCATGGAGCCGGGCCACCTCGACCTGAAGGC
AGTTATTGGTGCCCTTAAACGTCTGACGCTCAGTGGAACGAAAACTCACGTTAAGGGATTTT
GGTCATGAACAATAAAACTGTCTGCTTACATAAACAGTAATACAAGGGGTGTTATGAGCCAT
ATTCAACGGGAAACGTCTTGCTCTAGGCCGCGATTAAATTCCAACATGGATGCTGATTTATA
TGGGTATAAATGGGCTCGCGATAATGTCGGGCAATCAGGTGCGACAATCTATCGATTGTATG
GGAAGCCCGATGCGCCAGAGTTGTTTCTGAAACATGGCAAAGGTAGCGTTGCCAATGATGTT
ACAGATGAGATGGTCAGACTAAACTGGCTGACGGAATTTATGCCTCTTCCGACCATCAAGCA
TTTTATCCGTACTCCTGATGATGCATGGTTACTCACCACTGCGATCCCCGGGAAAACAGCAT
TCCAGGTATTAGAAGAATATCCTGATTCAGGTGAAAATATTGTTGATGCGCTGGCAGTGTTC
CTGCGCCGGTTGCATTCGATTCCTGTTTGTAATTGTCCTTTTAACAGCGATCGCGTATTTCG
TCTCGCTCAGGCGCAATCACGAATGAATAACGGTTTGGTTGATGCGAGTGATTTTGATGACG
AGCGTAATGGCTGGCCTGTTGAACAAGTCTGGAAAGAAATGCATAAACTTTTGCCATTCTCA
CCGGATTCAGTCGTCACTCATGGTGATTTCTCACTTGATAACCTTATTTTTGACGAGGGGAA
ATTAATAGGTTGTATTGATGTTGGACGAGTCGGAATCGCAGACCGATACCAGGATCTTGCCA
TCCTATGGAACTGCCTCGGTGAGTTTTCTCCTTCATTACAGAAACGGCTTTTTCAAAAATAT
GGTATTGATAATCCTGATATGAATAAATTGCAGTTTCATTTGATGCTCGATGAGTTTTTCTA
AAAGCTTAATTAGCTGATCTAGACGCGTGCTAGAGGCATCAAATAAAACGAAAGGCTCAGTC
GAAAGACTGGGCCTTTCGTTTTATCTGTTGTTTGTCGGTGAACGCTCTCCTGAGTAGGACAA
ATAGGTCGAGGGTGAAGTACTTGCTGACTTCCTTGAGGAACACATGATGCGTCCTACGGTTG
CTGCTACGCATATCATTGAGATGTCTGTGGGAGGAGTTGATGTGTACTCTGAGGACGATGAG
GGTTACGGTACGTCTTTCATTGAGTGGTGATTTATGCATTAGGACTGCATAGGGATGCACTA
TAGACCACGGATGGTCAGTTCTTTAAGTTACTGAAAAGACACGATAAATTAATACGACTCAC
TATAGGGAGAGGAGGGACGAAAGGTTACTATATAGATACTGAATGAATACTTATAGAGTGCA
TAAAGTATGCATAATGGTGTACCTAGAGTGACCTCTAAGAATGGTGATTATATTGTATTAGT
ATCACCTTAACTTAAGGCGGGATCGTCACCCTCAGCAGCGAAAGACAGCTGTCGGTCAGAGC
GTCATTGCGAAGCTGAGTGTGATCGATGCCATCAGCGAAGGGCCCAAACTCCGAGCGATTAA
GCGTTTGCTGGCTGTCACGCCTGCCTGTTGCTTGCTTGGACTTGCGATGTACGTGCTCAGCT
GTCTTTCGCTGCTGAGGGTGACGATCCCGCGAGGGCCTATGGAGTTCCTATAGGGTCCTTTA
AAATATACCATAAAAATCTGAGTGACTATCTCACAGTGTACGGACCTAAAGTTCCCCCATAG
GGGGTACCTAAAGCCCAGCCAATCACCTAAAGTCAACCTTCGGTTGACCTTGAGGGTTCCCT
AAGGGTTGGGGATGACCCTTGGGTTTGTCTTTGGGTGTTACCTTGAGTGTCTCTCTGTGTCC
CTATCTGTTACAGTCTCCTAAAGTATCCTCCTAAAGTCACCTCCTAACGCACATTTCCCCGA
AAAGTGCCACCTGGGTCCTTTTC

**Supplementary Data 6: Phagemid with EG pathway, pAN29**

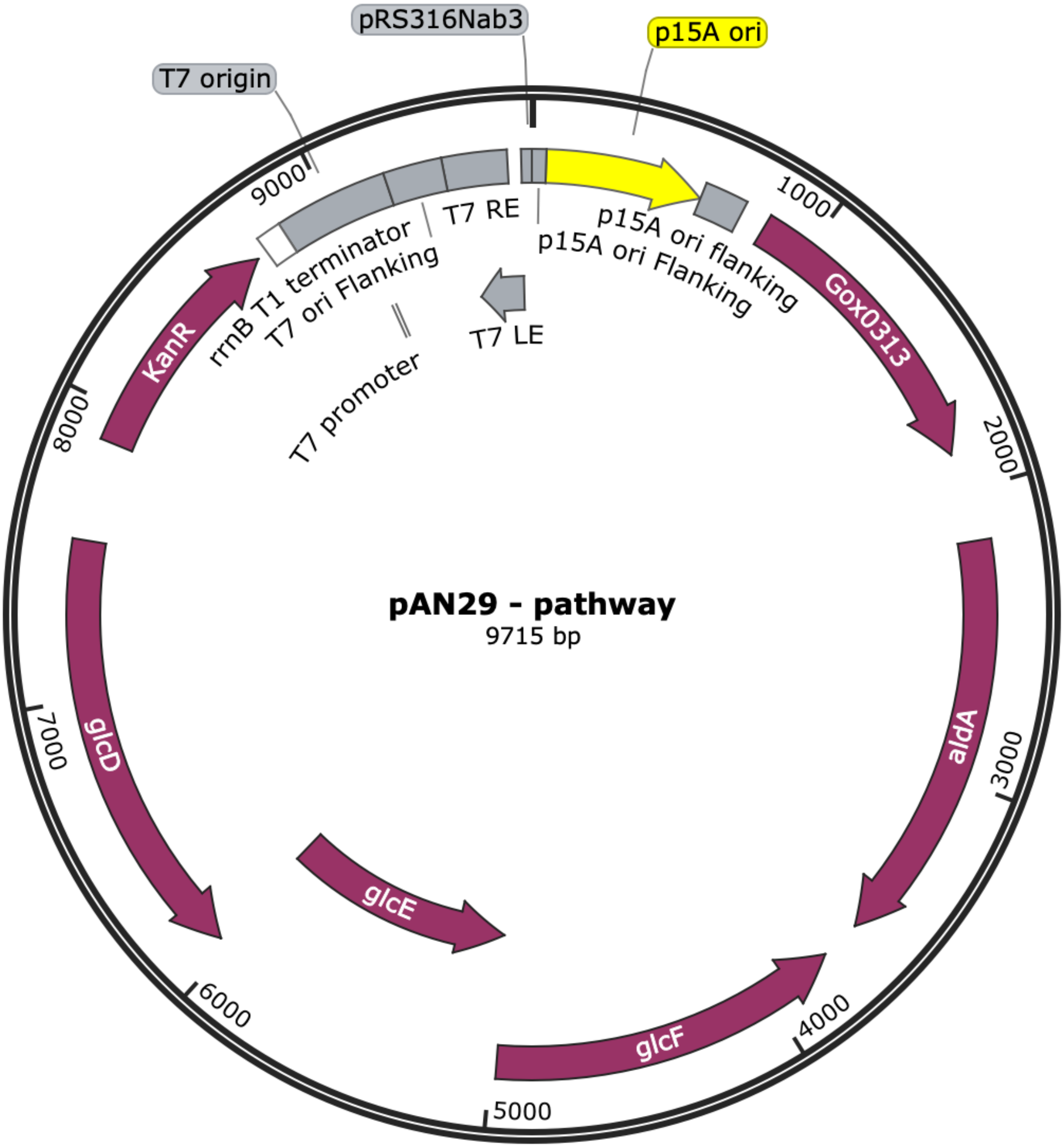


ACTTCGGGCTCATGAGCAAATATTTTATCTGATTAATAAGATGATCTTCTTGAGATCGTTTT
GGTCTGCGCGTAATCTCTTGCTCTGAAAACGAAAAAACCGCCTTGCAGGGCGGTTTTTCGAA
GGTTCTCTGAGCTACCAACTCTTTGAACCGAGGTAACTGGCTTGGAGGAGCGCAGTCACCAA
AACTTGTCCTTTCAGTTTAGCCTTAACCGGCGCATGACTTCAAGACTAACTCCTCTAAATCA
ATTACCAGTGGCTGCTGCCAGTGGTGCTTTTGCATGTCTTTCCGGGTTGGACTCAAGACGAT
AGTTACCGGATAAGGCGCAGCGGTCGGACTGAACGGGGGGTTCGTGCATACAGTCCAGCTTG
GAGCGAACTGCCTACCCGGAACTGAGTGTCAGGCGTGGAATGAGACAAACGCGGCCATAACA
GCGGAATGACACCGGTAAACCGAAAGGCAGGAACAGGAGAGCGCACGAGGGAGCCGCCAGGG
GGAAACGCCTGGTATCTTTATAGTCCTGTCGGGTTTCGCCACCACTGATTTGAGCGTCAGAT
TTCGTGATGCTTGTCAGGGGGGCGGAGCCTATGGAAAAACGGCTTTGCCGCGGCCCTCTCAC
TTCCCTGTTAAGTATCTTCCTGGCATCTTCCAGGAAATCTCCGCCCCGTTCGTAAGCCATTT
CCGCTCGCCGCAGTCGAACGACCGAGCGTAGCGAGTCAGTGAGCGAGGAAGCGGAATATATC

CTTGACAGCTAGCTCAGTCCTAGGTACTGTGCTAGCACCCGTTTTTACTAGTCTAGAAATAA
TAACAAAATGAGGAGGTACTGATATACATATGGCGGACACGATGCTGGCGGCGGTTGTACGT
GAATTTGGTAAACCTCTGTCTATCGAACGCCTGCCGATCCCAGACATTAAACCGCACCAGAT
CCTGGTTAAAGTTGATACCTGCGGCGTTTGTCACACTGACCTGCATGCAGCTCGTGGTGACT
GGCCTTCCAAGCCGAACCCGCCGTTTATCCCGGGTCACGAAGGCGTTGGCCACATCGTAGCT
GTTGGCAGCCAGGTGGGCGACTTTGTTAAGACTGGTGATGTTGTAGGTGTGCCTTGGCTGTA
CTCTGCATGTGGTCATTGTGAGCACTGTCTGGGTGGCTGGGAAACTCTGTGCGAGAAGCAGG
ACGACACTGGTTATACCGTGAACGGTTGTTTTGCAGAATACGTAGTTGCCGATCCGAATTAC
GTTGCACACCTGCCGAGCACCATTGATCCACTGCAGGCATCCCCGGTGCTGTGTGCTGGCCT
GACTGTTTACAAAGGTCTGAAGATGACTGAAGCTCGTCCAGGTCAGTGGGTTGCAGTTAGCG
GTGTAGGCGGCCTGGGTCAAATGGCAGTGCAGTATGCGGTGGCGATGGGCATGAACGTTGTT
GCCGTAGATATCGATGACGAAAAACTGGCCACCGCTAAAAAGCTGGGCGCTTCCCTGACTGT
GAACGCAAAAGATACCGATCCGGCTCGTTTTATCCAGCAACAGATCGGTGGTGCACATGGTG
CTCTGGTTACCGCAGTGGGTCGTACTGCCTTCTCCCAAGCAATGGGTTATGCCCGTCGTGGT
GGTACCATCGTTCTGAACGGTCTGCCGCCGGGTGACTTTCCGGTGTCCATTTTTGATATGGT
GATGAACGGCACGACCATCCGTGGTAGCATCGTGGGCACCCGTCTGGATATGATCGAAGCTA
TGGATTTCTTCGCGCGCGGCAAAGTCAAATCCGTTGTTACCCCGGGCAAACTGGAAAACATC
AACACGATCTTCGACGACCTGCAGAACGGTCGTCTGGAAGGTCGTACCGTTCTGGATTTTCG
TAGCTGAGAGACTCCCGGGACCTCAGAGTCCGCCACACCCGAACATAAGCTTGGCTGCGCAA
AAAACCCCGCTTCGGCGGGGTTTTTTCGCGATGCATACTTGCGATCGTTGACAGTAAAAATT
GCCCGTTTGTGAACCACTTGTTTGCAAACGGGCATGACTCCTGACTTTTATTTCTGCCTTTT
ATTCCTTTTACACTTGTTTTTATGAAGCCCTTCACAGAATTGTCCTTTCACGATTCCGTCTC
TCTGATGATTGATGTTAATTAACAATGTATTCACCGAAAACAAACATATAAATCACAGGAGT
CGCCCATGTCAGTACCCGTTCAACATCCTATGTATATCGATGGACAGTTTGTTACCTGGCGT
GGAGACGCATGGATTGATGTGGTAAACCCTGCTACAGAGGCTGTCATTTCCCGCATACCCGA
TGGTCAGGCCGAGGATGCCCGTAAGGCAATCGATGCAGCAGAACGTGCACAACCAGAATGGG
AAGCGTTGCCTGCTATTGAACGCGCCAGTTGGTTGCGCAAAATCTCCGCCGGGATCCGCGAA
CGCGCCAGTGAAATCAGTGCGCTGATTGTTGAAGAAGGGGGCAAGATCCAGCAGCTGGCTGA
AGTCGAAGTGGCTTTTACTGCCGACTATATCGATTACATGGCGGAGTGGGCACGGCGTTACG
AGGGCGAGATTATTCAAAGCGATCGTCCAGGAGAAAATATTCTTTTGTTTAAACGTGCGCTT
GGTGTGACTACCGGCATTCTGCCGTGGAACTTCCCGTTCTTCCTCATTGCCCGCAAAATGGC
TCCCGCTCTTTTGACCGGTAATACCATCGTCATTAAACCTAGTGAATTTACGCCAAACAATG
CGATTGCATTCGCCAAAATCGTCGATGAAATAGGCCTTCCGCGCGGCGTGTTTAACCTTGTA
CTGGGGCGTGGTGAAACCGTTGGGCAAGAACTGGCGGGTAACCCAAAGGTCGCAATGGTCAG
TATGACAGGCAGCGTCTCTGCAGGTGAGAAGATCATGGCGACTGCGGCGAAAAACATCACCA
AAGTGTGTCTGGAATTGGGGGGTAAAGCACCAGCTATCGTAATGGACGATGCCGATCTTGAA
CTGGCAGTCAAAGCCATCGTTGATTCACGCGTCATTAATAGTGGGCAAGTGTGTAACTGTGC
AGAACGTGTTTATGTACAGAAAGGCATTTATGATCAGTTCGTCAATCGGCTGGGTGAAGCGA
TGCAGGCGGTTCAATTTGGTAACCCCGCTGAACGCAACGACATTGCGATGGGGCCGTTGATT
AACGCCGCGGCGCTGGAAAGGGTCGAGCAAAAAGTGGCGCGCGCAGTAGAAGAAGGGGCGAG
AGTGGCGTTCGGTGGCAAAGCGGTAGAGGGGAAAGGATATTATTATCCGCCGACATTGCTGC
TGGATGTTCGCCAGGAAATGTCGATTATGCATGAGGAAACCTTTGGCCCGGTGCTGCCAGTT
GTCGCATTTGACACGCTGGAAGATGCTATCTCAATGGCTAATGACAGTGATTACGGCCTGAC
CTCATCAATCTATACCCAAAATCTGAACGTCGCGATGAAAGCCATTAAAGGGCTGAAGTTTG
GTGAAACTTACATCAACCGTGAAAACTTCGAAGCTATGCAAGGCTTCCACGCCGGATGGCGT
AAATCCGGTATTGGCGGCGCAGATGGTAAACATGGCTTGCATGAATATCTGCAGACCCAGGT
GGTTTATTTACAGTCTTAATGAGTGAAAGAGGCGGAGGTTTTTCCTCCGCCTGTGCGCAAT
GGAAACAGACCAGTTATTTTTCTGCGCCTCTTCCTGACCTGCGGCAATAATTGCACTCGCCA
TTTGCTGGCTAAGAATGACTTTAGTTTTCATTTTGTTATTCCTTTTCAAGGCTTGTTCTAC
AATTTCAATCCAGTGACGCACAGAGGTACGACCGGCGCTCGCCAGATGCGTCTGGCAACCAA
TGTTGGCGGTGACGATCATTTCCGGTTTGCCGCTTTCCAGCGCATTCATTTTGTTATCCCGC

68 AGCTGGCGTGCCAGATCGGGATGCGTTAACGCATATGTTCCCGCTGAACCGCAGCACAGATG
69 GCTGTCGGGAACGTCCGTTAAGGTAAATCCAAGACGAAGCAACACTTTTTCCACTTCGCCGT
70 TCAGCTTTTGCGCATGTTGTAGGGTACACGGACAGTGGAAGGCCAGCTTTTTATCGCCGCGA
71 ATTGCCAGTTTTTCCAGCGGTTCCTCGCGCAGAAGTTCGACTAAATCGACCGCCAGTTCACT
72 GACCTGACGTGCTTTATCGGCATATAACGCATCGTTTTTCAGCATCTGCCCATACTCTTTGA
73 CAAACGCGCCGCAGCCGCTGGCGGTTTGCAAAATTGCCTCGGCACCTGCTTCAATCGCGGGC
74 CACCAGGCATCAATATTATTGCGCGCCCGTGCCAGCCCTTTCTCCTGCGCATTAAGATGATA
75 GTCCACCGCGCCACAACAGCCTGCTTCGTTAGCTGGCATGACGCTGATCCCCAGACGATCCA
76 GCACTCGCGCAGTTGCCGCGTTGGTGTTGGGCGAAAGCGTAGGCTGGGCGCAGCCTTCCAAC
77 ATTAAAACCCGACGCTTATGGCGCAGCGGCGGACGCGGTTTAGCTTTCACCGTTTCAGCAGG
78 CAGTTTTGCTCTGACCTGTTCCGGTAAAAACGGTCGCAGCACCAGCCCTACCTGCGTCAGCG
79 CACGGAAGACCGCCGGACGCGGCACTACCTGGCGCAATCCTTCGCGCAGTATTCGCTCCGGC
80 AGTGGGCGTTTCACTTTCTGCTCGACAATATCACGCCCGATATCCAGCAAATTGTGATAGCG
81 CACACCAGAAGGACAGGTGGTTTCACAATTACGGCAAGTGAGGCAGCGATCGAGATGCTCCT
82 GTGTTTTAAGCGTGACTTCGTTGCCTTCCAGCACCTGTTTAATCAGATAGATGCGCCCGCGC
83 GGCCCGTCCAGTTCATCGCCCAGAAGCTGATAGGTTGGGCAGGTTGCGGTACAAAATCCGCA
84 GTGAACACAGGCGCGCAGGATGCTGTCGGCTTCCAGCGCGCGCGTTCTGCCGCATCTCTT
85 CAGTTAATTGGGTTTGCATAGCCTGCTCCTCAAAGTTCCGCGTACATGCGACCGGGGTTAAA
86 CACGCCGCAAGGGTCGAGCTGCTGTTTAAGCTGCTGGTGATAGCGGAATAAAGGAGCCGATA
87 GCGGGGCAAAGCCACCATCTCCGGCACTAAAGCGGGTCGCATGACCGCCAGCGTTGCGGGCG
88 ATGCGATGGATTTGATTGTCCTCGGCTGTCGATTTCAGCCAGCGTAACGCCCCGCCCCAGTC
89 GATCAGTTGCTCGCCGGGTAAATCCATCATCGGCGCATCACTGGGTAATGAAATGCGCCATA
90 AGGTACCTGGTAACGAGAAGAACGGCAGTTGTTGTTCACGCAATTGCTGCCAGAACTGACCG
91 GCAACCTCTTCGCCACCCAGCAGTTCACGCGCTGCTTTTACCGATCCTTCGCCGCCCTCAAG
92 GCGGATCCACAACGCATTGTCGAAGTAACATAAGCCACTAATGGGTAATGGCTGGAGTTGCC
93 ACTCGGCGATTTCACTCATGGCTTCTTGCAGGCTGATTTCCCGACGCAGGCTCAGGGAGGCG
94 CGCGGTCGCGGTAACACTTTCATTGAGATTTCAGTGAGCACGCCAAGACAACCGTAGCTTCC
95 GACCATTAACCGTGAGAGATCGTATCCGGCAACGTTTTTCATCACTTCGCCACCAAAACGCA
96 GATGTTTTCCAGCGCCGGTAATGATGCGCGTGCCGAGGACAAAATCGCGGACCGAACCGCTC
97 CACGGGCGACGCGGCCCCGCCAGCCCGCAGGCGACCATCCCGCCCCAGGTGGCTTCTTCACC
98 ATAATGCGGCGGCTCACAGGGGAGCATTTGCCCCGCGCTTTCCAGCGCCGCTTCAATTGTCA
99 CCAGCGGCGTTCCGACACGCGCGGTTATCACCAGCTCGGTCGGGTCGTAATTAACAATGCCG
100 CGATGACAACGAACATCCAGCGTTTGCCCGGTGACAGGGCGACCTAAAAAGGCTTTGCTATT
101 GCTGCCCTGAATCACCAGCGGCGTTTTATCGCTAATCGCCTGATTCACCTGCTCCAGCAGCG
102 CCTGGCTGTAATCACACTCGCGTAGCATCAGAAACGCTCCAGTTCAGGGAAAGGTAAATGAC
103 CGTGATGCACATGCATGGCACCAAATTCAGCACAGCGGTGTAGCGTGGGAATGTTTTTCCCA
104 GGGTTCAGCAAACCATCGGGGTCAAACGCCGCCTTGACCGCATGGAAGGTCGTGATTTCATC
105 GCTGTTGAACTGGGCGCACATTTGATTGATTTTTTCTCGCCCGATGCCATGTTCGCCACTGA
106 TGCTGCCGCCAACTTCAACGCAGAGTTCGAGGATCTTCCCGCCCAGCTCTTCCGCGCGGGCA
107 AATTCACCGGGTTCGTTGGCATCGAAAAGGATTAACGGGTGCATGTTGCCATCTCCGGCATG
108 AAAGACGTTGGCAACACGTAAATCATATTGCTGCGATAAACGGGCAATGCCTTCCAGTACGC
109 CAGGCAGGGCGCGACGCGGGATGGTGCCATCCATGCAGTAGTAATCCGGGGAGATACGTCCT
110 ACCGCCGGGAACGCATTTTTGCGACCGGCCCAGAAACGTACGCGCTCTGCTTCGTCCTGTGC
111 CAGACGGACGTCAGTCGCGCCCGCTTTCAACAAGATGTCGTTAACCCGCTCGCAGTCTTCCT
112 GTACGTCAGACTCCACGCCGTCCAGCTCGCATAACAAAATCGCTTCGGCGTCGACGGGATAA
113 CCGGCATGAATAAAATCTTCCGCCGCGCGGATCGACAGGTTATCCATCATCTCCAGCCCGCC
114 GGGGATAATGCCATTGGCGATGATGTCACCAACCGCAAGTCCGGCTTTTTCTACCGAGTCAA
115 AGCTGGCTAACAGAACCCGCGCCACGGGCGGCTTCGGCAGCAGTTTTACCGTCACTTCGGTG
116 GTCACGCCGAGCATACCTTCCGATCCGGTGAACAGCGCCAGCAGGTCAAAACCAGGTGAATC
117 CAGCGCGTCCGATCCAAGCGTCAGTGCCTCGCCGTCCAGCGTTTGCACTTCAATTTTCAGCA
118 GGTTATGTACGGTCAGACCATATTTCAGGCAGTGGACGCCGCCGGCATTTTCAGCCACATTG

CCGCCAATGGAACAGGCGATTTGTGAGGAAGGGTCCGGTGCGTAGTAGAGATTATGCGGTGC
AACGGCCTGGGAGATCGCCAGGTTACGCACGCCTGGCTGCACGCGCGCGCGGCGACCAACGG
GGTTAATGTCGAGGATCTCTTTAAAGCGCGCCATCACCAACAACACACCTTTTTCCAGCGGC
AGCGCGCCACCAGAAAGCCCGGTGCCTGCACCACGGGTCACCACCGGTACACGCAGGCGATG
GCAGACAGCCAGAATCGCTGTCACCTGTTCCATTTGCTTAGGCAGAACAACCAGTAATGGAC
GCGTGCGATACGCGCTCAACCCGTCACACTCGTAAGGAATGATCTCCTCATCGGTATGCAGG
ATCTCAAGTCCAGGGACATGCTCACGCAGTGCCATCAGTACCGATGTGCGGTCGACATCGGG
TAAAGCGCCATCAAGACGCTCTTCGTACAAGATGCTCATGAGTAGGCTTCGCTTTGTTGTGT
TGTGTGGCAGCTGATTTTTGCGCGCTGCTTCTGTGAACAGTTATTAAGCGGGCTTTTCGTTT
TCGTCTATCTCTTTAGCTACCGGTCAGACCATTTTTTTTCCAGCTCTGTGACCTTGTCTTGG
TTAACTCAATGTTAAATTGATGTAACATAATCACTTACGTGATGTGCGTGTTTTGCGAGTTA
AGAACAGAAAAAATTGGTCCTACCTGTGCACGAGGTCCGGGATCTGACGCTCAGTGGAACGAA
AACTCACGTTAAGGGATTTTGGTCATGAACAATAAAACTGTCTGCTTACATAAACAGTAATA
CAAGGGGTGTTATGAGCCATATTCAACGGGAAACGTCTTGCTCTAGGCCGCGATTAAATTCC
AACATGGATGCTGATTTATATGGGTATAAATGGGCTCGCGATAATGTCGGGCAATCAGGTGC
GACAATCTATCGATTGTATGGGAAGCCCGATGCGCCAGAGTTGTTTCTGAAACATGGCAAAG
GTAGCGTTGCCAATGATGTTACAGATGAGATGGTCAGACTAAACTGGCTGACGGAATTTATG
CCTCTTCCGACCATCAAGCATTTTATCCGTACTCCTGATGATGCATGGTTACTCACCACTGC
GATCCCCGGGAAAACAGCATTCCAGGTATTAGAAGAATATCCTGATTCAGGTGAAAATATTG
TTGATGCGCTGGCAGTGTTCCTGCGCCGGTTGCATTCGATTCCTGTTTGTAATTGTCCTTTT
AACAGCGATCGCGTATTTCGTCTCGCTCAGGCGCAATCACGAATGAATAACGGTTTGGTTGA
TGCGAGTGATTTTGATGACGAGCGTAATGGCTGGCCTGTTGAACAAGTCTGGAAAGAAATGC
ATAAACTTTTGCCATTCTCACCGGATTCAGTCGTCACTCATGGTGATTTCTCACTTGATAAC
CTTATTTTTGACGAGGGGAAATTAATAGGTTGTATTGATGTTGGACGAGTCGGAATCGCAGA
CCGATACCAGGATCTTGCCATCCTATGGAACTGCCTCGGTGAGTTTTCTCCTTCATTACAGA
AACGGCTTTTTCAAAAATATGGTATTGATAATCCTGATATGAATAAATTGCAGTTTCATTTG
ATGCTCGATGAGTTTTTCTAAAAGCTTAATTAGCTGATCTAGACGCGTGCTAGAGGCATCAA
ATAAAACGAAAGGCTCAGTCGAAAGACTGGGCCTTTCGTTTTATCTGTTGTTTGTCGGTGAA
CGCTCTCCTGAGTAGGACAAATAGGTCGAGGGTGAAGTACTTGCTGACTTCCTTGAGGAACA
CATGATGCGTCCTACGGTTGCTGCTACGCATATCATTGAGATGTCTGTGGGAGGAGTTGATG
TGTACTCTGAGGACGATGAGGGTTACGGTACGTCTTTCATTGAGTGGTGATTTATGCATTAG
GACTGCATAGGGATGCACTATAGACCACGGATGGTCAGTTCTTTAAGTTACTGAAAAGACAC
GATAAATTAATACGACTCACTATAGGGAGAGGAGGGACGAAAGGTTACTATATAGATACTGA
ATGAATACTTATAGAGTGCATAAAGTATGCATAATGGTGTACCTAGAGTGACCTCTAAGAAT
GGTGATTATATTGTATTAGTATCACCTTAACTTAAGGCGGGATCGTCACCCTCAGCAGCGAA
AGACAGCTGTCGGTCAGAGCGTCATTGCGAAGCTGAGTGTGATCGATGCCATCAGCGAAGGG
CCCAAACTCCGAGCGATTAAGCGTTTGCTGGCTGTCACGCCTGCCTGTTGCTTGCTTGGACT
TGCGATGTACGTGCTCAGCTGTCTTTCGCTGCTGAGGGTGACGATCCCGCGAGGGCCTATGG
AGTTCCTATAGGGTCCTTTAAAATATACCATAAAAATCTGAGTGACTATCTCACAGTGTACG
GACCTAAAGTTCCCCCATAGGGGGTACCTAAAGCCCAGCCAATCACCTAAAGTCAACCTTCG
GTTGACCTTGAGGGTTCCCTAAGGGTTGGGGATGACCCTTGGGTTTGTCTTTGGGTGTTACC
TTGAGTGTCTCTCTGTGTCCCTATCTGTTACAGTCTCCTAAAGTATCCTCCTAAAGTCACCT
CCTAACGCACATTTCCCCGAAAAGTGCCACCTGGGTCCTTTTC

**Supplementary Data 7: EGA1-8, nanopore sequencing**

pAN29 Wildtype

CCTATGGAACTGCCTCGGTGAGTTTTCTCCTTCATTACAGAAACGGCTTTTTCAAAAATATG
GTATTGATAATCCTGATATGAATAAATTGCAGTTTCATTTGATGCTCGATGAGTTTTTCTAA
AAGCTTAATTAGCTGATCTAGACGCGTGCTAGAGGCATCAAATAAAACGAAAGGCTCAGTCG
AAAGACTGGGCCTTTCGTTTTATCTGTTGTTTGTCGGTGAACGCTCTCCTGAGTAGGACAAA
TAGGTCGAGGGTGAAGTACTTGCTGACTTCCTTGAGGAACACATGATGCGTCCTACGGTTGC
TGCTACGCATATCATTGAGATGTCTGTGGGAGGAGTTGATGTGTACTCTGAGGACGATGAGG
GTTACGGTACGTCTTTCATTGAGTGGTGATTTATGCATTAGGACTGCATAGGGATGCACTAT
AGACCACGGATGGTCAGTTCTTTAAGTTACTGAAAAGACACGATAAATTAATACGACTCACT
ATAGGGAGAGGAGGGACGAAAGGTTACTATATAGATACTGAATGAATACTTATAGAGTGCAT
AAAGTATGCATAATGGTGTACCTAGAGTGACCTCTAAGAATGGTGATTATATTGTATTAGTA
TCACCTTAACTTAAGGCGGGATCGTCACCCTCAGCAGCGAAAGACAGCTGTCGGTCAGAGCG
TCATTGCGAAGCTGAGTGTGATCGATGCCATCAGCGAAGGGCCCAAACTCCGAGCGATTAAG
CGTTTGCTGGCTGTCACGCCTGCCTGTTGCTTGCTTGGACTTGCGATGTACGTGCTCAGCTG
TCTTTCGCTGCTGAGGGTGACGATCCCGCGAGGGCCTATGGAGTTCCTATAGGGTCCTTTAA
AATATACCATAAAAATCTGAGTGACTATCTCACAGTGTACGGACCTAAAGTTCCCCCATAGG
GGGTACCTAAAGCCCAGCCAATCACCTAAAGTCAACCTTCGGTTGACCTTGAGGGTTCCCTA
AGGGTTGGGGATGACCCTTGGGTTTGTCTTTGGGTGTTACCTTGAGTGTCTCTCTGTGTCCC
TATCTGTTACAGTCTCCTAAAGTATCCTCCTAAAGTCACCTCCTAACGCACATTTCCCCGAA
AAGTGCCACCTGGGTCCTTTTCACTTCGGGCTCATGAGCAAATATTTTATCTGATTAATAAG
ATGATCTTCTTGAGATCGTTTTGGTCTGCGCGTAATCTCTTGCTCTGAAAACGAAAAAACCG
CCTTGCAGGGCGGTTTTTCGAAGGTTCTCTGAGCTACCAACTCTTTGAACCGAGGTAACTGG
CTTGGAGGAGCGCAGTCACCAAAACTTGTCCTTTCAGTTTAGCCTTAACCGGCGCATGACTT
CAAGACTAACTCCTCTAAATCAATTACCAGTGGCTGCTGCCAGTGGTGCTTTTGCATGTCTT
TCCGGGTTGGACTCAAGACGATAGTTACCGGATAAGGCGCAGCGGTCGGACTGAACGGGGGG
TTCGTGCATACAGTCCAGCTTGGAGCGAACTGCCTACCCGGAACTGAGTGTCAGGCGTGGAA
TGAGACAAACGCGGCCATAACAGCGGAATGACACCGGTAAACCGAAAGGCAGGAACAGGAGA
GCGCACGAGGGAGCCGCCAGGGGGAAACGCCTGGTATCTTTATAGTCCTGTCGGGTTTCGCC
ACCACTGATTTGAGCGTCAGATTTCGTGATGCTTGTCAGGGGGGCGGAGCCTATGGAAAAAC
GGCTTTGCCGCGGCCCTCTCACTTCCCTGTTAAGTATCTTCCTGGCATCTTCCAGGAAATCT
CCGCCCCGTTCGTAAGCCATTTCCGCTCGCCGCAGTCGAACGACCGAGCGTAGCGAGTCAGT
GAGCGAGGAAGCGGAATATATCCTTGACAGCTAGCTCAGTCCTAGGTACTGTGCTAGCACCC
GTTTTTACTAGTCTAGAAATAATAACAAAATGAGGAGGTACTGATATACATATGGCGGACAC
GATGCTGGCGGCGGTTGTACGTGAATTTGGTAAACCTCTGTCTATCGAACGCCTGCCGATCC
CAGACATTAAACCGCACCAGATCCTGGTTAAAGTTGATACCTGCGGCGTTTGTCACACTGAC
CTGCATGCAGCTCGTGGTGACTGGCCTTCCAAGCCGAACCCGCCGTTTATCCCGGGTCACGA
AGGCGTTGGCCACATCGTAGCTGTTGGCAGCCAGGTGGGCGACTTTGTTAAGACTGGTGATG
TTGTAGGTGTGCCTTGGCTGTACTCTGCATGTGGTCATTGTGAGCACTGTCTGGGTGGCTGG
GAAACTCTGTGCGAGAAGCAGGACGACACTGGTTATACCGTGAACGGTTGTTTTGCAGAATA
CGTAGTTGCCGATCCGAATTACGTTGCACACCTGCCGAGCACCATTGATCCACTGCAGGCAT
CCCCGGTGCTGTGTGCTGGCCTGACTGTTTACAAAGGTCTGAAGATGACTGAAGCTCGTCCA
GGTCAGTGGGTTGCAGTTAGCGGTGTAGGCGGCCTGGGTCAAATGGCAGTGCAGTATGCGGT
GGCGATGGGCATGAACGTTGTTGCCGTAGATATCGATGACGAAAAACTGGCCACCGCTAAAA
AGCTGGGCGCTTCCCTGACTGTGAACGCAAAAGATACCGATCCGGCTCGTTTTATCCAGCAA
CAGATCGGTGGTGCACATGGTGCTCTGGTTACCGCAGTGGGTCGTACTGCCTTCTCCCAAGC
AATGGGTTATGCCCGTCGTGGTGGTACCATCGTTCTGAACGGTCTGCCGCCGGGTGACTTTC
CGGTGTCCATTTTTGATATGGTGATGAACGGCACGACCATCCGTGGTAGCATCGTGGGCACC
CGTCTGGATATGATCGAAGCTATGGATTTCTTCGCGCGCGGCAAAGTCAAATCCGTTGTTAC
CCCGGGCAAACTGGAAAACATCAACACGATCTTCGACGACCTGCAGAACGGTCGTCTGGAAG

| | |
|---|---|
| 52 | GTCGTACCGTTCTGGATTTTCGTAGCTGAGAGACTCCCGGGACCTCAGAGTCCGCCACACCC |
| 53 | GAACATAAGCTTGGCTGCGCAAAAAACCCCGCTTCGGCGGGGTTTTTTCGCGATGCATACTT |
| 54 | GCGATCGTTGACAGTAAAAATTGCCCGTTTGTGAACCACTTGTTTGCAAACGGGCATGACTC |
| 55 | CTGACTTTTATTTCTGCCTTTTATTCCTTTTACACTTGTTTTTATGAAGCCCTTCACAGAAT |
| 56 | TGTCCTTTCACGATTCCGTCTCTCTGATGATTGATGTTAATTAACAATGTATTCACCGAAAA |
| 57 | CAAACATATAAATCACAGGAGTCGCCCATGTCAGTACCCGTTCAACATCCTATGTATATCGA |
| 58 | TGGACAGTTTGTTACCTGGCGTGGAGACGCATGGATTGATGTGGTAAACCCTGCTACAGAGG |
| 59 | CTGTCATTTCCCGCATACCCGATGGTCAGGCCGAGGATGCCCGTAAGGCAATCGATGCAGCA |
| 60 | GAACGTGCACAACCAGAATGGGAAGCGTTGCCTGCTATTGAACGCGCCAGTTGGTTGCGCAA |
| 61 | AATCTCCGCCGGGATCCGCGAACGCGCCAGTGAAATCAGTGCGCTGATTGTTGAAGAAGGGG |
| 62 | GCAAGATCCAGCAGCTGGCTGAAGTCGAAGTGGCTTTTACTGCCGACTATATCGATTACATG |
| 63 | GCGGAGTGGGCACGGCGTTACGAGGGCGAGATTATTCAAAGCGATCGTCCAGGAGAAAATAT |
| 64 | TCTTTTGTTTAAACGTGCGCTTGGTGTGACTACCGGCATTCTGCCGTGGAACTTCCCGTTCT |
| 65 | TCCTCATTGCCCGCAAAATGGCTCCCGCTCTTTTGACCGGTAATACCATCGTCATTAAACCT |
| 66 | AGTGAATTTACGCCAAACAATGCGATTGCATTCGCCAAAATCGTCGATGAAATAGGCCTTCC |
| 67 | GCGCGGCGTGTTTAACCTTGTACTGGGGCGTGGTGAAACCGTTGGGCAAGAACTGGCGGGTA |
| 68 | ACCCAAAGGTCGCAATGGTCAGTATGACAGGCAGCGTCTCTGCAGGTGAGAAGATCATGGCG |
| 69 | ACTGCGGCGAAAAACATCACCAAAGTGTGTCTGGAATTGGGGGGTAAAGCACCAGCTATCGT |
| 70 | AATGGACGATGCCGATCTTGAACTGGCAGTCAAAGCCATCGTTGATTCACGCGTCATTAATA |
| 71 | GTGGGCAAGTGTGTAACTGTGCAGAACGTGTTTATGTACAGAAAGGCATTTATGATCAGTTC |
| 72 | GTCAATCGGCTGGGTGAAGCGATGCAGGCGGTTCAATTTGGTAACCCCGCTGAACGCAACGA |
| 73 | CATTGCGATGGGGCCGTTGATTAACGCCGCGGCGCTGGAAAGGGTCGAGCAAAAAGTGGCGC |
| 74 | GCGCAGTAGAAGAAGGGGCGAGAGTGGCGTTCGGTGGCAAAGCGGTAGAGGGGAAAGGATAT |
| 75 | TATTATCCGCCGACATTGCTGCTGGATGTTCGCCAGGAAATGTCGATTATGCATGAGGAAAC |
| 76 | CTTTGGCCCGGTGCTGCCAGTTGTCGCATTTGACACGCTGGAAGATGCTATCTCAATGGCTA |
| 77 | ATGACAGTGATTACGGCCTGACCTCATCAATCTATACCCAAAATCTGAACGTCGCGATGAAA |
| 78 | GCCATTAAAGGGCTGAAGTTTGGTGAAACTTACATCAACCGTGAAAACTTCGAAGCTATGCA |
| 79 | AGGCTTCCACGCCGGATGGCGTAAATCCGGTATTGGCGGCGCAGATGGTAAACATGGCTTGC |
| 80 | ATGAATATCTGCAGACCCAGGTGGTTTATTTACAGTCTTAATGAGTGAAAGAGGCGGAGGTT |
| 81 | TTTTCCTCCGCCTGTGCGCAATGGAAACAGACCAGTTATTTTTCTGCGCCTCTTCCTGACCT |
| 82 | GCGGCAATAATTGCACTCGCCATTTGCTGGCTAAGAATGACTTTAGTTTTCATTTTGTTATT |
| 83 | CCTTTTCAAGGGCTTGTTCTACAATTTCAATCCAGTGACGCACAGAGGTACGACCGGCGCTC |
| 84 | GCCAGATGCGTCTGGCAACCAATGTTGGCGGTGACGATCATTTCCGGTTTGCCGCTTTCCAG |
| 85 | CGCATTCATTTTGTTATCCCGCAGCTGGCGTGCCAGATCGGGATGCGTTAACGCATATGTTC |
| 86 | CCGCTGAACCGCAGCACAGATGGCTGTCGGGAACGTCCGTTAAGGTAAATCCAAGACGAAGC |
| 87 | AACACTTTTTCCACTTCGCCGTTCAGCTTTTGCGCATGTTGTAGGGTACACGGACAGTGGAA |
| 88 | GGCCAGCTTTTTATCGCCGCGAATTGCCAGTTTTTCCAGCGGTTCCTCGCGCAGAAGTTCGA |
| 89 | CTAAATCGACCGCCAGTTCACTGACCTGACGTGCTTTATCGGCATATAACGCATCGTTTTTC |
| 90 | AGCATCTGCCCATACTCTTTGACAAACGCGCCGCAGCCGCTGGCGGTTTGCAAAATTGCCTC |
| 91 | GGCACCTGCTTCAATCGCGGGCCACCAGGCATCAATATTATTGCGCGCCCGTGCCAGCCCTT |
| 92 | TCTCCTGCGCATTAAGATGATAGTCCACCGCGCCACAACAGCCTGCTTCGTTAGCTGGCATG |
| 93 | ACGCTGATCCCCAGACGATCCAGCACTCGCGCAGTTGCCGCGTTGGTGTTGGGCGAAAGCGT |
| 94 | AGGCTGGGCGCAGCCTTCCAACATTAAAACCCGACGCTTATGGCGCAGCGGCGGACGCGGTT |
| 95 | TAGCTTTCACCGTTTCAGCAGGCAGTTTTGCTCTGACCTGTTCCGGTAAAAACGGTCGCAGC |
| 96 | ACCAGCCCTACCTGCGTCAGCGCACGGAAGACCGCCGGACGCGGCACTACCTGGCGCAATCC |
| 97 | TTCGCGCAGTATTCGCTCCGGCAGTGGGCGTTTCACTTTCTGCTCGACAATATCACGCCCGA |
| 98 | TATCCAGCAAATTGTGATAGCGCACACCAGAAGGACAGGTGGTTTCACAATTACGGCAAGTG |
| 99 | AGGCAGCGATCGAGATGCTCCTGTGTTTTAAGCGTGACTTCGTTGCCTTCCAGCACCTGTTT |
| 100 | AATCAGATAGATGCGCCCGCGCGGCCCGTCCAGTTCATCGCCCAGAAGCTGATAGGTTGGGC |
| 101 | AGGTTGCGGTACAAAATCCGCAGTGAACACAGGCGCGCAGGATGCTGTCGGCTTCCAGCGCG |
| 102 | CGCGCGTTCTGCCGCATCTCTTCAGTTAATTGGGTTTGCATAGCCTGCTCCTCAAAGTTCCG |

CGTACATGCGACCGGGGTTAAACACGCCGCAAGGGTCGAGCTGCTGTTTAAGCTGCTGGTGA
TAGCGGAATAAAGGAGCCGATAGCGGGGCAAAGCCACCATCTCCGGCACTAAAGCGGGTCGC
ATGACCGCCAGCGTTGCGGGCGATGCGATGGATTTGATTGTCCTCGGCTGTCGATTTCAGCC
AGCGTAACGCCCCGCCCCAGTCGATCAGTTGCTCGCCGGGTAAATCCATCATCGGCGCATCA
CTGGGTAATGAAATGCGCCATAAGGTACCTGGTAACGAGAAGAACGGCAGTTGTTGTTCACG
CAATTGCTGCCAGAACTGACCGGCAACCTCTTCGCCACCCAGCAGTTCACGCGCTGCTTTTA
CCGATCCTTCGCCGCCCTCAAGGCGGATCCACAACGCATTGTCGAAGTAACATAAGCCACTA
ATGGGTAATGGCTGGAGTTGCCACTCGGCGATTTCACTCATGGCTTCTTGCAGGCTGATTTC
CCGACGCAGGCTCAGGGAGGCGCGCGGTCGCGGTAACACTTTCATTGAGATTTCAGTGAGCA
CGCCAAGACAACCGTAGCTTCCGACCATTAACCGTGAGAGATCGTATCCGGCAACGTTTTTC
ATCACTTCGCCACCAAAACGCAGATGTTTTCCAGCGCCGGTAATGATGCGCGTGCCGAGGAC
AAAATCGCGGACCGAACCGCTCCACGGGCGACGCGGCCCCGCCAGCCCGCAGGCGACCATCC
CGCCCCAGGTGGCTTCTTCACCATAATGCGGCGGCTCACAGGGGAGCATTTGCCCCGCGCTT
TCCAGCGCCGCTTCAATTGTCACCAGCGGCGTTCCGACACGCGCGGTTATCACCAGCTCGGT
CGGGTCGTAATTAACAATGCCGCGATGACAACGAACATCCAGCGTTTGCCCGGTGACAGGGC
GACCTAAAAAGGCTTTGCTATTGCTGCCCTGAATCACCAGCGGCGTTTTATCGCTAATCGCC
TGATTCACCTGCTCCAGCAGCGCCTGGCTGTAATCACACTCGCGTAGCATCAGAAACGCTCC
AGTTCAGGGAAAGGTAAATGACCGTGATGCACATGCATGGCACCAAATTCAGCACAGCGGTG
TAGCGTGGGAATGTTTTTCCCAGGGTTCAGCAAACCATCGGGGTCAAACGCCGCCTTGACCG
CATGGAAGGTCGTGATTTCATCGCTGTTGAACTGGGCGCACATTTGATTGATTTTTTCTCGC
CCGATGCCATGTTCGCCACTGATGCTGCCGCCAACTTCAACGCAGAGTTCGAGGATCTTCCC
GCCCAGCTCTTCCGCGCGGGCAAATTCACCGGGTTCGTTGGCATCGAAAAGGATTAACGGGT
GCATGTTGCCATCTCCGGCATGAAAGACGTTGGCAACACGTAAATCATATTGCTGCGATAAA
CGGGCAATGCCTTCCAGTACGCCAGGCAGGGCGCGACGCGGGATGGTGCCATCCATGCAGTA
GTAATCCGGGGAGATACGTCCTACCGCCGGGAACGCATTTTTGCGACCGGCCCAGAAACGTA
CGCGCTCTGCTTCGTCCTGTGCCAGACGGACGTCAGTCGCGCCCGCTTTCAACAAGATGTCG
TTAACCCGCTCGCAGTCTTCCTGTACGTCAGACTCCACGCCGTCCAGCTCGCATAACAAAAT
CGCTTCGGCGTCGACGGGATAACCGGCATGAATAAAATCTTCCGCCGCGCGGATCGACAGGT
TATCCATCATCTCCAGCCCGCCGGGGATAATGCCATTGGCGATGATGTCACCAACCGCAAGT
CCGGCTTTTTCTACCGAGTCAAAGCTGGCTAACAGAACCCGCGCCACGGGCGGCTTCGGCAG
CAGTTTTACCGTCACTTCGGTGGTCACGCCGAGCATACCTTCCGATCCGGTGAACAGCGCCA
GCAGGTCAAAACCAGGTGAATCCAGCGCGTCCGATCCAAGCGTCAGTGCCTCGCCGTCCAGC
GTTTGCACTTCAATTTTCAGCAGGTTATGTACGGTCAGACCATATTTCAGGCAGTGGACGCC
GCCGGCATTTTCAGCCACATTGCCGCCAATGGAACAGGCGATTTGTGAGGAAGGGTCCGGTG
CGTAGTAGAGATTATGCGGTGCAACGGCCTGGGAGATCGCCAGGTTACGCACGCCTGGCTGC
ACGCGCGCGCGGCGACCAACGGGGTTAATGTCGAGGATCTCTTTAAAGCGCGCCATCACCAA
CAACACACCTTTTTCCAGCGGCAGCGCGCCACCAGAAAGCCCGGTGCCTGCACCACGGGTCA
CCACCGGTACACGCAGGCGATGGCAGACAGCCAGAATCGCTGTCACCTGTTCCATTTGCTTA
GGCAGAACAACCAGTAATGGACGCGTGCGATACGCGCTCAACCCGTCACACTCGTAAGGAAT
GATCTCCTCATCGGTATGCAGGATCTCAAGTCCAGGGACATGCTCACGCAGTGCCATCAGTA
CCGATGTGCGGTCGACATCGGGTAAAGCGCCATCAAGACGCTCTTCGTACAAGATGCTCATG
AGTAGGCTTCGCTTTGTTGTGTTGTGTGGCAGCTGATTTTTGCGCGCTGCTTCTGTGAACAG
TTATTAAGCGGGCTTTTCGTTTTCGTCTATCTCTTTAGCTACCGGTCAGACCATTTTTTTTC
CAGCTCTGTGACCTTGTCTTGGTTAACTCAATGTTAAATTGATGTAACATAATCACTTACGT
GATGTGCGTGTTTTGCGAGTTAAGAACAGAAAAATTGGTCCTACCTGTGCACGAGGTCCGGG
ATCTGACGCTCAGTGGAACGAAAACTCACGTTAAGGGATTTTGGTCATGAACAATAAAACTG
TCTGCTTACATAAACAGTAATACAAGGGGTGTTATGAGCCATATTCAACGGG

EGA1_pAN29_1

CCTATGGAACTGCCTCGGTGAGTTTTCTCCTTCATTACAGAAACGGCTTTTTCAAAAATATG
GTATTGATAATCCTGATATGAATAAATTGCAGTTTCATTTGATGCTCGATGAGTTTTTCTAA
AAGCTTAATTAGCTGATCTAGACGCGTGCTAGAGGCATCAAATAAAACGAAAGGCTCAGTCG
AAAGACTGGGCCTTTCGTTTTATCTGTTGTTTGTCGGTGAACGCTCTCCTGAGTAGGACAAA
TAGGTCGAGGGTGAAGTACTTGCTGACTTCCTTGAGGAACACATGATGCGTCCTACGGTTGC
TGCTACGCATATCATTGAGATGTCTGTGGGAGGAGTTGATGTGTACTCTGAGGACGATGAGG
GTTACGGTACGTCTTTCATTGAGTGGTGATTTATGCATTAGGACTGCATAGGGATGCACTAT
AGACCACGGATGGTCAGTTCTTTAAGTTACTGAAAAGACACGATAAATTAATACGACTCACT
ATAGGGAGAGGAGGGACGAAAGGTTACTATATAGATACTGAATGAATACTTATAGAGTGCAT
AAAGTATGCATAATGGTGTACCTAGAGTGACCTCTAAGAATGGTGATTATATTGTATTAGTA
TCACCTTAACTTAAGGCGGGATCGTCACCCTCAGCAGCGAAAGACAGCTGTCGGTCAGAGCG
TCATTGCGAAGCTGAGTGTGATCGATGCCATCAGCGAAGGGCCCAAACTCCGAGCGATTAAG
CGTTTGCTGGCTGTCACGCCTGCCTGTTGCTTGCTTGGACTTGCGATGTACGTGCTCAGCTG
TCTTTCGCTGCTGAGGGTGACGATCCCGCGAGGGCCTATGGAGTTCCTATAGGGTCCTTTAA
AATATACCATAAAAATCTGAGTGACTATCTCACAGTGTACGGACCTAAAGTTCCCCCATAGG
GGGTACCTAAAGCCCAGCCAATCACCTAAAGTCAACCTTCGGTTGACCTTGAGGGTTCCCTA
AGGGTTGGGGATGACCCTTGGGTTTGTCTTTGGGTGTTACCTTGAGTGTCTCTCTGTGTCCC
TATCTGTTACAGTCTCCTAAAGTATCCTCCTAAAGTCACCTCCTAACGCACATTTCCCCGAA
AAGTGCCACCTGGGTCCTTTTCACTTCGGGCTCATGAGCAAATATTTTATCTGATTAATAAG
ATGATCTTCTTGAGATCGTTTTGGTCTGCGCGTAATCTCTTGCTCTGAAAACGAAAAAACCG
CCTTGCAGGGCGGTTTTTCGAAGGTTCTCTGAGCTACCAACTCTTTGAACCGAGGTAACTGG
CTTGGAGGAGCGCAGTCACCAAAACTTGTCCTTTCAGTTTAGCCTTAACCGGCGCATGACTT
CAAGACTAACTCCTCTAAATCAATTACCAGTGGCTGCTGCCAGTGGTGCTTTTGCATGTCTT
TCCGGGTTGGACTCAAGACGATAGTTACCGGATAAGGCGCAGCGGTCGGACTGAACGGGGGG
TTCGTGCATACAGTCCAGCTTGGAGCGAACTGCCTACCCGGAACTGAGTGTCAGGCGTGGAA
TGAGACAAACGCGGCCATAACAGCGGAATGACACCGGTAAACCGAAAGGCAGGAACAGGAGA
GCGCACGAGGGAGCCGCCAGGGGGAAACGCCTGGTATCTTTATAGTCCTGTCGGGTTTCGCC
ACCACTGATTTGAGCGTCAGATTTCGTGATGCTTGTCAGGGGGGCGGAGCCTATGGAAAAAC
GGCTTTGCCGCGGCCCTCTCACTTCCCTGTTAAGTATCTTCCTGGCATCTTCCAGGAAATCT
CCGCCCCGTTCGTAATCCATTTCCGCTCGCCGCAGTCGAACGACCGAGCGTAGCGAGTCAGT
GAGCGAGGAAGCGGAATATATCCTTGACAGCTAGCTCAGTCCTAGGTACTGTGCTAGCACCC
GTTTTTACTAGTCTAGAAATAATAACAAAATGAGGAGGTACTGATATACATATGGCGGACAC
GATGCTGGCGGCGGTTGTACGTGAATTTGGTAAACCTCTGTCTATCGAACGCCTGCCGATCC
CAGACATTAAACCGCACCAGATCCTGGTTAAAGTTGATACCTGCGGCGTTTGTCACACTGAC
CTGCATGCAGCTCGTGGTGACTGGCCTTCCAAGCCGAACCCGCCGTTTATCCCGGGTCACGA
AGGCGTTGGCCACATCGTAGCTGTTGGCAGCCAGGTGGGCGACTTTGTTAAGACTGGTGATG
TTGTAGGTGTGCCTTGGCTGTACTCTGCATGTGGTCATTGTGAGCACTGTCTGGGTGGCTGG
GAAACTCTGTGCGAGAAGCAGGACGACACTGGTTATACCGTGAACGGTTGTTTTGCAGAATA
CGTAGTTGCCGATCCGAATTACGTTGCACACCTGCCGAGCACCATTGATCCACTGCAGGCAT
CCCCGGTGCTGTGTGCTGGCCTGACTGTTTACAAAGGTCTGAAGATGACTGAAGCTCGTCCA
GGTCAGTGGGTTGCAGTTAGCGGTGTAGGCGGCCTGGGTCAAATGGCAGTGCAGTATGCGGT
GGCGATGGGCATGAACGTTGTTGCCGTAGATATCGATGACGAAAAACTGGCCACCGCTAAAA
AGCTGGGCGCTTCCCTGACTGTGAACGCAAAAGATACCGATCCGGCTCGTTTTATCCAGCAA
CAGATCGGTGGTGCACATGGTGCTCTGGTTACCGCAGTGGGTCGTACTGCCTTCTCCCAAGC
AATGGGTTATGCCCGTCGTGGTGGTACCATCGTTCTGAACGGTCTGCCGCCGGGTGACTTTC
CGGTGTCCATTTTTGATATGGTGATGAACGGCACGACCATCCGTGGTAGCATCGTGGGCACC
CGTCTGGATATGATCGAAGCTATGGATTTCTTCGCGCGCGGCAAAGTCAAATCCGTTGTTAC
CCCGGGCAAACTGGAAAACATCAACACGATCTTCGACGACCTGCAGAACGGTCGTCTGGAAG
GTCGTACCGTTCTGGATTTTCGTAGCTGAGAGACTCCCGGGACCTCAGAGTCCGCCACACCC
GAACATAAGCTTGGCTGCGCAAAAAACCCCGCTTCGGCGGGGTTTTTTCGCGATGCATACTT

GCGATCGTTGACAGTAAAAATTGCCCGTTTGTGAACCACTTGTTTGCAAACGGGCATGACTC
CTGACTTTTATTTCTGCCTTTTATTCCTTTTACACTTGTTTTTATGAAGCCCTTCACAGAAT
TGTCCTTTCACGATTCCGTCTCTCTGATGATTGATGTTAATTAACAATGTATTCACCGAAAA
CAAACATATAAATCACAGGAGTCGCCCATGTCAGTACCCGTTCAACATCCTATGTATATCGA
TGGACAGTTTGTTACCTGGCGTGGAGACGCATGGATTGATGTGGTAAACCCTGCTACAGAGG
CTGTCATTTCCCGCATACCCGATGGTCAGGCCGAGGATGCCCGTAAGGCAATCGATGCAGCA
GAACGTGCACAACCAGAATGGGAAGCGTTGCCTGCTATTGAACGCGCCAGTTGGTTGCGCAA
AATCTCCGCCGGGATCCGCGAACGCGCCAGTGAAATCAGTGCGCTGATTGTTGAAGAAGGGG
GCAAGATCCAGCAGCTGGCTGAAGTCGAAGTGGCTTTTACTGCCGACTATATCGATTACATG
GCGGAGTGGGCACGGCGTTACGAGGGCGAGATTATTCAAAGCGATCGTCCAGGAGAAAATAT
TCTTTTGTTTAAACGTGCGCTTGGTGTGACTACCGGCATTCTGCCGTGGAACTTCCCGTTCT
TCCTCATTGCCCGCAAAATGGCTCCCGCTCTTTTGACCGGTAATACCATCGTCATTAAACCT
AGTGAATTTACGCCAAACAATGCGATTGCATTCGCCAAAATCGTCGATGAAATAGGCCTTCC
GCGCGGCGTGTTTAACCTTGTACTGGGGCGTGGTGAAACCGTTGGGCAAGAACTGGCGGGTA
ACCCAAAGGTCGCAATGGTCAGTATGACAGGCAGCGTCTCTGCAGGTGAGAAGATCATGGCG
ACTGCGGCGAAAAACATCACCAAAGTGTGTCTGGAATTGGGGGGTAAAGCACCAGCTATCGT
AATGGACGATGCCGATCTTGAACTGGCAGTCAAAGCCATCGTTGATTCACGCGTCATTAATA
GTGGGCAAGTGTGTAACTGTGCAGAACGTGTTTATGTACAGAAAGGCATTTATGATCAGTTC
GTCAATCGGCTGGGTGAAGCGATGCAGGCGGTTCAATTTGGTAACCCCGCTGAACGCAACGA
CATTGCGATGGGGCCGTTGATTAACGCCGCGGCGCTGGAAAGGGTCGAGCAAAAAGTGGCGC
GCGCAGTAGAAGAAGGGGCGAGAGTGGCGTTCGGTGGCAAAGCGGTAGAGGGGAAAGGATAT
TATTATCCGCCGACATTGCTGCTGGATGTTCGCCAGGAAATGTCGATTATGCATGAGGAAAC
CTTTGGCCCGGTGCTGCCAGTTGTCGCATTTGACACGCTGGAAGATGCTATCTCAATGGCTA
ATGACAGTGATTACGGCCTGACCTCATCAATCTATACCCAAAATCTGAACGTCGCGATGAAA
GCCATTAAAGGGCTGAAGTTTGGTGAAACTTACATCAACCGTGAAAACTTCGAAGCTATGCA
AGGCTTCCACGCCGGATGGCGTAAATCCGGTATTGGCGGCGCAGATGGTAAACATGGCTTGC
ATGAATATCTGCAGACCCAGGTGGTTTATTTACAGTCTTAATGAGTGAAAGAGGCGGAGGTT
TTTTCCTCCGCCTGTGCGCAATGGAAACAGACCAGTTATTTTTCTGCGCCTCTTCCTGACCT
GCGGCAATAATTGCACTCGCCATTTGCTGGCTAAGAATGACTTTAGTTTTCATTTTGTTATT
CCTTTTCAAGGGCTTGTTCTACAATTTCAATCCAGTGACGCACAGAGGTACGACCGGCGCTC
GCCAGATGCGTCTGGCAACCAATGTTGGCGGTGACGATCATTTCCGGTTTGCCGCTTTCCAG
CGCATTCATTTTGTTATCCCGCAGCTGGCGTGCCAGATCGGGATGCGTTAACGCATATGTTC
CCGCTGAACCGCAGCACAGATGGCTGTCGGGAACGTCCGTTAAGGTAAATCCAAGACGAAGC
AACACTTTTTCCACTTCGCCGTTCAGCTTTTGCGCATGTTGTAGGGTACACGGACAGTGGAA
GGCCAGCTTTTTATCGCCGCGAATTGCCAGTTTTTCCAGCGGTTCCTCGCGCAGAAGTTCGA
CTAAATCGACCGCCAGTTCACTGACCTGACGTGCTTTATCGGCATATAACGCATCGTTTTTC
AGCATCTGCCCATACTCTTTGACAAACGCGCCGCAGCCGCTGGCGGTTTGCAAAATTGCCTC
GGCACCTGCTTCAATCGCGGGCCACCAGGCATCAATATTATTGCGCGCCCGTGCCAGCCCTT
TCTCCTGCGCATTAAGATGATAGTCCACCGCGCCACAACAGCCTGCTTCGTTAGCTGGCATG
ACGCTGATCCCCAGACGATCCAGCACTCGCGCAGTTGCCGCGTTGGTGTTGGGCGAAAGCGT
AGGCTGGGCGCAGCCTTCCAACATTAAAACCCGACGCTTATGGCGCAGCGGCGGACGCGGTT
TAGCTTTCACCGTTTCAGCAGGCAGTTTTGCTCTGACCTGTTCCGGTAAAAACGGTCGCAGC
ACCAGCCCTACCTGCGTCAGCGCACGGAAGACCGCCGGACGCGGCACTACCTGGCGCAATCC
TTCGCGCAGTATTCGCTCCGGCAGTGGGCGTTTCACTTTCTGCTCGACAATATCACGCCCGA
TATCCAGCAAATTGTGATAGCGCACACCAGAAGGACAGGTGGTTTCACAATTACGGCAAGTG
AGGCAGCGATCGAGATGCTCCTGTGTTTTAAGCGTGACTTCGTTGCCTTCCAGCACCTGTTT
AATCAGATAGATGCGCCCGCGCGGCCCGTCCAGTTCATCGCCCAGAAGCTGATAGGTTGGGC
AGGTTGCGGTACAAAATCCGCAGTGAACACAGGCGCGCAGGATGCTGTCGGCTTCCAGCGCG
CGCGCGTTCTGCCGCATCTCTTCAGTTAATTGGGTTTGCATAGCCTGCTCCTCAAAGTTCCG
CGTACATGCGACCGGGGTTAAACACGCCGCAAGGGTCGAGCTGCTGTTTAAGCTGCTGGTGA
TAGCGGAATAAAGGAGCCGATAGCGGGGCAAAGCCACCATCTCCGGCACTAAAGCGGGTCGC

ATGACCGCCAGCGTTGCGGGCGATGCGATGGATTTGATTGTCCTCGGCTGTCGATTTCAGCC
AGCGTAACGCCCCGCCCCAGTCGATCAGTTGCTCGCCGGGTAAATCCATCATCGGCGCATCA
CTGGGTAATGAAATGCGCCATAAGGTACCTGGTAACGAGAAGAACGGCAGTTGTTGTTCACG
CAATTGCTGCCAGAACTGACCGGCAACCTCTTCGCCACCCAGCAGTTCACGCGCTGCTTTTA
CCGATCCTTCGCCGCCCTCAAGGCGGATCCACAACGCATTGTCGAAGTAACATAAGCCACTA
ATGGGTAATGGCTGGAGTTGCCACTCGGCGATTTCACTCATGGCTTCTTGCAGGCTGATTTC
CCGACGCAGGCTCAGGGAGGCGCGCGGTCGCGGTAACACTTTCATTGAGATTTCAGTGAGCA
CGCCAAGACAACCGTAGCTTCCGACCATTAACCGTGAGAGATCGTATCCGGCAACGTTTTTC
ATCACTTCGCCACCAAAACGCAGATGTTTTCCAGCGCCGGTAATGATGCGCGTGCCGAGGAC
AAAATCGCGGACCGAACCGCTCCACGGGCGACGCGGCCCCGCCAGCCCGCAGGCGACCATCC
CGCCCCAGGTGGCTTCTTCACCATAATGCGGCGGCTCACAGGGGAGCATTTGCCCCGCGCTT
TCCAGCGCCGCTTCAATTGTCACCAGCGGCGTTCCGACACGCGCGGTTATCACCAGCTCGGT
CGGGTCGTAATTAACAATGCCGCGATGACAACGAACATCCAGCGTTTGCCCGGTGACAGGGC
GACCTAAAAAGGCTTTGCTATTGCTGCCCTGAATCACCAGCGGCGTTTTATCGCTAATCGCC
TGATTCACCTGCTCCAGCAGCGCCTGGCTGTAATCACACTCGCGTAGCATCAGAAACGCTCC
AGTTCAGGGAAAGGTAAATGACCGTGATGCACATGCATGGCACCAAATTCAGCACAGCGGTG
TAGCGTGGGAATGTTTTTCCCAGGGTTCAGCAAACCATCGGGGTCAAACGCCGCCTTGACCG
CATGGAAGGTCGTGATTTCATCGCTGTTGAACTGGGCGCACATTTGATTGATTTTTTCTCGC
CCGATGCCATGTTCGCCACTGATGCTGCCGCCAACTTCAACGCAGAGTTCGAGGATCTTCCC
GCCCAGCTCTTCCGCGCGGGCAAATTCACCGGGTTCGTTGGCATCGAAAAGGATTAACGGGT
GCATGTTGCCATCTCCGGCATGAAAGACGTTGGCAACACGTAAATCATATTGCTGCGATAAA
CGGGCAATGCCTTCCAGTACGCCAGGCAGGGCGCGACGCGGGATGGTGCCATCCATGCAGTA
GTAATCCGGGGAGATACGTCCTACCGCCGGGAACGCATTTTTGCGACCGGCCCAGAAACGTA
CGCGCTCTGCTTCGTCCTGTGCCAGACGGACGTCAGTCGCGCCCGCTTTCAACAAGATGTCG
TTAACCCGCTCGCAGTCTTCCTGTACGTCAGACTCCACGCCGTCCAGCTCGCATAACAAAAT
CGCTTCGGCGTCGACGGGATAACCGGCATGAATAAAATCTTCCGCCGCGCGGATCGACAGGT
TATCCATCATCTCCAGCCCGCCGGGGATAATGCCATTGGCGATGATGTCACCAACCGCAAGT
CCGGCTTTTTCTACCGAGTCAAAGCTGGCTAACAGAACCCGCGCCACGGGCGGCTTCGGCAG
CAGTTTTACCGTCACTTCGGTGGTCACGCCGAGCATACCTTCCGATCCGGTGAACAGCGCCA
GCAGGTCAAAACCAGGTGAATCCAGCGCGTCCGATCCAAGCGTCAGTGCCTCGCCGTCCAGC
GTTTGCACTTCAATTTTCAGCAGGTTATGTACGGTCAGACCATATTTCAGGCAGTGGACGCC
GCCGGCATTTTCAGCCACATTGCCGCCAATGGAACAGGCGATTTGTGAGGAAGGGTCCGGTG
CGTAGTAGAGATTATGCGGTGCAACGGCCTGGGAGATCGCCAGGTTACGCACGCCTGGCTGC
ACGCGCGCGCGGCGACCAACGGGGTTAATGTCGAGGATCTCTTTAAAGCGCGCCATCACCAA
CAACACACCTTTTTCCAGCGGCAGCGCGCCACCAGAAAGCCCGGTGCCTGCACCACGGGTCA
CCACCGGTACACGCAGGCGATGGCAGACAGCCAGAATCGCTGTCACCTGTTCCATTTGCTTA
GGCAGAACAACCAGTAATGGACGCGTGCGATACGCGCTCAACCCGTCACACTCGTAAGGAAT
GATCTCCTCATCGGTATGCAGGATCTCAAGTCCAGGGACATGCTCACGCAGTGCCATCAGTA
CCGATGTGCGGTCGACATCGGGTAAAGCGCCATCAAGACGCTCTTCGTACAAGATGCTCATG
AGTAGGCTTCGCTTTGTTGTGTTGTGTGGCAGCTGATTTTTGCGCGCTGCTTCTGTGAACAG
TTATTAAGCGGGCTTTTCGTTTTCGTCTATCTCTTTAGCTACCGGTCAGACCATTTTTTTTC
CAGCTCTGTGACCTTGTCTTGGTTAACTCAATGTTAAATTGATGTAACATAATCACTTACGT
GATGTGCGTGTTTTGCGAGTTAAGAACAGAAAAATTGGTCCTACCTGTGCACGAGGTCCGGG
ATCTGACGCTCAGTGGAACGAAAACTCACGTTAAGGGATTTTGGTCATGAACAATAAAACTG
TCTGCTTACATAAACAGTAATACAAGGGGTGTTATGAGCCATATTCAACGGG

EGA2_pAN29_2

CATCCTATGGAACTGCCTCGGTGAGTTTTCTCCTTCATTACAGAAACGGCTTTTTCAAAAAT
ATGGTATTGATAATCCTGATATGAATAAATTGCAGTTTCATTTGATGCTCGATGAGTTTTTC
TAAAAGCTTAATTAGCTGATCTAGACGCGTGCTAGAGGCATCAAATAAAACGAAAGGCTCAG
TCGAAAGACTGGGCCTTTCGTTTTATCTGTTGTTTGTCGGTGAACGCTCTCCTGAGTAGGAC
AAATAGGTCGAGGGTGAAGTACTTGCTGACTTCCTTGAGGAACACATGATGCGTCCTACGGT
TGCTGCTACGCATATCATTGAGATGTCTGTGGGAGGAGTTGATGTGTACTCTGAGGACGATG
AGGGTTACGGTACGTCTTTCATTGAGTGGTGATTTATGCATTAGGACTGCATAGGGATGCAC
TATAGACCACGGATGGTCAGTTCTTTAAGTTACTGAAAAGACACGATAAATTAATACGACTC
ACTATAGGGAGAGGAGGGACGAAAGGTTACTATATAGATACTGAATGAATACTTATAGAGTG
CATAAAGTATGCATAATGGTGTACCTAGAGTGACCTCTAAGAATGGTGATTATATTGTATTA
GTATCACCTTAACTTAAGGCGGGATCGTCACCCTCAGCAGCGAAAGACAGCTGTCGGTCAGA
GCGTCATTGCGAAGCTGAGTGTGATCGATGCCATCAGCGAAGGGCCCAAACTCCGAGCGATT
AAGCGTTTGCTGGCTGTCACGCCTGCCTGTTGCTTGCTTGGACTTGCGATGTACGTGCTCAG
CTGTCTTTCGCTGCTGAGGGTGACGATCCCGCGAGGGCCTATGGAGTTCCTATAGGGTCCTT
TAAAATATACCATAAAAATCTGAGTGACTATCTCACAGTGTACGGACCTAAAGTTCCCCCAT
AGGGGGTACCTAAAGCCCAGCCAATCACCTAAAGTCAACCTTCGGTTGACCTTGAGGGTTCC
CTAAGGGTTGGGGATGACCCTTGGGTTTGTCTTTGGGTGTTACCTTGAGTGTCTCTCTGTGT
CCCTATCTGTTACAGTCTCCTAAAGTATCCTCCTAAAGTCACCTCCTAACGCACATTTCCCC
GAAAAGTGCCACCTGGGTCCTTTTCACTTCGGGCTCATGAGCAAATATTTTATCTGATTAAT
AAGATGATCTTCTTGAGATCGTTTTGGTCTGCGCGTAATCTCTTGCTCTGAAAACGAAAAAA
CCGCCTTGCAGGGCGGTTTTTCGAAGGTTCTCTGAGCTACCAACTCTTTGAACCGAGGTAAC
TGGCTTGGAGGAGCGCAGTCACCAAAACTTGTCCTTTCAGTTTAGCCTTAACCGGCGCATGA
CTTCAAGACTAACTCCTCTAAATCAATTACCAGTGGCTGCTGCCAGTGGTGCTTTTGCATGT
CTTTCCGGGTTGGACTCAAGACGATAGTTACCGGATAAGGCGCAGCGGTCGGACTGAACGGG
GGGTTCGTGCATACAGTCCAGCTTGGAGCGAACTGCCTACCCGGAACTGAGTGTCAGGCGTG
GAATGAGACAAACGCGGCCATAACAGCGGAATGACACCGGTAAACCGAAAGGCAGGAACAGG
AGAGCGCACGAGGGAGCCGCCAGGGGGAAACGCCTGGTATCTTTATAGTCCTGTCGGGTTTC
GCCACCACTGATTTGAGCGTCAGATTTCGTGATGCTTGTCAGGGGGGCGGAGCCTATGGAAA
AACGGCTTTGCCGCGGCCCTCTCACTTCCCTGTTAAGTATCTTCCTGGCATCTTCCAGGAAA
TCTCCGCCCCGTTCGTAAGCCATTTCCGCTCGCCGCAGTCGAACGACCGAGCGTAGCGAGTC
AGTGAGCGAGGAAGCGGAATATATCCTTGACAGCTAGCTCAGTCCTAGGTACTGTGCTAGCA
CCCGTTTTTACTAGTCTAGAAATAATAACAAAATGAGGAGGTACTGATATACATATGGCGGA
CACGATGCTGGCGGCGGTTGTACGTGAATTTGGTAAACCTCTGTCTATCGAACGCCTGCCGA
TCCCAGACATTAAACCGCACCAGATCCTGGTTAAAGTTGATACCTGCGGCGTTTGTCACACT
GACCTGCATGCAGCTCGTGGTGACTGGCCTTCCTAGCCGAACCCGCCGTTTATCCCGGGTCA
CGAAGGCGTTGGCCACATCGTAGCTGTTGGCAGCCAGGTGGGCGACTTTGTTAAGACTGGTG
ATGTTGTAGGTGTGCCTTGGCTGTACTCTGCATGTGGTCATTGTGAGCACTGTCTGGGTGGC
TGGGAAACTCTGTGCGAGAAGCAGGACGACACTGGTTATACCGTGAACGGTTGTTTTGCAGA
ATACGTAGTTGCCGATCCGAATTACGTTGCACACCTGCCGAGCACCATTGATCCACTGCAGG
CATCCCCGGTGCTGTGTGCTGGCCTGACTGTTTACAAAGGTCTGAAGATGACTGAAGCTCGT
CCAGGTCAGTGGGTTGCAGTTAGCGGTGTAGGCGGCCTGGGTCAAATGGCAGTGCAGTATGC
GGTGGCGATGGGCATGAACGTTGTTGCCGTAGATATCGATGACGAAAAACTGGCCACCGCTA
AAAAGCTGGGCGCTTCCCTGACTGTGAACGCAAAAGATACCGATCCGGCTCGTTTTATCCAG
CAACAGATCGGTGGTGCACATGGTGCTCTGGTTACCGCAGTGGGTCGTACTGCCTTCTCCCA
AGCAATGGGTTATGCCCGTCGTGGTGGTACCATCGTTCTGAACGGTCTGCCGCCGGGTGACT
TTCCGGTGTCCATTTTTGATATGGTGATGAACGGCACGACCATCCGTGGTAGCATCGTGGGC
ACCCGTCTGGATATGATCGAAGCTATGGATTTCTTCGCGCGCGGCAAAGTCAAATCCGTTGT
TACCCCGGGCAAACTGGAAAACATCAACACGATCTTCGACGACCTGCAGAACGGTCGTCTGG
AAGGTCGTACCGTTCTGGATTTTCGTAGCTGAGAGACTCCCGGGACCTCAGAGTCCGCCACA
CCCGAACATAAGCTTGGCTGCGCAAAAAACCCCGCTTCGGCGGGGTTTTTCGCGATGCATA

351 CTTGCGATCGTTGACAGTAAAAATTGCCCGTTTGTGAACCACTTGTTTGCAAACGGGCATGA
352 CTCCTGACTTTTATTTCTGCCTTTTATTCCTTTTACACTTGTTTTTATGAAGCCCTTCACAG
353 AATTGTCCTTTCACGATTCCGTCTCTCTGATGATTGATGTTAATTAACAATGTATTCACCGA
354 AAACAAACATATAAATCACAGGAGTCGCCCATGTCAGTACCCGTTCAACATCCTATGTATAT
355 CGATGGACAGTTTGTTACCTGGCGTGGAGACGCATGGATTGATGTGGTAAACCCTGCTACAG
356 AGGCTGTCATTTCCCGCATACCCGATGGTCAGGCCGAGGATGCCCGTAAGGCAATCGATGCA
357 GCAGAACGTGCACAACCAGAATGGGAAGCGTTGCCTGCTATTGAACGCGCCAGTTGGTTGCG
358 CAAAATCTCCGCCGGGATCCGCGAACGCGCCAGTGAAATCAGTGCGCTGATTGTTGAAGAAG
359 GGGGCAAGATCCAGCAGCTGGCTGAAGTCGAAGTGGCTTTTACTGCCGACTATATCGATTAC
360 ATGGCGGAGTGGGCACGGCGTTACGAGGGCGAGATTATTCAAAGCGATCGTCCAGGAGAAAA
361 TATTCTTTTGTTTAAACGTGCGCTTGGTGTGACTACCGGCATTCTGCCGTGGAACTTCCCGT
362 TCTTCCTCATTGCCCGCAAAATGGCTCCCGCTCTTTTGACCGGTAATACCATCGTCATTAAA
363 CCTAGTGAATTTACGCCAAACAATGCGATTGCATTCGCCAAAATCGTCGATGAAATAGGCCT
364 TCCGCGCGGCGTGTTTAACCTTGTACTGGGGCGTGGTGAAACCGTTGGGCAAGAACTGGCGG
365 GTAACCCAAAGGTCGCAATGGTCAGTATGACAGGCAGCGTCTCTGCAGGTGAGAAGATCATG
366 GCGACTGCGGCGAAAAACATCACCAAAGTGTGTCTGGAATTGGGGGGTAAAGCACCAGCTAT
367 CGTAATGGACGATGCCGATCTTGAACTGGCAGTCAAAGCCATCGTTGATTCACGCGTCATTA
368 ATAGTGGGCAAGTGTGTAACTGTGCAGAACGTGTTTATGTACAGAAAGGCATTTATGATCAG
369 TTCGTCAATCGGCTGGGTGAAGCGATGCAGGCGGTTCAATTTGGTAACCCCGCTGAACGCAA
370 CGACATTGCGATGGGGCCGTTGATTAACGCCGCGGCGCTGGAAAGGGTCGAGCAAAAAGTGG
371 CGCGCGCAGTAGAAGAAGGGGCGAGAGTGGCGTTCGGTGGCAAAGCGGTAGAGGGGAAAGGA
372 TATTATTATCCGCCGACATTGCTGCTGGATGTTCGCCAGGAAATGTCGATTATGCATGAGGA
373 AACCTTTGGCCCGGTGCTGCCAGTTGTCGCATTTGACACGCTGGAAGATGCTATCTCAATGG
374 CTAATGACAGTGATTACGGCCTGACCTCATCAATCTATACCCAAAATCTGAACGTCGCGATG
375 AAAGCCATTAAAGGGCTGAAGTTTGGTGAAACTTACATCAACCGTGAAAACTTCGAAGCTAT
376 GCAAGGCTTCCACGCCGGATGGCGTAAATCCGGTATTGGCGGCGCAGATGGTAAACATGGCT
377 TGCATGAATATCTGCAGACCCAGGTGGTTTATTTACAGTCTTAATGAGTGAAAGAGGCGGAG
378 GTTTTTTCCTCCGCCTGTGCGCAATGGAAACAGACCAGTTATTTTTCTGCGCCTCTTCCTGA
379 CCTGCGGCAATAATTGCACTCGCCATTTGCTGGCTAAGAATGACTTTAGTTTTCATTTTGTT
380 ATTCCTTTTCAAGGGCTTGTTCTACAATTTCAATCCAGTGACGCACAGAGGTACGACCGGCG
381 CTCGCCAGATGCGTCTGGCAACCAATGTTGGCGGTGACGATCATTTCCGGTTTGCCGCTTTC
382 CAGCGCATTCATTTTGTTATCCCGCAGCTGGCGTGCCAGATCGGGATGCGTTAACGCATATG
383 TTCCCGCTGAACCGCAGCACAGATGGCTGTCGGGAACGTCCGTTAAGGTAAATCCAAGACGA
384 AGCAACACTTTTTCCACTTCGCCGTTCAGCTTTTGCGCATGTTGTAGGGTACACGGACAGTG
385 GAAGGCCAGCTTTTTATCGCCGCGAATTGCCAGTTTTTCCAGCGGTTCCTCGCGCAGAAGTT
386 CGACTAAATCGACCGCCAGTTCACTGACCTGACGTGCTTTATCGGCATATAACGCATCGTTT
387 TTCAGCATCTGCCCATACTCTTTGACAAACGCGCCGCAGCCGCTGGCGGTTTGCAAAATTGC
388 CTCGGCACCTGCTTCAATCGCGGGCCACCAGGCATCAATATTATTGCGCGCCCGTGCCAGCC
389 CTTTCTCCTGCGCATTAAGATGATAGTCCACCGCGCCACAACAGCCTGCTTCGTTAGCTGGC
390 ATGACGCTGATCCCCAGACGATCCAGCACTCGCGCAGTTGCCGCGTTGGTGTTGGGCGAAAG
391 CGTAGGCTGGGCGCAGCCTTCCAACATTAAAACCCGACGCTTATGGCGCAGCGGCGGACGCG
392 GTTTAGCTTTCACCGTTTCAGCAGGCAGTTTTGCTCTGACCTGTTCCGGTAAAAACGGTCGC
393 AGCACCAGCCCTACCTGCGTCAGCGCACGGAAGACCGCCGGACGCGGCACTACCTGGCGCAA
394 TCCTTCGCGCAGTATTCGCTCCGGCAGTGGGCGTTTCACTTTCTGCTCGACAATATCACGCC
395 CGATATCCAGCAAATTGTGATAGCGCACACCAGAAGGACAGGTGGTTTCACAATTACGGCAA
396 GTGAGGCAGCGATCGAGATGCTCCTGTGTTTTAAGCGTGACTTCGTTGCCTTCCAGCACCTG
397 TTTAATCAGATAGATGCGCCCGCGCGGCCCGTCCAGTTCATCGCCCAGAAGCTGATAGGTTG
398 GGCAGGTTGCGGTACAAAATCCGCAGTGAACACAGGCGCGCAGGATGCTGTCGGCTTCCAGC
399 GCGCGCGCGTTCTGCCGCATCTCTTCAGTTAATTGGGTTTGCATAGCCTGCTCCTCAAAGTT
400 CCGCGTACATGCGACCGGGGTTAAACACGCCGCAAGGGTCGAGCTGCTGTTTAAGCTGCTGG
401 TGATAGCGGAATAAAGGAGCCGATAGCGGGGCAAAGCCACCATCTCCGGCACTAAAGCGGGT

CGCATGACCGCCAGCGTTGCGGGCGATGCGATGGATTTGATTGTCCTCGGCTGTCGATTTCA
GCCAGCGTAACGCCCCGCCCCAGTCGATCAGTTGCTCGCCGGGTAAATCCATCATCGGCGCA
TCACTGGGTAATGAAATGCGCCATAAGGTACCTGGTAACGAGAAGAACGGCAGTTGTTGTTC
ACGCAATTGCTGCCAGAACTGACCGGCAACCTCTTCGCCACCCAGCAGTTCACGCGCTGCTT
TTACCGATCCTTCGCCGCCCTCAAGGCGGATCCACAACGCATTGTCGAAGTAACATAAGCCA
CTAATGGGTAATGGCTGGAGTTGCCACTCGGCGATTTCACTCATGGCTTCTTGCAGGCTGAT
TTCCCGACGCAGGCTCAGGGAGGCGCGCGGTCGCGGTAACACTTTCATTGAGATTTCAGTGA
GCACGCCAAGACAACCGTAGCTTCCGACCATTAACCGTGAGAGATCGTATCCGGCAACGTTT
TTCATCACTTCGCCACCAAAACGCAGATGTTTTCCAGCGCCGGTAATGATGCGCGTGCCGAG
GACAAAATCGCGGACCGAACCGCTCCACGGGCGACGCGGCCCCGCCAGCCCGCAGGCGACCA
TCCCGCCCCAGGTGGCTTCTTCACCATAATGCGGCGGCTCACAGGGGAGCATTTGCCCCGCG
CTTTCCAGCGCCGCTTCAATTGTCACCAGCGGCGTTCCGACACGCGCGGTTATCACCAGCTC
GGTCGGGTCGTAATTAACAATGCCGCGATGACAACGAACATCCAGCGTTTGCCCGGTGACAG
GGCGACCTAAAAAGGCTTTGCTATTGCTGCCCTGAATCACCAGCGGCGTTTTATCGCTAATC
GCCTGATTCACCTGCTCCAGCAGCGCCTGGCTGTAATCACACTCGCGTAGCATCAGAAACGC
TCCAGTTCAGGGAAAGGTAAATGACCGTGATGCACATGCATGGCACCAAATTCAGCACAGCG
GTGTAGCGTGGGAATGTTTTTCCCAGGGTTCAGCAAACCATCGGGGTCAAACGCCGCCTTGA
CCGCATGGAAGGTCGTGATTTCATCGCTGTTGAACTGGGCGCACATTTGATTGATTTTTTCT
CGCCCGATGCCATGTTCGCCACTGATGCTGCCGCCAACTTCAACGCAGAGTTCGAGGATCTT
CCCGCCCAGCTCTTCCGCGCGGGCAAATTCACCGGGTTCGTTGGCATCGAAAAGGATTAACG
GGTGCATGTTGCCATCTCCGGCATGAAAGACGTTGGCAACACGTAAATCATATTGCTGCGAT
AAACGGGCAATGCCTTCCAGTACGCCAGGCAGGGCGCGACGCGGGATGGTGCCATCCATGCA
GTAGTAATCCGGGGAGATACGTCCTACCGCCGGGAACGCATTTTTGCGACCGGCCCAGAAAC
GTACGCGCTCTGCTTCGTCCTGTGCCAGACGGACGTCAGTCGCGCCCGCTTTCAACAAGATG
TCGTTAACCCGCTCGCAGTCTTCCTGTACGTCAGACTCCACGCCGTCCAGCTCGCATAACAA
AATCGCTTCGGCGTCGACGGGATAACCGGCATGAATAAAATCTTCCGCCGCGCGGATCGACA
GGTTATCCATCATCTCCAGCCCGCCGGGGATAATGCCATTGGCGATGATGTCACCAACCGCA
AGTCCGGCTTTTTCTACCGAGTCAAAGCTGGCTAACAGAACCCGCGCCACGGGCGGCTTCGG
CAGCAGTTTTACCGTCACTTCGGTGGTCACGCCGAGCATACCTTCCGATCCGGTGAACAGCG
CCAGCAGGTCAAAACCAGGTGAATCCAGCGCGTCCGATCCAAGCGTCAGTGCCTCGCCGTCC
AGCGTTTGCACTTCAATTTTCAGCAGGTTATGTACGGTCAGACCATATTTCAGGCAGTGGAC
GCCGCCGGCATTTTCAGCCACATTGCCGCCAATGGAACAGGCGATTTGTGAGGAAGGGTCCG
GTGCGTAGTAGAGATTATGCGGTGCAACGGCCTGGGAGATCGCCAGGTTACGCACGCCTGGC
TGCACGCGCGCGCGGCGACCAACGGGGTTAATGTCGAGGATCTCTTTAAAGCGCGCCATCAC
CAACAACACACCTTTTTCCAGCGGCAGCGCGCCACCAGAAAGCCCGGTGCCTGCACCACGGG
TCACCACCGGTACACGCAGGCGATGGCAGACAGCCAGAATCGCTGTCACCTGTTCCATTTGC
TTAGGCAGAACAACCAGTAATGGACGCGTGCGATACGCGCTCAACCCGTCACACTCGTAAGG
AATGATCTCCTCATCGGTATGCAGGATCTCAAGTCCAGGGACATGCTCACGCAGTGCCATCA
GTACCGATGTGCGGTCGACATCGGGTAAAGCGCCATCAAGACGCTCTTCGTACAAGATGCTC
ATGAGTAGGCTTCGCTTTGTTGTGTTGTGTGGCAGCTGATTTTTGCGCGCTGCTTCTGTGAA
CAGTTATTAAGCGGGCTTTTCGTTTTCGTCTATCTCTTTAGCTACCGGTCAGACCATTTTTT
TTCCAGCTCTGTGACCTTGTCTTGGTTAACTCAATGTTAAATTGATGTAACATAATCACTTA
CGTGATGTGCGTGTTTTGCGAGTTAAGAACAGAAAAATTGGTCCTACCTGTGCACGAGGTCC
GGGATCTGACGCTCAGTGGAACGAAAACTCACGTTAAGGATTTTGGTCATGAACAATAAAA
CTGTCTGCTTACATAAACAGTAATACAAGGGGTGTTATGAGCCATATTCAACGGGA

EGA3_pAN29_3
CCTATGGAACTGCCTCGGTGAGTTTTCTCCTTCATTACAGAAACGGCTTTTTCAAAAATATG
GTATTGATAATCCTGATATGAATAAATTGCAGTTTCATTTGATGCTCGATGAGTTTTTCTAA
AAGCTTAATTAGCTGATCTAGACGCGTGCTAGAGGCATCAAATAAAACGAAAGGCTCAGTCG
AAAGACTGGGCCTTTCGTTTTATCTGTTGTTTGTCGGTGAACGCTCTCCTGAGTAGGACAAA
TAGGTCGAGGGTGAAGTACTTGCTGACTTCCTTGAGGAACACATGATGCGTCCTACGGTTGC
TGCTACGCATATCATTGAGATGTCTGTGGGAGGAGTTGATGTGTACTCTGAGGACGATGAGG
GTTACGGTACGTCTTTCATTGAGTGGTGATTTATGCATTAGGACTGCATAGGGATGCACTAT
AGACCACGGATGGTCAGTTCTTTAAGTTACTGAAAAGACACGATAAATTAATACGACTCACT
ATAGGGAGAGGAGGGACGAAAGGTTACTATATAGATACTGAATGAATACTTATAGAGTGCAT
AAAGTATGCATAATGGTGTACCTAGAGTGACCTCTAAGAATGGTGATTATATTGTATTAGTA
TCACCTTAACTTAAGGCGGGATCGTCACCCTCAGCAGCGAAAGACAGCTGTCGGTCAGAGCG
TCATTGCGAAGCTGAGTGTGATCGATGCCATCAGCGAAGGGCCCAAACTCCGAGCGATTAAG
CGTTTGCTGGCTGTCACGCCTGCCTGTTGCTTGCTTGGACTTGCGATGTACGTGCTCAGCTG
TCTTTCGCTGCTGAGGGTGACGATCCCGCGAGGGCCTATGGAGTTCCTATAGGGTCCTTTAA
AATATACCATAAAAATCTGAGTGACTATCTCACAGTGTACGGACCTAAAGTTCCCCCATAGG
GGGTACCTAAAGCCCAGCCAATCACCTAAAGTCAACCTTCGGTTGACCTTGAGGGTTCCCTA
AGGGTTGGGGATGACCCTTGGGTTTGTCTTTGGGTGTTACCTTGAGTGTCTCTCTGTGTCCC
TATCTGTTACAGTCTCCTAAAGTATCCTCCTAAAGTCACCTCCTAACGCACATTTCCCCGAA
AAGTGCCACCTGGGTCCTTTTCACTTCGGGCTCATGAGCAAATATTTTATCTGATTAATAAG
ATGATCTTCTTGAGATCGTTTTGGTCTGCGCGTAATCTCTTGCTCTGAAAACGAAAAAACCG
CCTTGCAGGGCGGTTTTTCGAAGGTTCTCTGAGCTACCAACTCTTTGAACCGAGGTAACTGG
CTTGGAGGAGCGCAGTCACCAAAACTTGTCCTTTCAGTTTAGCCTTAACCGGCGCATGACTT
CAAGACTAACTCCTCTAAATCAATTACCAGTGGCTGCTGCCAGTGGTGCTTTTGCATGTCTT
TCCGGGTTGGACTCAAGACGATAGTTACCGGATAAGGCGCAGCGGTCGGACTGAACGGGGGG
TTCGTGCATACAGTCCAGCTTGGAGCGAACTGCCTACCCGGAACTGAGTGTCAGGCGTGGAA
TGAGACAAACGCGGCCATAACAGCGGAATGACACCGGTAAACCGAAAGGCAGGAACAGGAGA
GCGCACGAGGGAGCCGCCAGGGGGAAACGCCTGGTATCTTTATAGTCCTGTCGGGTTTCGCC
ACCACTGATTTGAGCGTCAGATTTCGTGATGCTTGTCAGGGGGGCGGAGCCTATGGAAAAAC
GGCTTTGCCGCGGCCCTCTCACTTCCCTGTTAAGTATCTTCCTGGCATCTTCCAGGAAATCT
CCGCCCCGTTCGTAAGCCATTTCCGCTCGCCGCAGTCGAACGACCGAGCGTAGCGAGTCAGT
GAGCGAGGAAGCGGAATATATCCTTGACAGCTAGCTCAGTCCTAGGTACTGTGCTAGCACCC
GTTTTTACTAGTCTAGAAATAATAACAAAATGAGGAGGTACTGATATACATATGGCGGACAC
GATGCTGGCGGCGGTTGTACGTGAATTTGGTAAACCTCTGTCTATCGAACGCCTGCCGATCC
CAGACATTAAACCGCACCAGATCCTGGTTAAAGTTGATACCTGCGGCGTTTGTCACACTGAC
CTGCATGCAGCTCGTGGTGACTGGCCTTCCAAGCCGAACCCGCCGTTTATCCCGGGTCACGA
AGGCGTTGGCCACATCGTAGCTGTTGGCAGCCAGGTGGGCGACTTTGTTAAGACTGGTGATG
TTGTAGGTGTGCCTTGGCTGTTCTCTGCATGTGGTCATTGTGAGCACTGTCTGGGTGGCTGG
GAAACTCTGTGCGAGAAGCAGGACGACACTGGTTATACCGTGAACGGTTGTTTTGCAGAATA
CGTAGTTGCCGATCCGAATTACGTTGCACACCTGCCGAGCACCATTGATCCACTGCAGGCAT
CCCCGGTGCTGTGTGCTGGCCTGACTGTTTACAAAGGTCTGAAGATGACTGAAGCTCGTCCA
GGTCAGTGGGTTGCAGTTAGCGGTGTAGGCGGCCTGGGTCAAATGGCAGTGCAGTATGCGGT
GGCGATGGGCATGAACGTTGTTGCCGTAGATATCGATGACGAAAAACTGGCCACCGCTAAAA
AGCTGGGCGCTTCCCTGACTGTGAACGCAAAAGATACCGATCCGGCTCGTTTTATCCAGCAA
CAGATCGGTGGTGCACATGGTGCTCTGGTTACCGCAGTGGGTCGTACTGCCTTCTCCCAAGC
AATGGGTTATGCCCGTCGTGGTGGTACCATCGTTCTGAACGGTCTGCCGCCGGGTGACTTTC
CGGTGTCCATTTTTGATATGGTGATGAACGGCACGACCATCCGTGGTAGCATCGTGGGCACC
CGTCTGGATATGATCGAAGCTATGGATTTCTTCGCGCGCGGCAAAGTCAAATCCGTTGTTAC
CCCGGGCAAACTGGAAAACATCAACACGATCTTCGACGACCTGCAGAACGGTCGTCTGGAAG
GTCGTACCGTTCTGGATTTTCGTAGCTGAGAGACTCCCGGGACCTCAGAGTCCGCCACACCC
GAACATAAGCTTGGCTGCGCAAAAAACCCCGCTTCGGCGGGGTTTTTTCGCGATGCATACTT

499 GCGATCGTTGACAGTAAAAATTGCCCGTTTGTGAACCACTTGTTTGCAAACGGGCATGACTC
500 CTGACTTTTATTTCTGCCTTTTATTCCTTTTACACTTGTTTTTATGAAGCCCTTCACAGAAT
501 TGTCCTTTCACGATTCCGTCTCTCTGATGATTGATGTTAATTAACAATGTATTCACCGAAAA
502 CAAACATATAAATCACAGGAGTCGCCCATGTCAGTACCCGTTCAACATCCTATGTATATCGA
503 TGGACAGTTTGTTACCTGGCGTGGAGACGCATGGATTGATGTGGTAAACCCTGCTACAGAGG
504 CTGTCATTTCCCGCATACCCGATGGTCAGGCCGAGGATGCCCGTAAGGCAATCGATGCAGCA
505 GAACGTGCACAACCAGAATGGGAAGCGTTGCCTGCTATTGAACGCGCCAGTTGGTTGCGCAA
506 AATCTCCGCCGGGATCCGCGAACGCGCCAGTGAAATCAGTGCGCTGATTGTTGAAGAAGGGG
507 GCAAGATCCAGCAGCTGGCTGAAGTCGAAGTGGCTTTTACTGCCGACTATATCGATTACATG
508 GCGGAGTGGGCACGGCGTTACGAGGGCGAGATTATTCAAAGCGATCGTCCAGGAGAAAATAT
509 TCTTTTGTTTAAACGTGCGCTTGGTGTGACTACCGGCATTCTGCCGTGGAACTTCCCGTTCT
510 TCCTCATTGCCCGCAAAATGGCTCCCGCTCTTTTGACCGGTAATACCATCGTCATTAAACCT
511 AGTGAATTTACGCCAAACAATGCGATTGCATTCGCCAAAATCGTCGATGAAATAGGCCTTCC
512 GCGCGGCGTGTTTAACCTTGTACTGGGGCGTGGTGAAACCGTTGGGCAAGAACTGGCGGGTA
513 ACCCAAAGGTCGCAATGGTCAGTATGACAGGCAGCGTCTCTGCAGGTGAGAAGATCATGGCG
514 ACTGCGGCGAAAAACATCACCAAAGTGTGTCTGGAATTGGGGGGTAAAGCACCAGCTATCGT
515 AATGGACGATGCCGATCTTGAACTGGCAGTCAAAGCCATCGTTGATTCACGCGTCATTAATA
516 GTGGGCAAGTGTGTAACTGTGCAGAACGTGTTTATGTACAGAAAGGCATTTATGATCAGTTC
517 GTCAATCGGCTGGGTGAAGCGATGCAGGCGGTTCAATTTGGTAACCCCGCTGAACGCAACGA
518 CATTGCGATGGGGCCGTTGATTAACGCCGCGGCGCTGGAAAGGGTCGAGCAAAAAGTGGCGC
519 GCGCAGTAGAAGAAGGGGCGAGAGTGGCGTTCGGTGGCAAAGCGGTAGAGGGGAAAGGATAT
520 TATTATCCGCCGACATTGCTGCTGGATGTTCGCCAGGAAATGTCGATAATGCATGAGGAAAC
521 CTTTGGCCCGGTGCTGCCAGTTGTCGCATTTGACACGCTGGAAGATGCTATCTCAATGGCTA
522 ATGACAGTGATTACGGCCTGACCTCATCAATCTATACCCAAAATCTGAACGTCGCGATGAAA
523 GCCATTAAAGGGCTGAAGTTTGGTGAAACTTACATCAACCGTGAAAACTTCGAAGCTATGCA
524 AGGCTTCCACGCCGGATGGCGTAAATCCGGTATTGGCGGCGCAGATGGTAAACATGGCTTGC
525 ATGAATATCTGCAGACCCAGGTGGTTTATTTACAGTCTTAATGAGTGAAAGAGGCGGAGGTT
526 TTTTCCTCCGCCTGTGCGCAATGGAAACAGACCAGTTATTTTTCTGCGCCTCTTCCTGACCT
527 GCGGCAATAATTGCACTCGCCATTTGCTGGCTAAGAATGACTTTAGTTTTCATTTTGTTATT
528 CCTTTTCAAGGGCTTGTTCTACAATTTCAATCCAGTGACGCACAGAGGTACGACCGGCGCTC
529 GCCAGATGCGTCTGGCAACCAATGTTGGCGGTGACGATCATTTCCGGTTTGCCGCTTTCCAG
530 CGCATTCATTTTGTTATCCCGCAGCTGGCGTGCCAGATCGGGATGCGTTAACGCATATGTTC
531 CCGCTGAACCGCAGCACAGATGGCTGTCGGGAACGTCCGTTAAGGTAAATCCAAGACGAAGC
532 AACACTTTTTCCACTTCGCCGTTCAGCTTTTGCGCATGTTGTAGGGTACACGGACAGTGGAA
533 GGCCAGCTTTTTATCGCCGCGAATTGCCAGTTTTTCCAGCGGTTCCTCGCGCAGAAGTTCGA
534 CTAAATCGACCGCCAGTTCACTGACCTGACGTGCTTTATCGGCATATAACGCATCGTTTTTC
535 AGCATCTGCCCATACTCTTTGACAAACGCGCCGCAGCCGCTGGCGGTTTGCAAAATTGCCTC
536 GGCACCTGCTTCAATCGCGGGCCACCAGGCATCAATATTATTGCGCGCCCGTGCCAGCCCTT
537 TCTCCTGCGCATTAAGATGATAGTCCACCGCGCCACAACAGCCTGCTTCGTTAGCTGGCATG
538 ACGCTGATCCCCAGACGATCCAGCACTCGCGCAGTTGCCGCGTTGGTGTTGGGCGAAAGCGT
539 AGGCTGGGCGCAGCCTTCCAACATTAAAACCCGACGCTTATGGCGCAGCGGCGGACGCGGTT
540 TAGCTTTCACCGTTTCAGCAGGCAGTTTTGCTCTGACCTGTTCCGGTAAAAACGGTCGCAGC
541 ACCAGCCCTACCTGCGTCAGCGCACGGAAGACCGCCGGACGCGGCACTACCTGGCGCAATCC
542 TTCGCGCAGTATTCGCTCCGGCAGTGGGCGTTTCACTTTCTGCTCGACAATATCACGCCCGA
543 TATCCAGCAAATTGTGATAGCGCACACCAGAAGGACAGGTGGTTTCACAATTACGGCAAGTG
544 AGGCAGCGATCGAGATGCTCCTGTGTTTTAAGCGTGACTTCGTTGCCTTCCAGCACCTGTTT
545 AATCAGATAGATGCGCCCGCGCGGCCCGTCCAGTTCATCGCCCAGAAGCTGATAGGTTGGGC
546 AGGTTGCGGTACAAAATCCGCAGTGAACACAGGCGCGCAGGATGCTGTCGGCTTCCAGCGCG
547 CGCGCGTTCTGCCGCATCTCTTCAGTTAATTGGGTTTGCATAGCCTGCTCCTCAAAGTTCCG
548 CGTACATGCGACCGGGGTTAAACACGCCGCAAGGGTCGAGCTGCTGTTTAAGCTGCTGGTGA
549 TAGCGGAATAAAGGAGCCGATAGCGGGGCAAAGCCACCATCTCCGGCACTAAAGCGGGTCGC

ATGACCGCCAGCGTTGCGGGCGATGCGATGGATTTGATTGTCCTCGGCTGTCGATTTCAGCC
AGCGTAACGCCCCGCCCCAGTCGATCAGTTGCTCGCCGGGTAAATCCATCATCGGCGCATCA
CTGGGTAATGAAATGCGCCATAAGGTACCTGGTAACGAGAAGAACGGCAGTTGTTGTTCACG
CAATTGCTGCCAGAACTGACCGGCAACCTCTTCGCCACCCAGCAGTTCACGCGCTGCTTTTA
CCGATCCTTCGCCGCCCTCAAGGCGGATCCACAACGCATTGTCGAAGTAACATAAGCCACTA
ATGGGTAATGGCTGGAGTTGCCACTCGGCGATTTCACTCATGGCTTCTTGCAGGCTGATTTC
CCGACGCAGGCTCAGGGAGGCGCGCGGTCGCGGTAACACTTTCATTGAGATTTCAGTGAGCA
CGCCAAGACAACCGTAGCTTCCGACCATTAACCGTGAGAGATCGTATCCGGCAACGTTTTTC
ATCACTTCGCCACCAAAACGCAGATGTTTTCCAGCGCCGGTAATGATGCGCGTGCCGAGGAC
AAAATCGCGGACCGAACCGCTCCACGGGCGACGCGGCCCCGCCAGCCCGCAGGCGACCATCC
CGCCCCAGGTGGCTTCTTCACCATAATGCGGCGGCTCACAGGGGAGCATTTGCCCCGCGCTT
TCCAGCGCCGCTTCAATTGTCACCAGCGGCGTTCCGACACGCGCGGTTATCACCAGCTCGGT
CGGGTCGTAATTAACAATGCCGCGATGACAACGAACATCCAGCGTTTGCCCGGTGACAGGGC
GACCTAAAAAGGCTTTGCTATTGCTGCCCTGAATCACCAGCGGCGTTTTATCGCTAATCGCC
TGATTCACCTGCTCCAGCAGCGCCTGGCTGTAATCACACTCGCGTAGCATCAGAAACGCTCC
AGTTCAGGGAAAGGTAAATGACCGTGATGCACATGCATGGCACCAAATTCAGCACAGCGGTG
TAGCGTGGGAATGTTTTTCCCAGGGTTCAGCAAACCATCGGGGTCAAACGCCGCCTTGACCG
CATGGAAGGTCGTGATTTCATCGCTGTTGAACTGGGCGCACATTTGATTGATTTTTTCTCGC
CCGATGCCATGTTCGCCACTGATGCTGCCGCCAACTTCAACGCAGAGTTCGAGGATCTTCCC
GCCCAGCTCTTCCGCGCGGGCAAATTCACCGGGTTCGTTGGCATCGAAAAGGATTAACGGGT
GCATGTTGCCATCTCCGGCATGAAAGACGTTGGCAACACGTAAATCATATTGCTGCGATAAA
CGGGCAATGCCTTCCAGTACGCCAGGCAGGGCGCGACGCGGGATGGTGCCATCCATGCAGTA
GTAATCCGGGGAGATACGTCCTACCGCCGGGAACGCATTTTTGCGACCGGCCCAGAAACGTA
CGCGCTCTGCTTCGTCCTGTGCCAGACGGACGTCAGTCGCGCCCGCTTTCAACAAGATGTCG
TTAACCCGCTCGCAGTCTTCCTGTACGTCAGACTCCACGCCGTCCAGCTCGCATAACAAAAT
CGCTTCGGCGTCGACGGGATAACCGGCATGAATAAAATCTTCCGCCGCGCGGATCGACAGGT
TATCCATCATCTCCAGCCCGCCGGGGATAATGCCATTGGCGATGATGTCACCAACCGCAAGT
CCGGCTTTTTCTACCGAGTCAAAGCTGGCTAACAGAACCCGCGCCACGGGCGGCTTCGGCAG
CAGTTTTACCGTCACTTCGGTGGTCACGCCGAGCATACCTTCCGATCCGGTGAACAGCGCCA
GCAGGTCAAAACCAGGTGAATCCAGCGCGTCCGATCCAAGCGTCAGTGCCTCGCCGTCCAGC
GTTTGCACTTCAATTTTCAGCAGGTTATGTACGGTCAGACCATATTTCAGGCAGTGGACGCC
GCCGGCATTTTCAGCCACATTGCCGCCAATGGAACAGGCGATTTGTGAGGAAGGGTCCGGTG
CGTAGTAGAGATTATGCGGTGCAACGGCCTGGGAGATCGCCAGGTTACGCACGCCTGGCTGC
ACGCGCGCGCGGCGACCAACGGGGTTAATGTCGAGGATCTCTTTAAAGCGCGCCATCACCAA
CAACACACCTTTTTCCAGCGGCAGCGCGCCACCAGAAAGCCCGGTGCCTGCACCACGGGTCA
CCACCGGTACACGCAGGCGATGGCAGACAGCCAGAATCGCTGTCACCTGTTCCATTTGCTTA
GGCAGAACAACCAGTAATGGACGCGTGCGATACGCGCTCAACCCGTCACACTCGTAAGGAAT
GATCTCCTCATCGGTATGCAGGATCTCAAGTCCAGGGACATGCTCACGCAGTGCCATCAGTA
CCGATGTGCGGTCGACATCGGGTAAAGCGCCATCAAGACGCTCTTCGTACAAGATGCTCATG
AGTAGGCTTCGCTTTGTTGTGTTGTGTGGCAGCTGATTTTTGCGCGCTGCTTCTGTGAACAG
TTATTAAGCGGGCTTTTCGTTTTCGTCTATCTCTTTAGCTACCGGTCAGACCATTTTTTTTC
CAGCTCTGTGACCTTGTCTTGGTTAACTCAATGTTAAATTGATGTAACATAATCACTTACGT
GATGTGCGTGTTTTGCGAGTTAAGAACAGAAAAATTGGTCCTACCTGTGCACGAGGTCCGGG
ATCTGACGCTCAGTGGAACGAAAACTCACGTTAAGGGATTTTGGTCATGAACAATAAAACTG
TCTGCTTACATAAACAGTAATACAAGGGGTGTTATGAGCCATATTCAACGGG

EGA4_pAN29_4
ATCCTATGGAACTGCCTCGGTGAGTTTTCTCCTTCATTACAGAAACGGCTTTTTCAAAAATA
TGGTATTGATAATCCTGATATGAATAAATTGCAGTTTCATTTGATGCTCGATGAGTTTTTCT
AAAAGCTTAATTAGCTGATCTAGACGCGTGCTAGAGGCATCAAATAAAACGAAAGGCTCAGT
CGAAAGACTGGGCCTTTCGTTTTATCTGTTGTTTGTCGGTGAACGCTCTCCTGAGTAGGACA
AATAGGTCGAGGGTGAAGTACTTGCTGACTTCCTTGAGGAACACATGATGCGTCCTACGGTT
GCTGCTACGCATATCATTGAGATGTCTGTGGGAGGAGTTGATGTGTACTCTGAGGACGATGA
GGGTTACGGTACGTCTTTCATTGAGTGGTGATTTATGCATTAGGACTGCATAGGGATGCACT
ATAGACCACGGATGGTCAGTTCTTTAAGTTACTGAAAAGACACGATAAATTAACACGACTCA
CTATAGGGAGAGGAGGGACGAAAGGTTACTATATAGATACTGAATGAATACTTATAGAGTGC
ATAAAGTATGCATAATGGTGTACCTAGAGTGACCTCTAAGAATGGTGATTATATTGTATTAG
TATCACCTTAACTTAAGGCGGGATCGTCACCCTCAGCAGCGAAAGACAGCTGTCGGTCAGAG
CGTCATTGCGAAGCTGAGTGTGATCGATGCCATCAGCGAAGGGCCCAAACTCCGAGCGATTA
AGCGTTTGCTGGCTGTCACGCCTGCCTGTTGCTTGCTTGGACTTGCGATGTACGTGCTCAGC
TGTCTTTCGCTGCTGAGGGTGACGATCCCGCGAGGGCCTATGGAGTTCCTATAGGGTCCTTT
AAAATATACCATAAAAATCTGAGTGACTATCTCACAGTGTACGGACCTAAAGTTCCCCCATA
GGGGGTACCTAAAGCCCAGCCAATCACCTAAAGTCAACCTTCGGTTGACCTTGAGGGTTCCC
TAAGGGTTGGGGATGACCCTTGGGTTTGTCTTTGGGTGTTACCTTGAGTGTCTCTCTGTGTC
CCTATCTGTTACAGTCTCCTAAAGTATCCTCCTAAAGTCACCTCCTAACGCACATTTCCCCG
AAAAGTGCCACCTGGGTCCTTTTCACTTCGGGCTCATGAGCAAATATTTTATCTGATTAATA
AGATGATCTTCTTGAGATCGTTTTGGTCTGCGCGTAATCTCTTGCTCTGAAAACGAAAAAAC
CGCCTTGCAGGGCGGTTTTTCGAAGGTTCTCTGAGCTACCAACTCTTTGAACCGAGGTAACT
GGCTTGGAGGAGCGCAGTCACCAAAACTTGTCCTTTCAGTTTAGCCTTAACCGGCGCATGAC
TTCAAGACTAACTCCTCTAAATCAATTACCAGTGGCTGCTGCCAGTGGTGCTTTTGCATGTC
TTTCCGGGTTGGACTCAAGACGATAGTTACCGGATAAGGCGCAGCGGTCGGACTGAACGGGG
GGTTCGTGCATACAGTCCAGCTTGGAGCGAACTGCCTACCCGGAACTGAGTGTCAGGCGTGG
AATGAGACAAACGCGGCCATAACAGCGGAATGACACCGGTAAACCGAAAGGCAGGAACAGGA
GAGCGCACGAGGGAGCCGCCAGGGGGAAACGCCTGGTATCTTTATAGTCCTGTCGGGTTTCG
CCACCACTGATTTGAGCGTCAGATTTCGTGATGCTTGTCAGGGGGGCGGAGCCTATGGAAAA
ACGGCTTTGCCGCGGCCCTCTCACTTCCCTGTTAAGTATCTTCCTGGCATCTTCCAGGAAAT
CTCCGCCCCGTTCGTAAGCCATTTCCGCTCGCCGCAGTCGAACGACCGAGCGTAGCGAGTCA
GTGAGCGAGGAAGCGGAATATATCCTTGACAGCTAGCTCAGTCCTAGGTACTGTGCTAGCAC
CCGTTTTTACTAGTCTAGAAATAATAACAAAATGAGGAGGTACTGATATACATATGGCGGAC
ACGATGCTGGCGGCGGTTGTACGTGAATTTGGTAAACCTCTGTCTATCGAACGCCTGCCGAT
CCCAGACATTAAACCGCACCAGATCCTGGTTAAAGTTGATACCTGCGGCGTTTGTCACACTG
ACCTGCATGCAGCTCGTGGTGACTGGCCTTCCAAGCCGAACCCGCCGTTTATCCCGGGTCAC
GAAGGCGTTGGCCACATCGTAGCTGTTGGCAGCCAGGTGGGCGACTTTGTTAAGACTGGTGA
TGTTGTAGGTGTGCCTTGGCTGTACTCTGCATGTGGTCATTGTGAGCACTGTCTGGGTGGCT
GGGAAACTCTGTGCGAGAAGCAGGACGACACTGGTTATACCGTGAACGGTTGTTTTGCAGAA
TACGTAGTTGCCGATCCGAATTACGTTGCACACCTGCCGAGCACCATTGATCCACTGCAGGC
ATCCCCGGTGCTGTGTGCTGGCCTGACTGTTTACAAAGGTCTGAAGATGACTGAAGCTCGTC
CAGGTCAGTGGGTTGCAGTTAGCGGTGTAGGCGGCCTGGGTCAAATGGCAGTGCAGTATGCG
GTGGCGATGGGCATGAACGTTGTTGCCGTAGATATCGATGACGAAAAACTGGCCACCGCTAA
AAAGCTGGGCGCTTCCCTGACTGTGAACGCAAAAGATACCGATCCGGCTCGTTTTATCCAGC
AACAGATCGGTGGTGCACATGGTGCTCTGGTTACCGCAGTGGGTCGTACTGCCTTCTCCCAA
GCAATGGGTTATGCCCGTCGTGGTGGTACCATCGTTCTGAACGGTCTGCCGCCGGGTGACTT
TCCGGTGTCCATTTTTGATATGGTGATGAACGGCACGACCATCCGTGGTAGCATCGTGGGCA
CCCGTCTGGATATGATCGAAGCTATGGATTTCTTCGCGCGCGGCAAAGTCAAATCCGTTGTT
ACCCCGGGCAAACTGGAAAACATCAACACGATCTTCGACGACCTGCAGAACGGTCGTCTGGA
AGGTCGTACCGTTCTGGATTTTCGTAGCTGAGAGACTCCCGGGACCTCAGAGTCCGCCACAC
CCGAACATAAGCTTGGCTGCGCAAAAAACCCCGCTTCGGCGGGGTTTTTTCGCGATGCATAC

TTGCGATCGTTGACAGTAAAAATTGCCCGTTTGTGAACCACTTGTTTGCAAACGGGCATGAC
TCCTGACTTTTATTTCTGCCTTTTATTCCTTTTACACTTGTTTTTATGAAGCCCTTCACAGA
ATTGTCCTTTCACGATTCCGTCTCTCTGATGATTGATGTTAATTAACAATGTATTCACCGAA
AACAAACATATAAATCACAGGAGTCGCCCATGTCAGTACCCGTTCAACATCCTATGTATATC
GATGGACAGTTTGTTACCTGGCGTGGAGACGCATGGATTGATGTGGTAAACCCTGCTACAGA
GGCTGTCATTTCCCGCATACCCGATGGTCAGGCCGAGGATGCCCGTAAGGCAATCGATGCAG
CAGAACGTGCACAACCAGAATGGGAAGCGTTGCCTGCTATTGAACGCGCCAGTTGGTTGCGC
AAAATCTCCGCCGGGATCCGCGAACGCGCCAGTGAAATCAGTGCGCTGATTGTTGAAGAAGG
GGGCAAGATCCAGCAGCTGGCTGAAGTCGAAGTGGCTTTTACTGCCGACTATATCGATTACA
TGGCGGAGTGGGCACGGCGTTACGAGGGCGAGATTATTCAAAGCGATCGTCCAGGAGAAAAT
ATTCTTTTGTTTAAACGTGCGCTTGGTGTGACTACCGGCATTCTGCCGTGGAACTTCCCGTT
CTTCCTCATTGCCCGCAAAATGGCTCCCGCTCTTTTGACCGGTAATACCATCGTCATTAAAC
CTAGTGAATTTACGCCAAACAATGCGATTGCATTCGCCAAAATCGTCGATGAAATAGGCCTT
CCGCGCGGCGTGTTTAACCTTGTACTGGGGCGTGGTGAAACCGTTGGGCAAGAACTGGCGGG
TAACCCAAAGGTCGCAATGGTCAGTATGACAGGCAGCGTCTCTGCAGGTGAGAAGATCATGG
CGACTGCGGCGAAAAACATCACCAAAGTGTGTCTGGAATTGGGGGGTAAAGCACCAGCTATC
GTAATGGACGATGCCGATCTTGAACTGGCAGTCAAAGCCATCGTTGATTCACGCGTCATTAA
TAGTGGGCAAGTGTGTAACTGTGCAGAACGTGTTTATGTACAGAAAGGCATTTATGATCAGT
TCGTCAATCGGCTGGGTGAAGCGATGCAGGCGGTTCAATTTGGTAACCCCGCTGAACGCAAC
GACATTGCGATGGGGCCGTTGATTAACGCCGCGGCGCTGGAAAGGGTCGAGCAAAAAGTGGC
GCGCGCAGTAGAAGAAGGGGCGAGAGTGGCGTTCGGTGGCAAAGCGGTAGAGGGGAAAGGAT
ATTATTATCCGCCGACATTGCTGCTGGATGTTCGCCAGGAAATGTCGATTATGCATGAGGAA
ACCTTTGGCCCGGTGCTGCCAGTTGTCGCATTTGACACGCTGGAAGATGCTATCTCAATGGC
TAATGACAGTGATTACGGCCTGACCTCATCAATCTATACCCAAAATCTGAACGTCGCGATGA
AAGCCATTAAAGGGCTGAAGTTTGGTGAAACTTACATCAACCGTGAAAACTTCGAAGCTATG
CAAGGCTTCCACGCCGGATGGCGTAAATCCGGTATTGGCGGCGCAGATGGTAAACATGGCTT
GCATGAATATCTGCAGACCCAGGTGGTTTATTTACAGTCTTAATGAGTGAAAGAGGCGGAGG
TTTTTTCCTCCGCCTGTGCGCAATGGAAACAGACCAGTTATTTTTCTGCGCCTCTTCCTGAC
CTGCGGCAATAATTGCACTCGCCATTTGCTGGCTAAGAATGACTTTAGTTTTCATTTTGTTA
TTCCTTTTCAAGGGCTTGTTCTACAATTTCAATCCAGTGACGCACAGAGGTACGACCGGCGC
TCGCCAGATGCGTCTGGCAACCAATGTTGGCGGTGACGATCATTTCCGGTTTGCCGCTTTCC
AGCGCATTCATTTTGTTATCCCGCAGCTGGCGTGCCAGATCGGGATGCGTTAACGCATATGT
TCCCGCTGAACCGCAGCACAGATGGCTGTCGGGAACGTCCGTTAAGGTAAATCCAAGACGAA
GCAACACTTTTTCCACTTCGCCGTTCAGCTTTTGCGCATGTTGTAGGGTACACGGACAGTGG
AAGGCCAGCTTTTTATCGCCGCGAATTGCCAGTTTTTCCAGCGGTTCCTCGCGCAGAAGTTC
GACTAAATCGACCGCCAGTTCACTGACCTGACGTGCTTTATCGGCATATAACGCATCGTTTT
TCAGCATCTGCCCATACTCTTTGACAAACGCGCCGCAGCCGCTGGCGGTTTGCAAAATTGCC
TCGGCACCTGCTTCAATCGCGGGCCACCAGGCATCAATATTATTGCGCGCCCGTGCCAGCCC
TTTCTCCTGCGCATTAAGATGATAGTCCACCGCGCCACAACAGCCTGCTTCGTTAGCTGGCA
TGACGCTGATCCCCAGACGATCCAGCACTCGCGCAGTTGCCGCGTTGGTGTTGGGCGAAAGC
GTAGGCTGGGCGCAGCCTTCCAACATTAAAACCCGACGCTTATGGCGCAGCGGCGGACGCGG
TTTAGCTTTCACCGTTTCAGCAGGCAGTTTTGCTCTGACCTGTTCCGGTAAAAACGGTCGCA
GCACCAGCCCTACCTGCGTCAGCGCACGGAAGACCGCCGGACGCGGCACTACCTGGCGCAAT
CCTTCGCGCAGTATTCGCTCCGGCAGTGGGCGTTTCACTTTCTGCTCGACAATATCACGCCC
GATATCCAGCAAATTGTGATAGCGCACACCAGAAGGACAGGTGGTTTCACAATTACGGCAAG
TGAGGCAGCGATCGAGATGCTCCTGTGTTTTAAGCGTGACTTCGTTGCCTTCCAGCACCTGT
TTAATCAGATAGATGCGCCCGCGCGGCCCGTCCAGTTCATCGCCCAGAAGCTGATAGGTTGG
GCAGGTTGCGGTACAAAATCCGCAGTGAACACAGGCGCGCAGGATGCTGTCGGCTTCCAGCG
CGCGCGCGTTCTGCCGCATCTCTTCAGTTAATTGGGTTTGCATAGCCTGCTCCTCAAAGTTC
CGCGTACATGCGACCGGGGTTAAACACGCCGCAAGGGTCGAGCTGCTGTTTAAGCTGCTGGT
GATAGCGGAATAAAGGAGCCGATAGCGGGGCAAAGCCACCATCTCCGGCACTAAAGCGGGTC

GCATGACCGCCAGCGTTGCGGGCGATGCGATGGATTTGATTGTCCTCGGCTGTCGATTTCAG
CCAGCGTAACGCCCCGCCCCAGTCGATCAGTTGCTCGCCGGGTAAATCCATCATCGGCGCAT
CACTGGGTAATGAAATGCGCCATAAGGTACCTGGTAACGAGAAGAACGGCAGTTGTTGTTCA
CGCAATTGCTGCCAGAACTGACCGGCAACCTCTTCGCCACCCAGCAGTTCACGCGCTGCTTT
TACCGATCCTTCGCCGCCCTCAAGGCGGATCCACAACGCATTGTCGAAGTAACATAAGCCAC
TAATGGGTAATGGCTGGAGTTGCCACTCGGCGATTTCACTCATGGCTTCTTGCAGGCTGATT
TCCCGACGCAGGCTCAGGGAGGCGCGCGGTCGCGGTAACACTTTCATTGAGATTTCAGTGAG
CACGCCAAGACAACCGTAGCTTCCGACCATTAACCGTGAGAGATCGTATCCGGCAACGTTTT
TCATCACTTCGCCACCAAAACGCAGATGTTTTCCAGCGCCGGTAATGATGCGCGTGCCGAGG
ACAAAATCGCGGACCGAACCGCTCCACGGGCGACGCGGCCCCGCCAGCCCGCAGGCGACCAT
CCCGCCCCAGGTGGCTTCTTCACCATAATGCGGCGGCTCACAGGGGAGCATTTGCCCCGCGC
TTTCCAGCGCCGCTTCAATTGTCACCAGCGGCGTTCCGACACGCGCGGTTATCACCAGCTCG
GTCGGGTCGTAATTAACAATGCCGCGATGACAACGAACATCCAGCGTTTGCCCGGTGACAGG
GCGACCTAAAAAGGCTTTGCTATTGCTGCCCTGAATCACCAGCGGCGTTTTATCGCTAATCG
CCTGATTCACCTGCTCCAGCAGCGCCTGGCTGTAATCACACTCGCGTAGCATCAGAAACGCT
CCAGTTCAGGGAAAGGTAAATGACCGTGATGCACATGCATGGCACCAAATTCAGCACAGCGG
TGTAGCGTGGGAATGTTTTTCCCAGGGTTCAGCAAACCATCGGGGTCAAACGCCGCCTTGAC
CGCATGGAAGGTCGTGATTTCATCGCTGTTGAACTGGGCGCACATTTGATTGATTTTTTCTC
GCCCGATGCCATGTTCGCCACTGATGCTGCCGCCAACTTCAACGCAGAGTTCGAGGATCTTC
CCGCCCAGCTCTTCCGCGCGGGCAAATTCACCGGGTTCGTTGGCATCGAAAAGGATTAACGG
GTGCATGTTGCCATCTCCGGCATGAAAGACGTTGGCAACACGTAAATCATATTGCTGCGATA
AACGGGCAATGCCTTCCAGTACGCCAGGCAGGGCGCGACGCGGGATGGTGCCATCCATGCAG
TAGTAATCCGGGGAGATACGTCCTACCGCCGGGAACGCATTTTTGCGACCGGCCCAGAAACG
TACGCGCTCTGCTTCGTCCTGTGCCAGACGGACGTCAGTCGCGCCCGCTTTCAACAAGATGT
CGTTAACCCGCTCGCAGTCTTCCTGTACGTCAGACTCCACGCCGTCCAGCTCGCATAACAAA
ATCGCTTCGGCGTCGACGGGATAACCGGCATGAATAAAATCTTCCGCCGCGCGGATCGACAG
GTTATCCATCATCTCCAGCCCGCCGGGGATAATGCCATTGGCGATGATGTCACCAACCGCAA
GTCCGGCTTTTTCTACCGAGTCAAAGCTGGCTAACAGAACCCGCGCCACGGGCGGCTTCGGC
AGCAGTTTTACCGTCACTTCGGTGGTCACGCCGAGCATACCTTCCGATCCGGTGAACAGCGC
CAGCAGGTCAAAACCAGGTGAATCCAGCGCGTCCGATCCAAGCGTCAGTGCCTCGCCGTCCA
GCGTTTGCACTTCAATTTTCAGCAGGTTATGTACGGTCAGACCATATTTCAGGCAGTGGACG
CCGCCGGCATTTTCAGCCACATTGCCGCCAATGGAACAGGCGATTTGTGAGGAAGGGTCCGG
TGCGTAGTAGAGATTATGCGGTGCAACGGCCTGGGAGATCGCCAGGTTACGCACGCCTGGCT
GCACGCGCGCGCGGCGACCAACGGGGTTAATGTCGAGGATCTCTTTAAAGCGCGCCATCACC
AACAACACACCTTTTTCCAGCGGCAGCGCGCCACCAGAAAGCCCGGTGCCTGCACCACGGGT
CACCACCGGTACACGCAGGCGATGGCAGACAGCCAGAATCGCTGTCACCTGTTCCATTTGCT
TAGGCAGAACAACCAGTAATGGACGCGTGCGATACGCGCTCAACCCGTCACACTCGTAAGGA
ATGATCTCCTCATCGGTATGCAGGATCTCAAGTCCAGGGACATGCTCACGCAGTGCCATCAG
TACCGATGTGCGGTCGACATCGGGTAAAGCGCCATCAAGACGCTCTTCGTACAAGATGCTCA
TGAGTAGGCTTCGCTTTGTTGTGTTGTGTGGCAGCTGATTTTTGCGCGCTGCTTCTGTGAAC
AGTTATTAAGCGGGCTTTTCGTTTTCGTCTATCTCTTTAGCTACCGGTCAGACCATTTTTTT
TCCAGCTCTGTGACCTTGTCTTGGTTAACTCAATGTTAAATTGATGTAACATAATCACTTAC
GTGATGTGCGTGTTTTGCGAGTTAAGAACAGAAAAATTGGTCCTACCTGTGCACGAGGTCCG
GGATCTGACGCTCAGTGGAACGAAAACTCACGTTAAGGGATTTTGGTCATGAACAATAAAAC
TGTCTGCTTACATAAACAGTAATACAAGGGGTGTTATGAGCCATATTCAACGGG

EGA5_pAN29_5
CATCCTATGGAACTGCCTCGGTGAGTTTTCTCCTTCATTACAGAAACGGCTTTTTCAAAAAT
ATGGTATTGATAATCCTGATATGAATAAATTGCAGTTTCATTTGATGCTCGATGAGTTTTTC
TAAAAGCTTAATTAGCTGATCTAGACGCGTGCTAGAGGCATCAAATAAAACGAAAGGCTCAG
TCGAAAGACTGGGCCTTTCGTTTTATCTGTTGTTTGTCGGTGAACGCTCTCCTGAGTAGGAC
AAATAGGTCGAGGGTGAAGTACTTGCTGACTTCCTTGAGGAACACATGATGCGTCCTACGGT
TGCTGCTACGCATATCATTGAGATGTCTGTGGGAGGAGTTGATGTGTACTCTGAGGACGATG
AGGGTTACGGTACGTCTTTCATTGAGTGGTGATTTATGCATTAGGACTGCATAGGGATGCAC
TATAGACCACGGATGGTCAGTTCTTTAAGTTACTGAAAAGACACGATAAATTAATACGACTC
ACTATAGGGAGAGGAGGGACGAAAGGTTACTATATAGATACTGAATGAATACTTATAGAGTG
CATAAAGTATGCATAATGGTGTACCTAGAGTGACCTCTAAGAATGGTGATTATATTGTATTA
GTATCACCTTAACTTAAGGCGGGATCGTCACCCTCAGCAGCGAAAGACAGCTGTCGGTCAGA
GCGTCATTGCGAAGCTGAGTGTGATCGATGCCATCAGCGAAGGGCCCAAACTCCGAGCGATT
AAGCGTTTGCTGGCTGTCACGCCTGCCTGTTGCTTGCTTGGACTTGCGATGTACGTGCTCAG
CTGTCTTTCGCTGCTGAGGGTGACGATCCCGCGAGGGCCTATGGAGTTCCTATAGGGTCCTT
TAAAATATACCATAAAAATCTGAGTGACTATCTCACAGTGTACGGACCTAAAGTTCCCCCAT
AGGGGGTACCTAAAGCCCAGCCAATCACCTAAAGTCAACCTTCGGTTGACCTTGAGGGTTCC
CTAAGGGTTGGGGATGACCCTTGGGTTTGTCTTTGGGTGTTACCTTGAGTGTCTCTCTGTGT
CCCTATCTGTTACAGTCTCCTAAAGTATCCTCCTAAAGTCACCTCCTAACGCACATTTCCCC
GAAAAGTGCCACCTGGGTCCTTTTCACTTCGGGCTCATGAGCAAATATTTTATCTGATTAAT
AAGATGATCTTCTTGAGATCGTTTTGGTCTGCGCGTAATCTCTTGCTCTGAAAACGAAAAAA
CCGCCTTGCAGGGCGGTTTTTCGAAGGTTCTCTGAGCTACCAACTCTTTGAACCGAGGTAAC
TGGCTTGGAGGAGCGCAGTCACCAAAACTTGTCCTTTCAGTTTAGCCTTAACCGGCGCATGA
CTTCAAGACTAACTCCTCTAAATCAATTACCAGTGGCTGCTGCCAGTGGTGCTTTTGCATGT
CTTTCCGGGTTGGACTCAAGACGATAGTTACCGGATAAGGCGCAGCGGTCGGACTGAACGGG
GGGTTCGTGCATACAGTCCAGCTTGGAGCGAACTGCCTACCCGGAACTGAGTGTCAGGCGTG
GAATGAGACAAACGCGGCCATAACAGCGGAATGACACCGGTAAACCGAAAGGCAGGAACAGG
AGAGCGCACGAGGGAGCCGCCAGGGGGAAACGCCTGGTATCTTTATAGTCCTGTCGGGTTTC
GCCACCACTGATTTGAGCGTCAGATTTCGTGATGCTTGTCAGGGGGGCGGAGCCTATGGAAA
AACGGCTTTGCCGCGGCCCTCTCACTTCCCTGTTAAGTATCTTCCTGGCATCTTCCAGGAAA
TCTCCGCCCCGTTCGTAAGCCATTTCCGCTCGCCGCAGTCGAACGACCGAGCGTAGCGAGTC
AGTGAGCGAGGAAGCGGAATATATCCTTGACAGCTAGCTCAGTCCTAGGTACTGTGCTAGCA
CCCGTTTTTACTAGTCTAGAAATAATAACAAAATGAGGAGGTACTGATATACATATGGCGGA
CACGATGCTGGCGGCGGTTGTACGTGAATTTGGTAAACCTCTGTCTATCGAACGCCTGCCGA
TCCCAGACATTAAACCGCACCAGATCCTGGTTAAAGTTGATACCTGCGGCGTTTGTCACACT
GACCTGCATGCAGCTCGTGGTGACTGGCCTTCCAAGCCGAACCCGCCGTTTATCCCGGGTCA
CGAAGGCGTTGGCCACATCGTAGCTGTTGGCAGCCAGGTGGGCGACTTTGTTAAGACTGGTG
ATGTTGTAGGTGTGCCTTGGCTGTACTCTGCATGTGGTCATTGTGAGCACTGTCTGGGTGGC
TGGGAAACTCTGTGCGAGAAGCAGGACGACACTGGTTATACCGTGAACGGTTGTTTTGCAGA
ATACGTAGTTGCCGATCCGAATTACGTTGCACACCTGCCGAGCACCATTGATCCACTGCAGG
CATCCCCGGTGCTGTGTGCTGGCCTGACTGTTTACAAAGGTCTGAAGATGACTGAAGCTCGT
CCAGGTCAGTGGGTTGCAGTTAGCGGTGTAGGCGGCCTGGGTCAAATGGCAGTGCAGTATGC
GGTGGCGATGGGCATGAACGTTGTTGCCGTAGATATCGATGACGAAAAACTGGCCACCGCTA
AAAAGCTGGGCGCTTCCCTGACTGTGAACGCAAAAGATACCGATCCGGCTCGTTTTATCCAG
CAACAGATCGGTGGTGCACATGGTGCTCTGGTTACCGCAGTGGGTCGTACTGCCTTCTCCCA
AGCAATGGGTTATGCCCGTCGTGGTGGTACCATCGTTCTGAACGGTCTGCCGCCGGGTGACT
TTCCGGTGTCCATTTTTGATATGGTGATGAACGGCACGACCATCCGTGGTAGCATCGTGGGC
ACCCGTCTGGATATGATCGAAGCTATGGATTTCTTCGCGCGCGGCAAAGTCAAATCCGTTGT
TACCCCGGGCAAACTGGAAAACATCAACACGATCTTCGACGACCTGCAGAACGGTCGTCTGG
AAGGTCGTACCGTTCTGGATTTTCGTAGCTGAGAGACTCCCGGGACCTCAGAGTCCGCCACC
CCCGAACATAAGCTTGGCTGCGCAAAAAACCCCGCTTCGGCGGGGTTTTTTCGCGATGCATA

795 CTTGCGATCGTTGACAGTAAAAATTGCCCGTTTGTGAACCACTTGTTTGCAAACGGGCATGA
796 CTCCTGACTTTTATTTCTGCCTTTTATTCCTTTTACACTTGTTTTTATGAAGCCCTTCACAG
797 AATTGTCCTTTCACGATTCCGTCTCTCTGATGATTGATGTTAATTAACAATGTATTCACCGA
798 AAACAAACATATAAATCACAGGAGTCGCCCATGTCAGTACCCGTTCAACATCCTATGTATAT
799 CGATGGACAGTTTGTTACCTGGCGTGGAGACGCATGGATTGATGTGGTAAACCCTGCTACAG
800 AGGCTGTCATTTCCCGCATACCCGATGGTCAGGCCGAGGATGCCCGTAAGGCAATCGATGCA
801 GCAGAACGTGCACAACCAGAATGGGAAGCGTTGCCTGCTATTGAACGCGCCAGTTGGTTGCG
802 CAAAATCTCCGCCGGGATCCGCGAACGCGCCAGTGAAATCAGTGCGCTGATTGTTGAAGAAG
803 GGGGCAAGATCCAGCAGCTGGCTGAAGTCGAAGTGGCTTTTACTGCCGACTATATCGATTAC
804 ATGGCGGAGTGGGCACGGCGTTACGAGGGCGAGATTATTCAAAGCGATCGTCCAGGAGAAAA
805 TATTCTTTTGTTTAAACGTGCGCTTGGTGTGACTACCGGCATTCTGCCGTGGAACTTCCCGT
806 TCTTCCTCATTGCCCGCAAAATGGCTCCCGCTCTTTTGACCGGTAATACCATCGTCATTAAA
807 CCTAGTGAATTTACGCCAAACAATGCGATTGCATTCGCCAAAATCGTCGATGAAATAGGCCT
808 TCCGCGCGGCGTGTTTAACCTTGTACTGGGGCGTGGTGAAACCGTTGGGCAAGAACTGGCGG
809 GTAACCCAAAGGTCGCAATGGTCAGTATGACAGGCAGCGTCTCTGCAGGTGAGAAGATCATG
810 GCGACTGCGGCGAAAAACATCACCAAAGTGTGTCTGGAATTGGGGGGTAAAGCACCAGCTAT
811 CGTAATGGACGATGCCGATCTTGAACTGGCAGTCAAAGCCATCGTTGATTCACGCGTCATTA
812 ATAGTGGGCAAGTGTGTAACTGTGCAGAACGTGTTTATGTACAGAAAGGCATTTATGATCAG
813 TTCGTCAATCGGCTGGGTGAAGCGATGCAGGCGGTTCAATTTGGTAACCCCGCTGAACGCAA
814 CGACATTGCGATGGGGCCGTTGATTAACGCCGCGGCGCTGGAAAGGGTCGAGCAAAAAGTGG
815 CGCGCGCAGTAGAAGAAGGGGCGAGAGTGGCGTTCGGTGGCAAAGCGGTAGAGGGGAAAGGA
816 TATTATTATCCGCCGACATTGCTGCTGGATGTTCGCCAGGAAATGTCGATTATGCATGAGGA
817 AACCTTTGGCCCGGTGCTGCCAGTTGTCGCATTTGACACGCTGGAAGATGCTATCTCAATGG
818 CTAATGACAGTGATTACGGCCTGACCTCATCAATCTATACCCAAAATCTGAACGTCGCGATG
819 AAAGCCATTAAAGGGCTGAAGTTTGGTGAAACTTACATCAACCGTGAAAACTTCGAAGCTAT
820 GCAAGGCTTCCACGCCGGATGGCGTAAATCCGGTATTGGCGGCGCAGATGGTAAACATGGCT
821 TGCATGAATATCTGCAGACCCAGGTGGTTTATTTACAGTCTTAATGAGTGAAAGAGGCGGAG
822 GTTTTTTCCTCCGCCTGTGCGCAATGGAAACAGACCAGTTATTTTTCTGCGCCTCTTCCTGA
823 CCTGCGGCAATAATTGCACTCGCCATTTGCTGGCTAAGAATGACTTTAGTTTTCATTTTGTT
824 ATTCCTTTTCAAGGGCTTGTTCTACAATTTCAATCCAGTGACGCACAGAGGTACGACCGGCG
825 CTCGCCAGATGCGTCTGGCAACCAATGTTGGCGGTGACGATCATTTCCGGTTTGCCGCTTTC
826 CAGCGCATTCATTTTGTTATCCCGCAGCTGGCGTGCCAGATCGGGATGCGTTAACGCATATG
827 TTCCCGCTGAACCGCAGCACAGATGGCTGTCGGGAACGTCCGTTAAGGTAAATCCAAGACGA
828 AGCAACACTTTTTCCACTTCGCCGTTCAGCTTTTGCGCATGTTGTAGGGTACACGGACAGTG
829 GAAGGCCAGCTTTTTATCGCCGCGAATTGCCAGTTTTTCCAGCGGTTCCTCGCGCAGAAGTT
830 CGACTAAATCGACCGCCAGTTCACTGACCTGACGTGCTTTATCGGCATATAACGCATCGTTT
831 TTCAGCATCTGCCCATACTCTTTGACAAACGCGCCGCAGCCGCTGGCGGTTTGCAAAATTGC
832 CTCGGCACCTGCTTCAATCGCGGGCCACCAGGCATCAATATTATTGCGCGCCCGTGCCAGCC
833 CTTTCTCCTGCGCATTAAGATGATAGTCCACCGCGCCACAACAGCCTGCTTCGTTAGCTGGC
834 ATGACGCTGATCCCCAGACGATCCAGCACTCGCGCAGTTGCCGCGTTGGTGTTGGGCGAAAG
835 CGTAGGCTGGGCGCAGCCTTCCAACATTAAAACCCGACGCTTATGGCGCAGCGGCGGACGCG
836 GTTTAGCTTTCACCGTTTCAGCAGGCAGTTTTGCTCTGACCTGTTCCGGTAAAAACGGTCGC
837 AGCACCAGCCCTACCTGCGTCAGCGCACGGAAGACCGCCGGACGCGGCACTACCTGGCGCAA
838 TCCTTCGCGCAGTATTCGCTCCGGCAGTGGGCGTTTCACTTTCTGCTCGACAATATCACGCC
839 CGATATCCAGCAAATTGTGATAGCGCACACCAGAAGGACAGGTGGTTTCACAATTACGGCAA
840 GTGAGGCAGCGATCGAGATGCTCCTGTGTTTTAAGCGTGACTTCGTTGCCTTCCAGCACCTG
841 TTTAATCAGATAGATGCGCCCGCGCGGCCCGTCCAGTTCATCGCCCAGAAGCTGATAGGTTG
842 GGCAGGTTGCGGTACAAAATCCGCAGTGAACACAGGCGCGCAGGATGCTGTCGGCTTCCAGC
843 GCGCGCGCGTTCTGCCGCATCTCTTCAGTTAATTGGGTTTGCATAGCCTGCTCCTCAAAGTT
844 CCGCGTACATGCGACCGGGGTTAAACACGCCGCAAGGGTCGAGCTGCTGTTTAAGCTGCTGG
845 TGATAGCGGAATAAAGGAGCCGATAGCGGGGCAAAGCCACCATCTCCGGCACTAAAGCGGGT

CGCATGACCGCCAGCGTTGCGGGCGATGCGATGGATTTGATTGTCCTCGGCTGTCGATTTCA
GCCAGCGTAACGCCCCGCCCCAGTCGATCAGTTGCTCGCCGGGTAAATCCATCATCGGCGCA
TCACTGGGTAATGAAATGCGCCATAAGGTACCTGGTAACGAGAAGAACGGCAGTTGTTGTTC
ACGCAATTGCTGCCAGAACTGACCGGCAACCTCTTCGCCACCCAGCAGTTCACGCGCTGCTT
TTACCGATCCTTCGCCGCCCTCAAGGCGGATCCACAACGCATTGTCGAAGTAACATAAGCCA
CTAATGGGTAATGGCTGGAGTTGCCACTCGGCGATTTCACTCATGGCTTCTTGCAGGCTGAT
TTCCCGACGCAGGCTCAGGGAGGCGCGCGGTCGCGGTAACACTTTCATTGAGATTTCAGTGA
GCACGCCAAGACAACCGTAGCTTCCGACCATTAACCGTGAGAGATCGTATCCGGCAACGTTT
TTCATCACTTCGCCACCAAAACGCAGATGTTTTCCAGCGCCGGTAATGATGCGCGTGCCGAG
GACAAAATCGCGGACCGAACCGCTCCACGGGCGACGCGGCCCCGCCAGCCCGCAGGCGACCA
TCCCGCCCCAGGTGGCTTCTTCACCATAATGCGGCGGCTCACAGGGGAGCATTTGCCCCGCG
CTTTCCAGCGCCGCTTCAATTGTCACCAGCGGCGTTCCGACACGCGCGGTTATCACCAGCTC
GGTCGGGTCGTAATTAACAATGCCGCGATGACAACGAACATCCAGCGTTTGCCCGGTGACAG
GGCGACCTAAAAAGGCTTTGCTATTGCTGCCCTGAATCACCAGCGGCGTTTTATCGCTAATC
GCCTGATTCACCTGCTCCAGCAGCGCCTGGCTGTAATCACACTCGCGTAGCATCAGAAACGC
TCCAGTTCAGGGAAAGGTAAATGACCGTGATGCACATGCATGGCACCAAATTCAGCACAGCG
GTGTAGCGTGGGAATGTTTTTCCCAGGGTTCAGCAAACCATCGGGGTCAAACGCCGCCTTGA
CCGCATGGAAGGTCGTGATTTCATCGCTGTTGAACTGGGCGCACATTTGATTGATTTTTTCT
CGCCCGATGCCATGTTCGCCACTGATGCTGCCGCCAACTTCAACGCAGAGTTCGAGGATCTT
CCCGCCCAGCTCTTCCGCGCGGGCAAATTCACCGGGTTCGTTGGCATCGAAAAGGATTAACG
GGTGCATGTTGCCATCTCCGGCATGAAAGACGTTGGCAACACGTAAATCATATTGCTGCGAT
AAACGGGCAATGCCTTCCAGTACGCCAGGCAGGGCGCGACGCGGGATGGTGCCATCCATGCA
GTAGTAATCCGGGGAGATACGTCCTACCGCCGGGAACGCATTTTTGCGACCGGCCCAGAAAC
GTACGCGCTCTGCTTCGTCCTGTGCCAGACGGACGTCAGTCGCGCCCGCTTTCAACAAGATG
TCGTTAACCCGCTCGCAGTCTTCCTGTACGTCAGACTCCACGCCGTCCAGCTCGCATAACAA
AATCGCTTCGGCGTCGACGGGATAACCGGCATGAATAAAATCTTCCGCCGCGCGGATCGACA
GGTTATCCATCATCTCCAGCCCGCCGGGGATAATGCCATTGGCGATGATGTCACCAACCGCA
AGTCCGGCTTTTTCTACCGAGTCAAAGCTGGCTAACAGAACCCGCGCCACGGGCGGCTTCGG
CAGCAGTTTTACCGTCACTTCGGTGGTCACGCCGAGCATACCTTCCGATCCGGTGAACAGCG
CCAGCAGGTCAAAACCAGGTGAATCCAGCGCGTCCGATCCAAGCGTCAGTGCCTCGCCGTCC
AGCGTTTGCACTTCAATTTTCAGCAGGTTATGTACGGTCAGACCATATTTCAGGCAGTGGAC
GCCGCCGGCATTTTCAGCCACATTGCCGCCAATGGAACAGGCGATTTGTGAGGAAGGGTCCG
GTGCGTAGTAGAGATTATGCGGTGCAACGGCCTGGGAGATCGCCAGGTTACGCACGCCTGGC
TGCACGCGCGCGCGGCGACCAACGGGGTTAATGTCGAGGATCTCTTTAAAGCGCGCCATCAC
CAACAACACACCTTTTTCCAGCGGCAGCGCGCCACCAGAAAGCCCGGTGCCTGCACCACGGG
TCACCACCGGTACACGCAGGCGATGGCAGACAGCCAGAATCGCTGTCACCTGTTCCATTTGC
TTAGGCAGAACAACCAGTAATGGACGCGTGCGATACGCGCTCAACCCGTCACACTCGTAAGG
AATGATCTCCTCATCGGTATGCAGGATCTCAAGTCCAGGGACATGCTCACGCAGTGCCATCA
GTACCGATGTGCGGTCGACATCGGGTAAAGCGCCATCAAGACGCTCTTCGTACAAGATGCTC
ATGAGTAGGCTTCGCTTTGTTGTGTTGTGTGGCAGCTGATTTTTGCGCGCTGCTTCTGTGAA
CAGTTATTAAGCGGGCTTTTCGTTTTCGTCTATCTCTTTAGCTACCGGTCAGACCATTTTTT
TTCCAGCTCTGTGACCTTGTCTTGGTTAACTCAATGTTAAATTGATGTAACATAATCACTTA
CGTGATGTGCGTGTTTTGCGAGTTAAGAACAGAAAAATTGGTCCTACCTGTGCACGAGGTCC
GGGATCTGACGCTCAGTGGAACGAAAACTCACGTTAAGGGATTTTGGTCATGAACAATAAAA
CTGTCTGCTTACATAAACAGTAATACAAGGGGTGTTATGAGCCATATTCAACGGG

EGA6_pAN29_6
ATCCTATGGAACTGCCTCGGTGAGTTTTCTCCTTCATTACAGAAACGGCTTTTTCAAAAATA
TGGTATTGATAATCCTGATATGAATAAATTGCAGTTTCATTTGATGCTCGATGAGTTTTTCT
AAAAGCTTAATTAGCTGATCTAGACGCGTGCTAGAGGCATCAAATAAAACGAAAGGCTCAGT
CGAAAGACTGGGCCTTTCGTTTTATCTGTTGTTTGTCGGTGAACGCTCTCCTGAGTAGGACA
AATAGGTCGAGGGTGAAGTACTTGCTGACTTCCTTGAGGAACACATGATGCGTCCTACGGTT
GCTGCTACGCATATCATTGAGATGTCTGTGGGAGGAGTTGATGTGTACTCTGAGGACGATGA
GGGTTACGGTACGTCTTTCATTGAGTGGTGATTTATGCATTAGGACTGCATAGGGATGCACT
ATAGACCACGGATGGTCAGTTCTTTAAGTTACTGAAAAGACACGATAAATTAATACGACTCA
CTATAGGGAGAGGAGGGACGAAAGGTTACTATATAGATACTGAATGAATACTTATAGAGTGC
ATAAAGTATGCATAATGGTGTACCTAGAGTGACCTCTAAGAATGGTGATTATATTGTATTAG
TATCACCTTAACTTAAGGCGGGATCGTCACCCTCAGCAGCGAAAGACAGCTGAGCACGTACA
TCGCAAGTCCAAGCAAGCAACAGGCAGGCGTGACAGCCAGCAAACGCTTAATCGCTCGGAGT
TTGGGCCCTTCGCTGATGGCATCGATCACACTCAGCTTCGCAATGACGCTCTGACCGACAGC
TGTCTTTCGCTGCTGAGGGTGACGATCCCGCGAGGGCCTATGGAGTTCCTATAGGGTCCTTT
AAAATATACCATAAAAATCTGAGTGACTATCTCACAGTGTACGGACCTAAAGTTCCCCCATA
GGGGGTACCTAAAGCCCAGCCAATCACCTAAAGTCAACCTTCGGTTGACCTTGAGGGTTCCC
TAAGGGTTGGGGATGACCCTTGGGTTTGTCTTTGGGTGTTACCTTGAGTGTCTCTCTGTGTC
CCTATCTGTTACAGTCTCCTAAAGTATCCTCCTAAAGTCACCTCCTAACGCACATTTCCCCG
AAAAGTGCCACCTGGGTCCTTTTCACTTCGGGCTCATGAGCAAATATTTTATCTGATTAATA
AGATGATCTTCTTGAGATCGTTTTGGTCTGCGCGTAATCTCTTGCTCTGAAAACGAAAAAAC
CGCCTTGCAGGGCGGTTTTTCGAAGGTTCTCTGAGCTACCAACTCTTTGAACCGAGGTAACT
GGCTTGGAGGAGCGCAGTCACCAAAACTTGTCCTTTCAGTTTAGCCTTAACCGGCGCATGAC
TTCAAGACTAACTCCTCTAAATCAATTACCAGTGGCTGCTGCCAGTGGTGCTTTTGCATGTC
TTTCCGGGTTGGACTCAAGACGATAGTTACCGGATAAGGCGCAGCGGTCGGACTGAACGGGG
GGTTCGTGCATACAGTCCAGCTTGGAGCGAACTGCCTACCCGGAACTGAGTGTCAGGCGTGG
AATGAGACAAACGCGGCCATAACAGCGGAATGACACCGGTAAACCGAAAGGCAGGAACAGGA
GAGCGCACGAGGGAGCCGCCAGGGGGAAACGCCTGGTATCTTTATAGTCCTGTCGGGTTTCG
CCACCACTGATTTGAGCGTCAGATTTCGTGATGCTTGTCAGGGGGGCGGAGCCTATGGAAAA
ACGGCTTTGCCGCGGCCCTCTCACTTCCCTGTTAAGTATCTTCCTGGCATCTTCCAGGAAAT
CTCCGCCCCGTTCGTAAGCCATTTCCGCTCGCCGCAGTCGAACGACCGAGCGTAGCGAGTCA
GTGAGCGAGGAAGCGGAATATATCCTTGACAGCTAGCTCAGTCCTAGGTACTGTGCTAGCAC
CCGTTTTTACTAGTCTAGAAATAATAACAAAATGAGGAGGTACTGATATACATATGGCGGAC
ACGATGCTGGCGGCGGTTGTACGTGAATTTGGTAAACCTCTGTCTATCGAACGCCTGCCGAT
CCCAGACATTAAACCGCACCAGATCCTGGTTAAAGTTGATACCTGCGGCGTTTGTCACACTG
ACCTGCATGCAGCTCGTGGTGACTGGCCTTCCAAGCCGAACCCGCCGTTTATCCCGGGTCAC
GAAGGCGTTGGCCACATCGTAGCTGTTGGCAGCCAGGTGGGCGACTTTGTTAAGACTGGTGA
TGTTGTAGGTGTGCCTTGGCTGTACTCTGCATGTGGTCATTGTGAGCACTGTCTGGGTGGCT
GGGAAACTCTGTGCGAGAAGCAGGACGACACTGGTTATACCGTGAACGGTTGTTTTGCAGAA
TACGTAGTTGCCGATCCGAATTACGTTGCACACCTGCCGAGCACCATTGATCCACTGCAGGC
ATCCCCGGTGCTGTGTGCTGGCCTGACTGTTTACAAAGGTCTGAAGATGACTGAAGCTCGTC
CAGGTCAGTGGGTTGCAGTTAGCGGTGTAGGCGGCCTGGGTCAAATGGCAGTGCAGTATGCG
GTGGCGATGGGCATGAACGTTGTTGCCGTAGATATCGATGACGAAAAACTGGCCACCGCTAA
AAAGCTGGGCGCTTCCCTGACTGTGAACGCAAAAGATACCGATCCGGCTCGTTTTATCCAGC
AACAGATCGGTGGTGCACATGGTGCTCTGGTTACCGCAGTGGGTCGTACTGCCTTCTCCCAA
GCAATGGGTTATGCCCGTCGTGGTGGTACCATCGTTCTGAACGGTCTGCCGCCGGGTGACTT
TCCGGTGTCCATTTTTGATATGGTGATGAACGGCACGACCATCCGTGGTAGCATCGTGGGCA
CCCGTCTGGATATGATCGAAGCTATGGATTTCTTCGCGCGCGGCAAAGTCAAATCCGTTGTT
ACCCCGGGCAAACTGGAAAACATCAACACGATCTTCGACGACCTGCAGAACGGTCGTCTGGA
AGGTCGTACCGTTCTGGATTTTCGTAGCTGAGAGACTCCCGGGACCTCAGAGTCCGCCACAC
CCGAACATAAGCTTGGCTGCGCAAAAAACCCCGCTTCGGCGGGGTTTTTTCGCGATGCATAC

TTGCGATCGTTGACAGTAAAAATTGCCCGTTTGTGAACCACTTGTTTGCAAACGGGCATGAC
TCCTGACTTTTATTTCTGCCTTTTATTCCTTTTACACTTGTTTTTATGAAGCCCTTCACAGA
ATTGTCCTTTCACGATTCCGTCTCTCTGATGATTGATGTTAATTAACAATGTATTCACCGAA
AACAAACATATAAATCACAGGAGTCGCCCATGTCAGTACCCGTTCAACATCCTATGTATATC
GATGGACAGTTTGTTACCTGGCGTGGAGACGCATGGATTGATGTGGTAAACCCTGCTACAGA
GGCTGTCATTTCCCGCATACCCGATGGTCAGGCCGAGGATGCCCGTAAGGCAATCGATGCAG
CAGAACGTGCACAACCAGAATGGGAAGCGTTGCCTGCTATTGAACGCGCCAGTTGGTTGCGC
AAAATCTCCGCCGGGATCCGCGAACGCGCCAGTGAAATCAGTGCGCTGATTGTTGAAGAAGG
GGGCAAGATCCAGCAGCTGGCTGAAGTCGAAGTGGCTTTTACTGCCGACTATATCGATTACA
TGGCGGAGTGGGCACGGCGTTACGAGGGCGAGATTATTCAAAGCGATCGTCCAGGAGAAAAT
ATTCTTTTGTTTAAACGTGCGCTTGGTGTGACTACCGGCATTCTGCCGTGGAACTTCCCGTT
CTTCCTCATTGCCCGCAAAATGGCTCCCGCTCTTTTGACCGGTAATACCATCGTCATTAAAC
CTAGTGAATTTACGCCAAACAATGCGATTGCATTCGCCAAAATCGTCGATGAAATAGGCCTT
CCGCGCGGCGTGTTTAACCTTGTACTGGGGCGTGGTGAAACCGTTGGGCAAGAACTGGCGGG
TAACCCAAAGGTCGCAATGGTCAGTATGACAGGCAGCGTCTCTGCAGGTGAGAAGATCATGG
CGACTGCGGCGAAAAACATCACCAAAGTGTGTCTGGAATTGGGGGGTAAAGCACCAGCTATC
GTAATGGACGATGCCGATCTTGAACTGGCAGTCAAAGCCATCGTTGATTCACGCGTCATTAA
TAGTGGGCAAGTGTGTAACTGTGCAGAACGTGTTTATGTACAGAAAGGCATTTATGATCAGT
TCGTCAATCGGCTGGGTGAAGCGATGCAGGCGGTTCAATTTGGTAACCCCGCTGAACGCAAC
GACATTGCGATGGGGCCGTTGATTAACGCCGCGGCGCTGGAAAGGGTCGAGCAAAAAGTGGC
GCGCGCAGTAGAAGAAGGGGCGAGAGTGGCGTTCGGTGGCAAAGCGGTAGAGGGGAAAGGAT
ATTATTATCCGCCGACATTGCTGCTGGATGTTCGCCAGGAAATGTCGATTATGCATGAGGAA
ACCTTTGGCCCGGTGCTGCCAGTTGTCGCATTTGACACGCTGGAAGATGCTATCTCAATGGC
TAATGACAGTGATTACGGCCTGACCTCATCAATCTATACCCAAAATCTGAACGTCGCGATGA
AAGCCATTAAAGGGCTGAAGTTTGGTGAAACTTACATCAACCGTGAAAACTTCGAAGCTATG
CAAGGCTTCCACGCCGGATGGCGTAAATCCGGTATTGGCGGCGCAGATGGTAAACATGGCTT
GCATGAATATCTGCAGACCCAGGTGGTTTATTTACAGTCTTAATGAGTGAAAGAGGCGGAGG
TTTTTTCCTCCGCCTGTGCGCAATGGAAACAGACCAGTTATTTTTCTGCGCCTCTTCCTGAC
CTGCGGCAATAATTGCACTCGCCATTTGCTGGCTAAGAATGACTTTAGTTTTCATTTTGTTA
TTCCTTTTCAAGGGCTTGTTCTACAATTTCAATCCAGTGACGCACAGAGGTACGACCGGCGC
TCGCCAGATGCGTCTGGCAACCAATGTTGGCGGTGACGATCATTTCCGGTTTGCCGCTTTCC
AGCGCATTCATTTTGTTATCCCGCAGCTGGCGTGCCAGATCGGGATGCGTTAACGCATATGT
TCCCGCTGAACCGCAGCACAGATGGCTGTCGGGAACGTCCGTTAAGGTAAATCCAAGACGAA
GCAACACTTTTTCCACTTCGCCGTTCAGCTTTTGCGCATGTTGTAGGGTACACGGACAGTGG
AAGGCCAGCTTTTTATCGCCGCGAATTGCCAGTTTTTCCAGCGGTTCCTCGCGCAGAAGTTC
GACTAAATCGACCGCCAGTTCACTGACCTGACGTGCTTTATCGGCATATAACGCATCGTTTT
TCAGCATCTGCCCATACTCTTTGACAAACGCGCCGCAGCCGCTGGCGGTTTGCAAAATTGCC
TCGGCACCTGCTTCAATCGCGGGCCACCAGGCATCAATATTATTGCGCGCCCGTGCCAGCCC
TTTCTCCTGCGCATTAAGATGATAGTCCACCGCGCCACAACAGCCTGCTTCGTTAGCTGGCA
TGACGCTGATCCCCAGACGATCCAGCACTCGCGCAGTTGCCGCGTTGGTGTTGGGCGAAAGC
GTAGGCTGGGCGCAGCCTTCCAACATTAAAACCCGACGCTTATGGCGCAGCGGCGGACGCGG
TTTAGCTTTCACCGTTTCAGCAGGCAGTTTTGCTCTGACCTGTTCCGGTAAAAACGGTCGCA
GCACCAGCCCTACCTGCGTCAGCGCACGGAAGACCGCCGGACGCGGCACTACCTGGCGCAAT
CCTTCGCGCAGTATTCGCTCCGGCAGTGGGCGTTTCACTTTCTGCTCGACAATATCACGCCC
GATATCCAGCAAATTGTGATAGCGCACACCAGAAGGACAGGTGGTTTCACAATTACGGCAAG
TGAGGCAGCGATCGAGATGCTCCTGTGTTTTAAGCGTGACTTCGTTGCCTTCCAGCACCTGT
TTAATCAGATAGATGCGCCCGCGCGGCCCGTCCAGTTCATCGCCCAGAAGCTGATAGGTTGG
GCAGGTTGCGGTACAAAATCCGCAGTGAACACAGGCGCGCAGGATGCTGTCGGCTTCCAGCG
CGCGCGCGTTCTGCCGCATCTCTTCAGTTAATTGGGTTTGCATAGCCTGCTCCTCAAAGTTC
CGCGTACATGCGACCGGGGTTAAACACGCCGCAAGGGTCGAGCTGCTGTTTAAGCTGCTGGT
GATAGCGGAATAAAGGAGCCGATAGCGGGGCAAAGCCACCATCTCCGGCACTAAAGCGGGTC

| | |
|---|---|
| 994 | GCATGACCGCCAGCGTTGCGGGCGATGCGATGGATTTGATTGTCCTCGGCTGTCGATTTCAG |
| 995 | CCAGCGTAACGCCCCGCCCCAGTCGATCAGTTGCTCGCCGGGTAAATCCATCATCGGCGCAT |
| 996 | CACTGGGTAATGAAATGCGCCATAAGGTACCTGGTAACGAGAAGAACGGCAGTTGTTGTTCA |
| 997 | CGCAATTGCTGCCAGAACTGACCGGCAACCTCTTCGCCACCCAGCAGTTCACGCGCTGCTTT |
| 998 | TACCGATCCTTCGCCGCCCTCAAGGCGGATCCACAACGCATTGTCGAAGTAACATAAGCCAC |
| 999 | TAATGGGTAATGGCTGGAGTTGCCACTCGGCGATTTCACTCATGGCTTCTTGCAGGCTGATT |
| 1000 | TCCCGACGCAGGCTCAGGGAGGCGCGCGGTCGCGGTAACACTTTCATTGAGATTTCAGTGAG |
| 1001 | CACGCCAAGACAACCGTAGCTTCCGACCATTAACCGTGAGAGATCGTATCCGGCAACGTTTT |
| 1002 | TCATCACTTCGCCACCAAAACGCAGATGTTTTCCAGCGCCGGTAATGATGCGCGTGCCGAGG |
| 1003 | ACAAAATCGCGGACCGAACCGCTCCACGGGCGACGCGGCCCCGCCAGCCCGCAGGCGACCAT |
| 1004 | CCCGCCCCAGGTGGCTTCTTCACCATAATGCGGCGGCTCACAGGGGAGCATTTGCCCCGCGC |
| 1005 | TTTCCAGCGCCGCTTCAATTGTCACCAGCGGCGTTCCGACACGCGCGGTTATCACCAGCTCG |
| 1006 | GTCGGGTCGTAATTAACAATGCCGCGATGACAACGAACATCCAGCGTTTGCCCGGTGACAGG |
| 1007 | GCGACCTAAAAAGGCTTTGCTATTGCTGCCCTGAATCACCAGCGGCGTTTTATCGCTAATCG |
| 1008 | CCTGATTCACCTGCTCCAGCAGCGCCTGGCTGTAATCACACTCGCGTAGCATCAGAAACGCT |
| 1009 | CCAGTTCAGGGAAAGGTAAATGACCGTGATGCACATGCATGGCACCAAATTCAGCACAGCGG |
| 1010 | TGTAGCGTGGGAATGTTTTTCCCAGGGTTCAGCAAACCATCGGGGTCAAACGCCGCCTTGAC |
| 1011 | CGCATGGAAGGTCGTGATTTCATCGCTGTTGAACTGGGCGCACATTTGATTGATTTTTTCTC |
| 1012 | GCCCGATGCCATGTTCGCCACTGATGCTGCCGCCAACTTCAACGCAGAGTTCGAGGATCTTC |
| 1013 | CCGCCCAGCTCTTCCGCGCGGGCAAATTCACCGGGTTCGTTGGCATCGAAAAGGATTAACGG |
| 1014 | GTGCATGTTGCCATCTCCGGCATGAAAGACGTTGGCAACACGTAAATCATATTGCTGCGATA |
| 1015 | AACGGGCAATGCCTTCCAGTACGCCAGGCAGGGCGCGACGCGGGATGGTGCCATCCATGCAG |
| 1016 | TAGTAATCCGGGGAGATACGTCCTACCGCCGGGAACGCATTTTTGCGACCGGCCCAGAAACG |
| 1017 | TACGCGCTCTGCTTCGTCCTGTGCCAGACGGACGTCAGTCGCGCCCGCTTTCAACAAGATGT |
| 1018 | CGTTAACCCGCTCGCAGTCTTCCTGTACGTCAGACTCCACGCCGTCCAGCTCGCATAACAAA |
| 1019 | ATCGCTTCGGCGTCGACGGGATAACCGGCATGAATAAAATCTTCCGCCCGCGCGGCCCGTCC |
| 1020 | AGTTCATCGCCCAGAAGCTGATAGGTTGGGCAGGTTGCGGTACAAAATCCGCAGTGAACACA |
| 1021 | GGCGCGCAGGATGCTGTCGGCTTCCAGCGCGCGCGCGTTCTGCCGCATCTCTTCAGTTAATT |
| 1022 | GGGTTTGCATAGCCTGCTCCTCAAAGTTCCGCGTACATGCGACCGGGGTTAAACACGCCGCA |
| 1023 | AGGGTCGAGCTGCTGTTTAAGCTGCTGGTGATAGCGGAATAAAGGAGCCGATAGCGGGGCAA |
| 1024 | AGCCACCATCTCCGGCACTAAAGCGGGTCGCATGACCGCCAGCGTTGCGGGCGATGCGATGG |
| 1025 | ATTTGATTGTCCTCGGCTGTCGATTTCAGCCAGCGTAACGCCCCGCCCCAGTCGATCAGTTG |
| 1026 | CTCGCCGGGTAAATCCATCATCGGCGCATCACTGGGTAATGAAATGCGCCATAAGGTACCTG |
| 1027 | GTAACGAGAAGAACGGCAGTTGTTGTTCACGCAATTGCTGCCAGAACTGACCGGCAACCTCT |
| 1028 | TCGCCACCCAGCAGTTCACGCGCTGCTTTTACCGATCCTTCGCCGCCCTCAAGGCGGATCCA |
| 1029 | CAACGCATTGTCGAAGTAACATAAGCCACTAATGGGTAATGGCTGGAGTTGCCACTCGGCGA |
| 1030 | TTTCACTCATGGCTTCTTGCAGGCTGATTTCCCGACGCAGGCTCAGGGAGGCGCGCGGTCGC |
| 1031 | GGTAACACTTTCATTGAGATTTCAGTGAGCACGCCAAGACAACCGTAGCTTCCGACCATTAA |
| 1032 | CCGTGAGAGATCGTATCCGGCAACGTTTTTCATCACTTCGCCACCAAAACGCAGATGTTTTC |
| 1033 | CAGCGCCGGTAATGATGCGCGTGCCGAGGACAAAATCGCGGACCGAACCGCTCCACGGGCGA |
| 1034 | CGCGGCCCCGCCAGCCCGCAGGCGACCATCCCGCCCCAGGTGGCTTCTTCACCATAATGCGG |
| 1035 | CGGCTCACAGGGGAGCATTTGCCCCGCGCTTTCCAGCGCCGCTTCAATTGTCACCAGCGGCG |
| 1036 | TTCCGACACGCGCGGTTATCACCAGCTCGGTCGGGTCGTAATTAACAATGCCGCGATGACAA |
| 1037 | CGAACATCCAGCGTTTGCCCGGTGACAGGGCGACCTAAAAAGGCTTTGCTATTGCTGCCCTG |
| 1038 | AATCACCAGCGGCGTTTTATCGCTAATCGCCTGATTCACCTGCTCCAGCAGCGCCTGGCTGT |
| 1039 | AATCACACTCGCGTAGCATCAGAAACGCTCCAGTTCAGGGAAAGGTAAATGACCGTGATGCA |
| 1040 | CATGCATGGCACCAAATTCAGCACAGCGGTGTAGCGTGGGAATGTTTTTCCCAGGGTTCAGC |
| 1041 | AAACCATCGGGGTCAAACGCCGCCTTGACCGCATGGAAGGTCGTGATTTCATCGCTGTTGAA |
| 1042 | CTGGGCGCACATTTGATTGATTTTTTCTCGCCCGATGCCATGTTCGCCACTGATGCTGCCGC |
| 1043 | CAACTTCAACGCAGAGTTCGAGGATCTTCCCGCCCAGCTCTTCCGCGCGGGCAAATTCACCG |
| 1044 | GGTTCGTTGGCATCGAAAAGGATTAACGGGTGCATGTTGCCATCTCCGGCATGAAAGACGTT |

GGCAACACGTAAATCATATTGCTGCGATAAACGGGCAATGCCTTCCAGTACGCCAGGCAGGG
CGCGACGCGGGATGGTGCCATCCATGCAGTAGTAATCCGGGGAGATACGTCCTACCGCCGGG
AACGCATTTTTGCGACCGGCCCAGAAACGTACGCGCTCTGCTTCGTCCTGTGCCAGACGGAC
GTCAGTCGCGCCCGCTTTCAACAAGATGTCGTTAACCCGCTCGCAGTCTTCCTGTACGTCAG
ACTCCACGCCGTCCAGCTCGCATAACAAAATCGCTTCGGCGTCGACGGGATAACCGGCATGA
ATAAAATCTTCCGCCGCGCGGATCGACAGGTTATCCATCATCTCCAGCCCGCCGGGGATAAT
GCCATTGGCGATGATGTCACCAACCGCAAGTCCGGCTTTTTCTACCGAGTCAAAGCTGGCTA
ACAGAACCCGCGCCACGGGCGGCTTCGGCAGCAGTTTTACCGTCACTTCGGTGGTCACGCCG
AGCATACCTTCCGATCCGGTGAACAGCGCCAGCAGGTCAAAACCAGGTGAATCCAGCGCGTC
CGATCCAAGCGTCAGTGCCTCGCCGTCCAGCGTTTGCACTTCAATTTTCAGCAGGTTATGTA
CGGTCAGACCATATTTCAGGCAGTGGACGCCGCCGGCATTTTCAGCCACATTGCCGCCAATG
GAACAGGCGATTTGTGAGGAAGGGTCCGGTGCGTAGTAGAGATTATGCGGTGCAACGGCCTG
GGAGATCGCCAGGTTACGCACGCCTGGCTGCACGCGCGCGCGGCGACCAACGGGGTTAATGT
CGAGGATCTCTTTAAAGCGCGCCATCACCAACAACACACCTTTTTCCAGCGGCAGCGCGCCA
CCAGAAAGCCCGGTGCCTGCACCACGGGTCACCACCGGTACACGCAGGCGATGGCAGACAGC
CAGAATCGCTGTCACCTGTTCCATTTGCTTAGGCAGAACAACCAGTAATGGACGCGTGCGAT
ACGCGCTCAACCCGTCACACTCGTAAGGAATGATCTCCTCATCGGTATGCAGGATCTCAAGT
CCAGGGACATGCTCACGCAGTGCCATCAGTACCGATGTGCGGTCGACATCGGGTAAAGCGCC
ATCAAGACGCTCTTCGTACAAGATGCTCATGAGTAGGCTTCGCTTTGTTGTGTTGTGTGGCA
GCTGATTTTTGCGCGCTGCTTCTGTGAACAGTTATTAAGCGGGCTTTTCGTTTTCGTCTATC
TCTTTAGCTACCGGTCAGACCATTTTTTTTCCAGCTCTGTGACCTTGTCTTGGTTAACTCAA
TGTTAAATTGATGTAACATAATCACTTACGTGATGTGCGTGTTTTGCGAGTTAAGAACAGAA
AAATTGGTCCTACCTGTGCACGAGGTCCGGGATCTGACGCTCAGTGGAACGAAAACTCACGT
TAAGGATTTTGGTCATGAACAATAAAACTGTCTGCTTACATAAACAGTAATACAAGGGGTG
TTATGAGCCATATTCAACGG

EGA7_pAN29_7

CATCCTATGGAACTGCCTCGGTGAGTTTTCTCCTTCATTACAGAAACGGCTTTTTCAAAAAT
ATGGTATTGATAATCCTGATATGAATAAATTGCAGTTTCATTTGATGCTCGATGAGTTTTTC
TAAAAGCTTAATTAGCTGATCTAGACGCGTGCTAGAGGCATCAAATAAAACGAAAGGCTCAG
TCGAAAGACTGGGCCTTTCGTTTTATCTGTTGTTTGTCGGTGAACGCTCTCCTGAGTAGGAC
AAATAGGTCGAGGGTGAAGTACTTGCTGACTTCCTTGAGGAACACATGATGCGTCCTACGGT
TGCTGCTACGCATATCATTGAGATGTCTGTGGGAGGAGTTGATGTGTACTCTGAGGACGATG
AGGGTTACGGTACGTCTTTCATTGAGTGGTGATTTATGCATTAGGACTGCATAGGGATGCAC
TATAGACCACGGATGGTCAGTTCTTTAAGTTACTGAAAAGACACGATAAATTAATACGACTC
ACTATAGGGAGAGGAGGGACGAAAGGTTACTATATAGATACTGAATGAATACTTATAGAGTG
CATAAAGTATGCATAATGGTGTACCTAGAGTGACCTCTAAGAATGGTGATTATATTGTATTA
GTATCACCTTAACTTAAGGCGGGATCGTCACCCTCAGCAGCGAAAGACAGCTGTCGGTCAGA
GCGTCATTGCGAAGCTGAGTGTGATCGATGCCATCAGCGAAGGGCCCAAACTCCGAGCGATT
AAGCGTTTGCTGGCTGTCACGCCTGCCTGTTGCTTGCTTGGACTTGCGATGTACGTGCTCAG
CTGTCTTTCGCTGCTGAGGGTGACGATCCCGCGAGGGCCTATGGAGTTCCTATAGGGTCCTT
TAAAATATACCATAAAAATCTGAGTGACTATCTCACAGTGTACGGACCTAAAGTTCCCCCAT
AGGGGGTACCTAAAGCCCAGCCAATCACCTAAAGTCAACCTTCGGTTGACCTTGAGGGTTCC
CTAAGGGTTGGGGATGACCCTTGGGTTTGTCTTTGGGTGTTACCTTGAGTGTCTCTCTGTGT
CCCTATCTGTTACAGTCTCCTAAAGTATCCTCCTAAAGTCACCTCCTAACGCACATTTCCCC
GAAAAGTGCCACCTGGGTCCTTTTCACTTCGGGCTCATGAGCAAATATTTTATCTGATTAAT
AAGATGATCTTCTTGAGATCGTTTTGGTCTGCGCGTAATCTCTTGCTCTGAAAACGAAAAAA
CCGCCTTGCAGGGCGGTTTTTCGAAGGTTCTCTGAGCTACCAACTCTTTGAACCGAGGTAAC
TGGCTTGGAGGAGCGCAGTCACCAAAACTTGTCCTTTCAGTTTAGCCTTAACCGGCGCATGA
CTTCAAGACTAACTCCTCTAAATCAATTACCAGTGGCTGCTGCCAGTGGTGCTTTTGCATGT
CTTTCCGGGTTGGACTCAAGACGATAGTTACCGGATAAGGCGCAGCGGTCGGACTGAACGGG
GGGTTCGTGCATACAGTCCAGCTTGGAGCGAACTGCCTACCCGGAACTGAGTGTCAGGCGTG
GAATGAGACAAACGCGGCCATAACAGCGGAATGACACCGGTAAACCGAAAGGCAGGAACAGG
AGAGCGCACGAGGGAGCCGCCAGGGGGAAACGCCTGGTATCTTTATAGTCCTGTCGGGTTTC
GCCACCACTGATTTGAGCGTCAGATTTCGTGATGCTTGTCAGGGGGGCGGAGCCTATGGAAA
AACGGCTTTGCCGCGGCCCTCTCACTTCCCTGTTAAGTATCTTCCTGGCATCTTCCAGGAAA
TCTCCGCCCCGTTCGTAAGCCATTTCCGCTCGCCGCAGTCGAACGACCGAGCGTAGCGAGTC
AGTGAGCGAGGAAGCGGAATATATCCTTGACAGCTAGCTCAGTCCTAGGTACTGTGCTAGCA
CCCGTTTTTACTAGTCTAGAAATAATAACAAAATGAGGAGGTACTGATATACATATGGCGGA
CACGATGCTGGCGGCGGTTGTACGTGAATTTGGTAAACCTCTGTCTATCGAACGCCTGCCGA
TCCCAGACATTAAACCGCACCAGATCCTGGTTAAAGTTGATACCTGCGGCGTTTGTCACACT
GACCTGCATGCAGCTCGTGGTGACTGGCCTTCCAAGCCGAACCCGCCGTTTATCCCGGGTCA
CGAAGGCGTTGGCCACATCGTAGCTGTTGGCAGCCAGGTGGGCGACTTTGTTAAGACTGGTG
ATGTTGTAGGTGTGCCTTGGCTGTACTCTGCATGTGGTCATTGTGAGCACTGTCTGGGTGGC
TGGGAAACTCTGTGCGAGAAGCAGGACGACACTGGTTATACCGTGAACGGTTGTTTTGCAGA
ATACGTAGTTGCCGATCCGAATTACGTTGCACACCTGCCGAGCACCATTGATCCACTGCAGG
CATCCCCGGTGCTGTGTGCTGGCCTGACTGTTTACAAAGGTCTGAAGATGACTGAAGCTCGT
CCAGGTCAGTGGGTTGCAGTTAGCGGTGTAGGCGGCCTGGGTCAAATGGCAGTGCAGTATGC
GGTGGCGATGGGCATGAACGTTGTTGCCGTAGATATCGATGACGAAAAACTGGCCACCGCTA
AAAAGCTGGGCGCTTCCCTGACTGTGAACGCAAAAGATACCGATCCGGCTCGTTTTATCCAG
CAACAGATCGGTGGTGCACATGGTGCTCTGGTTACCGCAGTGGGTCGTACTGCCTTCTCCCA
AGCAATGGGTTATGCCCGTCGTGGTGGTACCATCGTTCTGAACGGTCTGCCGCCGGGTGACT
TTCCGGTGTCCATTTTTGATATGGTGATGAACGGCACGACCATCCGTGGTAGCATCGTGGGC
ACCCGTCTGGATATGATCGAAGCTATGGATTTCTTCGCGCGCGGCAAAGTCAAATCCGTTGT
TACCCCGGGCAAACTGGAAAACATCAACACGATCTTCGACGACCTGCAGAACGGTCGTCTGG
AAGGTCGTACCGTTCTGGATTTTCGTAGCTGAGAGACTCCCGGGACCTCAGAGTCCGCCACA
CCCGAACATAAGCTTGGCTGCGCAAAAAACCCCGCTTCGGCGGGGTTTTTTCGCGATGCATA

1122 CTTGCGATCGTTGACAGTAAAAATTGCCCGTTTGTGAACCACTTGTTTGCAAACGGGCATGA
1123 CTCCTGACTTTTATTTCTGCCTTTTATTCCTTTTACACTTGTTTTTATGAAGCCCTTCACAG
1124 AATTGTCCTTTCACGATTCCGTCTCTCTGATGATTGATGTTAATTAACAATGTATTCACCGA
1125 AAACAAACATATAAATCACAGGAGTCGCCCATGTCAGTACCCGTTCAACATCCTATGTATAT
1126 CGATGGACAGTTTGTTACCTGGCGTGGAGACGCATGGATTGATGTGGTAAACCCTGCTACAG
1127 AGGCTGTCATTTCCCGCATACCCGATGGTCAGGCCGAGGATGCCCGTAAGGCAATCGATGCA
1128 GCAGAACGTGCACAACCAGAATGGGAAGCGTTGCCTGCTATTGAACGCGCCAGTTGGTTGCG
1129 CAAAATCTCCGCCGGGATCCGCGAACGCGCCAGTGAAATCAGTGCGCTGATTGTTGAAGAAG
1130 GGGGCAAGATCCAGCAGCTGGCTGAAGTCGAAGTGGCTTTTACTGCCGACTATATCGATTAC
1131 ATGGCGGAGTGGGCACGGCGTTACGAGGGCGAGATTATTCAAAGCGATCGTCCAGGAGAAAA
1132 TATTCTTTTGTTTAAACGTGCGCTTGGTGTGACTACCGGCATTCTGCCGTGGAACTTCCCGT
1133 TCTTCCTCATTGCCCGCAAAATGGCTCCCGCTCTTTTGACCGGTAATACCATCGTCATTAAA
1134 CCTAGTGAATTTACGCCAAACAATGCGATTGCATTCGCCAAAATCGTCGATGAAATAGGCCT
1135 TCCGCGCGGCGTGTTTAACCTTGTACTGGGGCGTGGTGAAACCGTTGGGCAAGAACTGGCGG
1136 GTAACCCAAAGGTCGCAATGGTCAGTATGACAGGCAGCGTCTCTGCAGGTGAGAAGATCATG
1137 GCGACTGCGGCGAAAAACATCACCAAAGTGTGTCTGGAATTGGGGGGTAAAGCACCAGCTAT
1138 CGTAATGGACGATGCCGATCTTGAACTGGCAGTCAAAGCCATCGTTGATTCACGCGTCATTA
1139 ATAGTGGGCAAGTGTGTAACTGTGCAGAACGTGTTTATGTACAGAAAGGCATTTATGATCAG
1140 TTCGTCAATCGGCTGGGTGAAGCGATGCAGGCGGTTCAATTTGGTAACCCCGCTGAACGCAA
1141 CGACATTGCGATGGGGCCGTTGATTAACGCCGCGGCGCTGGAAAGGGTCGAGCAAAAAGTGG
1142 CGCGCGCAGTAGAAGAAGGGGCGAGAGTGGCGTTCGGTGGCAAAGCGGTAGAGGGGAAAGGA
1143 TATTATTATCCGCCGACATTGCTGCTGGATGTTCGCCAGGAAATGTCGATTATGCATGAGGA
1144 AACCTTTGGCCCGGTGCTGCCAGTTGTCGCATTTGACACGCTGGAAGATGCTATCTCAATGG
1145 CTAATGACAGTGATTACGGCCTGACCTCATCAATCTATACCCAAAATCTGAACGTCGCGATG
1146 AAAGCCATTAAAGGGCTGAAGTTTGGTGAAACTTACATCAACCGTGAAAACTTCGAAGCTAT
1147 GCAAGGCTTCCACGCCGGATGGCGTAAATCCGGTATTGGCGGCGCAGATGGTAAACATGGCT
1148 TGCATGAATATCTGCAGACCCAGGTGGTTTATTTACAGTCTTAATGAGTGAAAGAGGCGGAG
1149 GTTTTTTCCTCCGCCTGTGCGCAATGGAAACAGACCAGTTATTTTTCTGCGCCTCTTCCTGA
1150 CCTGCGGCAATAATTGCACTCGCCATTTGCTGGCTAAGAATGACTTTAGTTTTCATTTTGTT
1151 ATTCCTTTTCAAGGGCTTGTTCTACAATTTCAATCCAGTGACGCACAGAGGTACGACCGGCG
1152 CTCGCCAGATGCGTCTGGCAACCAATGTTGGCGGTGACGATCATTTCCGGTTTGCCGCTTTC
1153 CAGCGCATTCATTTTGTTATCCCGCAGCTGGCGTGCCAGATCGGGATGCGTTAACGCATATG
1154 TTCCCGCTGAACCGCAGCACAGATGGCTGTCGGGAACGTCCGTTAAGGTAAATCCAAGACGA
1155 AGCAACACTTTTTCCACTTCGCCGTTCAGCTTTTGCGCATGTTGTAGGGTACACGGACAGTG
1156 GAAGGCCAGCTTTTTATCGCCGCGAATTGCCAGTTTTTCCAGCGGTTCCTCGCGCAGAAGTT
1157 CGACTAAATCGACCGCCAGTTCACTGACCTGACGTGCTTTATCGGCATATAACGCATCGTTT
1158 TTCAGCATCTGCCCATACTCTTTGACAAACGCGCCGCAGCCGCTGGCGGTTTGCAAAATTGC
1159 CTCGGCACCTGCTTCAATCGCGGGCCACCAGGCATCAATATTATTGCGCGCCCGTGCCAGCC
1160 CTTTCTCCTGCGCATTAAGATGATAGTCCACCGCGCCACAACAGCCTGCTTCGTTAGCTGGC
1161 ATGACGCTGATCCCCAGACGATCCAGCACTCGCGCAGTTGCCGCGTTGGTGTTGGGCGAAAG
1162 CGTAGGCTGGGCGCAGCCTTCCAACATTAAAACCCGACGCTTATGGCGCAGCGGCGGACGCG
1163 GTTTAGCTTTCACCGTTTCAGCAGGCAGTTTTGCTCTGACCTGTTCCGGTAAAAACGGTCGC
1164 AGCACCAGCCCTACCTGCGTCAGCGCACGGAAGACCGCCGGACGCGGCACTACCTGGCGCAA
1165 TCCTTCGCGCAGTATTCGCTCCGGCAGTGGGCGTTTCACTTTCTGCTCGACAATATCACGCC
1166 CGATATCCAGCAAATTGTGATAGCGCACACCAGAAGGACAGGTGGTTTCACAATTACGGCAA
1167 GTGAGGCAGCGATCGAGATGCTCCTGTGTTTTAAGCGTGACTTCGTTGCCTTCCAGCACCTG
1168 TTTAATCAGATAGATGCGCCCGCGCGGCCCGTCCAGTTCATCGCCCAGAAGCTGATAGGTTG
1169 GGCAGGTTGCGGTACAAAATCCGCAGTGAACACAGGCGCGCAGGATGCTGTCGGCTTCCAGC
1170 GCGCGCGCGTTCTGCCGCATCTCTTCAGTTAATTGGGTTTGCATAGCCTGCTCCTCAAAGTT
1171 CCGCGTACATGCGACCGGGGTTAAACACGCCGCAAGGGTCGAGCTGCTGTTTAAGCTGCTGG
1172 TGATAGCGGAATAAAGGAGCCGATAGCGGGGCAAAGCCACCATCTCCGGCACTAAAGCGGGT

CGCATGACCGCCAGCGTTGCGGGCGATGCGATGGATTTGATTGTCCTCGGCTGTCGATTTCA
GCCAGCGTAACGCCCCGCCCCAGTCGATCAGTTGCTCGCCGGGTAAATCCATCATCGGCGCA
TCACTGGGTAATGAAATGCGCCATAAGGTACCTGGTAACGAGAAGAACGGCAGTTGTTGTTC
ACGCAATTGCTGCCAGAACTGACCGGCAACCTCTTCGCCACCCAGCAGTTCACGCGCTGCTT
TTACCGATCCTTCGCCGCCCTCAAGGCGGATCCACAACGCATTGTCGAAGTAACATAAGCCA
CTAATGGGTAATGGCTGGAGTTGCCACTCGGCGATTTCACTCATGGCTTCTTGCAGGCTGAT
TTCCCGACGCAGGCTCAGGGAGGCGCGCGGTCGCGGTAACACTTTCATTGAGATTTCAGTGA
GCACGCCAAGACAACCGTAGCTTCCGACCATTAACCGTGAGAGATCGTATCCGGCAACGTTT
TTCATCACTTCGCCACCAAAACGCAGATGTTTTCCAGCGCCGGTAATGATGCGCGTGCCGAG
GACAAAATCGCGGACCGAACCGCTCCACGGGCGACGCGGCCCCGCCAGCCCGCAGGCGACCA
TCCCGCCCCAGGTGGCTTCTTCACCATAATGCGGCGGCTCACAGGGGAGCATTTGCCCCGCG
CTTTCCAGCGCCGCTTCAATTGTCACCAGCGGCGTTCCGACACGCGCGGTTATCACCAGCTC
GGTCGGGTCGTAATTAACAATGCCGCGATGACAACGAACATCCAGCGTTTGCCCGGTGACAG
GGCGACCTAAAAAGGCTTTGCTATTGCTGCCCTGAATCACCAGCGGCGTTTTATCGCTAATC
GCCTGATTCACCTGCTCCAGCAGCGCCTGGCTGTAATCACACTCGCGTAGCATCAGAAACGC
TCCAGTTCAGGGAAAGGTAAATGACCGTGATGCACATGCATGGCACCAAATTCAGCACAGCG
GTGTAGCGTGGGAATGTTTTTCCCAGGGTTCAGCAAACCATCGGGGTCAAACGCCGCCTTGA
CCGCATGGAAGGTCGTGATTTCATCGCTGTTGAACTGGGCGCACATTTGATTGATTTTTTCT
CGCCCGATGCCATGTTCGCCACTGATGCTGCCGCCAACTTCAACGCAGAGTTCGAGGATCTT
CCCGCCCAGCTCTTCCGCGCGGGCAAATTCACCGGGTTCGTTGGCATCGAAAAGGATTAACG
GGTGCATGTTGCCATCTCCGGCATGAAAGACGTTGGCAACACGTAAATCATATTGCTGCGAT
AAACGGGCAATGCCTTCCAGTACGCCAGGCAGGGCGCGACGCGGGATGGTGCCATCCATGCA
GTAGTAATCCGGGGAGATACGTCCTACCGCCGGGAACGCATTTTTGCGACCGGCCCAGAAAC
GTACGCGCTCTGCTTCGTCCTGTGCCAGACGGACGTCAGTCGCGCCCGCTTTCAACAAGAAG
TCGTTAACCCGCTCGCAGTCTTCCTGTACGTCAGACTCCACGCCGTCCAGCTCGCATAACAA
AATCGCTTCGGCGTCGACGGGATAACCGGCATGAATAAAATCTTCCGCCGCGCGGATCGACA
GGTTATCCATCATCTCCAGCCCGCCGGGGATAATGCCATTGGCGATGATGTCACCAACCGCA
AGTCCGGCTTTTTCTACCGAGTCAAAGCTGGCTAACAGAACCCGCGCCACGGGCGGCTTCGG
CAGCAGTTTTACCGTCACTTCGGTGGTCACGCCGAGCATACCTTCCGATCCGGTGAACAGCG
CCAGCAGGTCAAAACCAGGTGAATCCAGCGCGTCCGATCCAAGCGTCAGTGCCTCGCCGTCC
AGCGTTTGCACTTCAATTTTCAGCAGGTTATGTACGGTCAGACCATATTTCAGGCAGTGGAC
GCCGCCGGCATTTTCAGCCACATTGCCGCCAATGGAACAGGCGATTTGTGAGGAAGGGTCCG
GTGCGTAGTAGAGATTATGCGGTGCAACGGCCTGGGAGATCGCCAGGTTACGCACGCCTGGC
TGCACGCGCGCGCGGCGACCAACGGGGTTAATGTCGAGGATCTCTTTAAAGCGCGCCATCAC
CAACAACACACCTTTTTCCAGCGGCAGCGCGCCACCAGAAAGCCCGGTGCCTGCACCACGGG
TCACCACCGGTACACGCAGGCGATGGCAGACAGCCAGAATCGCTGTCACCTGTTCCATTTGC
TTAGGCAGAACAACCAGTAATGGACGCGTGCGATACGCGCTCAACCCGTCACACTCGTAAGG
AATGATCTCCTCATCGGTATGCAGGATCTCAAGTCCAGGGACATGCTCACGCAGTGCCATCA
GTACCGATGTGCGGTCGACATCGGGTAAAGCGCCATCAAGACGCTCTTCGTACAAGATGCTC
ATGAGTAGGCTTCGCTTTGTTGTGTTGTGTGGCAGCTGATTTTTGCGCGCTGCTTCTGTGAA
CAGTTATTAAGCGGGCTTTTCGTTTTCGTCTATCTCTTTAGCTACCGGTCAGACCATTTTTT
TTCCAGCTCTGTGACCTTGTCTTGGTTAACTCAATGTTAAATTGATGTAACATAATCACTTA
CGTGATGTGCGTGTTTTGCGAGTTAAGAACAGAAAAATTGGTCCTACCTGTGCACGAGGTCC
GGGATCTGACGCTCAGTGGAACGAAAACTCACGTTAAGGGATTTTGGTCATGAACAATAAAA
CTGTCTGCTTACATAAACAGTAATACAAGGGGTGTTATGAGCCATATTCAACGGG

EGA8_pAN29_8
CCTATGGAACTGCCTCGGTGAGTTTTCTCCTTCATTACAGAAACGGCTTTTTCAAAAATATG
GTATTGATAATCCTGATATGAATAAATTGCAGTTTCATTTGATGCTCGATGAGTTTTTCTAA
AAGCTTAATTAGCTGATCTAGACGCGTGCTAGAGGCATCAAATAAAACGAAAGGCTCAGTCG
AAAGACTGGGCCTTTCGTTTTATCTGTTGTTTGTCGGTGAACGCTCTCCTGAGTAGGACAAA
TAGGTCGAGGGTGAAGTACTTGCTGACTTCCTTGAGGAACACATGATGCGTCCTACGGTTGC
TGCTACGCATATCATTGAGATGTCTGTGGGAGGAGTTGATGTGTACTCTGAGGACGATGAGG
GTTACGGTACGTCTTTCATTGAGTGGTGATTTATGCATTAGGACTGCATAGGGATGCACTAT
AGACCACGGATGGTCAGTTCTTTAAGTTACTGAAAAGACACGATAAATTAATACGACTCACT
ATAGGGAGAGGAGGGACGAAAGGTTACTATATAGATACTGAATGAATACTTATAGAGTGCAT
AAAGTATGCATAATGGTGTACCTAGAGTGACCTCTAAGAATGGTGATTATATTGTATTAGTA
TCACCTTAACTTAAGGCGGGATCGTCACCCTCAGCAGCGAAAGACAGCTGTCGGTCAGAGCG
TCATTGCGAAGCTGAGTGTGATCGATGCCATCAGCGAAGGGCCCAAACTCCGAGCGATTAAG
CGTTTGCTGGCTGTCACGCCTGCCTGTTGCTTGCTTGGACTTGCGATGTACGTGCTCAGCTG
TCTTTCGCTGCTGAGGGTGACGATCCCGCGAGGGCCTATGGAGTTCCTATAGGGTCCTTTAA
AATATACCATAAAAATCTGAGTGACTATCTAACAGTGTACGGACCTAAAGTTCCCCCATAGG
GGGTACCTAAAGCCCAGCCAATCACCTAAAGTCAACCTTCGGTTGACCTTGAGGGTTCCCTA
AGGGTTGGGGATGACCCTTGGGTTTGTCTTTGGGTGTTACCTTGAGTGTCTCTCTGTGTCCC
TATCTGTTACAGTCTCCTAAAGTATCCTCCTAAAGTCACCTCCTAACGCACATTTCCCCGAA
AAGTGCCACCTGGGTCCTTTTCACTTCGGGCTCATGAGCAAATATTTTATCTGATTAATAAG
ATGATCTTCTTGAGATCGTTTTGGTCTGCGCGTAATCTCTTGCTCTGAAAACGAAAAAACCG
CCTTGCAGGGCGGTTTTTCGAAGGTTCTCTGAGCTACCAACTCTTTGAACCGAGGTAACTGG
CTTGGAGGAGCGCAGTCACCAAAACTTGTCCTTTCAGTTTAGCCTTAACCGGCGCATGACTT
CAAGACTAACTCCTCTAAATCAATTACCAGTGGCTGCTGCCAGTGGTGCTTTTGCATGTCTT
TCCGGGTTGGACTCAAGACGATAGTTACCGGATAAGGCGCAGCGGTCGGACTGAACGGGGGG
TTCGTGCATACAGTCCAGCTTGGAGCGAACTGCCTACCCGGAACTGAGTGTCAGGCGTGGAA
TGAGACAAACGCGGCCATAACAGCGGAATGACACCGGTAAACCGAAAGGCAGGAACAGGAGA
GCGCACGAGGGAGCCGCCAGGGGGAAACGCCTGGTATCTTTATAGTCCTGTCGGGTTTCGCC
ACCACTGATTTGAGCGTCAGATTTCGTGATGCTTGTCAGGGGGGCGGAGCCTATGGAAAAAC
GGCTTTGCCGCGGCCCTCTCACTTCCCTGTTAAGTATCTTCCTGGCATCTTCCAGGAAATCT
CCGCCCCGTTCGTAAGCCATTTCCGCTCGCCGCAGTCGAACGACCGAGCGTAGCGAGTCAGT
GAGCGAGGAAGCGGAATATATCCTTGACAGCTAGCTCAGTCCTAGGTACTGTGCTAGCACCC
GTTTTTACTAGTCTAGAAATAATAACAAAATGAGGAGGTACTGATATACATATGGCGGACAC
GATGCTGGCGGCGGTTGTACGTGAATTTGGTAAACCTCTGTCTATCGAACGCCTGCCGATCC
CAGACATTAAACCGCACCAGATCCTGGTTAAAGTTGATACCTGCGGCGTTTGTCACACTGAC
CTGCATGCAGCTCGTGGTGACTGGCCTTCCAAGCCGAACCCGCCGTTTATCCCGGGTCACGA
AGGCGTTGGCCACATCGTAGCTGTTGGCAGCCAGGTGGGCGACTTTGTTAAGACTGGTGATG
TTGTAGGTGTGCCTTGGCTGTACTCTGCATGTGGTCATTGTGAGCACTGTCTGGGTGGCTGG
GAAACTCTGTGCGAGAAGCAGGACGACACTGGTTATACCGTGAACGGTTGTTTTGCAGAATA
CGTAGTTGCCGATCCGAATTACGTTGCACACCTGCCGAGCACCATTGATCCACTGCAGGCAT
CCCCGGTGCTGTGTGCTGGCCTGACTGTTTACAAAGGTCTGAAGATGACTGAAGCTCGTCCA
GGTCAGTGGGTTGCAGTTAGCGGTGTAGGCGGCCTGGGTCAAATGGCAGTGCAGTATGCGGT
GGCGATGGGCATGAACGTTGTTGCCGTAGATATCGATGACGAAAAACTGGCCACCGCTAAAA
AGCTGGGCGCTTCCCTGACTGTGAACGCAAAAGATACCGATCCGGCTCGTTTTATCCAGCAA
CAGATCGGTGGTGCACATGGTGCTCTGGTTACCGCAGTGGGTCGTACTGCCTTCTCCCAAGC
AATGGGTTATGCCCGTCGTGGTGGTACCATCGTTCTGAACGGTCTGCCGCCGGGTGACTTTC
CGGTGTCCATTTTTGATATGGTGATGAACGGCACGACCATCCGTGGTAGCATCGTGGGCACC
CGTCTGGATATGATCGAAGCTATGGATTTCTTCGCGCGCGGCAAAGTCAAATCCGTTGTTAC
CCCGGGCAAACTGGAAAACATCAACACGATCTTCGACGACCTGCAGAACGGTCGTCTGGAAG
GTCGTACCGTTCTGGATTTTCGTAGCTGAGAGACTCCCGGGACCTCAGAGTCCGCCACACCC
GAACATAAGCTTGGCTGCGCAAAAAACCCCGCTTCGGCGGGGTTTTTTCGCGATGCATACTT

1270 GCGATCGTTGACAGTAAAAATTGCCCGTTTGTGAACCACTTGTTTGCAAACGGGCATGACTC
1271 CTGACTTTTATTTCTGCCTTTTATTCCTTTTACACTTGTTTTTATGAAGCCCTTCACAGAAT
1272 TGTCCTTTCACGATTCCGTCTCTCTGATGATTGATGTTAATTAACAATGTATTCACCGAAAA
1273 CAAACATATAAATCACAGGAGTCGCCCATGTCAGTACCCGTTCAACATCCTATGTATATCGA
1274 TGGACAGTTTGTTACCTGGCGTGGAGACGCATGGATTGATGTGGTAAACCCTGCTACAGAGG
1275 CTGTCATTTCCCGCATACCCGATGGTCAGGCCGAGGATGCCCGTAAGGCAATCGATGCAGCA
1276 GAACGTGCACAACCAGAATGGGAAGCGTTGCCTGCTATTGAACGCGCCAGTTGGTTGCGCAA
1277 AATCTCCGCCGGGATCCGCGAACGCGCCAGTGAAATCAGTGCGCTGATTGTTGAAGAAGGGG
1278 GCAAGATCCAGCAGCTGGCTGAAGTCGAAGTGGCTTTTACTGCCGACTATATCGATTACATG
1279 GCGGAGTGGGCACGGCGTTACGAGGGCGAGATTATTCAAAGCGATCGTCCAGGAGAAAATAT
1280 TCTTTTGTTTAAACGTGCGCTTGGTGTGACTACCGGCATTCTGCCGTGGAACTTCCCGTTCT
1281 TCCTCATTGCCCGCAAAATGGCTCCCGCTCTTTTGACCGGTAATACCATCGTCATTAAACCT
1282 AGTGAATTTACGCCAAACAATGCGATTGCATTCGCCAAAATCGTCGATGAAATAGGCCTTCC
1283 GCGCGGCGTGTTTAACCTTGTACTGGGGCGTGGTGAAACCGTTGGGCAAGAACTGGCGGGTA
1284 ACCCAAAGGTCGCAATGGTCAGTATGACAGGCAGCGTCTCTGCAGGTGAGAAGATCATGGCG
1285 ACTGCGGCGAAAAACATCACCAAAGTGTGTCTGGAATTGGGGGGTAAAGCACCAGCTATCGT
1286 AATGGACGATGCCGATCTTGAACTGGCAGTCAAAGCCATCGTTGATTCACGCGTCATTAATA
1287 GTGGGCAAGTGTGTAACTGTGCAGAACGTGTTTATGTACAGAAAGGCATTTATGATCAGTTC
1288 GTCAATCGGCTGGGTGAAGCGATGCAGGCGGTTCAATTTGGTAACCCCGCTGAACGCAACGA
1289 CATTGCGATGGGGCCGTTGATTAACGCCGCGGCGCTGGAAAGGGTCGAGCAAAAAGTGGCGC
1290 GCGCAGTAGAAGAAGGGGCGAGAGTGGCGTTCGGTGGCAAAGCGGTAGAGGGGAAAGGATAT
1291 TATTATCCGCCGACATTGCTGCTGGATGTTCGCCAGGAAATGTCGATTATGCATGAGGAAAC
1292 CTTTGGCCCGGTGCTGCCAGTTGTCGCATTTGACACGCTGGAAGATGCTATCTCAATGGCTA
1293 ATGACAGTGATTACGGCCTGACCTCATCAATCTATACCCAAAATCTGAACGTCGCGATGAAA
1294 GCCATTAAAGGGCTGAAGTTTGGTGAAACTTACATCAACCGTGAAAACTTCGAAGCTATGCA
1295 AGGCTTCCACGCCGGATGGCGTAAATCCGGTATTGGCGGCGCAGATGGTAAACATGGCTTGC
1296 ATGAATATCTGCAGACCCAGGTGGTTTATTTACAGTCTTAATGAGTGAAAGAGGCGGAGGTT
1297 TTTTCCTCCGCCTGTGCGCAATGGAAACAGACCAGTTATTTTTCTGCGCCTCTTCCTGACCT
1298 GCGGCAATAATTGCACTCGCCATTTGCTGGCTAAGAATGACTTTAGTTTTCATTTTGTTATT
1299 CCTTTTCAAGGGCTTGTTCTACAATTTCAATCCAGTGACGCACAGAGGTACGACCGGCGCTC
1300 GCCAGATGCGTCTGGCAACCAATGTTGGCGGTGACGATCATTTCCGGTTTGCCGCTTTCCAG
1301 CGCATTCATTTTGTTATCCCGCAGCTGGCGTGCCAGATCGGGATGCGTTAACGCATATGTTC
1302 CCGCTGAACCGCAGCACAGATGGCTGTCGGGAACGTCCGTTAAGGTAAATCCAAGACGAAGC
1303 AACACTTTTTCCACTTCGCCGTTCAGCTTTTGCGCATGTTGTAGGGTACACGGACAGTGGAA
1304 GGCCAGCTTTTTATCGCCGCGAATTGCCAGTTTTTCCAGCGGTTCCTCGCGCAGAAGTTCGA
1305 CTAAATCGACCGCCAGTTCACTGACCTGACGTGCTTTATCGGCATATAACGAATCGTTTTTC
1306 AGCATCTGCCCATACTCTTTGACAAACGCGCCGCAGCCGCTGGCGGTTTGCAAAATTGCCTC
1307 GGCACCTGCTTCAATCGCGGGCCACCAGGCATCAATATTATTGCGCGCCCGTGCCAGCCCTT
1308 TCTCCTGCGCATTAAGATGATAGTCCACCGCGCCACAACAGCCTGCTTCGTTAGCTGGCATG
1309 ACGCTGATCCCCAGACGATCCAGCACTCGCGCAGTTGCCGCGTTGGTGTTGGGCGAAAGCGT
1310 AGGCTGGGCGCAGCCTTCCAACATTAAAACCCGACGCTTATGGCGCAGCGGCGGACGCGGTT
1311 TAGCTTTCACCGTTTCAGCAGGCAGTTTTGCTCTGACCTGTTCCGGTAAAAACGGTCGCAGC
1312 ACCAGCCCTACCTGCGTCAGCGCACGGAAGACCGCCGGACGCGGCACTACCTGGCGCAATCC
1313 TTCGCGCAGTATTCGCTCCGGCAGTGGGCGTTTCACTTTCTGCTCGACAATATCACGCCCGA
1314 TATCCAGCAAATTGTGATAGCGCACACCAGAAGGACAGGTGGTTTCACAATTACGGCAAGTG
1315 AGGCAGCGATCGAGATGCTCCTGTGTTTTAAGCGTGACTTCGTTGCCTTCCAGCACCTGTTT
1316 AATCAGATAGATGCGCCCGCGCGGCCCGTCCAGTTCATCGCCCAGAAGCTGATAGGTTGGGC
1317 AGGTTGCGGTACAAAATCCGCAGTGAACACAGGCGCGCAGGATGCTGTCGGCTTCCAGCGCG
1318 CGCGCGTTCTGCCGCATCTCTTCAGTTAATTGGGTTTGCATAGCCTGCTCCTCAAAGTTCCG
1319 CGTACATGCGACCGGGGTTAAACACGCCGCAAGGGTCGAGCTGCTGTTTAAGCTGCTGGTGA
1320 TAGCGGAATAAAGGAGCCGATAGCGGGGCAAAGCCACCATCTCCGGCACTAAAGCGGGTCGC

ATGACCGCCAGCGTTGCGGGCGATGCGATGGATTTGATTGTCCTCGGCTGTCGATTTCAGCC
AGCGTAACGCCCCGCCCCAGTCGATCAGTTGCTCGCCGGGTAAATCCATCATCGGCGCATCA
CTGGGTAATGAAATGCGCCATAAGGTACCTGGTAACGAGAAGAACGGCAGTTGTTGTTCACG
CAATTGCTGCCAGAACTGACCGGCAACCTCTTCGCCACCCAGCAGTTCACGCGCTGCTTTTA
CCGATCCTTCGCCGCCCTCAAGGCGGATCCACAACGCATTGTCGAAGTAACATAAGCCACTA
ATGGGTAATGGCTGGAGTTGCCACTCGGCGATTTCACTCATGGCTTCTTGCAGGCTGATTTC
CCGACGCAGGCTCAGGGAGGCGCGCGGTCGCGGTAACACTTTCATTGAGATTTCAGTGAGCA
CGCCAAGACAACCGTAGCTTCCGACCATTAACCGTGAGAGATCGTATCCGGCAACGTTTTTC
ATCACTTCGCCACCAAAACGCAGATGTTTTCCAGCGCCGGTAATGATGCGCGTGCCGAGGAC
AAAATCGCGGACCGAACCGCTCCACGGGCGACGCGGCCCCGCCAGCCCGCAGGCGACCATCC
CGCCCCAGGTGGCTTCTTCACCATAATGCGGCGGCTCACAGGGGAGCATTTGCCCCGCGCTT
TCCAGCGCCGCTTCAATTGTCACCAGCGGCGTTCCGACACGCGCGGTTATCACCAGCTCGGT
CGGGTCGTAATTAACAATGCCGCGATGACAACGAACATCCAGCGTTTGCCCGGTGACAGGGC
GACCTAAAAAGGCTTTGCTATTGCTGCCCTGAATCACCAGCGGCGTTTTATCGCTAATCGCC
TGATTCACCTGCTCCAGCAGCGCCTGGCTGTAATCACACTCGCGTAGCATCAGAAACGCTCC
AGTTCAGGGAAAGGTAAATGACCGTGATGCACATGCATGGCACCAAATTCAGCACAGCGGTG
TAGCGTGGGAATGTTTTTCCCAGGGTTCAGCAAACCATCGGGGTCAAACGCCGCCTTGACCG
CATGGAAGGTCGTGATTTCATCGCTGTTGAACTGGGCGCACATTTGATTGATTTTTTCTCGC
CCGATGCCATGTTCGCCACTGATGCTGCCGCCAACTTCAACGCAGAGTTCGAGGATCTTCCC
GCCCAGCTCTTCCGCGCGGGCAAATTCACCGGGTTCGTTGGCATCGAAAAGGATTAACGGGT
GCATGTTGCCATCTCCGGCATGAAAGACGTTGGCAACACGTAAATCATATTGCTGCGATAAA
CGGGCAATGCCTTCCAGTACGCCAGGCAGGGCGCGACGCGGGATGGTGCCATCCATGCAGTA
GTAATCCGGGGAGATACGTCCTACCGCCGGGAACGCATTTTTGCGACCGGCCCAGAAACGTA
CGCGCTCTGCTTCGTCCTGTGCCAGACGGACGTCAGTCGCGCCCGCTTTCAACAAGATGTCG
TTAACCCGCTCGCAGTCTTCCTGTACGTCAGACTCCACGCCGTCCAGCTCGCATAACAAAAT
CGCTTCGGCGTCGACGGGATAACCGGCATGAATAAAATCTTCCGCCGCGCGGATCGACAGGT
TATCCATCATCTCCAGCCCGCCGGGGATAATGCCATTGGCGATGATGTCACCAACCGCAAGT
CCGGCTTTTTCTACCGAGTCAAAGCTGGCTAACAGAACCCGCGCCACGGGCGGCTTCGGCAG
CAGTTTTACCGTCACTTCGGTGGTCACGCCGAGCATACCTTCCGATCCGGTGAACAGCGCCA
GCAGGTCAAAACCAGGTGAATCCAGCGCGTCCGATCCAAGCGTCAGTGCCTCGCCGTCCAGC
GTTTGCACTTCAATTTTCAGCAGGTTATGTACGGTCAGACCATATTTCAGGCAGTGGACGCC
GCCGGCATTTTCAGCCACATTGCCGCCAATGGAACAGGCGATTTGTGAGGAAGGGTCCGGTG
CGTAGTAGAGATTATGCGGTGCAACGGCCTGGGAGATCGCCAGGTTACGCACGCCTGGCTGC
ACGCGCGCGCGGCGACCAACGGGGTTAATGTCGAGGATCTCTTTAAAGCGCGCCATCACCAA
CAACACACCTTTTTCCAGCGGCAGCGCGCCACCAGAAAGCCCGGTGCCTGCACCACGGGTCA
CCACCGGTACACGCAGGCGATGGCAGACAGCCAGAATCGCTGTCACCTGTTCCATTTGCTTA
GGCAGAACAACCAGTAATGGACGCGTGCGATACGCGCTCAACCCGTCACACTCGTAAGGAAT
GATCTCCTCATCGGTATGCAGGATCTCAAGTCCAGGGACATGCTCACGCAGTGCCATCAGTA
CCGATGTGCGGTCGACATCGGGTAAAGCGCCATCAAGACGCTCTTCGTACAAGATGCTCATG
AGTAGGCTTCGCTTTGTTGTGTTGTGTGGCAGCTGATTTTTGCGCGCTGCTTCTGTGAACAG
TTATTAAGCGGGCTTTTCGTTTTCGTCTATCTCTTTAGCTACCGGTCAGACCATTTTTTTTC
CAGCTCTGTGACCTTGTCTTGGTTAACTCAATGTTAAATTGATGTAACATAATCACTTACGT
GATGTGCGTGTTTTGCGAGTTAAGAACAGAAAAATTGGTCCTACCTGTGCACGAGGTCCGGG
ATCTGACGCTCAGTGGAACGAAAACTCACGTTAAGGGATTTTGGTCATGAACAATAAAACTG
TCTGCTTACATAAACAGTAATACAAGGGGTGTTATGAGCCATATTCAACGGG

**Supplementary Data 8: Recombination plasmid, pPBG01**

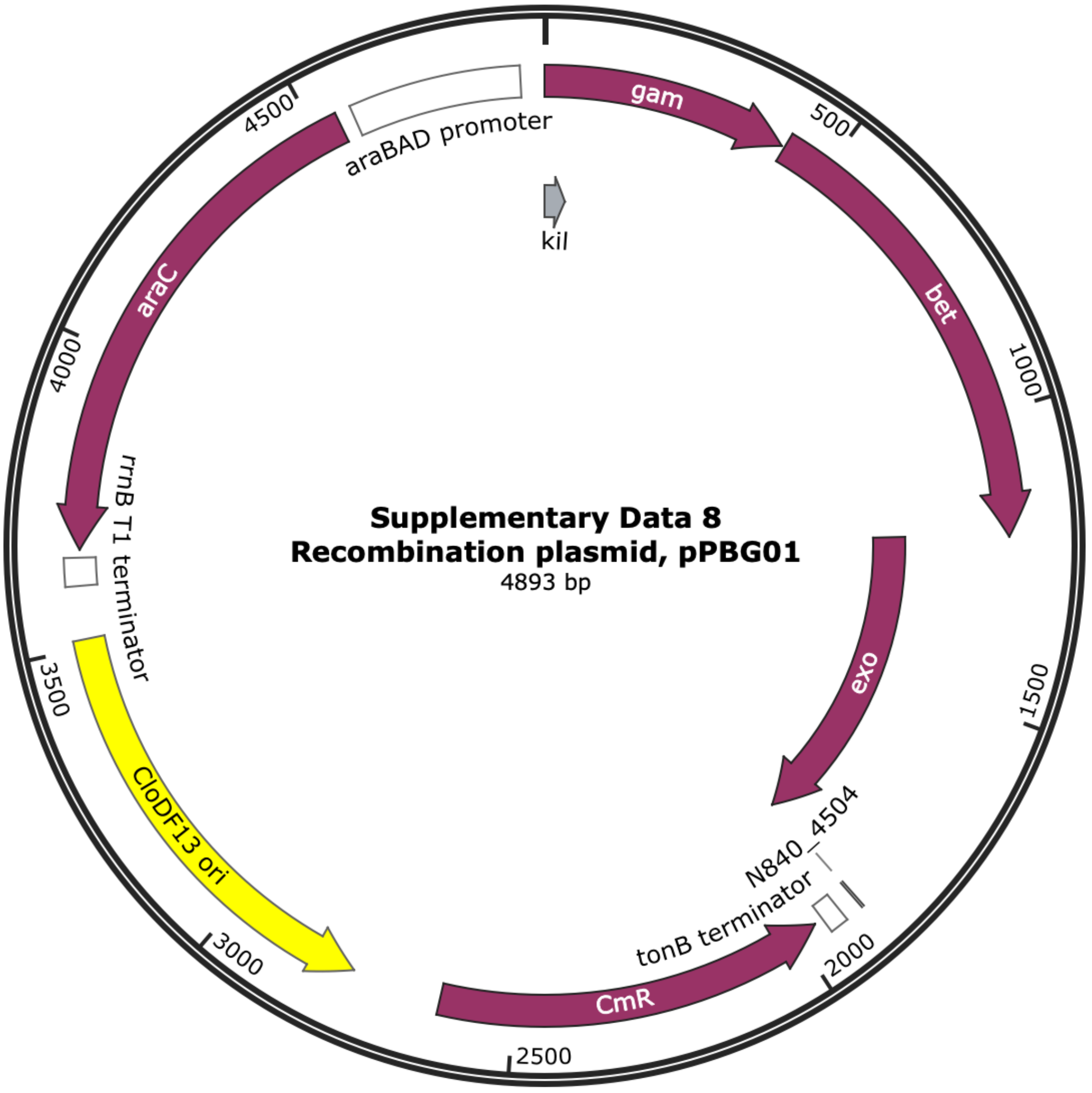


ATGGATATTAATACTGAAACTGAGATCAAGCAAAAGCATTCACTAACCCCCTTTCCTGTTTT
CCTAATCAGCCCGGCATTTCGCGGGCGATATTTTCACAGCTATTTCAGGAGTTCAGCCATGA
ACGCTTATTACATTCAGGATCGTCTTGAGGCTCAGAGCTGGGCGCGTCACTACCAGCAGCTC
GCCCGTGAAGAGAAAGAGGCAGAACTGGCAGACGACATGGAAAAAGGCCTGCCCCAGCACCT
GTTTGAATCGCTATGCATCGATCATTTGCAACGCCACGGGGCCAGCAAAAAATCCATTACCC
GTGCGTTTGATGACGATGTTGAGTTTCAGGAGCGCATGGCAGAACACATCCGGTACATGGTT
GAAACCATTGCTCACCACCAGGTTGATATTGATTCAGAGGTATAAAACGAATGAGTACTGCA
CTCGCAACGCTGGCTGGGAAGCTGGCTGAACGTGTCGGCATGGATTCTGTCGACCCACAGGA
ACTGATCACCACTCTTCGCCAGACGGCATTTAAAGGTGATGCCAGCGATGCGCAGTTCATCG
CATTACTGATCGTTGCCAACCAGTACGGCCTTAATCCGTGGACGAAAGAAATTTACGCCTTT
CCTGATAAGCAGAATGGCATCGTTCCGGTGGTGGGCGTTGATGGCTGGTCCCGCATCATCAA
TGAAAACCAGCAGTTTGATGGCATGGACTTTGAGCAGGACAATGAATCCTGTACATGCCGGA
TTTACCGCAAGGACCGTAATCATCCGATCTGCGTTACCGAATGGATGGATGAATGCCGCCGC
GAACCATTCAAAACTCGCGAAGGCAGAGAAATCACGGGGCCGTGGCAGTCGCATCCCAAACG

19 GATGTTACGTCATAAAGCCATGATTCAGTGTGCCCGTCTGGCCTTCGGATTTGCTGGTATCT
20 ATGACAAGGATGAAGCCGAGCGCATTGTCGAAAATACTGCATACACTGCAGAACGTCAGCCG
21 GAACGCGACATCACTCCGGTTAACGATGAAACCATGCAGGAGATTAACACTCTGCTGATCGC
22 CCTGGATAAAACATGGGATGACGACTTATTGCCGCTCTGTTCCCAGATATTTCGCCGCGACA
23 TTCGTGCATCGTCAGAACTGACACAGGCCGAAGCAGTAAAAGCTCTTGGATTCCTGAAACAG
24 AAAGCCGCAGAGCAGAAGGTGGCAGCATGACACCGGACATTATCCTGCAGCGTACCGGGATC
25 GATGTGAGAGCTGTCGAACAGGGGGAAGATGCGTGGCACAAATTACGGCTCGGCGTCATCAC
26 CGCTTCAGAAGTTCACAACGTGATAGCAAAACCCCGCTCCGGAAAGAAGTGGCCTGACATGA
27 AAATGTCCTACTTCCACACCCTGCTTGCTGAGGTTTGCACCGGTGTGGCTCCGGAAGTTAAC
28 GCTAAAGCACTGGCCTGGGGAAAACAGTACGAGAACGACGCCAGAACCCTGTTTGAATTTAC
29 TTCCGGCGTGAATGTTACTGAATCCCCGATCATCTATCGCGACGAAAGTATGCGTACCGCCT
30 GCTCTCCCGATGGTTTATGCAGTGACGGCAACGGCCTTGAACTGAAATGCCCGTTTACCTCC
31 CGGGATTTCATGAAGTTCCGGCTCGGTGGTTTCGAGGCCATAAAGTCAGCTTACATGGCCCA
32 GGTGCAGTACAGCATGTGGGTGACGCGAAAAAATGCCTGGTACTTTGCCAACTATGACCCGC
33 GTATGAAGCGTGAAGGCCTGCATTATGTCGTGATTGAGCGGGATGAAAAGTACATGGCGAGT
34 TTTGACGAGATCGTGCCGGAGTTCATCGAAAAAATGGACGAGGCACTGGCTGAAATTGGTTT
35 TGTATTTGGGGAGCAATGGCGATGACTAGTACTTAATTAACGGCACTCCTCAGCCAAGTCAA
36 AAGCCTCCGGTCGGAGGCTTTTGACTACATGCCCATGGCGTTTACGCCCCGCCCTGCCACTC
37 ATCGCAGTACTGTTGTAATTCATTAAGCATTCTGCCGACATGGAAGCCATCACAAACGGCAT
38 GATGAACCTGAATCGCCAGCGGCATCAGCACCTTGTCGCCTTGCGTATAATATTTGCCCATA
39 GTGAAAACGGGGGCGAAGAAGTTGTCCATATTGGCCACGTTTAAATCAAAACTGGTGAAACT
40 CACCCAGGGATTGGCTGAGACGAAAAACATATTCTCAATAAACCCTTTAGGGAAATAGGCCA
41 GGTTTTCACCGTAACACGCCACATCTTGCGAATATATGTGTAGAAACTGCCGGAAATCGTCG
42 TGGTATTCACTCCAGAGCGATGAAAACGTTTCAGTTTGCTCATGGAAAACGGTGTAACAAGG
43 GTGAACACTATCCCATATCACCAGCTCACCGTCTTTCATTGCCATACGGAACTCCGGATGAG
44 CATTCATCAGGCGGGCAAGAATGTGAATAAAGGCCGGATAAAACTTGTGCTTATTTTTCTTT
45 ACGGTCTTTAAAAAGGCCGTAATATCCAGCTGAACGGTCTGGTTATAGGTACATTGAGTAAC
46 TGACTGAAATGCCTCAAAATGTTCTTTACGATGCCATTGGGATATATCAACGGTGGTATATC
47 CAGTGATTTTTTTCTCCATTTTAGCTTCCTTAGCTCCTGAAAATCTCGATAACTCAAAAAAT
48 ACGCCCGGTAGTGATCTTATTTCATTATGGTGAAAGTTGGAACCTCTTACGTGCCAAGCCAA
49 ATAGGCCGTCACTCGGTCGCTACGCTCCGGGCGTGAGACTGCGGCGGGCGCTGCGGACACAT
50 ACAAAGTTACCCACAGATTCCGTGGATAAGCAGGGGACTAACATGTGAGGCAAAACAGCAGG
51 GCCGCGCCGGTGGCGTTTTTCCATAGGCTCCGCCCTCCTGCCAGAGTTCACATAAACAGACG
52 CTTTTCCGGTGCATCTGTGGGAGCCGTGAGGCTCAACCATGAATCTGACAGTACGGGCGAAA
53 CCCGACAGGACTTAAAGATCCCCACCGTTTCCGGCGGGTCGCTCCCTCTTGCGCTCTCCTGT
54 TCCGACCCTGCCGTTTACCGGATACCTGTTCCGCCTTTCTCCCTTACGGGAAGTGTGGCGCT
55 TTCTCATAGCTCACACACTGGTATCTCGGCTCGGTGTAGGTCGTTCGCTCCAAGCTGGGCTG
56 TAAGCAAGAACTCCCCGTTCAGCCCGACTGCTGCGCCTTATCCGGTAACTGTTCACTTGAGT
57 CCAACCCGGAAAAGCACGGTAAAACGCCACTGGCAGCAGCCATTGGTAACTGGGAGTTCGCA
58 GAGGATTTGTTTAGCTAAACACGCGGTTGCTCTTGAAGTGTGCGCCAAAGTCCGGCTACACT
59 GGAAGGACAGATTTGGTTGCTGTGCTCTGCGAAAGCCAGTTACCACGGTTAAGCAGTTCCCC
60 AACTGACTTAACCTTCGATCAAACCACCTCCCCAGGTGGTTTTTTCGTTTACAGGGCAAAAG
61 ATTACGCGCAGAAAAAAAGGATCTCAAGAAGATCCTTTGATCTTTTCTACTGAACCGCTCTA
62 GATTTCAGTGCAATTTATCTCTTCAAATGTAGCACCTGAAGTCAGCCCAGGAGGAAGAGGAC
63 ATCCGGTCAAATAAAACGAAAGGCTCAGTCGAAAGACTGGGCCTTTCGTTTTAGACTTAGGG
64 ACCCTTTATGACAACTTGACGGCTACATCATTCACTTTTTCTTCACAACCGGCACGGAACTC
65 GCTCGGGCTGGCCCCGGTGCATTTTTTAAATACCCGCGAGAAATAGAGTTGATCGTCAAAAC
66 CAACATTGCGACCGACGGTGGCGATAGGCATCCGGGTGGTGCTCAAAAGCAGCTTCGCCTGG
67 CTGATACGTTGGTCCTCGCGCCAGCTTAAGACGCTAATCCCTAACTGCTGGCGGAAAAGATG
68 TGACAGACGCGACGGCGACAAGCAAACATGCTGTGCGACGCTGGCGATATCAAAATTGCTGT
69 CTGCCAGGTGATCGCTGATGTACTGACAAGCCTCGCGTACCCGATTATCCATCGGTGGATGG

70 AGCGACTCGTTAATCGCTTCCATGCGCCGCAGTAACAATTGCTCAAGCAGATTTATCGCCAG
71 CAGCTCCGAATAGCGCCCTTCCCCTTGCCCGGCGTTAATGATTTGCCCAAACAGGTCGCTGA
72 AATGCGGCTGGTGCGCTTCATCCGGGCGAAAGAACCCCGTATTGGCAAATATTGACGGCCAG
73 TTAAGCCATTCATGCCAGTAGGCGCGCGGACGAAAGTAAACCCACTGGTGATACCATTCGCG
74 AGCCTCCGGATGACGACCGTAGTGATGAATCTCTCCTGGCGGGAACAGCAAAATATCACCCG
75 GTCGGCAAACAAATTCTCGTCCCTGATTTTTCACCACCCCCTGACCGCGAATGGTGAGATTG
76 AGAATATAACCTTTCATTCCCAGCGGTCGGTCGATAAAAAAATCGAGATAACCGTTGGCCTC
77 AATCGGCGTTAAACCCGCCACCAGATGGGCATTAAACGAGTATCCCGGCAGCAGGGGATCAT
78 TTTGCGCTTCAGCCATACTTTTCATACTCCCACCATTCAGAGAAGAAACCAATTGTCCATAT
79 TGCATCAGACATTGCCGTCACTGCGTCTTTTACTGGCTCTTCTCGCTAACCCAACCGGTAAC
80 CCCGCTTATTAAAAGCATTCTGTAACAAAGCGGGACCAAAGCCATGACAAAAACGCGTAACA
81 AAAGTGTCTATAATCACGGCAGAAAAGTCCACATTGATTATTTGCACGGCGTCACACTTTGC
82 TATGCCATAGCATTTTTATCCATAAGATTAGCGGATCCTACCTGACGCTTTTTATCGCAACT
83 CTCTACTGTTTCTCCATACCCGTTTTTTTGGACGCGTACAACTCAAGTCTGACATAA

### Strains created in this study (LySE)

| Strain | Genotype or relevant features |
|---|---|
| *E. coli* MG1655 *rpsL* K43R | F- lambda- ilvG- rfb-50 rph-1 rpsLK43R |
| *E. coli* MG1655 Δ*trxA* | Scarless deletion of *trxA* |
| *E. coli* MG1655 Δ*lacIZYA* | Scarless deletion of lacIZYA |
| Phage T7 ΔDNAP *trxA* | Replacement of gp5 (T7 DNAP) with MG1655 *trxA* |

| Function |
| --- |
| Host strain for LySE |
| Selection strain for TXTL phage engineering |
| Host strain for lacZ inactivation assay |
| Biocontained T7 |

<u>Supplementary Table 2a. All T7 DNAP variants tested.</u> The functionality of each engineered T7DNAP variant was tested by complementation assay with T7ΔDNAP. Efficient cell lysis shown by reduction in cell density indicates functional T7DNAP (green curves), while lack of lysis shown by increase in cell density indicates non-functional T7DNAP (red curves). Displayed are representative curves of at least three independent replicates.

| T7 DNAP | Variant type | Functional ? | Lysis curve |
|---|---|---|---|
| wt | Wildtype | Yes | 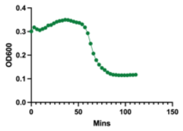 |
| v1 | Y64C/F120L/S399T | Yes | 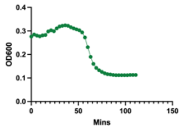 |
| v2.1 | L479N/H506Y/T523R/P560H | No | 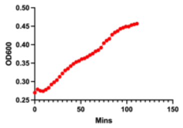 |
| v2.2 | Y64C/F120L/S399T/L479N | Yes | 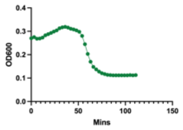 |
| v2.3 | Y64C/F120L/S399T/H506Y | No | 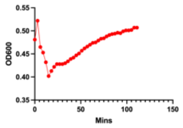 |
| v2.4 | Y64C/F120L/S399T/T523R | Yes | 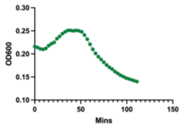 |
| v2.5 | Y64C/F120L/S399T/P560H | Yes | 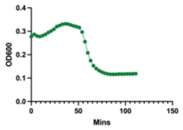 |
| v2.6 | Y64C/F120L/S399T/L479N/H506Y | No | 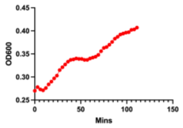 |
| v2.7 | Y64C/F120L/S399T/L479N/T523R | Yes | 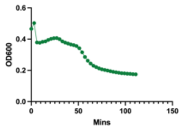 |
| v2.8 | Y64C/F120L/S399T/L479N/P560H | Yes | 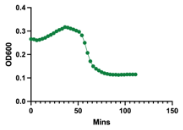 |
| v2.9 | Y64C/F120L/S399T/H506Y/T523R | Yes | 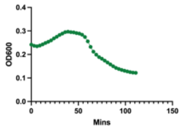 |

| | | | |
|---|---|---|---|
| v2.10 | Y64C/F120L/S399T/H506Y/P560H | No | 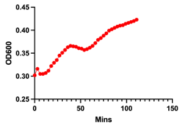 |
| v2.11 | Y64C/F120L/S399T/T523R/P560H | Yes | 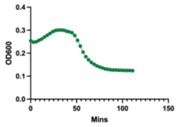 |
| v2.12 | Y64C/F120L/S399T/H506Y/T523R/P560H | No | 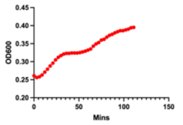 |
| v2.13 | Y64C/F120L/S399T/L479N/T523R/P560H | Yes | 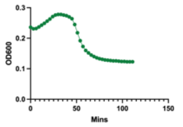 |
| v2.14 | Y64C/F120L/S399T/L479N/H506Y/P560H | No | 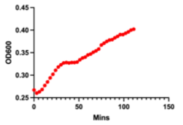 |
| v2.15 | Y64C/F120L/S399T/L479N/H506Y/T523R | No | 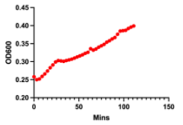 |
| v2.16 | Y64C/F120L/S399T/L479N/H506Y/T523R/P560H | No | 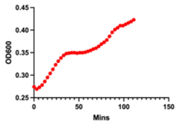 |
| v3 | D5A/E7A/Y64C/F120L/S399T | Yes | 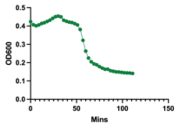 |
| v4 | D5A/E7A/Y64C/F120L/S399T/T523R | Yes | 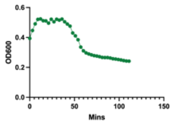 |
| v5.1 | TadA-8e-15aa linker-T7DNAP + UGI | No | 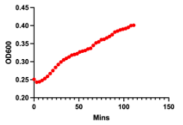 |
| v5.2 | TadA-8e-21aa linker-T7DNAP + UGI | No | 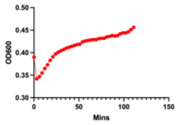 |
| v5.3 | TadA-8e-24aa linker-T7DNAP + UGI | Yes | 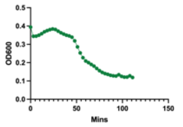 |
| v5.4 | TadA-8e-29aa linker-T7DNAP + UGI | No | 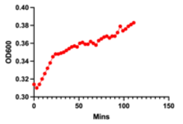 |
| v5.5 | TadA-8e-33aa linker-T7DNAP + UGI | Yes | 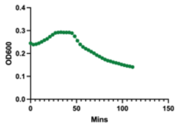 |

| | | | |
|---|---|---|---|
| v6.1 | PmCDA1-24aa linker-T7DNAP + UGI | Yes | 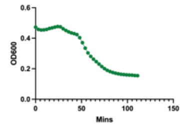 |
| v6.2 | PmCDA1-33aa linker-T7DNAP + UGI | No | 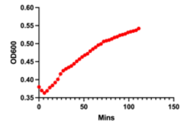 |
| v7 | TadA-8e-24aa linker-T7DNAP(Y64C/F120L/S399T/T523R) + UGI | Yes | 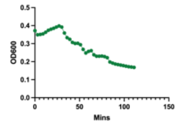 |
| v8 | TadA-8e-24aa linker-T7DNAP(D5A/E7A/Y64C/F120L/S399T/T523R) + UGI | Yes | 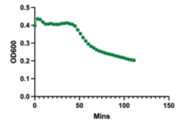 |
| v9 | TadDE-24aa linker-T7DNAP(D5A/E7A/Y64C/F120L/S399T/T523R) + UGI | Yes | 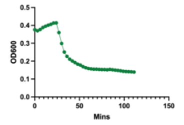 |

Supplementary Table 2b. Protein sequence of T7 DNAP variants

Deaminase; Linker

| T7 DNAP | Protein sequence |
|---|---|
| wt | MIVSDIEANALLESVTKFHCGVIYDYSTAEYVSYRPSDFGAYLDALEAEVAR<br>GGLIVFHNGHKYDVPALTKLAKLQLNREFHLPRENCIDTLVLSRLIHSNLKD<br>TDMGLLRSGKLPGKRFGSHALEAWGYRLGEMKGEYKDDFKRMLEEQGE<br>EYVDGMEWWNFNEEMMDYNVQDVVVTKALLEKLLSDKHYFPPEIDFTDV<br>GYTTFWSESLEAVDIEHRAAWLLAKQERNGFPFDTKAIEELYVELAARRSE<br>LLRKLTETFGSWYQPKGGTEMFCHPRTGKPLPKYPRIKTPKVGGIFKKPKN<br>KAQREGREPCELDTREYVAGAPYTPVEHVVFNPSSRDHIQKKLQEAGWV<br>PTKYTDKGAPVVDDEVLEGVRVDDPEKQAAIDLIKEYLMIQKRIGQSAEGD<br>KAWLRYVAEDGKIHGSVNPNGAVTGRATHAFPNLAQIPGVRSPYGEQCRA<br>AFGAEHHLDGITGKPWVQAGIDASGLELRCLAHFMARFDNGEYAHEILNG<br>DIHTKNQIAAELPTRDNAKTFIYGFLYGAGDEKIGQIVGAGKERGKELKKKF<br>LENTPAIAALRESIQQTLVESSQWVAGEQQVKWKRRWIKGLDGRKVHVRS<br>PHAALNTLLQSAGALICKLWIIKTEEMLVEKGLKHGWDGDFAYMAWVHDEI<br>QVGCRTEEIAQVVIETAQEAMRWVGDHWNFRCLLDTEGKMGPNWAICH |
| v5.1 | MKSSMSEVEFSHEYWMRHALTLAKRARDEREVPVGAVLVLNNRVIGEGW<br>NRAIGLHDPTAHAEIMALRQGGLVMQNYRLIDATLYVTFEPCVMCAGAMIH<br>SRIGRVVFGVRNSKRGAAGSLMNVLNYPGMNHRVEITEGILADECAALLC<br>DFYRMPRQVFNAQKKAQSSINSGSETPGTSESATPEIVSDIEANALLESVT<br>KFHCGVIYDYSTAEYVSYRPSDFGAYLDALEAEVARGGLIVFHNGHKYDVP<br>ALTKLAKLQLNREFHLPRENCIDTLVLSRLIHSNLKDTDMGLLRSGKLPGKR<br>FGSHALEAWGYRLGEMKGEYKDDFKRMLEEQGEEYVDGMEWWNFNEE<br>MMDYNVQDVVVTKALLEKLLSDKHYFPPEIDFTDVGYTTFWSESLEAVDIE<br>HRAAWLLAKQERNGFPFDTKAIEELYVELAARRSELLRKLTETFGSWYQPK<br>GGTEMFCHPRTGKPLPKYPRIKTPKVGGIFKKPKNKAQREGREPCELDTR<br>EYVAGAPYTPVEHVVFNPSSRDHIQKKLQEAGWVPTKYTDKGAPVVDDE<br>VLEGVRVDDPEKQAAIDLIKEYLMIQKRIGQSAEGDKAWLRYVAEDGKIHG<br>SVNPNGAVTGRATHAFPNLAQIPGVRSPYGEQCRAAFGAEHHLDGITGKP<br>WVQAGIDASGLELRCLAHFMARFDNGEYAHEILNGDIHTKNQIAAELPTRD<br>NAKTFIYGFLYGAGDEKIGQIVGAGKERGKELKKKFLENTPAIAALRESIQQ<br>TLVESSQWVAGEQQVKWKRRWIKGLDGRKVHVRSPHAALNTLLQSAGALI<br>CKLWIIKTEEMLVEKGLKHGWDGDFAYMAWVHDEIQVGCRTEEIAQVVIET<br>AQEAMRWVGDHWNFRCLLDTEGKMGPNWAICH |
| v5.2 | MKSSMSEVEFSHEYWMRHALTLAKRARDEREVPVGAVLVLNNRVIGEGW<br>NRAIGLHDPTAHAEIMALRQGGLVMQNYRLIDATLYVTFEPCVMCAGAMIH<br>SRIGRVVFGVRNSKRGAAGSLMNVLNYPGMNHRVEITEGILADECAALLC<br>DFYRMPRQVFNAQKKAQSSINSGGGSETPGTSESATPESGGSIVSDIEAN<br>ALLESVTKFHCGVIYDYSTAEYVSYRPSDFGAYLDALEAEVARGGLIVFHN<br>GHKYDVPALTKLAKLQLNREFHLPRENCIDTLVLSRLIHSNLKDTDMGLLRS<br>GKLPGKRFGSHALEAWGYRLGEMKGEYKDDFKRMLEEQGEEYVDGME<br>WWNFNEEMMDYNVQDVVVTKALLEKLLSDKHYFPPEIDFTDVGYTTFWS<br>ESLEAVDIEHRAAWLLAKQERNGFPFDTKAIEELYVELAARRSELLRKLTET<br>FGSWYQPKGGTEMFCHPRTGKPLPKYPRIKTPKVGGIFKKPKNKAQREG<br>REPCELDTREYVAGAPYTPVEHVVFNPSSRDHIQKKLQEAGWVPTKYTDK<br>GAPVVDDEVLEGVRVDDPEKQAAIDLIKEYLMIQKRIGQSAEGDKAWLRYV<br>AEDGKIHGSVNPNGAVTGRATHAFPNLAQIPGVRSPYGEQCRAAFGAEHH<br>LDGITGKPWVQAGIDASGLELRCLAHFMARFDNGEYAHEILNGDIHTKNQI<br>AAELPTRDNAKTFIYGFLYGAGDEKIGQIVGAGKERGKELKKKFLENTPAIA<br>ALRESIQQTLVESSQWVAGEQQVKWKRRWIKGLDGRKVHVRSPHAALNT<br>LLQSAGALICKLWIIKTEEMLVEKGLKHGWDGDFAYMAWVHDEIQVGCRTE<br>EIAQVVIETAQEAMRWVGDHWNFRCLLDTEGKMGPNWAICH |

| | |
|---|---|
| v5.3 | MKSSMSEVEFSHEYWMRHALTLAKRARDEREVPVGAVLVLNNRVIGEGW<br>NRAIGLHDPTAHAEIMALRQGGLVMQNYRLIDATLYVTFEPCVMCAGAMIH<br>SRIGRVVFGVRNSKRGAAGSLMNVLNYPGMNHRVEITEGILADECAALLC<br>DFYRMPRQVFNAQKKAQSSINSGGGSETPGTSESATPESGGSIKGIVSDIE<br>ANALLESVTKFHCGVIYDYSTAEYVSYRPSDFGAYLDALEAEVARGGLIVFH<br>NGHKYDVPALTKLAKLQLNREFHLPRENCIDTLVLSRLIHSNLKDTDMGLLR<br>SGKLPGKRFGSHALEAWGYRLGEMKGEYKDDFKRMLEEQGEEYVDGME<br>WWNFNEEMMDYNVQDVVVTKALLEKLLSDKHYFPPEIDFTDVGYTTFWS<br>ESLEAVDIEHRAAWLLAKQERNGFPFDTKAIEELYVELAARRSELLRKLTET<br>FGSWYQPKGGTEMFCHPRTGKPLPKYPRIKTPKVGGIFKKPKNKAQREG<br>REPCELDTREYVAGAPYTPVEHVVFNPSSRDHIQKKLQEAGWVPTKYTDK<br>GAPVVDDEVLEGVRVDDPEKQAAIDLIKEYLMIQKRIGQSAEGDKAWLRYV<br>AEDGKIHGSVNPNGAVTGRATHAFPNLAQIPGVRSPYGEQCRAAFGAEHH<br>LDGITGKPWVQAGIDASGLELRCLAHFMARFDNGEYAHEILNGDIHTKNQI<br>AAELPTRDNAKTFIYGFLYGAGDEKIGQIVGAGKERGKELKKKFLENTPAIA<br>ALRESIQQTLVESSQWVAGEQQVKWKRRWIKGLDGRKVHVRSPHAALNT<br>LLQSAGALICKLWIIKTEEMLVEKGLKHGWDGDFAYMAWVHDEIQVGCRTE<br>EIAQVVIETAQEAMRWVGDHWNFRCLLDTEGKMGPNWAICH |
| v5.4 | MKSSMSEVEFSHEYWMRHALTLAKRARDEREVPVGAVLVLNNRVIGEGW<br>NRAIGLHDPTAHAEIMALRQGGLVMQNYRLIDATLYVTFEPCVMCAGAMIH<br>SRIGRVVFGVRNSKRGAAGSLMNVLNYPGMNHRVEITEGILADECAALLC<br>DFYRMPRQVFNAQKKAQSSINSGGSSGGSSGGGSETPGTSESATPESG<br>GSIVSDIEANALLESVTKFHCGVIYDYSTAEYVSYRPSDFGAYLDALEAEVA<br>RGGLIVFHNGHKYDVPALTKLAKLQLNREFHLPRENCIDTLVLSRLIHSNLK<br>DTDMGLLRSGKLPGKRFGSHALEAWGYRLGEMKGEYKDDFKRMLEEQG<br>EEYVDGMEWWNFNEEMMDYNVQDVVVTKALLEKLLSDKHYFPPEIDFTD<br>VGYTTFWSESLEAVDIEHRAAWLLAKQERNGFPFDTKAIEELYVELAARRS<br>ELLRKLTETFGSWYQPKGGTEMFCHPRTGKPLPKYPRIKTPKVGGIFKKPK<br>NKAQREGREPCELDTREYVAGAPYTPVEHVVFNPSSRDHIQKKLQEAGW<br>VPTKYTDKGAPVVDDEVLEGVRVDDPEKQAAIDLIKEYLMIQKRIGQSAEG<br>DKAWLRYVAEDGKIHGSVNPNGAVTGRATHAFPNLAQIPGVRSPYGEQCR<br>AAFGAEHHLDGITGKPWVQAGIDASGLELRCLAHFMARFDNGEYAHEILN<br>GDIHTKNQIAAELPTRDNAKTFIYGFLYGAGDEKIGQIVGAGKERGKELKKK<br>FLENTPAIAALRESIQQTLVESSQWVAGEQQVKWKRRWIKGLDGRKVHVR<br>SPHAALNTLLQSAGALICKLWIIKTEEMLVEKGLKHGWDGDFAYMAWVHDE<br>IQVGCRTEEIAQVVIETAQEAMRWVGDHWNFRCLLDTEGKMGPNWAICH |
| v5.5 | MKSSMSEVEFSHEYWMRHALTLAKRARDEREVPVGAVLVLNNRVIGEGW<br>NRAIGLHDPTAHAEIMALRQGGLVMQNYRLIDATLYVTFEPCVMCAGAMIH<br>SRIGRVVFGVRNSKRGAAGSLMNVLNYPGMNHRVEITEGILADECAALLC<br>DFYRMPRQVFNAQKKAQSSINSGSETPGTSESATPESGGSDYKDDDDKG<br>SLIKGIVSDIEANALLESVTKFHCGVIYDYSTAEYVSYRPSDFGAYLDALEAE<br>VARGGLIVFHNGHKYDVPALTKLAKLQLNREFHLPRENCIDTLVLSRLIHSN<br>LKDTDMGLLRSGKLPGKRFGSHALEAWGYRLGEMKGEYKDDFKRMLEE<br>QGEEYVDGMEWWNFNEEMMDYNVQDVVVTKALLEKLLSDKHYFPPEIDF<br>TDVGYTTFWSESLEAVDIEHRAAWLLAKQERNGFPFDTKAIEELYVELAAR<br>RSELLRKLTETFGSWYQPKGGTEMFCHPRTGKPLPKYPRIKTPKVGGIFKK<br>PKNKAQREGREPCELDTREYVAGAPYTPVEHVVFNPSSRDHIQKKLQEAG<br>WVPTKYTDKGAPVVDDEVLEGVRVDDPEKQAAIDLIKEYLMIQKRIGQSAE<br>GDKAWLRYVAEDGKIHGSVNPNGAVTGRATHAFPNLAQIPGVRSPYGEQC<br>RAAFGAEHHLDGITGKPWVQAGIDASGLELRCLAHFMARFDNGEYAHEIL<br>NGDIHTKNQIAAELPTRDNAKTFIYGFLYGAGDEKIGQIVGAGKERGKELKK<br>KFLENTPAIAALRESIQQTLVESSQWVAGEQQVKWKRRWIKGLDGRKVHV<br>RSPHAALNTLLQSAGALICKLWIIKTEEMLVEKGLKHGWDGDFAYMAWVHD<br>EIQVGCRTEEIAQVVIETAQEAMRWVGDHWNFRCLLDTEGKMGPNWAICH |

| | |
|---|---|
| v6.1 | MKSSMMTDAEYVRIHEKLDIYTFKKQFFNNKKSVSHRCYVLFELKRRGER<br>RACFWGYAVNKPQSGTERGIHAEIFSIRKVEEYLRDNPGQFTINWYSSWS<br>PCADCAEKILEWYNQELRGNGHTLKIWACKLYYEKNARNQIGLWNLRDNG<br>VGLNVMVSEHYQCCRKIFIQSSHNQLNENRWLEKTLKRAEKWRSELSIMI<br>QVKILHTTKSPAVSSGGGSETPGTSESATPESGGSIKGIVSDIEANALLESV<br>TKFHCGVIYDYSTAEYVSYRPSDFGAYLDALEAEVARGGLIVFHNGHKYDV<br>PALTKLAKLQLNREFHLPRENCIDTLVLSRLIHSNLKDTDMGLLRSGKLPGK<br>RFGSHALEAWGYRLGEMKGEYKDDFKRMLEEQGEEYVDGMEWWNFNE<br>EMMDYNVQDVVVTKALLEKLLSDKHYFPPEIDFTDVGYTTFWSESLEAVDI<br>EHRAAWLLAKQERNGFPFDTKAIEELYVELAARRSELLRKLTETFGSWYQP<br>KGGTEMFCHPRTGKPLPKYPRIKTPKVGGIFKKPKNKAQREGREPCELDT<br>REYVAGAPYTPVEHVVFNPSSRDHIQKKLQEAGWVPTKYTDKGAPVVDD<br>EVLEGVRVDDPEKQAAIDLIKEYLMIQKRIGQSAEGDKAWLRYVAEDGKIH<br>GSVNPNGAVTGRATHAFPNLAQIPGVRSPYGEQCRAAFGAEHHLDGITGK<br>PWVQAGIDASGLELRCLAHFMARFDNGEYAHEILNGDIHTKNQIAAELPTR<br>DNAKTFIYGFLYGAGDEKIGQIVGAGKERGKELKKKFLENTPAIAALRESIQ<br>QTLVESSQWVAGEQQVKWKRRWIKGLDGRKVHVRSPHAALNTLLQSAGA<br>LICKLWIIKTEEMLVEKGLKHGWDGDFAYMAWVHDEIQVGCRTEEIAQVVIE<br>TAQEAMRWVGDHWNFRCLLDTEGKMGPNWAICH |
| v6.2 | MKSSMMTDAEYVRIHEKLDIYTFKKQFFNNKKSVSHRCYVLFELKRRGER<br>RACFWGYAVNKPQSGTERGIHAEIFSIRKVEEYLRDNPGQFTINWYSSWS<br>PCADCAEKILEWYNQELRGNGHTLKIWACKLYYEKNARNQIGLWNLRDNG<br>VGLNVMVSEHYQCCRKIFIQSSHNQLNENRWLEKTLKRAEKWRSELSIMI<br>QVKILHTTKSPAVSSGSETPGTSESATPESGGSDYKDDDDKGSLIKGIVSDI<br>EANALLESVTKFHCGVIYDYSTAEYVSYRPSDFGAYLDALEAEVARGGLIVF<br>HNGHKYDVPALTKLAKLQLNREFHLPRENCIDTLVLSRLIHSNLKDTDMGLL<br>RSGKLPGKRFGSHALEAWGYRLGEMKGEYKDDFKRMLEEQGEEYVDGM<br>EWWNFNEEMMDYNVQDVVVTKALLEKLLSDKHYFPPEIDFTDVGYTTFW<br>SESLEAVDIEHRAAWLLAKQERNGFPFDTKAIEELYVELAARRSELLRKLTE<br>TFGSWYQPKGGTEMFCHPRTGKPLPKYPRIKTPKVGGIFKKPKNKAQRE<br>GREPCELDTREYVAGAPYTPVEHVVFNPSSRDHIQKKLQEAGWVPTKYTD<br>KGAPVVDDEVLEGVRVDDPEKQAAIDLIKEYLMIQKRIGQSAEGDKAWLRY<br>VAEDGKIHGSVNPNGAVTGRATHAFPNLAQIPGVRSPYGEQCRAAFGAEH<br>HLDGITGKPWVQAGIDASGLELRCLAHFMARFDNGEYAHEILNGDIHTKNQ<br>IAAELPTRDNAKTFIYGFLYGAGDEKIGQIVGAGKERGKELKKKFLENTPAIA<br>ALRESIQQTLVESSQWVAGEQQVKWKRRWIKGLDGRKVHVRSPHAALNT<br>LLQSAGALICKLWIIKTEEMLVEKGLKHGWDGDFAYMAWVHDEIQVGCRTE<br>EIAQVVIETAQEAMRWVGDHWNFRCLLDTEGKMGPNWAICH |
| v7 | MKSSMSEVEFSHEYWMRHALTLAKRARDEREVPVGAVLVLNNRVIGEGW<br>NRAIGLHDPTAHAEIMALRQGGLVMQNYRLIDATLYVTFEPCVMCAGAMIH<br>SRIGRVVFGVRNSKRGAAGSLMNVLNYPGMNHRVEITEGILADECAALLC<br>DFYRMPRQVFNAQKKAQSSINSGGGSETPGTSESATPESGGSIKGIVSDIE<br>ANALLESVTKFHCGVIYDYSTAEYVSYRPSDFGAYLDALEAEVARGGLIVFH<br>NGHKCDVPALTKLAKLQLNREFHLPRENCIDTLVLSRLIHSNLKDTDMGLLR<br>SGKLPGKRLGSHALEAWGYRLGEMKGEYKDDFKRMLEEQGEEYVDGME<br>WWNFNEEMMDYNVQDVVVTKALLEKLLSDKHYFPPEIDFTDVGYTTFWS<br>ESLEAVDIEHRAAWLLAKQERNGFPFDTKAIEELYVELAARRSELLRKLTET<br>FGSWYQPKGGTEMFCHPRTGKPLPKYPRIKTPKVGGIFKKPKNKAQREG<br>REPCELDTREYVAGAPYTPVEHVVFNPSSRDHIQKKLQEAGWVPTKYTDK<br>GAPVVDDEVLEGVRVDDPEKQAAIDLIKEYLMIQKRIGQTAEGDKAWLRYV<br>AEDGKIHGSVNPNGAVTGRATHAFPNLAQIPGVRSPYGEQCRAAFGAEHH<br>LDGITGKPWVQAGIDASGLELRCLAHFMARFDNGEYAHEILNGDIHTKNQI<br>AAELPTRDNAKRFIYGFLYGAGDEKIGQIVGAGKERGKELKKKFLENTPAIA<br>ALRESIQQTLVESSQWVAGEQQVKWKRRWIKGLDGRKVHVRSPHAALNT |

| | |
|---|---|
| | LLQSAGALICKLWIIKTEEMLVEKGLKHGWDGDFAYMAWVHDEIQVGCRTE<br>EIAQVVIETAQEAMRWVGDHWNFRCLLDTEGKMGPNWAICH |
| v8 | MKSSMSEVEFSHEYWMRHALTLAKRARDEREVPVGAVLVLNNRVIGEGW<br>NRAIGLHDPTAHAEIMALRQGGLVMQNYRLIDATLYVTFEPCVMCAGAMIH<br>SRIGRVVFGVRNSKRGAAGSLMNVLNYPGMNHRVEITEGILADECAALLC<br>DFYRMPRQVFNAQKKAQSSINSGGGSETPGTSESATPESGGSIKGIVSAIA<br>ANALLESVTKFHCGVIYDYSTAEYVSYRPSDFGAYLDALEAEVARGGLIVFH<br>NGHKCDVPALTKLAKLQLNREFHLPRENCIDTLVLSRLIHSNLKDTDMGLLR<br>SGKLPGKRLGSHALEAWGYRLGEMKGEYKDDFKRMLEEQGEEYVDGME<br>WWNFNEEMMDYNVQDVVVTKALLEKLLSDKHYFPPEIDFTDVGYTTFWS<br>ESLEAVDIEHRAAWLLAKQERNGFPFDTKAIEELYVELAARRSELLRKLTET<br>FGSWYQPKGGTEMFCHPRTGKPLPKYPRIKTPKVGGIFKKPKNKAQREG<br>REPCELDTREYVAGAPYTPVEHVVFNPSSRDHIQKKLQEAGWVPTKYTDK<br>GAPVVDDEVLEGVRVDDPEKQAAIDLIKEYLMIQKRIGQTAEGDKAWLRYV<br>AEDGKIHGSVNPNGAVTGRATHAFPNLAQIPGVRSPYGEQCRAAFGAEHH<br>LDGITGKPWVQAGIDASGLELRCLAHFMARFDNGEYAHEILNGDIHTKNQI<br>AAELPTRDNAKRFIYGFLYGAGDEKIGQIVGAGKERGKELKKKFLENTPAIA<br>ALRESIQQTLVESSQWVAGEQQVKWKRRWIKGLDGRKVHVRSPHAALNT<br>LLQSAGALICKLWIIKTEEMLVEKGLKHGWDGDFAYMAWVHDEIQVGCRTE<br>EIAQVVIETAQEAMRWVGDHWNFRCLLDTEGKMGPNWAICH |
| v9 | MKSSMSEVEFSHEYWMRHALTLAKRARDEGEAPVGAVLVLNNRVIGEGW<br>NRRIGLHDPTAHAEIMALRQGGLVMQNSRLIDATLYVTFEPCVMCAGAMIN<br>SRIGRVVFGVRNSKRGAAGSLMNVLNYPGMNVIGLADECAALCDFYRMP<br>RQVFNAQKKAQSSINSGGGSETPGTSESATPESGGSIKGIVSAIAANALLE<br>SVTKFHCGVIYDYSTAEYVSYRPSDFGAYLDALEAEVARGGLIVFHNGHKC<br>DVPALTKLAKLQLNREFHLPRENCIDTLVLSRLIHSNLKDTDMGLLRSGKLP<br>GKRLGSHALEAWGYRLGEMKGEYKDDFKRMLEEQGEEYVDGMEWWNF<br>NEEMMDYNVQDVVVTKALLEKLLSDKHYFPPEIDFTDVGYTTFWSESLEA<br>VDIEHRAAWLLAKQERNGFPFDTKAIEELYVELAARRSELLRKLTETFGSW<br>YQPKGGTEMFCHPRTGKPLPKYPRIKTPKVGGIFKKPKNKAQREGREPCE<br>LDTREYVAGAPYTPVEHVVFNPSSRDHIQKKLQEAGWVPTKYTDKGAPVV<br>DDEVLEGVRVDDPEKQAAIDLIKEYLMIQKRIGQTAEGDKAWLRYVAEDGKI<br>HGSVNPNGAVTGRATHAFPNLAQIPGVRSPYGEQCRAAFGAEHHLDGITG<br>KPWVQAGIDASGLELRCLAHFMARFDNGEYAHEILNGDIHTKNQIAAELPT<br>RDNAKRFIYGFLYGAGDEKIGQIVGAGKERGKELKKKFLENTPAIAALRESI<br>QQTLVESSQWVAGEQQVKWKRRWIKGLDGRKVHVRSPHAALNTLLQSA<br>GALICKLWIIKTEEMLVEKGLKHGWDGDFAYMAWVHDEIQVGCRTEEIAQV<br>VIETAQEAMRWVGDHWNFRCLLDTEGKMGPNWAICH |
